# Mapping variation in the morphological landscape of human cells with optical pooled CRISPRi screening

**DOI:** 10.1101/2022.12.27.522042

**Authors:** Ramon Lorenzo D. Labitigan, Adrian L. Sanborn, Cynthia V. Hao, Caleb K. Chan, Nathan M. Belliveau, Eva M. Brown, Mansi Mehrotra, Julie A. Theriot

**Affiliations:** Department of Biology and Howard Hughes Medical Institute, University of Washington, Seattle, WA 98195, USA; Department of Biochemistry, Stanford University, Stanford, CA 94305, USA; Department of Computer Science, Stanford University, Stanford, CA 94305, USA; Department of Structural Biology, Stanford University, Stanford, CA 94305, USA; Department of Bioengineering, Stanford University, Stanford, CA 94305, USA; Allen Institute for Cell Science, Seattle, WA 98125, USA; Information School, University of Washington, Seattle, WA 98195, USA

## Abstract

The contributions of individual genes to cell-scale morphology and cytoskeletal organization are challenging to define due to the wide intercellular variation of these complex phenotypes. We leveraged the controlled nature of image-based pooled screening to assess the impact of CRISPRi knockdown of 366 genes on cell and nuclear morphology in human U2OS osteosarcoma cells. Screen scale-up was facilitated by a new, efficient barcode readout method that successfully genotyped 85% of cells. Phenotype analysis using a deep learning algorithm, the β-variational autoencoder, produced a feature embedding space distinct from one derived from conventional morphological profiling, but detected similar gene hits while requiring minimal design decisions. We found 45 gene hits and visualized their effect by rationally constrained sampling of cells along the direction of phenotypic shift. By relating these phenotypic shifts to each other, we construct a quantitative and interpretable space of morphological variation in human cells.

## INTRODUCTION

Image-based screening of genetic perturbations is a powerful platform for biological discovery because it can simultaneously identify the molecular players involved in specific biological phenomena and provide high-content spatial phenotypic information for each perturbed population^1,2^. While classic genetic screens typically examined exceptionally stereotyped phenotypes where mutants of interest could be robustly identified by eye, even at the level of individual specimens^3,4^, modern image-based screens can detect increasingly subtle effects within phenotypic spaces of growing complexity, enabled by innovations in screening technology and data analysis^5^.

Conventionally, cellular screens have been executed in an arrayed format, wherein each perturbation is applied to a population of cells in its own separate compartment^6,7^. However, arrayed genetic screens are both labor-intensive and susceptible to technical variation, as well-to-well differences in conditions and handling (such as from seeding, staining, or imaging) can obscure subtle phenotypic effects^8,9^. Pooled screens overcome these limitations by introducing a genetic perturbation library within a single mixed sample, then using sequencing to identify the genetic perturbations present in phenotypic subpopulations generated by physical separation or enrichment. For example, this is commonly done for one-dimensional optical phenotypes, such as total fluorescence from a reporter, using fluorescence-activated cell sorting (FACS) followed by bulk sequencing^10,11^. Recent papers extended this separation principle to microscopy-based phenotypes by photoactivating a fluorophore within a phenotype-defined subset of cells using precise laser targeting^12–14^. However, these classification-dependent screening approaches are not possible when phenotypic filters cannot be specified in advance, such as when the important modes of variation are high-dimensional, continuous, or unknown *a priori*. Furthermore, these bulksequencing-based approaches do not generate genotypephenotype mappings at single-cell resolution.

Only recently has it become feasible to do pooled, image-based screens with single-cell resolution (Fig. 1a). In this approach, associating the visual phenotype with the perturbation genotype is made possible by new, more efficient methods to amplify perturbation-associated DNA barcodes within fixed cells and sequence them on a standard fluorescence microscope^15,16^. The first applications of pooled optical screens employed clearly-defined phenotypes such as nuclear localization of a fluorescently tagged protein^16^ or focused on essential genes whose knockdowns were expected to have drastic effects on known parameters such as cell size, the presence of DNA damage, and mitotic defect prevalence^17^. However, the detection and characterization of more subtle changes within a complex phenotypic space have yet to be fully explored.

**Fig. 1:**
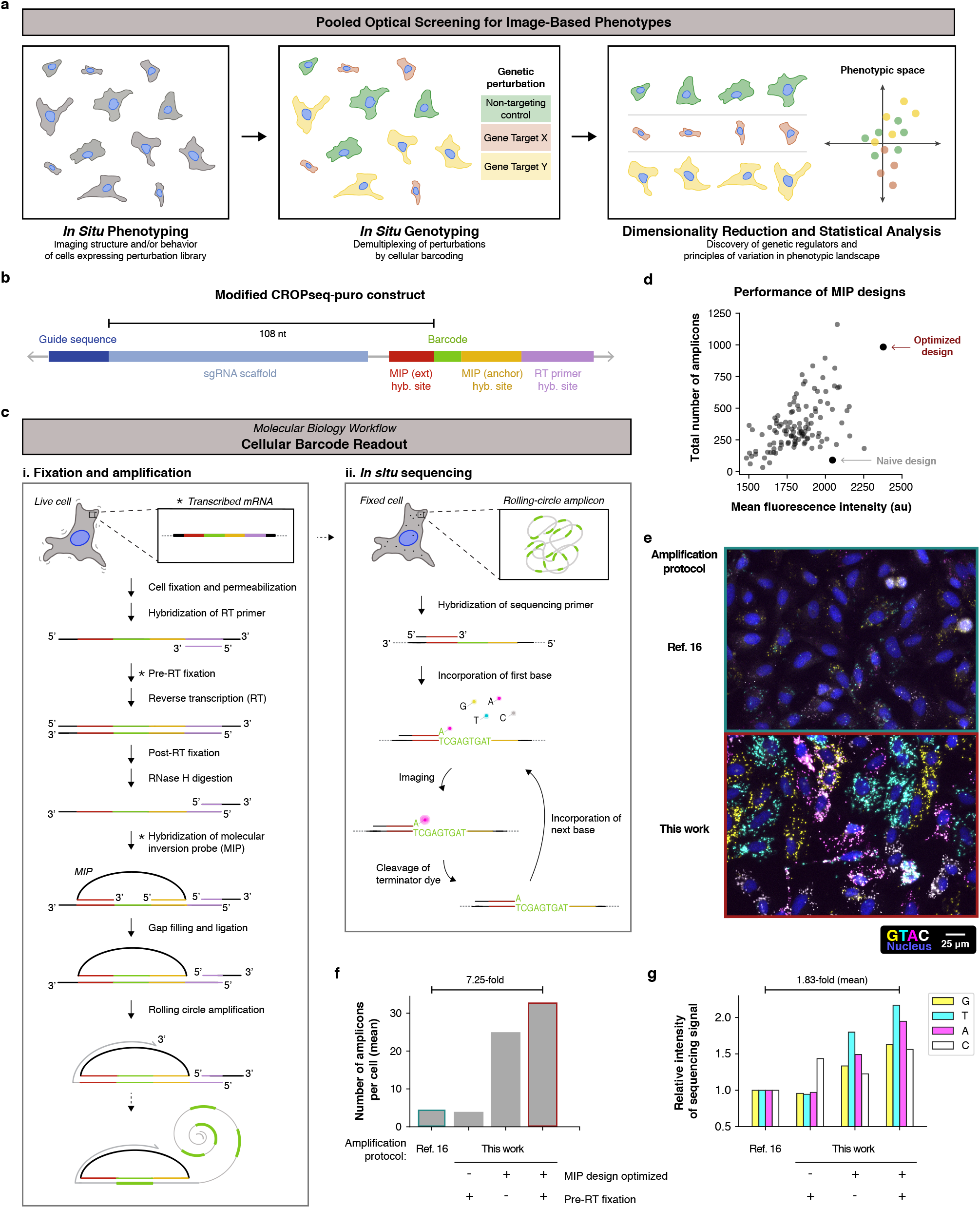
Improved cell barcode readout enables larger-scale pooled screens. **a**, Stages of a pooled screen for high-dimensional image-based phenotypes. **b**, Schematic of the proximally-barcoded sgRNA expression cassette. Minimizing the spacing between guide and barcode sequences reduces lentivirus template switching and simplifies cloning. **c**, Molecular biology workflow for reading out DNA barcodes from single cells, grouped into two stages: (i) cell fixation and *in situ* barcode amplification and (ii) *in situ* sequencing (ISS). Asterisks mark aspects of the protocol for which key optimizations were made in this work. **d**, Number of amplicons (y-axis) and mean fluorescence intensity (x-axis) of barcode readout from each of 113 molecular inversion probes (MIPs, gray circles), tested in a pooled experiment. **e**, Representative ISS images resulting from amplification protocols from reference 16 (top) and this work (bottom), performed side-by-side. Nuclei (blue, Hoechst 33342 stain) and rolling circle amplicons that incorporated fluorescently-labeled nucleotides (spots in yellow, cyan, magenta, and white) are shown. **f**,**g**, Quantification of the mean number of amplicons per cell (**f**) and fluorescence intensity of sequencing signal (**g**) resulting from changes to the *in situ* amplification protocol. The first and fourth conditions correspond to the upper and lower images in (**e**), respectively.

A major challenge in making the best use of optical screen data lies in the data analysis. A screen for cell morphology may show changes in many interdependent cell characteristics like shape, actin distribution, actin intensity, nucleus localization, etc. To enable quantitative comparison between images of different cells, image data must be distilled down to a manageable number of dimensions that capture biologically meaningful axes of variation^18^. The ideal encoding should faithfully and fully capture the rich information contained within cell images; this is especially important for phenotypes whose axes of variation are not easily defined *a priori* or whose perturbation-induced shifts are small compared to the range of natural variation. The ideal method should also yield human-interpretable outputs, so that these phenotypic dimensions can be given meaning and reasoned about, not just used for statistical analysis.

For more than a decade, morphological profiling has been used to capture variation in cell image data by extracting thousands of predefined single-cell features and passing them through one of a growing suite of dimensionality reduction algorithms^19^. While they are versatile and powerful tools, it remains unclear how efficiently these human-curated feature sets capture biologically relevant information and how choices of features or subsequent dimensionality reduction methods may bias the final results. Consequently, deep learning algorithms have recently gained traction as a way to simultaneously extract features and reduce dimensionality of cell images in a data-driven manner without significant imposition of human design. For example, a β-variational autoencoder (β-VAE) based on neural networks has been used to encode 3D images of single cells in a low-dimensional latent space with sufficient fidelity to accurately reconstruct the original images^20^. These latent encodings were then used to uncover principles of 3D organization in cultured human induced pluripotent stem cells. However, these two types of approaches for cell image analysis have not yet been directly compared.

Individual gene contributions to cell morphology and organization have been historically difficult to dissect due to wide intercellular variation even in a single cell type, which can make identification of altered phenotypes challenging. We used pooled optical screening to characterize the genetic factors regulating cell morphology and actin organization (Fig. 1a), taking advantage of the method’s controlled nature and ability to link genotype to phenotype at the single-cell level. We conducted our screen in the human osteosarcoma cell line U2OS, which naturally exhibits wide and continuous morphological variation, in part arising from varied, complex motile behaviors. Investigation of cell-intrinsic shape and organization requires cells to be seeded sparsely, placing practical limits on the scale of our screen. To minimize the effect of these constraints, we developed an optimized barcode amplification method that significantly increases the proportion of cells that can be matched to their genetic perturbation. We employed two methods to analyze cell images – one using morphological profiling features and one using a β-VAE trained on cell images – and compared their performance in identifying gene hits and visualizing phenotypic effects. Altogether, we established a quantitative and interpretable map of the U2OS cell morphological landscape.

## RESULTS

### Improved cell barcode readout enables larger-scale pooled screens

Human U2OS osteosarcoma cells, a well-established model system for mechanistic study of cellular actin networks^21–23^, are an ideal medium to probe genetic determinants of cell morphology and actin organization. In addition to exhibiting a wide range of shapes and complex motile behavior, U2OS cells spread thin (<5 μm in height) over large areas (several tens of micrometers in diameter) when plated on stiff substrates, enabling the use of widefield fluorescence microscopy to visualize fine internal structure^24–26^. To investigate the cell-intrinsic determinants of cell shape and internal organization, plating at low densities is necessary to limit the potentially dominant influence of physical packing among neighboring cells.

However, large cell sizes and low densities also constrain the number of cells per imaged sample area and therefore the total number of genetic perturbations that can be screened (Extended Data Fig. 1). To maximize cell numbers in a screen, common practice is to plate cells in a confluent monolayer, but our application required seeding cells one order of magnitude more sparsely. Low cell density can be compensated for by increasing the surface area of the sample, but only to a limited extent because of the significant handling and microscope time required to image the sample once per nucleotide in the genetic perturbation-associated DNA barcode. Furthermore, in previous protocols, cells were often discarded due to insufficient genotyping information and low barcode amplification efficiency^16^. We therefore optimized the genotyping workflow to increase the number of barcode amplicons generated per cell and thereby maximize the percentage of usable cells.

The established method for identifying the sgRNA in each cell is as follows (Fig. 1b,c). First, cells are fixed and an mRNA containing an sgRNA-associated barcode sequence is reverse-transcribed (RT) to complementary DNA (cDNA) *in situ*. After digesting the RNA, a DNA molecular inversion probe (MIP) is hybridized to the cDNA with 5’ and 3’ ends flanking the barcode. The barcode is added by primer extension onto the MIP, which is then circularized by ligation and copied by rolling circle amplification to produce long, single-stranded DNA amplicons. Finally, the barcode sequence is read by *in situ* sequencing (ISS), one base at a time, by applying Illumina reagents to the sample and imaging the amplicons, which appear as diffraction-limited spots on the fluorescence microscope.

Within this approach, we improved sgRNA identification efficiency by optimizing the readout strategy, MIP design, and cell fixation. First, sgRNA readout can be done in two ways: by amplifying and sequencing either the 20-nt sgRNA targeting sequence itself or a guide-associated DNA barcode^16^. Barcodes are advantageous because they can be designed to distinguish between guides using fewer bases, requiring fewer sequencing rounds, and because the shorter length is thought to allow the MIP to hybridize, elongate, and circularize more efficiently^15^. However, due to template switching during lentivirus generation, barcodes lose their one-to-one association with sgRNAs with frequency proportional to the number of nucleotides between the barcode and sgRNA^27^. We designed a new construct based on the CROP-seq approach^28^ with the barcode placed immediately downstream of the sgRNA (Fig. 1b). This reduced levels of barcode swapping to acceptable levels (7.5% by a digestion-qPCR assay, 7.6% by genomic DNA sequencing, Extended Data Fig. 2a,b), eliminating the need for alternative workflows to minimize template switching using large volumes of low-titer virus^29^.

Next, we tested 113 MIP sequences and selected an optimized design that maximized barcode amplification. To do this, we pooled the probes in equal amounts, used the pool to amplify barcodes integrated into U2OS cells, and distinguished amplicons resulting from each MIP by performing ISS on a MIP-specific barcode^16^ (Supplementary Methods). Performance of each MIP design was determined based on amplicon number and average signal strength, measured by hybridizing a fluorescent oligo to the probe, and was highly reproducible between replicates (Extended Data Fig. 2c). The best MIP from this experiment had 11-fold more amplicons and 1.2-fold higher signal than our naive first design (Fig. 1d). To measure the effect of MIP length, 41 probes had similar sequences but ranged from 72 to 112 nt. The longest and shortest MIPs yielded fewer amplicons (Extended Data Fig. 2d). Interestingly, the 83-nt and 92-nt MIPs performed best but all intermediate lengths were worse, producing up to 3.1-fold fewer amplicons at the midpoint of 88 nt. This matches previous observations that rolling circle amplification yield has a sinusoidal dependence on template length, suggesting that torsional stress of the MIP during hybridization, circularization, and amplification may be a key limiting factor^30^.

Finally, we varied the fixation conditions to reduce the number of RNA or cDNA molecules lost from cells during incubations. Specifically, we added a fixation step after hybridizing the RT primer but before adding enzyme, and we also added a 5’ primary amine to the RT primer to allow direct cross-linking to the cDNA^15^. Altogether, our final protocol yielded an average of 33 amplicons per U2OS cell, which was 7.3-fold more than previously published conditions^16^ run side-by-side, and had 1.8-fold mean increase in sequencing signal intensity across the four channels (Fig. 1f,g). MIP optimization had a large effect on both amplicon number and signal, while pre-RT fixation had a modest but significant effect. This optimized protocol simplified library generation and improved sgRNA readout efficiency, supporting more scalable optical screens.

### A pooled screen assays 366 genes for regulators of cell morphology in over one million cells

To characterize the genetic influences on U2OS cell shape and cytoskeletal structure, we performed a CRISPRi screen focused on genes with known or hypothesized roles in actin organization and related cellular processes, such as cell shape determination, adhesion, and migration. The target gene set was selected via GO-term filtering and complemented by a review of relevant literature (Supplementary Methods). In total, we targeted 366 genes in triplicate using 1098 sgRNA sequences from a list of previously-validated guides^31^, plus 20 non-targeting control guides. Guides were synthesized in a pool and cloned into a library of CROPseq plasmids, each uniquely associated with a 9-nt barcode^28^. The collection of barcodes was designed with a pairwise edit distance of three to allow correction of one sequencing error.

The CRISPRi system enables specific, efficient knockdown of target genes^32^. We generated a U2OS cell line constitutively expressing dCas9-XTEN-KRAB^33,34^, and transduced lentivirus containing our sgRNA library at a low multiplicity of infection (~20%) to ensure that the majority of cells received at most one guide (Fig. 2a(i)). After generating the U2OS CRISPRi cellular library and inducing knockdown for eight days, we seeded cells in two 6-well plates (a total area of 115 cm^2^; Fig. 2a(ii)). In preliminary tests, we found that seeding 100,000 cells per well (~10,400 cells per cm^2^), would maximize cell density without overly crowding the cells (Extended Data Fig. 3), and found a technique for seeding cells that reproducibly gave high uniformity across the well (Supplementary Methods). Thirteen hours after seeding, cells were fixed and cellular barcodes were amplified *in situ* as described above (Fig. 2a(iii)). Actin and nuclei in fixed cells were stained and phenotyping images were collected at 20x magnification, yielding 104,476 fields of view (FOVs) (Fig. 2a(iv)). Next, genotyping images (four nucleotide channels and nucleus channel) were collected at 10x magnification over nine rounds of *in situ* sequencing, yielding 168,948 FOVs (Fig. 2a(v),b).

**Fig. 2:**
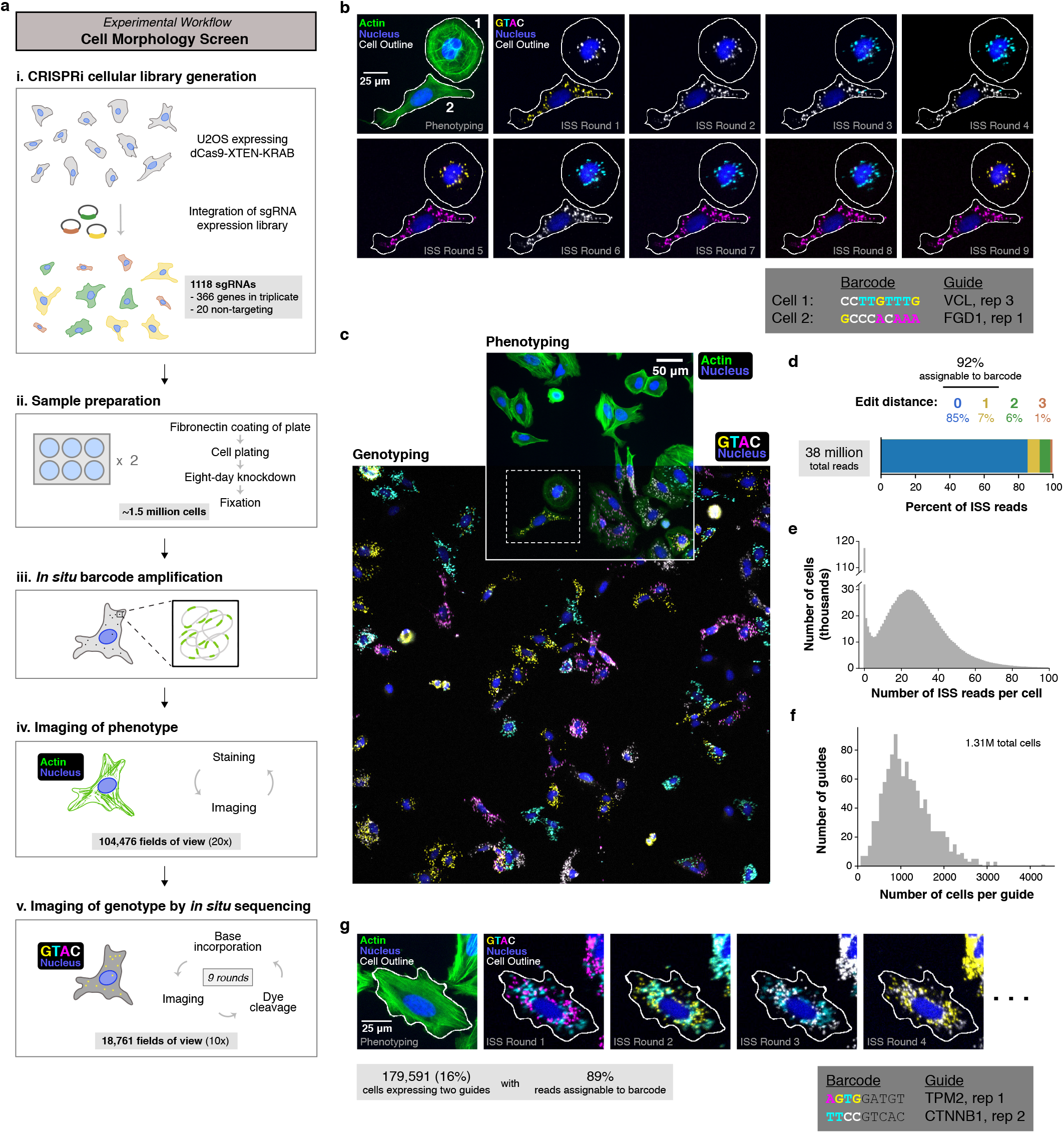
A pooled screen for 366 genes regulating cell morphology in over one million cells. **a**, Experimental workflow of the pooled screen. **b**, Phenotyping image (upper left) and genotyping images over nine rounds of sequencing for two example cells. **c**, Full phenotyping and genotyping fields of view (FOVs) at 20x and 10x magnification, respectively, aligned and overlaid. Dashed box marks the region shown in (**b**). **d**, Distribution of edit distance to a designed barcode for each of the 38M *in situ* sequencing (ISS) reads. **e**, Histogram of the number of ISS reads per cell. **f**, Histogram of the number of cells assigned to each CRISPR guide. **g**, Phenotyping image (left) and genotyping images over four rounds of sequencing for one example cell that contains two distinct barcodes. **b**, **c**,**g**, Images show nuclei in blue (Hoechst 3342), actin in green (Alexa Fluor 488 phalloidin), cell segmentations outlined in white, and amplicons containing fluorescently-labeled nucleotides in yellow, cyan, magenta, and white.

The sgRNAs present in each cell were determined using a custom computational pipeline (Supplementary Methods, Extended Data Fig. 4 and 5). First, amplicons were detected from genotyping images by finding pixels with high variance over all rounds of sequencing, and nucleotide identity of each amplicon in each round was called using a linear classifier. Second, cells and nuclei were segmented from phenotyping images using the generalist deep learning-based segmentation algorithm Cellpose^35^. Third, genotyping and phenotyping images were aligned based on nucleus position, and barcode reads were assigned to each cell (Fig. 2c, Extended Data Fig. 5d). Finally, in each cell, a barcode was designated as consensus if it appeared in at least five independent reads and represented at least 30% of reads in the cell. In total, we obtained 38M sequencing reads, of which 85% perfectly matched a designed barcode and 7% were one mismatch away and therefore error-correctable (97% per-nucleotide sequencing accuracy, Fig. 2d). Consistent with our optimization experiments above (Fig. 1f), we obtained a median of 25 reads per cell (Fig. 2e). The number of reads per cell followed a bimodal distribution, suggesting that cells with no amplicons had mutated or suppressed expression of the barcode. Requiring five independent reads ensured high-confidence consensus calls, and 85.4% of cells were successfully assigned a consensus barcode. sgRNA integration by lentivirus is expected to create a subset of cells containing multiple sgRNA sequences, and indeed, 16% of cells were assigned two consensus barcodes (Extended Data Fig. 5e). Despite the close spatial proximity of amplicons with different sequences (Fig. 2g), reads from cells with two consensus barcodes were highly accurate (89% of these reads could be assigned to a designed barcode), showing efficient and accurate multiplexing of barcode readout.

Altogether, 1.49M cells were imaged and 1.13M cells (76%) were assigned a consensus guide and passed all quality filters (Supplementary Methods). Because we expected guides producing strong perturbation phenotypes to be rare, we included both guides in double-guide cells during downstream analysis, for a total of 1.31M analyzable cellguide pairs. Each guide was represented by a median of 1079 cells, with 99% of guides represented by at least 250 cells (Fig. 2f).

### Principal components of cell morphological profiles can be assigned biological interpretations by visualizing constrained walkthroughs

To reduce the dimensionality of the screen image data, we extracted cell features using morphological profiling and performed principal component analysis (PCA), which together we refer to as feature-based PCA (fPCA) (Fig. 3a, left). We used the image analysis software CellProfiler to extract single-cell morphological profiles consisting of 1054 cell and nucleus features^36^. We identified groups of highly correlated or anti-correlated features using hierarchical clustering and discarded all but one representative feature per cluster. We then performed PCA on the 708 remaining features and derived principal components, which we refer to as fPCs. For downstream analysis, we used the top 25 fPCs, which encapsulate the most prominent modes of variation in the data and together explain 63.9% of the total variance of the input features (Fig. 3c).

**Fig. 3:**
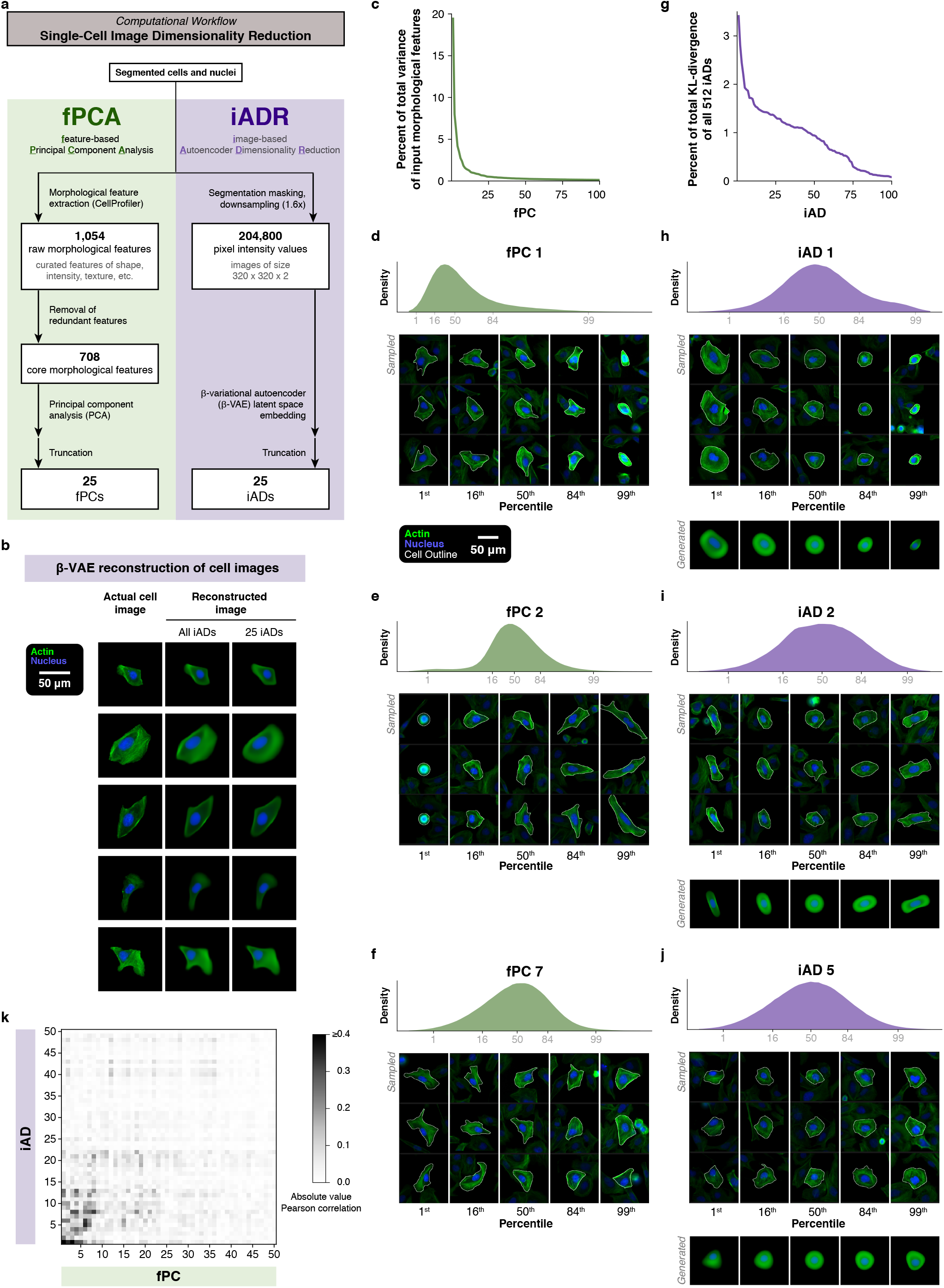
Dimensionality reduction of cell images by morphological feature-based PCA and image-based β-variational autoencoder (β-VAE). **a**, Computational workflow for capturing the important axes of variation using morphological feature-based PCA (green, left) and imagebased β-VAE (purple, right). **b**, For five example cell images (left column), *de novo* reconstruction by the β-VAE using the entire latent space (middle column) or only the top 25 iADs (right column). Nuclear and actin signals outside the segmented nuclei and cells, respectively, were masked out. These cells were not seen by the β-VAE during training. **c**, Percent of total variance of input morphological features explained by each of the top 100 fPCs. **g**, Percent of total KL-divergence of all 512 iADs captured by each of the top 100 iADs. **d-f**,**h-j** Visual Interpretation of Embeddings by constrained Walkthrough Sampling (VIEWS) of three fPCs (**d-f**) and three iADs (**h-j**). Top: density distribution along each dimension for all cells in the dataset, with 1^st^, 16^th^, 50^th^, 84^th^, and 99^th^ percentiles marked. Bottom: images of three cells that fall at each of these percentiles along the given dimension but have near-average values in each other dimension. Images show nuclei in blue (Hoechst 33342), actin in green (Alexa Fluor 488 phalloidin), and cell segmentations outlined in white. Surrounding cells are shown at 50% brightness. In (**h-j**), β-VAE-generated synthetic images of cells traversing each iAD are also shown below. **k**, Absolute value of Pearson’s correlation across all cells in the dataset for each iAD paired with each fPC is shown as a heatmap.

Two aspects of this approach make biological interpretation of fPCs challenging. First, interpreting an fPC based on its composite input features is non-trivial because tens or hundreds of features can contribute to each fPC and because many features themselves lack intuitive interpretations. Second, this approach is not generative, so synthetic cell images, which can serve as aids for visual interpretation, cannot be generated directly from fPC encodings.

We therefore developed a general method to visualize a defined direction in high-dimensional space, which we used to interpret fPCs. We achieve this by sampling images of real cells from the dataset (belonging to any sgRNA population) at specified landmark points along a given fPC axis. Because the information encoded by a given fPC axis can be obscured by high variability among cells along the other axes, we constrain the sampling to only cells that are close to the average along all other fPCs, producing a clearer visualization of trends (Extended Data Fig. 6). Applying this Visual Interpretation of Embeddings by constrained Walkthrough Sampling (VIEWS) to fPC 1, we find variation in cell area, showing a continuum from small oblong cells with bright actin signal throughout the cell area to larger cells with more complex shapes and dimmer actin signal, especially in the perinuclear region (Fig. 3d). VIEWS of fPC 2 suggests efficient encoding of a complex set of characteristics associated with cell length (Fig. 3e). Inspection of VIEWS for other fPCs using this method revealed variation along diverse biologically meaningful axes, such as nuclear positioning (fPC 7 and 6; Fig. 3f and Extended Data Fig. 7a, respectively), actin stain intensity (fPC 1 and 4; Extended Data Fig. 7b), and actin stain distribution (fPC 19 and 22; Extended Data Fig. 7c,d).

### A β-variational autoencoder accurately reconstructs cell images and produces an embedding space distinct from morphological feature-based PCA

For an alternative approach to image analysis, we turned to the β-variational autoencoder (β-VAE), a deep learning algorithm consisting of an encoder and decoder that learns to embed input data in a latent space of reduced dimensionality while balancing reconstruction accuracy with embedding simplicity^37^. This image-based autoencoder dimensionality reduction (iADR) (Fig. 3a, right) has three potential advantages compared to fPCA. First, the β-VAE directly maps image data to a learned, low-dimensional representation, without the need to curate hundreds of human-defined input features. Second, this mapping is nonlinear, which may result in a more expressive, compact phenotypic encoding than PCA, which is constrained to be linear with orthogonal axes. Third, a β-VAE is generative—synthetic cell images can be directly generated from encodings–which provides a powerful tool for human interpretation of what the model has learned.

We adapted a β-VAE architecture originally developed for analysis of 3D cell images^20^ to accept 2D images in two fluorescent channels (actin and DNA) as input. After training on segmented images of individual cells, the model accurately reconstructed cells in the held-out test set (Fig. 3b, Extended Data Fig. 8). Reconstruction was accurate for large-scale features such as cell size and shape; nuclear size, shape, and positioning; and actin intensity and distribution within the cell. However, fine-grained features such as individual stress fibers and nuclear texture were reconstructed less accurately. We next examined the 512 image-based autoencoder dimensions (iADs) in the network’s latent space, sorted in order by the magnitude of the Kullback-Leibler (KL) divergence term in the objective function used for training (Donovan-Maiye et al., 2022). The first 25 iADs together captured 43.2% of the total KL-divergence summed across all dimensions of the latent space (Fig. 3g). (Note however that this metric is not directly comparable to the 63.9% of variance explained by the first 25 fPCs.) The autoencoder also distributed information more evenly across dimensions than did PCA (Fig. 3c,g).

VIEWS of iADs revealed that individual iADs encoded biologically interpretable features, and many appeared to capture information similar to individual fPCs. For example, iAD 1 encoded information related to cell area (Fig. 3h, similar to fPC 1), iAD 2 corresponded with cell length (Fig. 3i, similar to fPC 2), and iAD 5 captured information related to cell polarity and nuclear position (Fig. 3j, similar to fPC 7). Generated synthetic images of cells traversing each iAD recapitulated patterns observed in the traversals of real cells, although with high-frequency information smoothed out (Fig. 3h–j, bottom). In contrast with fPCs, which were derived from rotationally-invariant CellProfiler features, many iADs included information about cell orientation, which is not a biologically significant form of phenotypic variation in our experiment.

To assess the interrelationships between individual axes of the embedding spaces derived from these two methods, we quantified the absolute-valued correlations between each fPC and each iAD across all cells (Fig. 3k). Modest correlations were observed between many pairs of axes but were not one-to-one, suggesting that the two spaces capture some similar quantitative information encoded in the images, but distribute it across axes differently. For example, although VIEWS of fPC 1 and iAD 1 both show encoding of cell size-related information, iAD 1 correlates with all five of the top-ranked fPCs (with absolute Pearson’s r ranging from 0.28 to 0.53). Thus, fPCA and iADR offer complementary options for dimensionality reduction of single-cell U2OS image data, resulting in distinct phenotypic embedding spaces.

### Two dimensionality reduction methods identify many overlapping gene hits

To identify hits in our screen, we used the data-driven dimensions defined by fPCA and iADR to identify guide populations with altered phenotypes. For each dimensionality reduction method, a “gene hit” was called if at least two of its three sgRNAs each had a statistically significant difference relative to cells containing nontargeting control sgRNAs in at least one of the top 25 dimensions (p < 10^-6^ by two-sample Kolmogorov-Smirnov test, per-guide false discovery rate of 2.7%). The individual guides contributing to each gene hit were referred to as “guide hits.” We identified 31 gene hits using fPCs and 16 gene hits using iADs, 15 of which were shared (Fig. 4a). Shared gene hits included those with known roles in cell morphology and actin organization, such as in adhesion (e.g. vinculin – VCL), contractility (e.g. nonmuscle myosin IIA heavy chain 9 – MYH9), actin filament nucleation and branching (e.g. actin-related protein 2 – ACTR2), and actin filament capping (e.g. capping actin protein of muscle Z-line subunit beta – CAPZB). Additionally, 33 and 24 genes had only one significant guide by fPCA and iADR, respectively, which we denote as “non-replicated significant genes” and revisit in the next section.

**Fig. 4:**
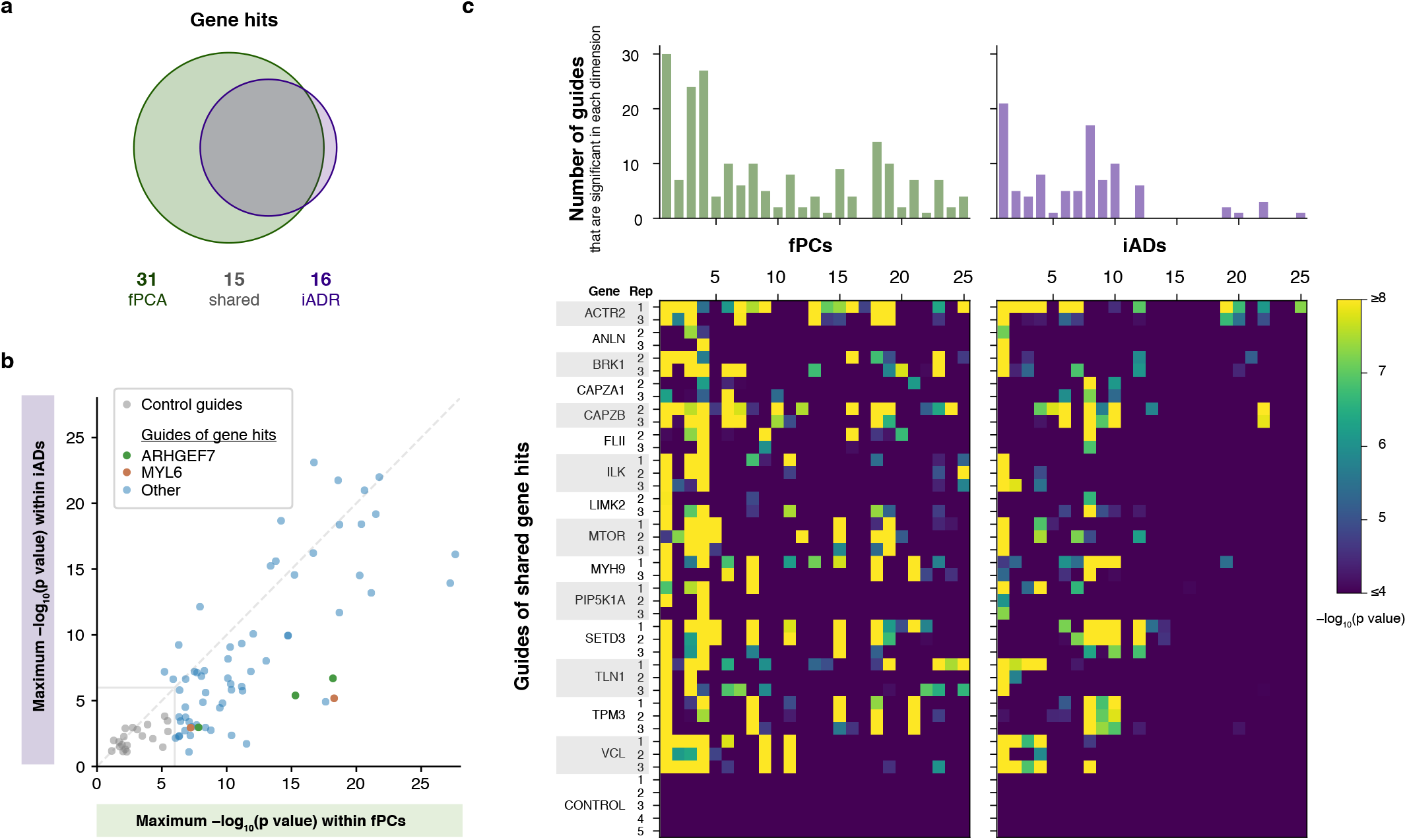
Two dimensionality reduction methods identify many overlapping gene hits. **a**, Venn diagram of gene hits identified using fPCA (green) and iADR (purple). **b**, For each hit (blue) and non-targeting control (gray) guide, the maximum -log_10_(p value) across all fPCs and across all iADs are plotted. Guide hits for two genes, ARHGEF7 (green) and MYL6 (orange), are highlighted. The diagonal dashed line shows equality; the square in the lower left shows the significance threshold for calling hits (p < 10^-6^). Eleven guides have x- or y-values higher than 30 and are not shown. p values were calculated by two-sample Kolmogorov-Smirnov test relative to control guides. **c**, Bottom: for each of the 37 guide hits shared by fPCA and iADR, significance (-log_10_(p value)) in each of the top 25 fPCs (left) and iADs (right) is plotted as a heatmap. Five non-targeting control guides are also shown. Top: the number of guides that are significant (p < 10^-6^) in each dimension is shown as a bar chart.

To explore differences in hit calling between fPCA and iADR, we compared the most significant p value produced by each method for each guide hit (Fig. 4b). p values in iADs were equally or less significant than in fPCs. Notably, all the replicates of some gene hits, such as nonmuscle myosin II regulatory light chain 6 (MYL6) and Rho guanine nucleotide exchange factor 7 (ARHGEF7), were consistently less significant by iADR. We hypothesized that this could be due to certain modes of variation that fPCA is sensitive to but that iADR is insensitive to or fails to capture. Consistent with this, when we computed p values for these two genes across each input CellProfiler feature, features corresponding to texture, a property which the β-VAE reconstructs poorly, were the most significant (over-represented 4.8-fold within the top 20 most significant features).

To better understand which fPCs and iADs were important for identifying the hits, we examined fPC and iAD p value profiles for the 37 guide hits that comprise the set of shared gene hits (Fig. 4c). Most of these guides exhibited significant effects in multiple fPCs and iADs. For both embedding spaces, the rank of the dimension did not exactly correspond with the frequency of being altered across shared guide hits; for example, the top three most frequently affected iADs were iADs 1, 8, and 10. The two embedding spaces differed in that significant guides were detected throughout the top 25 fPCs but were heavily concentrated in just the top 12 iADs, suggesting that iADR more compactly encodes the distinguishing biological information necessary to identify altered phenotypes.

### The linear discriminant vector defines the direction of phenotypic shift and facilitates calling 13 additional gene hits

Replicate guides were significant in the same sets of dimensions, suggesting that they caused a similar shift in phenotype (Fig. 4c). Some guides targeting different genes also shared the same significant dimensions, suggesting that they produced similar or opposite phenotypes. To quantify this, we summarized the direction of phenotypic shift caused by each guide by computing the direction in embedding space along which the knockdown population was best separated from the control population, also known as the linear discriminant (LD) (Fig. 5a). For each guide, this yielded a unit-length LD vector that points in the direction of the knockdown phenotype and encodes a weighted importance of each fPC or each iAD in describing the phenotype. LD vectors were highly correlated between pairs of replicate guide hits (mean Pearson’s r = 0.806 and 0.886 in fPC and iAD space, respectively) but largely uncorrelated for pairs of control guides (mean Pearson’s r = −0.043 and −0.036 in fPC and iAD space, respectively; Fig. 5b, Extended Data Fig. 9a). Thus, significant guides targeting the same gene had high phenotypic concordance.

**Fig. 5:**
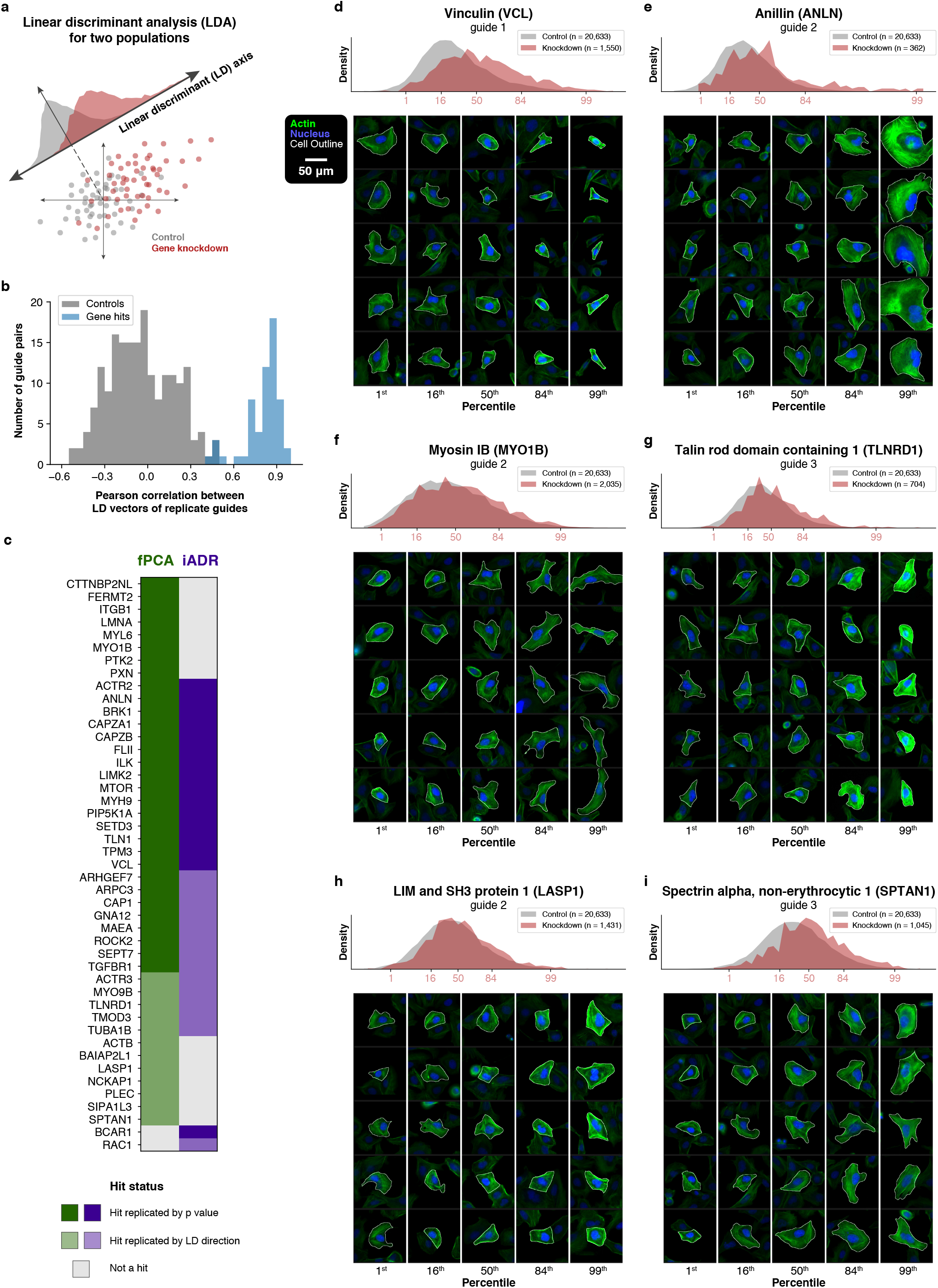
Definition and visualization of phenotypic shifts caused by gene knockdowns. **a**, Schematic: the linear discriminant (LD) axis (thick diagonal line) is the direction that produces the best separation between two populations in high-dimensional space (gray and maroon dots). **b**, Histogram of Pearson correlation values between LD vectors of two non-targeting control guides (gray) or between LD vectors of two replicate guide hits (blue). LD vectors are calculated in fPCA space. **c**, All gene hits are listed, with color indicating if they were detected using fPCA (green) or iADR (purple) and whether they were found by two or more replicate guides with significant p value (dark color) or by correlated LD vectors (light color). **d-i**, VIEWS plots of the LD vector between control cells (gray) and knockdown cells (maroon) for six different guide hits. Density plots show the distribution of each cell population projected onto the LD axis. Cell images are sampled from the entire dataset and are displayed in the same manner as in Fig. 3. n, number of cells in each population. In all panels, LD vectors are calculated using the first 25 fPCs.

Because the magnitude of gene knockdown varies between guides^31^, we suspected that some non-replicated significant genes only had one significant guide because the other replicates only had weak phenotypic shifts. We therefore asked whether the direction of the LD vectors could reveal a concordant phenotypic shift between replicates, confirming reproducibility of the hit based on direction rather than magnitude of effect. Specifically, for each nonreplicated significant guide, we checked whether its LD vector had a Pearson correlation of at least 0.6 with the LD vector of any of its replicate sgRNAs. In this way, we confirmed 12 additional “LD-validated” gene hits from fPCA and 14 from iADR (Fig. 5c). Indeed, eight of the 14 LD-validated gene hits from iADR were replicated gene hits from fPCA, and five of the remaining six LD-validated gene hits from iADR were also LD-validated gene hits from fPCA. These observations show that LD-validation can reliably identify biologically meaningful gene hits, even when the wild-type phenotype is highly variable and the phenotypes of control and knockdown cells largely overlap.

### Precise, interpretable visualization of phenotypic shifts in cell populations

Having summarized hit phenotypes in a quantitative manner, we then sought to visualize and interpret these phenotypes. Inspecting cells sampled from the knockdown and control populations sometimes suggested potential phenotypic differences, but this often failed to definitively present the key difference between the two populations (Extended Data Fig. 9c). We therefore applied VIEWS to the axis defined by each guide hit’s LD vector, using landmark points defined by specified percentiles in the knockdown population but still sampling cell images from the full dataset. Because LD VIEWS in fPC and iAD space were visually similar, except that LD VIEWS in iAD space tended to show rotationally symmetric cells, we focused on LD VIEWS in fPC space. VIEWS for vinculin (VCL, Fig. 5d), a focal adhesion protein important for cell spreading^38^, revealed a clear phenotypic shift upon knockdown from large, well-spread cells with well-organized actin networks towards smaller, slightly elongated or triangular cells with actin accumulation near their edges, consistent with impaired cell spreading. VIEWS for all three VCL replicate sgRNAs were visually similar, consistent with the high pairwise Pearson correlations of their LD vectors (Extended Data Fig. 9b). Thus, VIEWS applied to LD axes provides focused, intuitive visualizations of knockdown phenotypes that enable human interpretation.

VIEWS across all hits (included in Supplemental Data 1) exhibit a wide range of phenotypes. For example, VIEWS for knockdown of anillin (ANLN, Fig. 5e) reveals a shift toward extremely large, round cells with multilobed nuclei, consistent with the known requirement for anillin for normal completion of cytokinesis^39^. VIEWS for unconventional myosin-Ib (MYO1B, Fig. 5f) depicts a shift from small round cells with centered nuclei and bright actin signal near the cell edge toward elongated, crescent-shaped cells with wide lamellipodium-like structures and nuclei positioned along one of the cell’s long edges, resembling fast, migratory U2OS cells^40^. This observation is consistent with a potential role for MYO1B in suppressing U2OS cell polarization or migration, in contrast with a previously described pro-migratory role in cervical cancer cells^41^. VIEWS for talin rod domain containing 1 (TLNRD1, Fig. 5g) exhibits a shift from dim actin signal to brighter, more prominent stress fibers throughout the entire cell, with no notable change in cell shape. Given past work characterizing TLNRD1 as an actin bundler that localizes to thick stress fibers^42^, the VIEWS result suggests a role for TLNRD1 in limiting stress fiber thickness. Inspection of the LIM and SH3 domain protein 1 (LASP1, Fig. 5h) phenotype suggests a shift toward cells with multilobed nuclei, as well as bright peripheral actin with fewer lamellipodium-like cell edges. This actin phenotype is consistent with reports supporting a role for LASP1 in activating branched actin network growth^43^, but the nuclear phenotype suggests a yet undescribed role in nuclear division or nuclear envelope integrity. VIEWS for spectrin alpha, non-erythrocytic 1 (SPTAN1, Fig. 5i), a component of the actin-associated, filamentous-network-forming heterodimer fodrin, depicts a shift from cells with less actin in the perinuclear regions toward cells with “fuzzier” actin distributed throughout the entire cell area consisting of less distinct stress fibers. This suggests SPTAN1 may govern stress fiber formation and distribution throughout the cell, consistent with prior work in human foreskin fibroblasts suggesting a mutually exclusive distribution of fodrin and focal adhesion proteins^44^. Thus, VIEWS can be used for visual representation and interpretation of diverse morphological phenotypes, including complex combinations of variations in cell shape; actin intensity, organization, and distribution; and nuclear morphology and localization.

### A morphological landscape of U2OS cells

With an LD encoding the phenotype of each sgRNA population as a directional vector in both fPC space and iAD space, we reasoned that we could identify groups of gene hits with similar phenotypes by comparing these vectors. Starting with the 15 replicated gene hits identified as significant using both methods, we calculated the pairwise Pearson correlations for each of the individual sgRNA hits in both spaces and performed hierarchical clustering (Fig. 6a). Inspection of the clusters indicated that this method reliably grouped genes whose knockdown would be expected to lead to similar phenotypes; for example, both the alpha and beta subunits of the heterodimeric actin filament capping protein CapZ (CAPZA1 and CAPZB, respectively) were clustered together (Fig. 6a, cyan). Similarly, genes encoding three proteins known to be major structural components of focal adhesions, talin (TLN1), integrin-linked kinase (ILK), and vinculin (VCL), also clustered tightly together (Fig. 6a, red). Importantly, the pattern of similarity between pairs of sgRNAs in fPC space was highly congruent with their similarity in iAD space. This further confirms our finding that the two independent phenotypic analysis methods lead to similar biological conclusions.

**Fig. 6:**
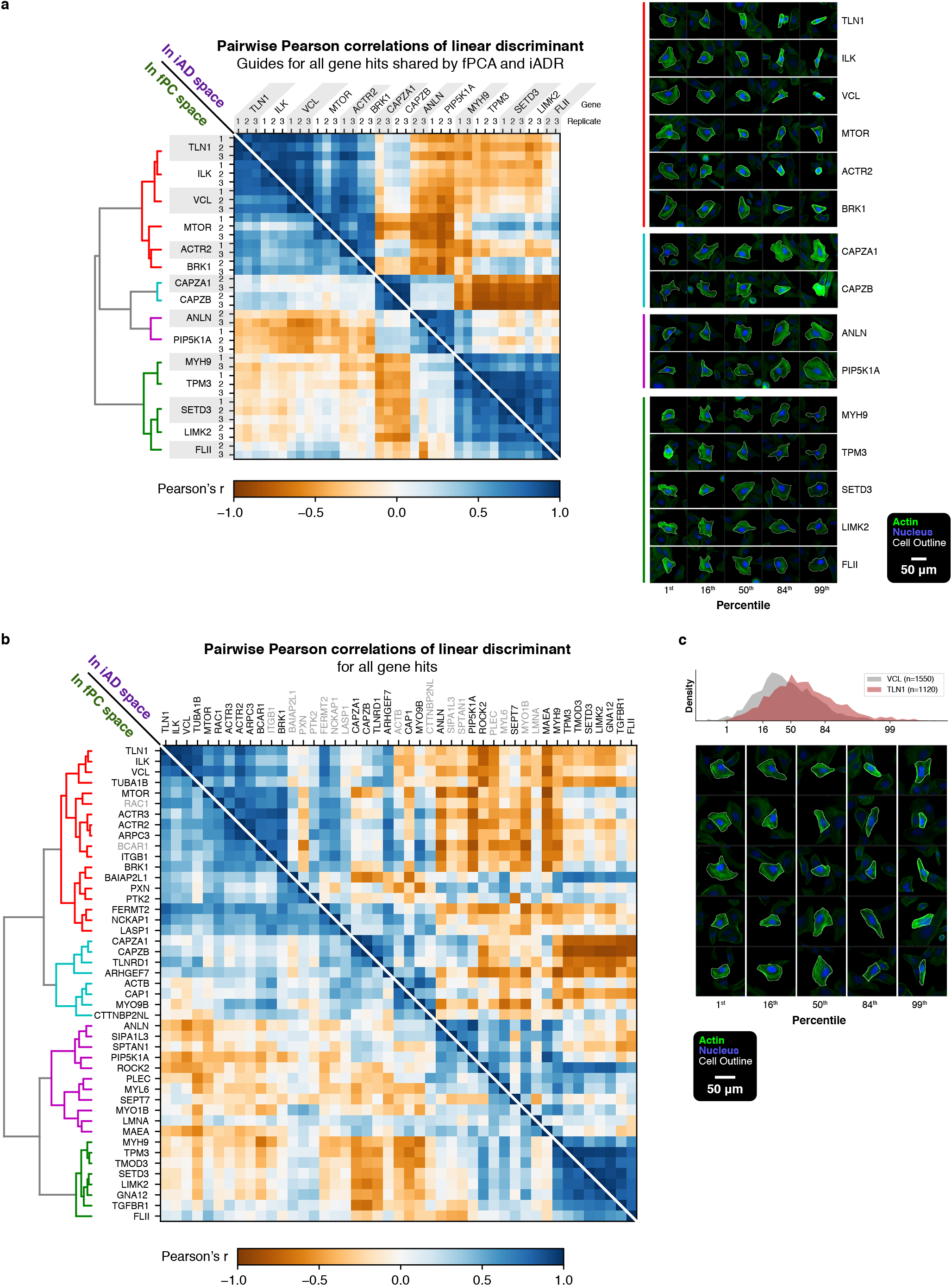
A morphological landscape of U2OS cells. **a**, Left: heatmap of Pearson correlation between LD vectors of all pairs of sgRNAs of replicated gene hits. Values below and above the diagonal use LD vectors calculated with the first 25 fPCs and the first 15 iADs, respectively. Only guide hits shared by fPCA and iADR are shown. Dendrogram from hierarchical clustering in fPC space is shown with the four major clusters highlighted in color. Right: VIEWS of LD vectors of one guide from each gene, shown in the same order as in the heatmap and grouped into the four clusters. **b**, Same as (**a**), but showing all gene hits identified by fPCA or iADR. Genes that are not hits by either fPCA or iADR are displayed in gray text on the corresponding half of the heatmap. Dendrogram from hierarchical clustering in fPC space is shown with the same four major clusters highlighted in color. **c**, VIEWS of the LD vector describing the phenotypic shift between VCL knockdown cells (replicate guide 1) and TLN1 knockdown cells (replicate guide 1).

VIEWS generated for the LD vector of one sgRNA for each of the replicated hits confirmed that the hits that were clustered together did indeed cause similar phenotypic shifts in U2OS cells (Fig. 6a, right). The major cluster of six genes indicated in red all displayed knockdown phenotypes with poorly spread cells that had straight borders and few lamellipodia, consistent with a primary defect in adhesion to the substrate. The cyan cluster representing CapZ displayed a phenotypic shift toward cells with an increased density of filamentous actin, as would be expected for cells defective in the actin capping activity that normally limits filament elongation. The purple cluster displayed an overrepresentation of large cells with multiple nuclei, typified by cytokinetic protein anillin (ANLN)^39^. Finally, the green cluster, typified by MYH9, displayed a phenotypic shift toward cells with irregular borders and lower densities of contractile actin stress fibers, consistent with a primary defect in contractility. Quantitatively, the cluster relating to cellular contractility displayed phenotypes that were strongly negatively correlated with the CapZ phenotypes, consistent with the simple interpretation that these gene knockdowns had the overall opposite effects of decreasing versus increasing the number and size of actin-rich stress fibers, respectively.

In addition to the expected linkages that we found between genes whose products are known to interact, our similarity analysis generated several unexpected correlations that could generate hypotheses for further experimentation. For example, the gene knockdown showing the strongest similarity to ANLN is PIP5K1A, encoding phosphatidylinositol-4-phosphate 5-kinase type 1 alpha, which is involved in the PI3K-AKT signaling pathway. Overexpression of PIP5K1A has been correlated with poor outcomes in several types of metastatic cancer^45,46^, but a role in cytokinesis has not previously been described. A second particularly intriguing replicated hit is SETD3, a histidine methyltransferase whose only known target is actin, but whose exact role in regulation of the actin cytoskeleton is unclear^47^. Our phenotypic analysis reveals that the cellular-level effect of SETD3 knockdown is similar to the effect of other sgRNAs targeting proteins involved in stress fiber formation or contraction, including MYH9, TPM3, which encodes an isoform of tropomyosin, and LIMK2, which encodes a downstream target of RhoA, the master regulator of myosin-based contractility^48^. The possibility that SETD3 methylation of actin might be involved in regulating cellular contractility is particularly intriguing given its recently described role in regulation of smooth muscle contraction, resulting in primary dystocia (delayed labor) in pregnant mice deficient in this gene^49^. Finally, the mechanistic target of rapamycin kinase (MTOR) has been implicated in general regulation of actin cytoskeleton dynamics in several cell types including neurons^50,51^. We find that its knockdown in U2OS cells causes a phenotype similar to knockdown of ACTR2, which encodes one of the subunits of the Arp2/3 complex responsible for branched actin filament nucleation, and knockdown of BRK1, which encodes one of the subunits of the SCAR/WAVE complex that is known to be an activator of Arp2/3 activity. Specific connections between mTOR and Arp2/3 activation have recently been implicated in pancreatic cancer^52^.

Encouraged by these findings, we expanded our phenotypic similarity analysis (Fig. 6b) to include all genes identified as replicated hits by either the fPCA or iADR method (as shown in Fig. 5c). This analysis revealed additional examples confirming the specificity of our quantitative phenotypic comparisons, for example clustering ACTR2 with genes encoding two other subunits of the stable heptameric Arp2/3 protein complex, ACTR3 and ARPC3. It has previously been shown that disruption of individual subunits of this complex destabilizes the entire complex^53,54^, so it is to be expected that the knockdown phenotypes of distinct subunits should be virtually identical.

A closer examination of these phenotypic clusters suggests potentially interesting functional relationships. For example, the phenotype caused by knockdown of the gene encoding an isoform of alpha-tubulin (TUBA1B) is most similar to the phenotype caused by knockdown of genes encoding proteins that are structural components of focal adhesions (TLN1, ILK and VCL). It is intriguing to speculate that this phenotypic similarity may be related to specific interactions with dynamic microtubule ends that regulate focal adhesion stability^55^. In addition, despite the overall similarity between TLN1 and VCL knockdown phenotypes, intergene VIEWS on the LD axis between the TLN1 and VCL knockdown populations revealed that TLN1 knockdown resulted in brighter actin signal at the cell periphery, compared to VCL knockdown (Fig. 6c). This effect was reproducible between all replicate guides, suggesting that these two directly interacting focal adhesion proteins have distinct roles in actin dynamics at the cell periphery (Extended Data Fig. 9d,e).

## DISCUSSION

In this work, we demonstrate the power of optical pooled screening for investigating single-cell shape and actin cytoskeletal organization, where subtle phenotypes were revealed by minimizing technical variation, scaling the number of cells, and detecting small shifts using linear discriminant analysis. The substantial improvements in *in situ* amplification efficiency yielding an average of 33 amplicons per cell allowed the large majority of cells to be successfully genotyped here, and in future applications, may enable readout of combinatorial knockdowns and screens in challenging cell types or organoids. These improvements could also be adapted to perform spatially-resolved lineage tracing in tissues, which also requires high-throughput microscopic readout of multiple distinct DNA barcodes from each cell^56,57^.

In our analysis of our large cell biological image dataset, we provide a direct comparison between conventional morphological profiling and a deep learning-based variational autoencoder for dimensionality reduction. Despite generating distinct mappings for cellular phenotypes, both methods detected a largely overlapping set of gene hits and distributed them similarly within the respective phenotypic spaces. The autoencoder approach captured a subset of the hits identified by the morphological profiling approach, which we attribute to poorer encoding of fine-grained features like texture and over-encoding of orientation-based information. With additional developments in texture encoding and rotational invariance^58^, we expect the variational autoencoder can serve as a generalizable dimensionality reduction approach for the unbiased featurization of cell images.

Although the 366 genes screened here were chosen based on validated or hypothesized association with actin-based processes, only around 10% were hits. What should be made of this? This hit rate could be partially attributable to limitations in the technology or format of the high-throughput screen. First, despite continuous improvements in CRISPRi technology, knockdown efficiency is still widely variable between different guides, target genes, and cell types; using more replicate guides in a future screen could mitigate this problem. Second, even if strong knockdown is achieved, homeostatic adaptations in gene expression over the eight-day knockdown period could obscure the primary effects. Third, even a high-content screen only captures a portion of the physical state of the cell; here, we focused on actin and nuclei of fixed cells, but staining additional structures or time-lapse imaging could reveal new phenotypic effects and identify new hits. Even considering these limitations, the small proportion of hits may also reflect a biological truth of functional redundancy or functional sparsity. In other words, most of the genes linked to actin-based processes in the literature may have significant functional overlap with other genes and/or have roles evident only in specific cell systems and contexts.

As the number of cells and the complexity of phenotypes measured in screens has rapidly increased, analysis methods have accordingly emphasized quantitative description and dissection of phenotypes over direct visualization. However, while quantitative rigor is indeed necessary when calling hits, our ultimate goal was to understand the major axes of variation and to communicate them in a clear and intuitive way. Here, in developing the VIEWS approach to precisely visualize axes of high-dimensional spaces, we have taken care to keep interpretability at the forefront without sacrificing rigor. We used VIEWS to demonstrate that meaningful biological information was encoded in the dimensions, interpret altered phenotypes, and generate hypotheses about the cell as a system. In this new era of big data in cell biology, as exemplified by scalable optical screening, we anticipate a balance of quantitation and interpretation will be key to deriving lasting, biological insights.

## ACKNOWLEDGEMENTS

We thank Aaron Straight, James Spudich, Linda Song, Rene Ladurner, and Chao Liu for sharing use of and assisting with the fluorescence microscope; Roger Kornberg for providing equipment and lab space; Matt Footer for providing invaluable support; Marco Jost, Jonathan Weissman, and Sean Collins for providing the dCas9-XTEN-KRAB expression plasmid; Erez Lieberman Aiden for providing sequencing; Christopher Prinz for assistance with figures; James Spudich, Alex Dunn, James Ferrell, Zachary Pincus, Roger Kornberg, Peter Geiduschek, Benjamin Yeh, and members of the Theriot and Kornberg Labs for helpful discussions. R.L.D.L. was supported by Stanford University Medical Scientist Training Program grant T32-GM007365 and T32-GM145402. A.L.S. was supported by a National Defense Science & Engineering Graduate (NDSEG) Fellowship. J.A.T. was supported by the Howard Hughes Medical Institute (HHMI) and the Washington Research Foundation. Microscopy instrumentation was supported by NIH grant S10RR026775.

This article is subject to HHMI’s Open Access to Publications policy. HHMI lab heads have previously granted a nonexclusive CC BY 4.0 license to the public and a sublicensable license to HHMI in their research articles. Pursuant to those licenses, the author-accepted manuscript of this article can be made freely available under a CC BY 4.0 license immediately upon publication.

## Author contributions

A.L.S. and R.L.D.L. conceived of this work; C.V.H., R.L.D.L., A.L.S., and N.M.B. generated cell lines; C.V.H. and A.L.S. optimized barcode readout conditions; R.L.D.L., A.L.S., and C.V.H. designed and performed the screen experiment; C.V.H. and M.M. established computational infrastructure with assistance from E.M.B., R.L.D.L., and A.L.S.; C.V.H., A.L.S., and R.L.D.L. processed screen image data; A.L.S. and R.L.D.L. analyzed screen results with assistance from C.V.H.; R.L.D.L., A.L.S., and J.A.T. interpreted results; C.K.C. and A.L.S. developed the variational autoencoder; J.A.T. and A.L.S. supervised research; J.A.T. provided funding and resources; R.L.D.L and A.L.S. wrote the paper with input from all authors.

