## Supplemental Methods for "Mapping variation in the morphological landscape of human cells with optical pooled CRISPRi screening"

### Extended Data Figures

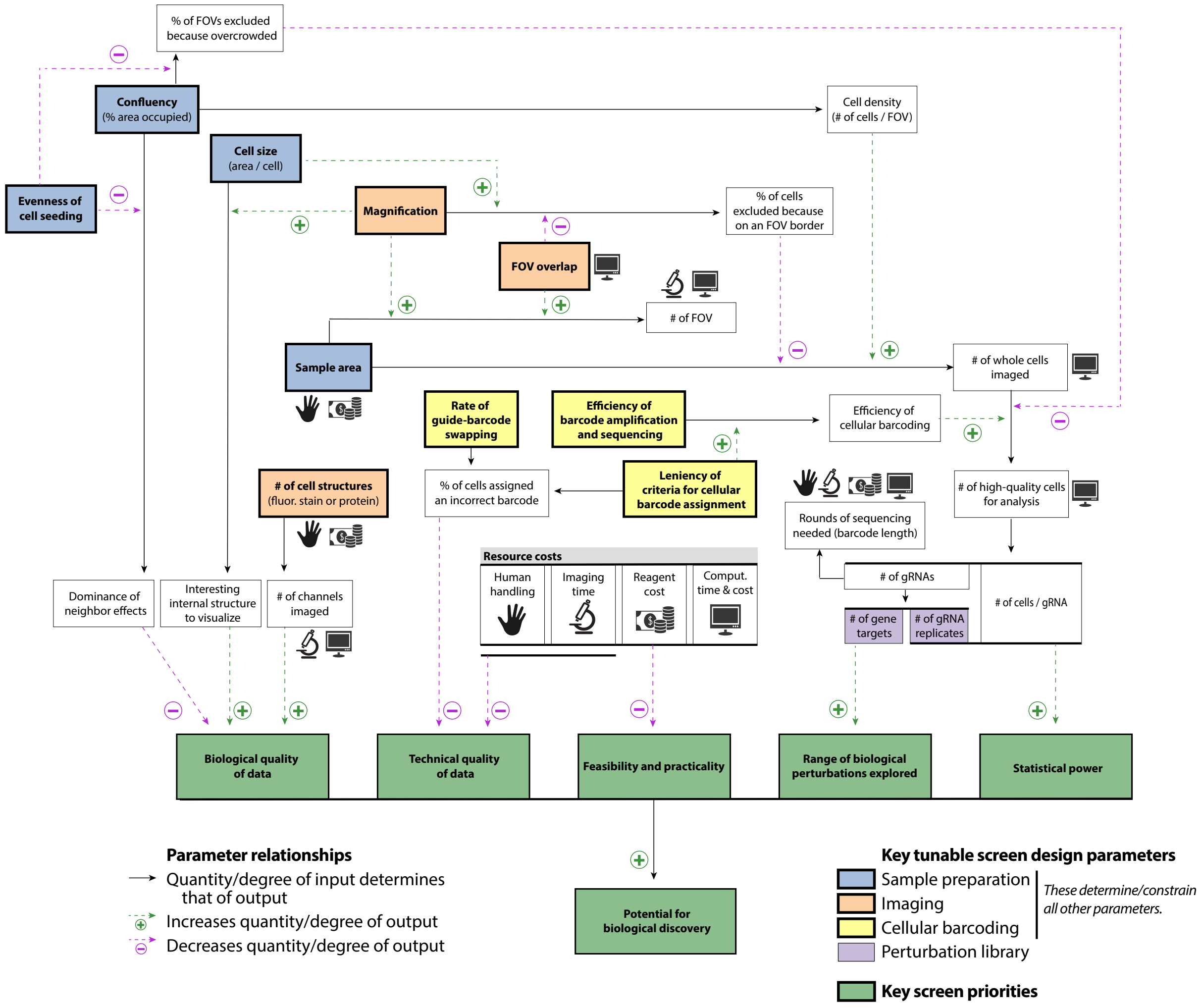


**Extended Data Fig. 1: Priorities and tradeoffs in the design of screens for image-based cellular phenotypes.**

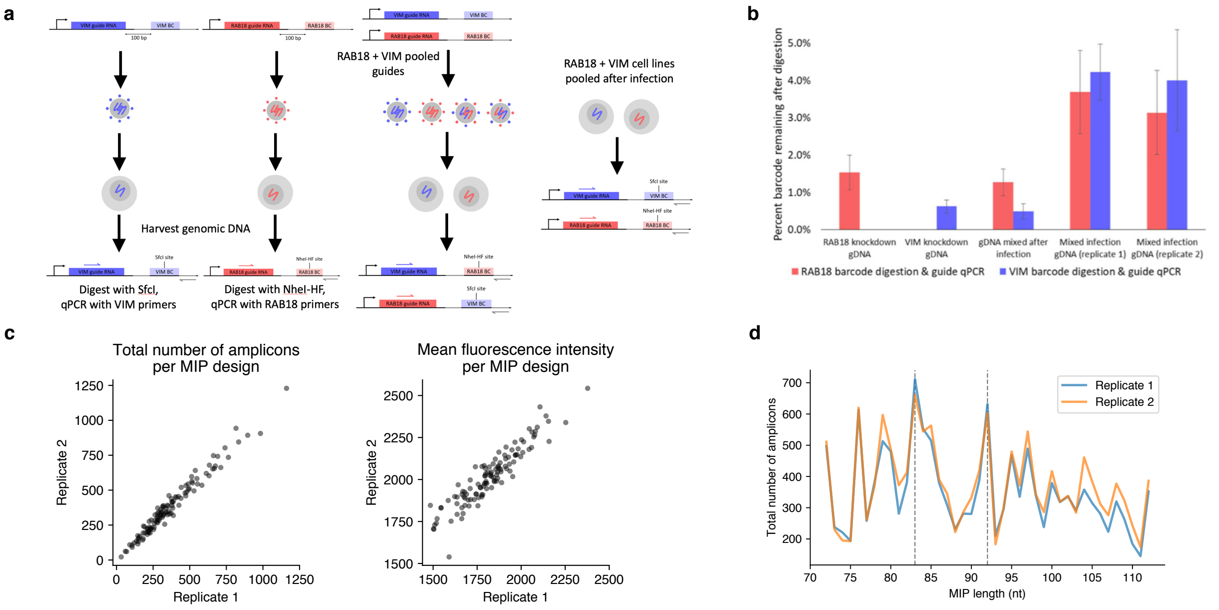


**Extended Data Fig. 2: Optimization of barcode construct design and *in situ* amplification.**
**a**, Schematic of the RT-qPCR assay to measure barcode swapping due to lentivirus template switching. See the Supplementary Methods for details. **b**, Data from the barcode swapping RT-qPCR assay. Y-axis shows the amount of template remaining after digestion, normalized to the amount of input (mock digested) template. Error bars show standard deviation of measurements from triplicate digestions. The average percentage of undigested barcode in mixed infection cells was 3.4% for RAB18 and 4.1% for VIM. Since only half of swapping events are observed in each assay condition, this indicates that swapping occurs in around 7.5% of integration events. **c**, The number (left) and mean fluorescence intensity (right) of amplicons produced by each MIP design from two replicate wells of the pooled MIP optimization experiment. **d**, The number of amplicons produced by a series of MIPs with similar sequence ranging from 72 to 112 nt. Two replicates are shown. The lengths of the two best MIPs are emphasized with dashed lines.

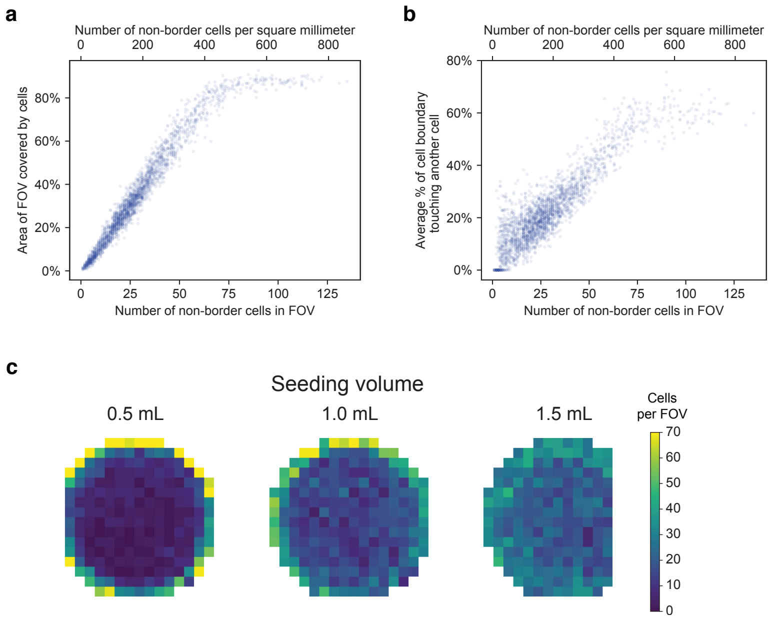


**Extended Data Fig. 3: Optimization of cell seeding conditions.**
**a**,**b**, U2OS cells were seeded in 6-well plates at a variety of densities, stained, imaged at 20x magnification, and computationally segmented (Supplementary Methods). Shown here is the relationship between the number of cells in a field of view (FOV) and (**a**) the percentage of the FOV surface covered by cells, or (**b**) the average percentage of cell boundaries that are touching each other. Each point represents one FOV. Counts exclude cells that touch the border of the image. **c**, Heatmap of cell densities across three wells of a 6-well plate in which an equal number of total cells was seeded in 0.5, 1.0, or 1.5 mL of media. Each pixel represents one FOV. The left and right edges of the 0.5 mL and 1.5 mL wells, respectively, were not imaged due to limited movement of the microscope stage.

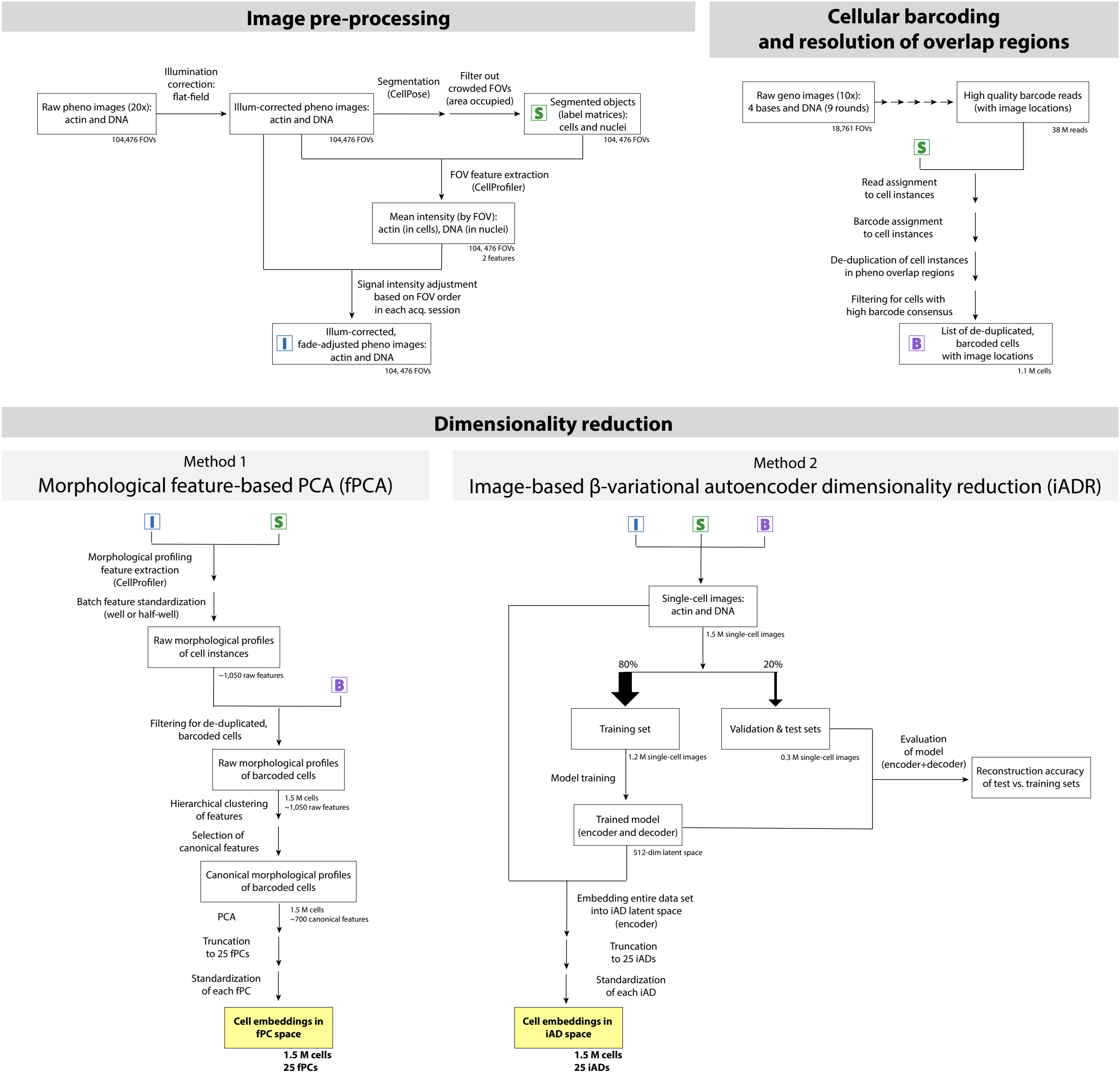


**Extended Data Fig. 4: Overview of the screen data processing and analysis pipeline.**

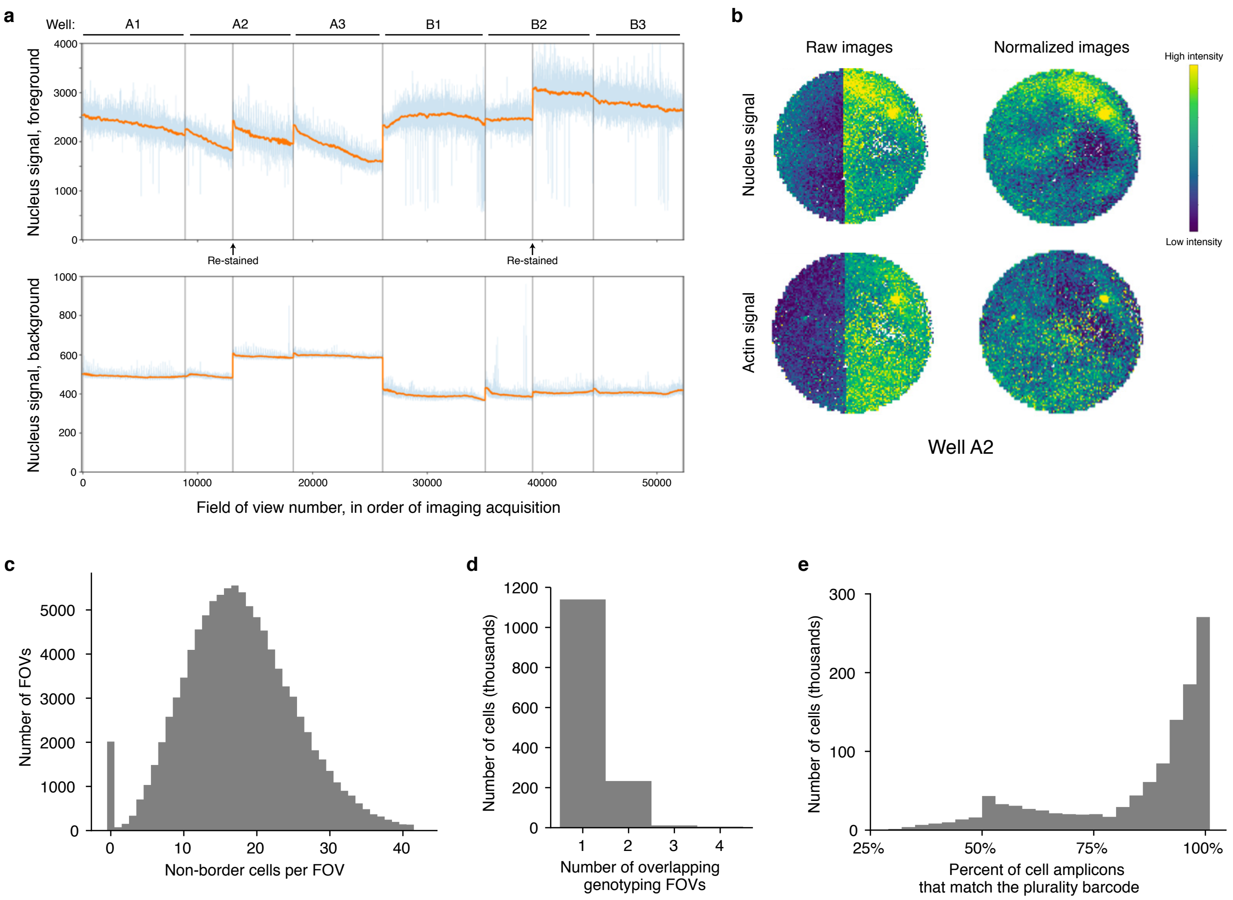


**Extended Data Fig. 5: Imaging and barcode-calling statistics for the screen.**
**a**, Plot of the average signal from the DNA stain inside cells ("foreground", above) and outside cells ("background", below) for each phenotyping FOV in a full 6-well plate. FOVs are displayed in the order in which their images were acquired on the microscope. Boundaries between different wells and two mid-well re-staining events are marked by vertical lines. Raw signal values are shown in light blue and a 200-FOV moving average is shown in orange (the moving average treats vertical lines as a boundary). **b**, Heatmap of the average intra-cellular signal intensity in the nucleus (above) and actin (below) channels for each phenotyping FOV before (left) and after (right) normalizing images based on the signal drift calculated from (**a**). Shown here is well A2, which had a mid-well re-staining event. **c**, Histogram of the number of non-border cells in each phenotyping FOVs. **d**, Histogram of the number of genotyping FOVs that each cell appeared fully within. **e**, Histogram of the percentage of amplicons in each cell that were the plurality barcode in the cell. The peak at 50% represents cells with two distinct and clearly-measurable barcodes.

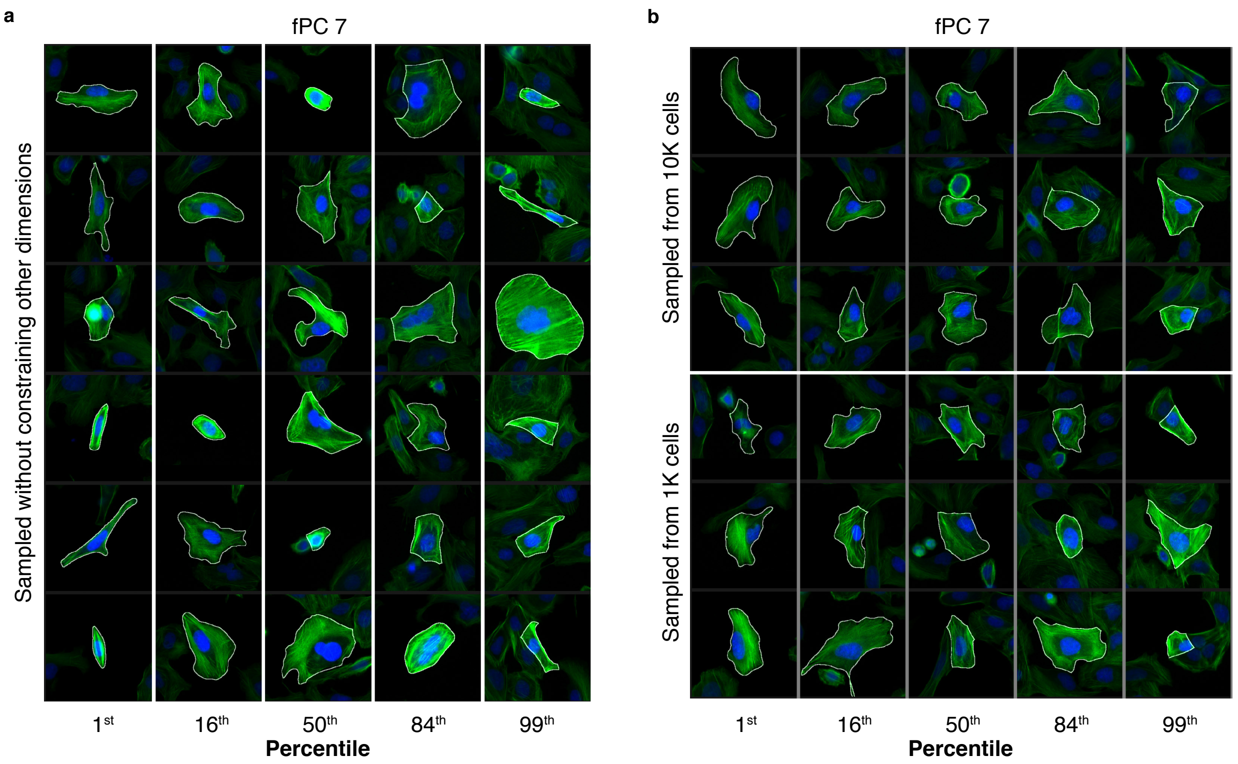


**Extended Data Fig. 6: Alternative ways of generating VIEWS.**
VIEWS of fPC 7, except (**a**) cells were sampled without minimizing all orthogonal dimensions or (**b**) cells were sampled from a random subset of 10K (above) or 1K (below) cells instead of the full dataset of >1M cells. Cells are displayed in the same manner as in Fig. 3; compare with Fig. 3f.

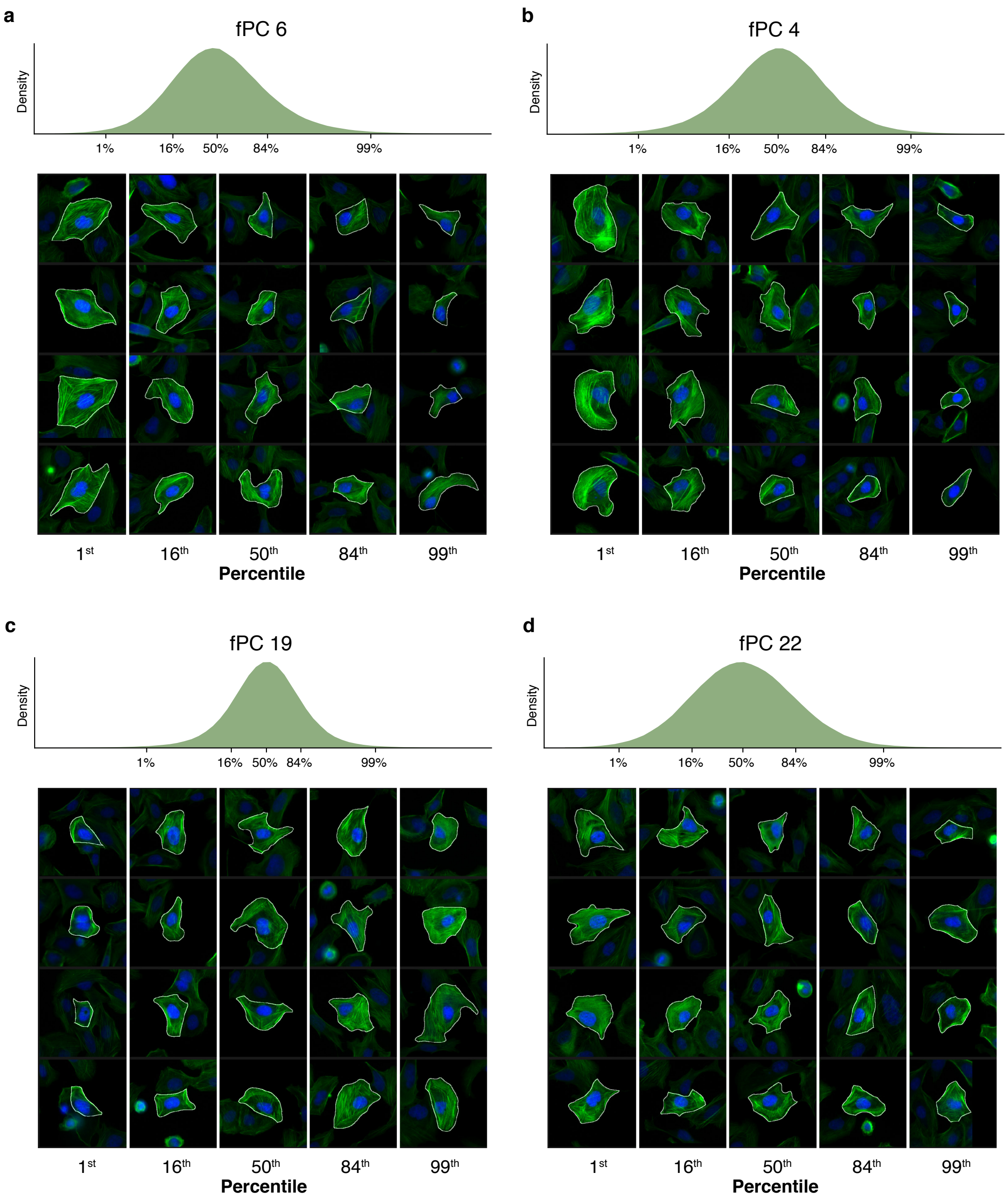


**Extended Data Fig. 7: VIEWS of additional fPCs.**

VIEWS of (**a**) fPC 6, (**b**) fPC 4, (**c**) fPC 19, and (**d**) fPC 22, displayed in the same manner as in Fig. 3.

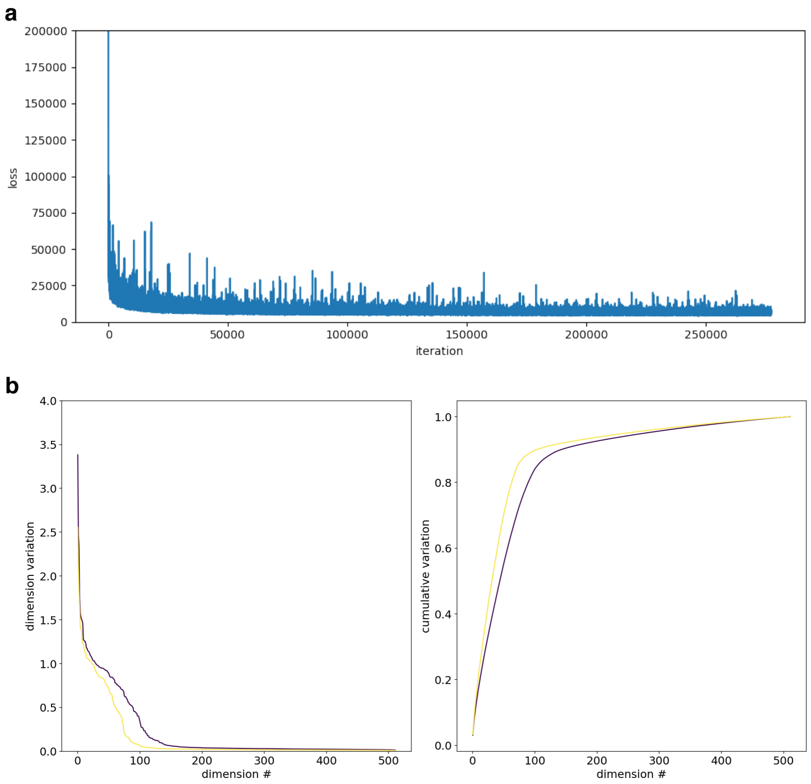


**Extended Data Fig. 8: β-VAE training.**

**a,** β-VAE loss over 270K iterations of training. **b**, For each of the 512 latent dimensions of the β-VAE, the variation of each dimension (left) or as a cumulative fraction of total variation (right) is plotted, with the dimensions in sorted order. Two models are shown, one with β = 0.5 (purple) and one with β = 0.99 (yellow). The latter was used for all analysis of knockdowns.

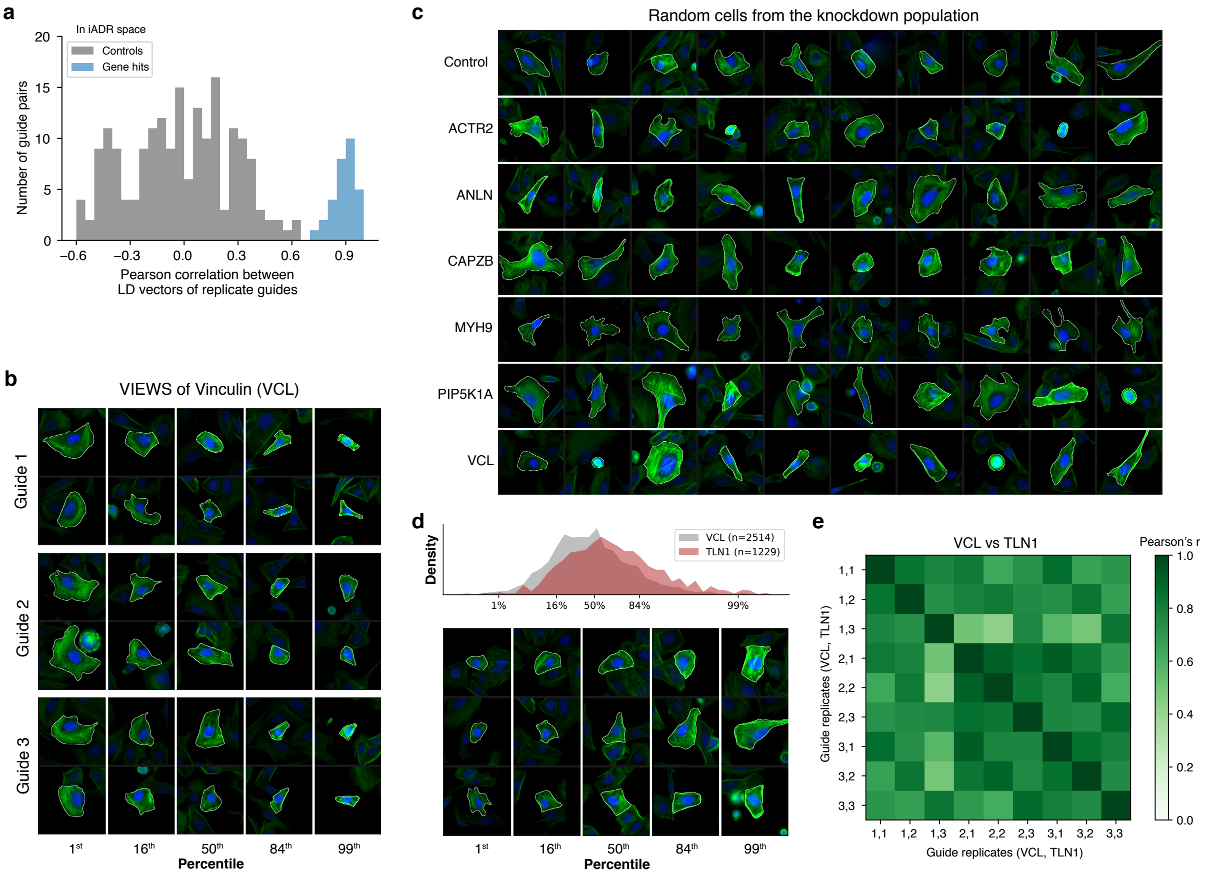


**Extended Data Fig. 9: Additional analysis of gene hits.**
**a**, Histogram of Pearson correlation values between LD vectors of two non-targeting control guides (gray) or between LD vectors of two replicate guide hits (blue). LD vectors are calculated in iADR space. Compare with Fig. 5b. **b**, VIEWS of three replicate guides targeting Vinculin (VCL). **c**, Ten cells randomly sampled from cells containing control guides or guides targeting one of six different genes. **d**, VIEWS of the LD vector describing the phenotypic shift from VCL knockdown cells (replicate 3) to TLN1 knockdown cells (replicate 3). Compare to Fig. 6c, which shows replicate 1 for each gene. **e**, Heatmap of pairwise Pearson's r values for all LD vectors describing the phenotypic shift from a VCL replicate and a TLN1 replicate. The labels "*x*,*y*" refer to VCL replicate *x* and TLN1 replicate *y*.

### Tables and Data

**Supplementary Table 1. DNA sequences, reagents, and equipment.**

**Supplementary Table 2. Pooled MIP optimization experiment results.**
Sequence and amplification efficiency for each MIP tested in the pooled optimization experiment. Number of amplicons ("count") and mean amplicon intensity ("signal") are provided in two replicate wells for each MIP. Related to Fig. 1d and Extended Data Fig. 2c,d.

**Supplementary Table 3. CellProfiler features.**
A list of all features calculated by CellProfiler for fPCA analysis. Features removed due to redundancy are marked with an asterisk. Related to Fig. 3 and Extended Data Fig. 4.

**Supplementary Table 4. Summary of screen results for each guide.**
For each guide, the sgRNA sequence, barcode sequence, number of cells measured, and statistical significance in each fPC and each iAD are provided. Statistical significance values are provided as $-\log_{10}$(p value). Also indicated is whether each guide was called as a hit by fPC or iAD and whether it was a replicated hit or a significant singleton verified using LD analysis. Related to Fig. 4 and 5.

**Supplementary Table 5. Screen data.**
For each cell, the associated guides and latent space embedding values for the top 25 fPCs and top 25 iADs are provided. Additional metadata is provided on the cell's position (plate, well, FOV, and coordinate within the FOV) and ISS statistics (number of total amplicons and fraction assigned to each guide). Cells associated with both a control guide and another guide were not used as control cells, so these control guides are marked here with an additional trailing underscore.

**Supplementary Data 1. VIEWS plots for each fPC 1-25 and each iAD 1-15.**

**Supplementary Data 2. VIEWS plots for each guide hit.**

### Supplemental Methods

#### CELL CULTURE

##### Cell culture

U2OS cells were obtained from ATCC (HTB-96, Lot# 70008732) and were cultured in McCoy's 5A modified media (Thermo, Gibco #16600-082) supplemented with 10% fetal bovine serum (EMD, TMS-013-B) at 37 C and 5% carbon dioxide. HEK293T cells were obtained from ATCC (ATCC, CRL-3216) and were cultured in DMEM media (Thermo, Gibco #11965-092) supplemented with 10% fetal bovine serum at 37 C and 5% carbon dioxide.

##### Construction of the CRISPRi U2OS cell line

U2OS cells expressing dCas9 were generated by lentiviral transduction with plasmid pHR-UCOE-Ef1a-dCas9-HA-2xNLS-XTEN80-KRAB-P2A-Bls (referred to as pHR-dCas9-XTEN-KRAB), which was a gift from Dr. Marco Jost and Dr. Jonathan Weissman, and modified by Dr. Sean Collins to include a blasticidin resistance marker for selection. The dCas9 transfer plasmid along with packaging plasmids psPAX2 (Addgene #12260, a gift from Didier Trono) and pMD2.G (Addgene #12259, a gift from Didier Trono) were miniprepped using the GeneJET plasmid miniprep kit (Thermo, K0503). For lentivirus production, 15 µg pHR-dCas9-XTEN-KRAB, 18.5 µg psPAX2, and 1.85 µg pMD2.G were diluted in 3.5 mL Opti-MEM I reduced-serum media (Thermo, Gibco #31985070) and then combined with 19 µL TransIT-Lenti Transfection Reagent (Mirus, MIR6600). Following a 10 minute incubation, this mixture was added dropwise to confluent HEK293T cells in a T175 flask containing 35 mL DMEM media supplemented with 1 mM sodium pyruvate (Thermo, Gibco #11360070). Lentivirus was recovered by collecting the media 48 hr later, with centrifugation at 500 g for 10 minutes to remove any residual cells and debris. Finally, the lentivirus was concentrated approximately 60-fold using Lenti-X Concentrator reagent (Takara Bio Inc., 631231). Here, the lentivirus was combined with 12 mL concentrator reagent and incubated overnight at 4 C. The mixture was then centrifuged for 45 min at 4 C and then following removal of the supernatant, the remaining pellet containing lentivirus was resuspended in 550 µL McCoy's 5A media. The concentrated lentivirus was snap-frozen in liquid nitrogen and stored at -80 C prior to use.

Wild-type U2OS cells were grown to approximately 40% confluence in a T25 flask in media supplemented with 8 µg/mL polybrene (Sigma, TR-1003-G) and infected with 300 µL of lentivirus. After 24 hours, cells were changed to selection media containing 10 µg/mL blasticidin (Sigma, SBR00022-1ML). Cells were selected for 7 days and then maintained in media supplemented with blasticidin for downstream use.

##### sgRNA cell line generation and knockdown

The library of knockdown cells expressing dCas9-XTEN-KRAB and a proximally-barcoded CROPseq-guide-proxBC plasmid (see below) were generated by low multiplicity of infection (MOI) lentiviral transduction following the CROP-seq protocol (Datlinger et al. 2017). HEK293T cells seeded in a 6-well plate in Opti-MEM with 200 µM sodium pyruvate and were transfected using Lipofectamine 3000 (Thermo, L3000001) with 0.9 µg each of the lentiviral packaging plasmids pMD2.G, pMDLg/pRRE (Addgene #12251, a gift from Didier Trono), and pRSV-Rev (Addgene #12253, a gift from Didier Trono)[59], and 1.7 µg of the pooled CROPseq-guide-proxBC plasmid library (per well). The supernatant was collected after 48 hours and passed through a 0.45 µm filter.

A T150 flask of dCas9-XTEN-KRAB-expressing U2OS cells in media supplemented with 8 µg/mL polybrene at approximately 50% confluence was infected with 400 µL of lentivirus. After 24 hours, cells were changed to selection media containing 2 µg/mL puromycin. Cells were selected for 7 days before plating for fixation, barcode amplification, and phenotyping and genotyping imaging.

U2OS cell lines (without dCas9-XTEN-KRAB) expressing one sgRNA or a smaller library of sgRNAs were generated similarly, but with scaled-down volumes.

#### CRISPR GUIDE CONSTRUCT

##### Creating a proximally-barcoded CRISPR sgRNA expression plasmid

We created an altered version of the CROPseq-guide-puro plasmid (Addgene #86708, a gift from Christoph Bock) (Datlinger et al., 2017) that removed the sgRNA scaffold and inserted a hybridization sequence for the MIP, which we call CROPseq-NOguide-puro. This was done using the Q5 Site-Directed Mutagenesis Kit (NEB, E0554). The plasmid sequence outside of the sgRNA scaffold was amplified using Phusion HotStart Master Mix (2x) (NEB, M0531L) and primers CROPseq-NOguide-PCR-F and CROPseq-NOguide-PCR-R. The PCR product was KLD treated following the NEB kit protocol, and transformed into NEBstable competent cells (NEB, C3040), which were plated on LB agar plates containing 100 µg/µL carbenicillin and grown overnight at 30 C. Plasmids were purified from picked colony cultures, miniprepped, and the insert was verified with Sanger sequencing.

To create a proximally-barcoded sgRNA expression plasmid, CROPseq-guide-proxBC, an insert was designed that contained the sgRNA sequence (targeting sequence and backbone) and associated barcode region, flanked by PCR primers. Inserts were synthesized by IDT as gBlocks and inserted into CROPseq-NOguide-puro using the NEB BsmBI-v2 Golden Gate assembly kit (NEB, E1602). 10 fmol of pooled insert and 20 fmol of plasmid was used in a 20 µL Golden Gate reaction with incubation at 42 C for 1 hour and 60 C for 5 minutes, which was then transformed into NEBStable competent cells.

To construct a library of sgRNA plasmids for pooled MIP experiments or for the full-scale screen, inserts were designed in the same manner, then synthesized by IDT as an oPool (Screen_Insert_Library). The oligo pool was converted to double-stranded DNA by hybridizing the Screen_Elongation-R primer to the 3' end and extending the primer by incubating with Phusion polymerase master mix at 98 C for 2 minutes, 65C for 30 seconds, and 72 C for 5 minutes. The double stranded pool was then purified in a column cleanup (QIAGEN, 28104) and inserted into CROPseq-NOguide-puro as above. An 8 µL aliquot was transformed into 200 µL NEBStable competent cells and grown overnight in LB media with carbenicillin at 30 C to minimize homologous recombination. Plasmids were midiprepped from 50 mL culture and sequence-verified with next generation sequencing to check representation of each guide and correct association between guides and barcodes. Note that attempts to clone these libraries using Gibson assembly (NEB, EE5510) produced a high rate of molecules with incorrect guide-barcode pairs, possibly due to template-switching of the Gibson assembly polymerase within the sgRNA scaffold, which has significant secondary structure.

**Measuring barcode swapping by digestion-qPCR assay**

To measure the level of swapping between guide-barcode pairs during lentiviral infection of the proximally-barcoded CROPseq-guide-proxBC plasmid, we used two guide sequences targeting vimentin (VIM) and Ras-related protein Rab-1 (RAB18), each associated with a unique 8 nt barcode. Lentivirus was generated by transfecting HEK293T cells with VIM plasmid, RAB18 plasmid, or an equal mixture of both (two replicates), and corresponding U2OS cell lines were generated, as described above. A fifth condition was made by mixing together the VIM-only and RAB18-only cell lines in equal cell counts right before genomic DNA (gDNA) extraction.

gDNA was harvested from 1M cells from each condition using the PureLink Genomic DNA mini kit (K182001) and eluted in 50 µL. gDNA was digested in triplicate 30 µL reactions overnight for 37 C with a combination of NheI-HF (NEB, R3131) and BmtI-HF (NEB, R3658) (these cut in the RAB18 barcode), for 1 hour at 22 C and overnight at 37 C with a combination of SfcI (NEB, R0561) and Hpy188III (NEB, R0622) (these cut in the VIM barcode), or a mock digestion with no enzyme (Extended Data Fig. 2a). qPCR measurements were performed on an ABI 7900HT system using SYBR FAST ABI qPCR mix (KAPA, KK4605) in triplicate reactions with annealing at 66C. One primer pair specific to each guide sequence that amplifies across the digestion site was used (CROP-RAB18-qPCR-F + CROP-sgRNA-qPCR-R and CROP-VIM-qPCR-F + CROP-sgRNA-qPCR-R). The total barcode-guide swapping rate is expected to equal twice the percentage of barcode DNA left undigested (as detected by this qPCR) since barcode-guide swap events in lentiviral particles with two VIM copies or two RAB18 copies would go undetected.

**Measuring barcode swapping by Illumina sequencing**

For measuring barcode swapping during plasmid library cloning, the region containing the sgRNA and barcode was amplified from 10 ng of BsmBI-digested plasmid library in a 100 µL PCR reaction with Phusion polymerase for 10 cycles with primers CROP-sgBC-PCRf-R1p and CROP-sgBC-PCRr-R2p at a 61 C annealing temperature. For measuring barcode swapping after lentivirus transduction, gDNA was extracted as described above, and the same PCR reaction was performed with 50 ng of input gDNA. After a column clean-up, a second PCR was performed with 5 ng of template to add Illumina sequencing adapter and multiplexing barcode sequences. Samples underwent 1.0x size selection with Agencourt AMPure XP beads (Beckman, A63880) and 2 x 80 nt paired-end sequencing on an Illumina NextSeq.

Reads were trimmed and aligned to the expected sequences between the sgRNA targeting sequence and the barcode using bwa-mem, and the previous 20 bp and the subsequent 9 bp were taken to be the sgRNA targeting sequence and the barcode, respectively. Reads that did not perfectly match either a sgRNA sequence or a barcode were excluded. Barcode swapping was calculated as the percentage of remaining reads that matched a sgRNA sequence and a barcode that were not designed to be associated. For the final pooled screen, the plasmid library and gDNA-integrated construct had barcode swapping rates of 0.41% and 7.6%, respectively.

#### SCREENING METHODS

##### Amplification of the barcode sequence inside fixed cells

For all experiments involving fluorescence microscopy, cells were seeded in glass-bottomed plates (Cellvis P06-1.5H-N and P96-1.5H-N). Before seeding, plates were incubated in 10 µg/mL fibronectin (Thermo, 33016015) in PBS (Thermo, AM9624) for 90 minutes, then rinsed with PBS and McCoy's 5A media with 10% fetal bovine serum.

For 6-well plates, all washes were performed in a volume of 2 mL per well, and all molecular biology reactions were performed in a volume of 500 µL per well. Volumes were scaled down proportionally for plates with smaller wells. To minimize evaporation during overnight incubations, any adjacent wells were filled with water and plates were placed in a sealed humidified container. For shorter incubations that required elevated temperatures, a microplate incubator with a heated lid (BT Lab Systems, BT1101) was used to minimize evaporation.

Cells were fixed with 3.6% formaldehyde (Thermo, 28906) in PBS, washed twice with RNase-free PBS plus 0.05% Tween (PBST), dehydrated with 70% ethanol on ice for 30 minutes, washed with PBS, and permeabilized with PBST. When removing ethanol, 80% of the volume was first exchanged with PBST to avoid drying out the cells before a full volume exchange was performed. A reverse transcription (RT) primer (CROP-RT2-LNA-5Am), which was modified with locked nucleic acid (LNA) bases and a 5' amino group, was hybridized at 1 µM in PBST for 30 minutes at room temperature. After primer hybridization but before RT, cells were fixed again with 3% formaldehyde and 0.1% glutaraldehyde (Sigma, G5882-10x1ML) in PBST for 30 minutes at room temperature. The reaction was briefly quenched with 1M Tris-HCl (pH 8), and cells were washed twice with PBST and once with 1x RevertAid RT buffer. The RT reaction – consisting of 5 U/µL RevertAid H-Minus reverse transcriptase (Fisher, EP0452) in 1x RevertAid RT buffer, 1 µM RT primer, 0.2 mg/mL bovine serum albumin (BSA; NEB, B9000), 250 µM dNTPs (Thermo, 18427088), and 1U/µL RNaseOUT RNase inhibitor (Thermo, 10777-019) – was run overnight at 37 C.

After RT, cells were washed 3 times in PBST, fixed again with 3% formaldehyde and 0.1% glutaraldehyde in PBST for 30 minutes at room temperature, quenched briefly with 1M Tris-HCl, and washed twice with PBST. Wells were washed with 1x AmpLigase buffer and then gap-filling was performed with 10 nM molecular inversion probe (MIP) BestMIP1_L81+8_A1R4 for 5 minutes at 37 C and 90 minutes at 45 C, with TaqIT polymerase (Enzymatics, P7620L) at 0.002 U/µL, RNase H (NEB, M0297S) at 0.4 U/µL, BSA at 0.2 mg/mL, dNTPs at 50 nM, and AmpLigase at 0.5 U/µL (Lucigen, A32750) in 1x AmpLigase buffer. After gap-filling, the cells were washed twice with PBST and once with 1x phi29 buffer. The rolling circle amplification reaction – consisting of 1 U/µL phi29 polymerase (NEB, M0269L), 250 µM dNTPs, 0.2 mg/mL BSA, and 5% glycerol, in 1x phi29 buffer – was performed overnight at 30 C.

##### *In situ* sequencing (ISS) of barcode DNA

The sequencing primer ISSeq-BestMIP1 was hybridized to amplicons at 1 uM in 2x saline-sodium citrate (SSC) buffer (Thermo, 15557044) with 10% formamide (Sigma, F9037-100ML) for 30 minutes at room temperature before the first round of sequencing. The cells were washed with 500 µL of Illumina Nano Kit (MS-103-1003) incorporation buffer (PR2 reagent), incubated in Illumina Nano Kit incorporation mix for 3 minutes at 60 C, and washed thoroughly with 500 µL of PR2 6 times for 3 minutes each at 60 C. Cell nuclei were stained with 1 µg/mL Hoechst 33342 (Thermo, H3570) for 15 minutes at room temperature and washed for 5 minutes in 2x SSC buffer, then placed in 1 mL 2x SSC buffer for fluorescent imaging. After each round of imaging, the fluorophores and terminators were cleaved with Illumina Nano Kit cleavage buffer for 6 minutes at 60 C and thoroughly washed again with PR2 6 times for 3 minutes each before the next incorporation round.

**Microscopy**

Sequencing images were acquired on a Nikon Ti-E Eclipse microscope using a CFI Plan Apo Lambda 10x/0.45NA objective (Nikon) or a Plan Apo 20x/0.75NA objective (Nikon) and illuminated by a Lambda XL lamp (Sutter). Bases were imaged with the following filters: 534/20 excitation, 572/28 emission, and 552 dichroic for base G; 555/25 excitation, 605/52 emission for base T; 635/18 excitation, 680/42 emission, and 635 dichroic for base A; and 650/60 excitation, 775/140 emission, and 700 dichroic for base C; all filters were obtained from Semrock. Nuclei were stained with Hoechst 33342 and imaged using ET490/20x excitation and ET525/36m emission filters (Chroma). Actin was stained with Alexa Fluor 488 phalloidin and imaged using D350/50x excitation and ET455/50m emission filters (Chroma). Micromanager software was used for automated microscope control.

**Testing of 113 MIP designs in a pooled experiment**

This experiment followed the method described in (Feldman et al., 2019). To minimize secondary structure, backbone sequences were randomly generated and selected to contain only A, C, and T nucleotides with 45-55% Cs, and without homopolymers 4 nt or longer. For each of three different 5' hybridization sites and three different backbone lengths, many backbone sequences were generated, the sequences were run through the secondary structure predictor Zipfold[60], and eight sequences with the highest minimum free energy were selected, for a total of 72 sequences. Another 41 sequences were designed with a fixed backbone sequence but with total amplicon length varying from 80 nt to 120 nt. A unique 5 nt barcode was designed into the backbone region of each MIP probe to allow them to be distinguished by *in situ* sequencing. All designs are listed in Supplemental Table 2.

Oligos were synthesized from IDT as an oPool. A naive probe design (MIP-BC-AE9) used in previous tests was also separately added to the pool. A cell line of U2OS cells was created as described above with 100 unique sgRNAs with associated 8 nt barcode regions (CROPseq-guide-proxBC) integrated into genomic DNA. These cells were plated in replicate wells of a 96-well plate and amplified *in situ* as described above, using the pool of MIPs. Amplicon count and intensity were determined by hybridizing an Alexa Fluor 647-labeled DNA probe (PoolMIP-AF647-probe) to a constant region on the probe backbone; intensity values were corrected for differences in illumination across the field of view (FOV). The identity of the MIP corresponding to each amplicon was then determined by 5 rounds of *in situ* sequencing, as described above, using a mix of four primers (PoolMIP-ISSeq-A{1,2,3} and ISSeq-BC-AE9). Each entire well was imaged.

**Side-by-side comparison of different barcode amplification protocols**

The U2OS cell line expressing 100 unique sgRNAs and associated 8 nt barcodes was plated in a 96-well plate, and five different conditions for barcode amplification were run side-by-side in two replicate wells each. Conditions 1-3 followed our protocol above and used one of two different MIPs, which amplify the 8 nt barcode, and associated ISS primers: our first probe MIP-BC_AE9 with ISS primer ISSeq-BC-AE9, or our optimized probe BestMIP1_L81+8_A1R4 with ISS primer ISSeq-BestMIP1. These conditions also either included or skipped the pre-RT primer hybridization, fixation, and quench steps. Two conditions were performed exactly as described in (Feldman et al., 2019) using the RT primer oRT_CROPseq, the MIP oPD_CROPseq, which amplifies the 20 nt guide targeting region, and ISS primer oSBS_CROPseq. In one condition, the pre-RT primer hybridization, fixation, and quench steps from our protocol were added, and to allow fixation, an RT primer with an added 5' amino group was used. Specifically, conditions were as follows:

1. MIP-BC_AE9 probe with pre-RT fixation
2. BestMIP1_L81+8_A1R4 probe without pre-RT fixation
3. BestMIP1_L81+8_A1R4 probe with pre-RT fixation
4. Feldman et al. protocol (no pre-RT fixation)
5. Feldman et al. protocol with additional pre-RT fixation

A single round of ISS was performed and the four bases and cell nuclei were imaged as described above. Around 10,000 cells in 137 FOVs were imaged for each condition and each replicate. Signal from the four bases were used to count amplicons and quantify their signal intensity, all using Fiji. Specifically, amplicon locations were determined using the "Find maxima" function with Prominence values of 400, 350, 125, and 150 for bases G, T, A, and C respectively. Sequencing signal intensity was measured as the value of the pixel of the amplicon center minus the background, which was calculated as the median pixel value across the image in that channel. The number of cells was counted by segmenting nuclei. Specifically, images were subjected to a one-pixel Gaussian blur, thresholding with default parameters, filling holes, and watershed, and finally nuclei were counted using the "Analyze particles" function with a lower size limit of 500 pixels (approximately 320 $\mu m^2$). In addition to results reported in the main text, adding a pre-RT fixation to the Feldman et al. method increased amplicon number by 1.8-fold (conditions 4 versus 5).

#### SCREEN – EXPERIMENT

**Seeding experiment tests**

Tests were performed to determine a reproducible method of obtaining even seeding of cells at the correct density. U2OS cells were seeded at a variety of densities in glass-bottom 6-well plates (coated with fibronectin as described above), actin and nuclei were stained as described below, and cells imaged at 20x magnification in FOVs evenly spaced across each well, all as described above. CellProfiler was used to segment and count cells and compute the percentage of each cell's boundary that was touching another cell (Extended Data Fig. 3a,b).

Seeding cells using low volumes of media gave very uneven distributions across the well because the curvature of the meniscus caused more cells to fall along the periphery of the well

(Extended Data Fig. 3c). Best results were found using large volumes (~5 mL for a 6-well plate). However, slight movements of the liquid or plate after addition of the media caused cells to collect towards the center, similar to how tea leaf fragments settle in the center of the cup after swirling the liquid (see the "tea leaf paradox"). To avoid these issues, cells were added slowly and dropwise to each well, the plate was left to sit for 30 min after seeding to allow cells to settle and begin attaching to the surface, and the plate was moved very gently to the incubator.

**Choice of gene targets, guide sequences, and barcodes**

The gene target library was designed as follows. First, human genes not expressed in U2OS were filtered out based on RNA sequencing data from the Human Protein Atlas (https://v19.proteinatlas.org/about/download)[61] . The second filtering step was based on GO term annotations related to the actin cytoskeleton, including terms such as “stress fiber assembly”, “actomyosin”, and “regulation of cell shape” [62,63]. Specifically, if a gene was annotated with the “actin-binding” term or at least two of the other selected GO terms, then the gene passed this filtering step. Lastly, manual inclusion or exclusion on an individual gene basis was executed, based on a broad literature search and considering expression level in U2OS. The end result was a library of 366 genes enriched for those with known or hypothesized involvement in actin-related cellular processes. Note that talin rod domain containing 1 (TLNRD1, Fig. 5 and 6) is also known as mesoderm development candidate 1 (MESDC1).

Triplicate CRISPRi guide RNA sequences for each of these genes were taken from the DolcettoA set of validated designs[31], for a total of 1098 guides. An additional 20 control guides were also randomly selected from this set. A set of 1118 barcodes 9 bp long were randomly sampled and constrained to have either four or five total Gs and Cs and to have a minimum edit distance of 3 between any two barcodes.

**The optical pooled screen**

A U2OS cell line expressing dCas9-XTEN-KRAB was made fresh from wild-type cells and selected for seven days as described above. A library of 1118 barcoded sgRNA expression plasmids were cloned and lentivirus was made as described above. Cells were cultured for eight days after transducing the dCas9-expressing cells with sgRNA expression lentivirus to allow integration and full knockdown to be achieved. Cells were plated in two full 6-well plates (fibronectin-coated as described above) at a density of 100,000 cells/well, fixed 13 hours after plating, and then barcodes were amplified *in situ*, as described above.

After completion of rolling circle amplification, fixed cells were stained with 1 µg/mL Hoechst 33342 and 165 nM Alexa Fluor 488 phalloidin in 1x PBST for 20 min at room temperature and washed two times in 1x PBST, and imaged at 20x magnification as described above. Because both stains dissociated over the many hours required for imaging, staining and imaging was done in batches; specifically, two wells were stained and one and a half wells were imaged continuously in each batch. By re-staining and resuming imaging in the middle of a well, we were able to verify that intensity normalization performed in post-processing completely adjusted for the change in intensity over the course of imaging time.

After phenotyping imaging, nuclear stain was removed by two washes of 70% ethanol for 5 minutes each and the actin stain was removed during the 60 C washes required for ISS.

Nine rounds of ISS were performed and each plate was imaged at 10x magnification in one continuous session. Phenotyping and genotyping images were taken with 22% and 10% overlap between adjacent FOVs, respectively, in order to maximize the number of cells completely contained within at least one FOV. Duplicate cells—cells completely contained within more than one field of view—were resolved computationally in downstream processing steps (see below).

#### SCREEN – DATA PROCESSING

##### Genotype image processing, amplicon detection, and base calling

Each genotyping image was corrected for uneven illumination by averaging together >10,000 images and dividing each channel by the averaged image. For each genotyping FOV, images from each sequencing round were aligned to the first round using the nucleus channel, then cropped to leave only pixels that overlapped all sequencing rounds. For each FOV, round, and sequencing channel, pixel intensities were normalized by dividing by the 95th percentile value, a Laplacian of Gaussian (LoG) transformation was performed, and a maximum filter in a 3 x 3 grid was performed.

Pixels corresponding to amplicons are the pixels that vary the most between channels and sequencing rounds, so to call amplicons, for each pixel, the standard deviation was calculated across all four base channels and all nine sequencing rounds. An initial amplicon position list was compiled by taking the pixels with standard deviation within 0.19 and 0.45 and keeping only the pixels whose standard deviation matched the maximum standard deviation in a 5 x 5 grid of pixels around it. This thresholding method has advantages over only finding local intensity maxima, because it avoids detecting false positives in autofluorescent regions with high intensity in all channels, but with low variance.

Due to small errors in nucleus-based image alignment, chromatic aberration, and other systematic imaging aberrations, the called amplicon positions were often systematically a few pixels shifted from the maximum intensity pixel in some images and some channels. Thus, the amplicon signal itself was used to refine the image alignments to obtain single-pixel resolution. For each FOV and round, a 9 x 9 area around each amplicon was averaged, producing one grid per channel that showed the aggregate signal relative to the amplicon position. Then each channel within each round was shifted by a few pixels to place the highest intensity pixel (in this 9 x 9 area) at the center. Once again, amplicons were called and alignment was refined, and this was repeated for a total of three times. The final list of amplicons was called with standard deviation thresholds widened to 0.07 and 0.45.

For each amplicon and each sequencing round, the identity of the nucleotide was determined based on the maximum intensity in a 3 x 3 grid around the amplicon center in each of the four base channels. A linear classifier was fit to intensity data from eight sequencing rounds from a cell line expressing a single known barcode, then used to call nucleotide identities in the screen. To adjust for systematic changes in sequencing signal and background signal that occurred between rounds, new linear classifiers were trained per-round on the screen data itself. Specifically, a subset of amplicons whose called sequences matched a designed barcode with 0 or 1 errors were used as training data, with the error-corrected barcodes as labels. These per round classifiers were then used for a final iteration of base calling over all of the amplicons, with each classifier used to predict the bases from its corresponding sequencing round. This increased the per-round accuracy on a held-out test set of amplicons by ~1.5% (based on edit distance to designed barcodes).

**Cell segmentation and deduplication**

Each phenotyping image was corrected for uneven illumination by averaging together >10,000 images and dividing each channel by the averaged image. Cells and nuclei were each segmented using the "cyto2" model of CellPose version 0.6.5[35]. Cells were segmented with object diameter of 120 and flow threshold of 0.6, while nuclei were segmented with object diameter of 44, default flow threshold, and cellprob_threshold of 1.0. Best results were obtained when the nucleus channel was background-subtracted (25th percentile pixel value) and gamma corrected with exponent 1/3 prior to segmentation.

Cells and nuclei containing fewer than 225 or 64 pixels, respectively, were removed. Each nucleus was associated with a cell if it contained at least 90% of the nucleus, and nuclei were masked to be entirely contained within their associated cell. Nuclei that were not associated with any cells were removed. Some cells were associated with multiple nuclei, so the number of nuclei associated with each cell was counted and kept as a feature, and then the nuclei of each cell were merged into one segmented object. Label matrices of cells and associated nuclei were constructed and used for feature extraction in CellProfiler. Cells without nuclei were removed. Finally, cells touching the border of the image (and their associated nuclei) were removed.

Adjacent phenotyping FOVs overlapped by 22%, so some cells were fully contained within multiple FOVs. To associate duplicate images of cells with each other, each FOV was aligned with the FOVs occurring directly below it and to the right based the nucleus channel, with each image pre-cropped to the expected overlap region based on the microscope stage position list. Two cell instances, one from each image, were considered duplicates of each other if their areas overlapped by at least 50%. In practice, duplicate cell instances were very similar and often overlapped by >90%. Duplicates were resolved after assigning amplicons to cells.

**Correcting fluorescence changes over imaging time**

Due to differences in staining between wells and steady dissociation of stains during imaging, intensity of the nucleus and actin signal were shifted in systematic ways between FOVs and wells (Extended Data Fig. 5a). To measure and account for this shift, cells were segmented in each FOV, and the mean nucleus and actin intensity in each FOV were calculated for the foreground (within cells) and background (outside of cells). The FOVs were sorted based on the order of imaging and a rolling average of signal intensity was calculated across groups of 200 consecutively-imaged FOVs. Finally, the intensity of original images was normalized by the rolling average value so that all images had approximately the same mean intensity as the first well, and cells were re-segmented from normalized images.

**Assigning barcodes and consensus guides to cells**

Each phenotyping image (acquired at 20x magnification) was aligned to at least one, and up to four, genotyping images (acquired at 10x magnification), due to overlap of genotyping images. Alignment was done using the nucleus channel with the phenotyping image shrunk by two-fold in each direction, and with both images pre-cropped to the expected overlap region based on microscope XY-coordinates of the FOVs. Each cell instance in the phenotyping image was then assigned all the amplicons that fell within its cell boundaries from each aligned genotyping image. Finally, if two cell instances were duplicates of each other (see above), the instance that was assigned the most error-correctable amplicons was chosen as the canonical instance and used for all downstream analysis.

Note that some amplicons were captured in two genotyping images, and these were not deduplicated due to the difficulty of computing pixel-perfect alignments. However, fewer than 20% of cells overlapped more than one genotyping FOV (Extended Data Fig. 5d), so the double-counting was not expected to be impactful. To avoid counting duplicate amplicons, the histogram of ISS reads per cell (Fig. 2e) was calculated from a subset of cells that only overlapped one genotyping FOV.

A guide was considered present in a cell if the cell contained at least 5 error-corrected amplicons that matched the guide's barcode and if this represented at least 30% of all error-correctable amplicons in the cell. Each cell was assigned up to two guides satisfying these criteria; cells that were not assigned any consensus guide were omitted from all downstream analysis.

#### SCREEN – FEATURE EXTRACTION

##### Selection and calculation of CellProfiler features

Features were calculated using CellProfiler version 3.1.5[36]. Direct inputs to the CellProfiler pipeline included actin and nucleus raw images, actin and nucleus illumination correction functions (calculated from a subset of the screen data), and cell and nucleus segmentations (generated using CellPose as explained above).

Orientation-specific features were averaged across different orientations to collapse into single rotationally-invariant features. A total of 1152 features were calculated. First, 98 features were omitted because they involved DNA signal in the cytoplasm region. Features were then clustered by Ward's method of hierarchical clustering using a Pearson's r cutoff of 0.95, and a single representative feature was chosen from each cluster based roughly on interpretability of the feature. Specifically, cell features were preferred over nucleus features, which were preferred over cytoplasm features. Similarly, feature categories followed this priority order: AreaShape, Intensity, Neighbors, RadialDistribution, Granularity, Texture, Correlation, AreaOccupied, and Count. This process removed 346 highly-redundant features. Afterwards, there were 305, 336, and 67 features describing the cell, nucleus, and cytoplasm respectively; and 109, 9, 38, 55, 13, 345, and 139 features describing AreaShape, Correlation, Granularity, Intensity, Neighbors, RadialDistribution, and Texture respectively. All features are listed in Supplementary Table 3.

##### Filtering, normalization, and PCA on CellProfiler features

To remove FOVs containing debris or other imaging aberrations, phenotyping FOVs with greater than 99.5th percentile average mean intensity in the DNA or actin channels were removed from downstream analysis. Furthermore, to avoid phenotypic changes due to cell crowding, FOVs with greater than 99.5th percentile cell numbers (42 cells or more) were removed from downstream analysis. In total, these excluded 1324 phenotyping FOVs out of 101,903 total (1.3%).

Because staining and phenotype imaging was done in batches corresponding to 1.5 wells, different wells and plates varied slightly in their mean intensity, particularly for the DNA stain. To correct for this, DNA stain intensity-dependent CellProfiler features (namely, the Intensity and Texture features) were normalized within each batch (i.e. full well or half well) to have the same mean and standard deviation.

PCA was performed on all cells and all CellProfiler features remaining after the filtering described above. Each feature was standardized to have mean of 0 and variance of 1 prior to PCA, and the top 100 PCs were recorded. Cells with highly extreme values (greater than six standard deviations from the mean) in any of the top 30 PCs were excluded from downstream analysis.

**Training the β-variational autoencoder (β-VAE)**

The β-VAE is employed as an alternative approach to disentangle morphological variations of cell and nuclear shapes as well as the distribution of actin. The β-VAE is based on the variational autoencoder (VAE) architecture[64,65], which maps images into a multidimensional latent space via the deconstruction and reconstruction of the original image using an encoder and a decoder. The β-VAE introduces a hyperparameter, β, that modulates the balance between different learning constraints applied to the model[37].

In this application, the β hyperparameter modulates the trade-off between the capacity of the latent space used (sparsity) versus image reconstruction fidelity[20]. Specifically, the model is trained to optimize the Evidence Lower Bound (ELBO) given an input image, which comprises of two components:

$$\mathrm{ELBO}(x) = (1-\beta)\mathbb{E}_{q(z|x)}[\log p(x|z)] - \beta KL(q(z|x)|p(z)).$$

with β ranging between 0 and 1. The term on the left represents the reconstruction error between the input and reconstructed image, while the term on the right represents the KL-divergence between the embeddings produced by the encoder (q(z|x) and the Gaussian prior (p(z)).

A single 501 x 501 pixel image (198 x 198 µm) image centered around the cell centroid was created for each of the 1,514,673 cells. This size was chosen so that large and elongated cells would not be cropped out, but to speed training time, the images were rescaled to 320 x 320 pixels. The area outside of the nucleus segmentation was masked in the nucleus channel and the area outside of the cell segmentation was masked in the actin channel. Pixel intensities of the actin and nucleus channels were capped at 6000 and 7500 respectively and rescaled to 8-bit images; these thresholds were chosen so that less than 10% of cells contained a pixel exceeding the maximum. Cell images were split into 80% training, 10% validation and 10% test sets. The model used is trained with a β value of 0.99 and a batch size of 32, with a learning rate of 0.0002, for 10 epochs. The Adam optimizer is used for gradient-descent. The maximum latent space dimensionality was set to 512 dimensions. After training, latent space embedding vectors were each cell, the latent dimensions were sorted in order of decreasing variance, and only the top 25 dimensions were used for analyses.

#### SCREEN – HIT CALLING, ANALYSIS, AND VISUALIZATION

##### Calling guide and gene hits

To determine if a guide of interest caused a statistically significant shift along a single given axis (such as a fPC or iAD), a two-sample Kolmogorov-Smirnov test was performed comparing all cells containing the guide of interest with all cells containing any of the 20 control guides. If $p < 10^{-6}$, the guide was considered a statistically significant hit. To ensure consistency of the control cells, cells that contained two distinct guides were excluded from the set. If the guide of interest was a control guide (such as in the bottom five rows of Fig. 4c), cells containing that control guide were also excluded from the control set. A gene was considered a hit if at least two of the three replicate guides targeting that gene were hits.

##### VIEWS

VIEWS plots were calculated given an embedding space (e.g. the first 25 fPCs or the first 25 iADs), a vector within the embedding space (e.g. the 5th fPC or the linear discriminant vector that best separates VIM knockdown cells from controls), a population of cells (e.g. the entire dataset or the VIM knockdown cells), and a percentile value *k* to highlight (e.g. 1st, 16th, 50th, 84th, or 99th). First, cells in the chosen population were mapped to the embedding space and projected onto the chosen vector, yielding one value per cell (i.e. the signed distance from the origin along the vector), and values $v_{low}$ and $v_{high}$ were calculated corresponding to the ($k$ - 0.5)-th and (k + 0.5)-th percentile of this distribution. Second, cells from the entire dataset were similarly projected onto the chosen vector, and all cells falling between $v_{low}$ and $v_{high}$ were selected. Finally, for each of these selected cells, the Euclidean distance in the embedding space from the cell to the vector was calculated, and the cells with the lowest distances were displayed. For example, when the embedding space is the first 25 fPCs and the chosen vector is the 5th fPC, this procedure picks out cells with 5th fPC value between $v_{low}$ and $v_{high}$ and all other fPC values close to 0.

##### Linear discriminant (LD) analysis

LD vectors were calculated in an embedding space consisting of the first 25 fPCs or the first 25 iADs, standardized so that each dimension had mean equal to zero and variance equal to one, using the LinearDiscriminantAnalysis function in scikit-learn version 0.20.1 in python. Similar to hit calling, LD vectors were computed for each guide relative to all control guides combined. LD vectors were rescaled to have unit length and to point in the direction of the knockdown population.

##### Cross-correlation and clustering

Agglomerative hierarchical clustering in Figure 6 was performed using a distance metric consisting of Pearson's correlation and Ward's method to combine clusters. For each gene, a single representative LD vector was calculated by averaging the LD vectors of all guide hits of the gene, and Pearson's correlation was calculated between all pairs of gene LD vectors. These analyses were performed using LD vectors calculated in fPCA space.
