## Supplementary figures and images for "Mapping variation in the morphological landscape of human cells with optical pooled CRISPRi screening"

### VIEWS_fPC1.pdf

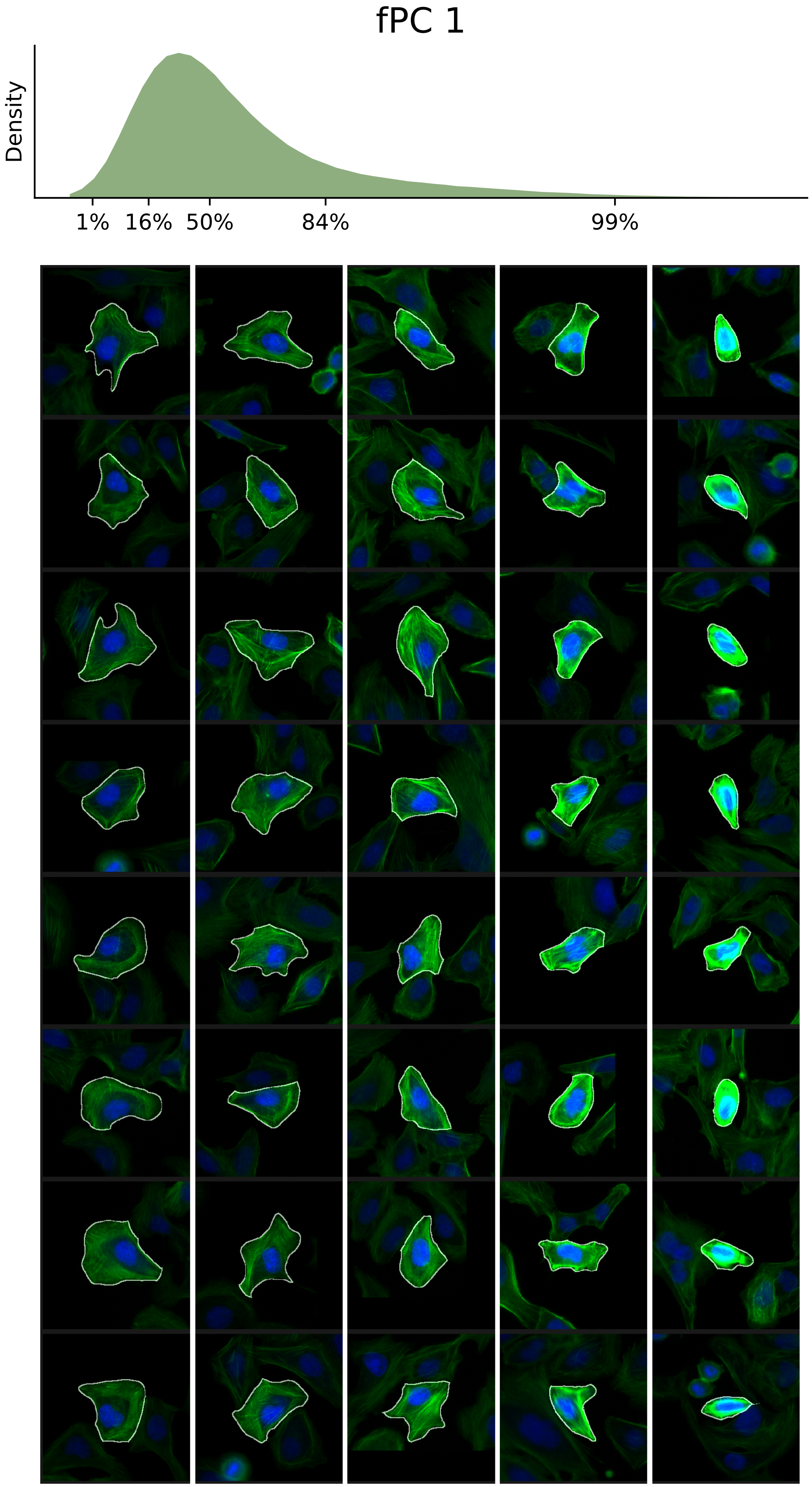

### VIEWS_fPC2.pdf

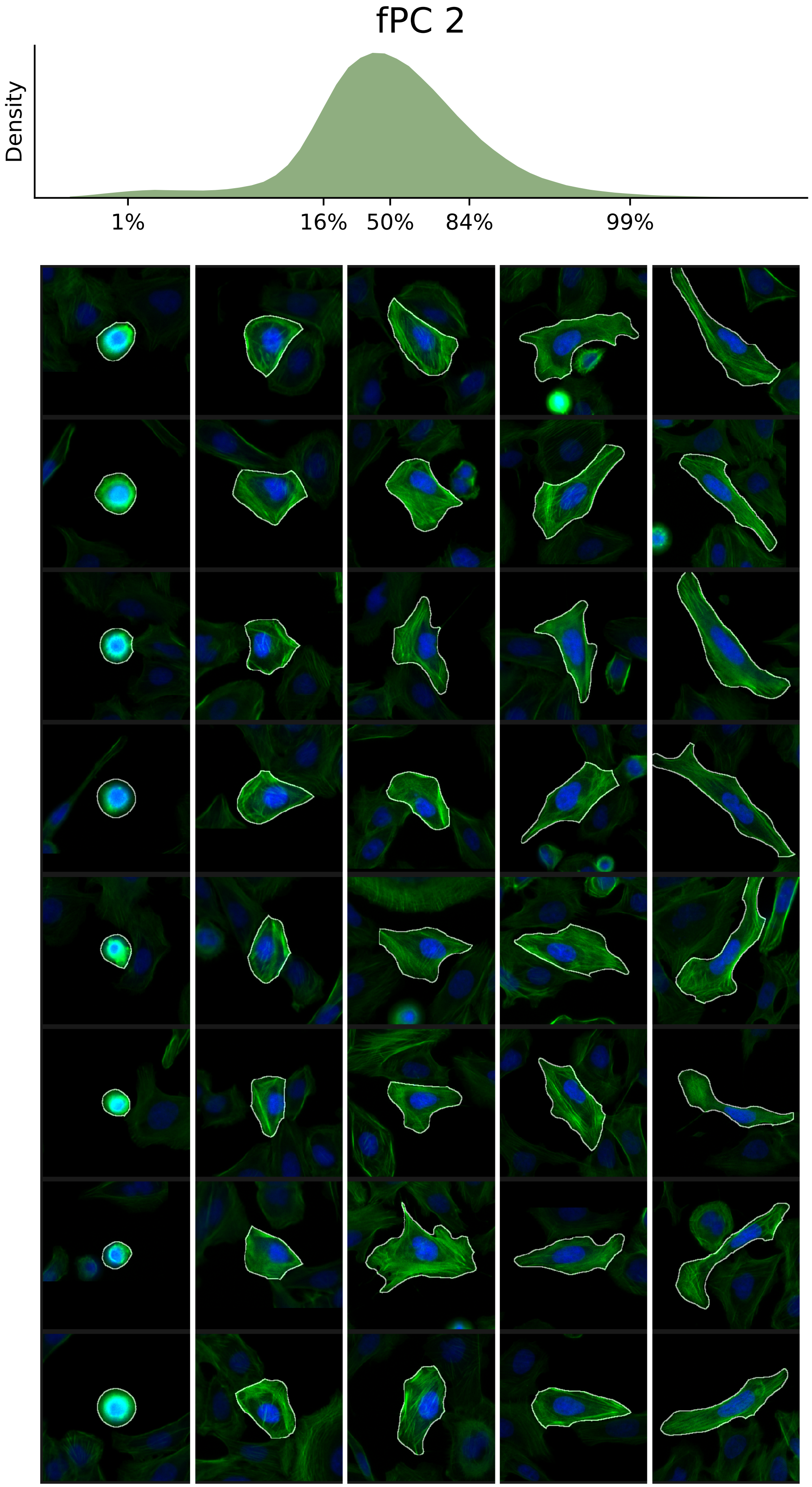

### VIEWS_fPC3.pdf

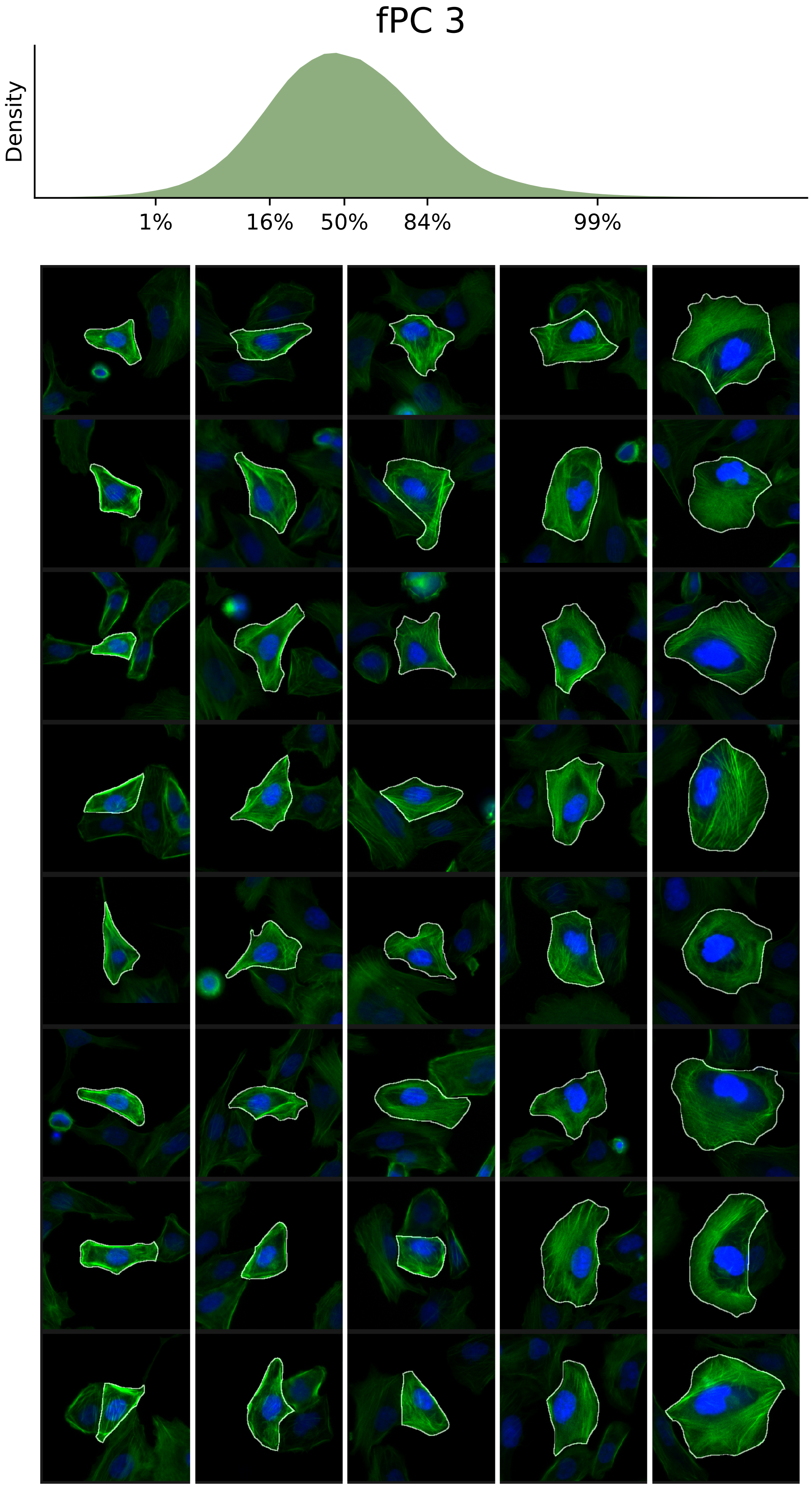

### VIEWS_fPC4.pdf

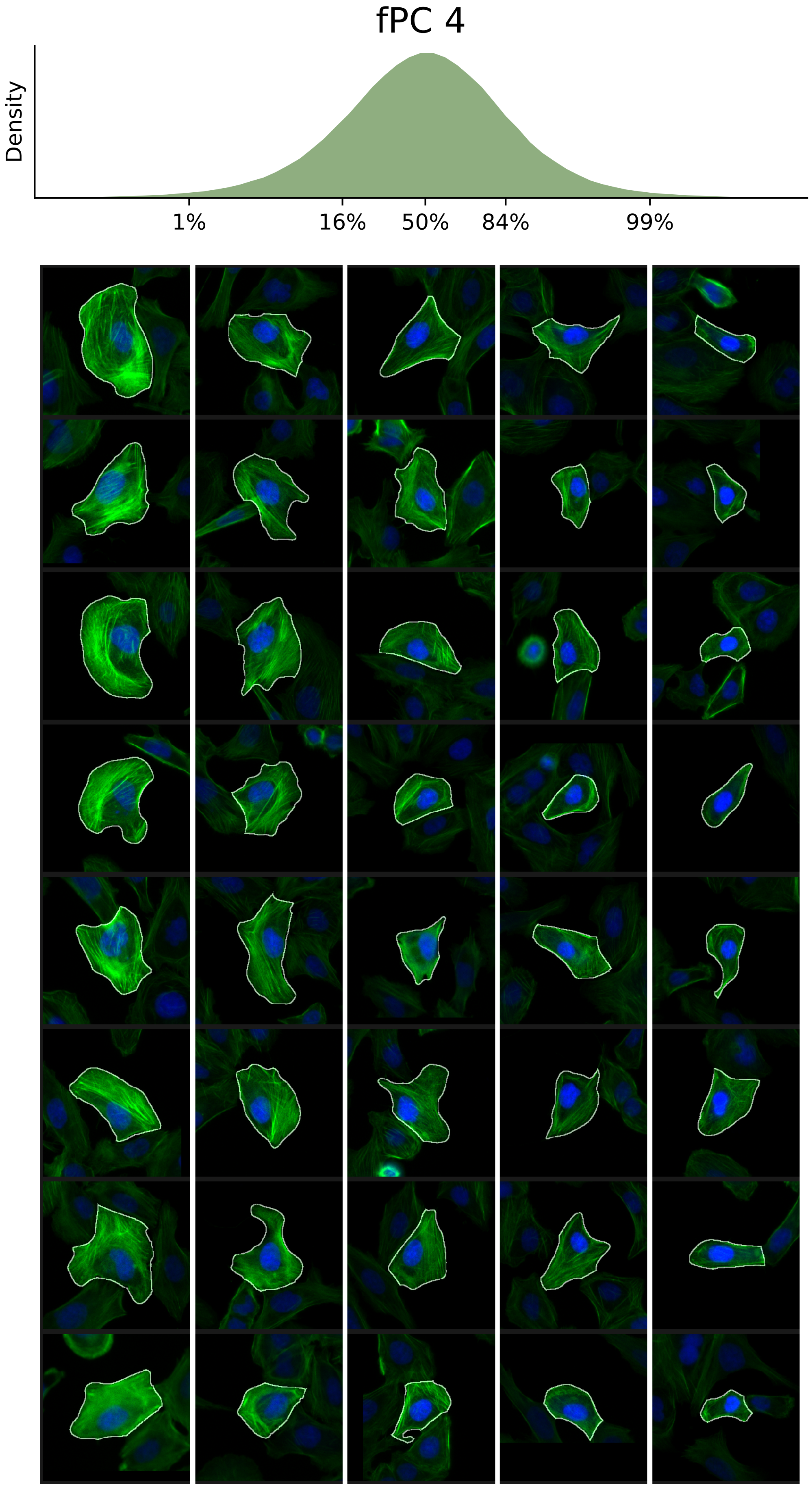

### VIEWS_fPC5.pdf

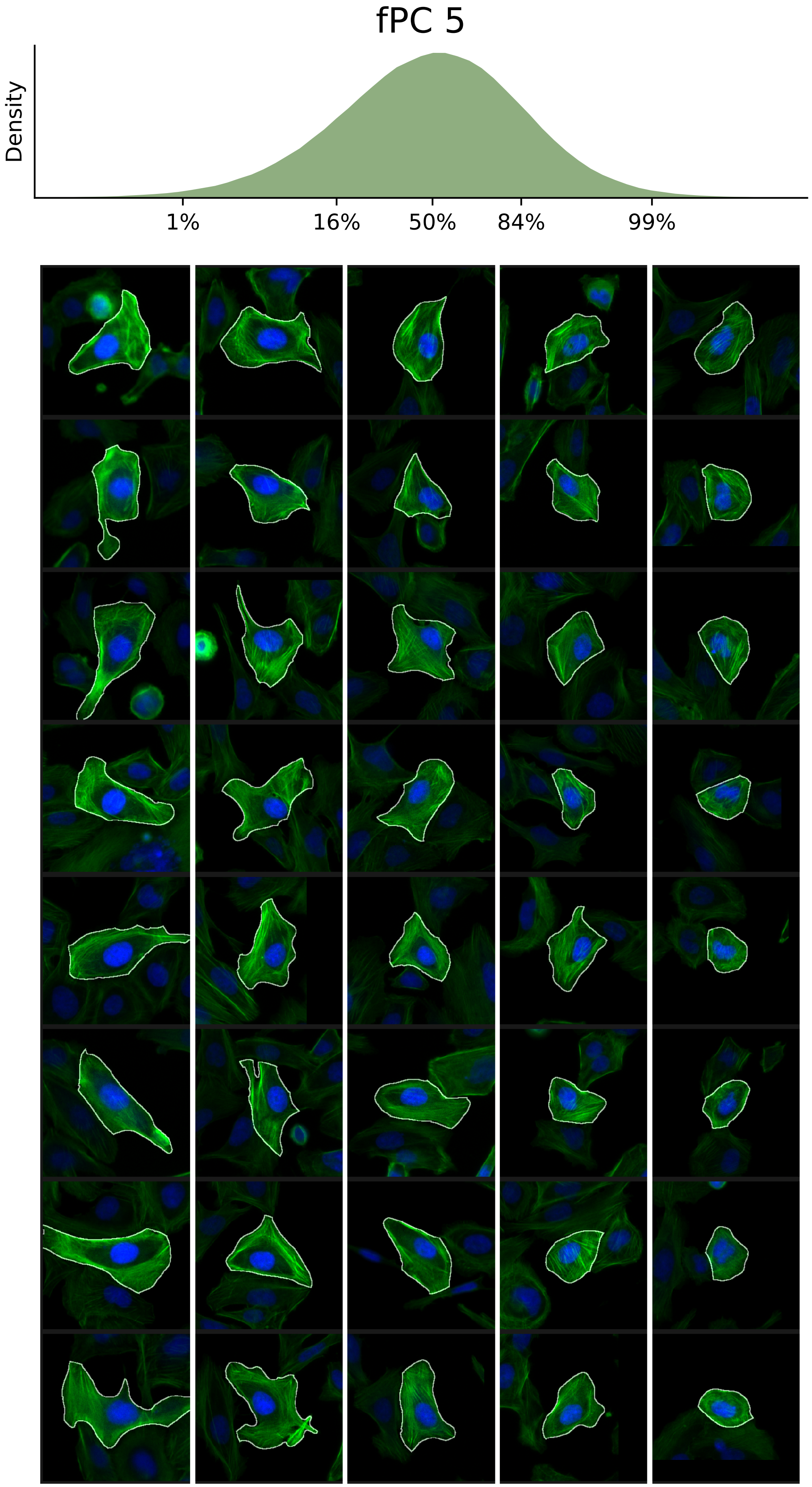

### VIEWS_fPC6.pdf

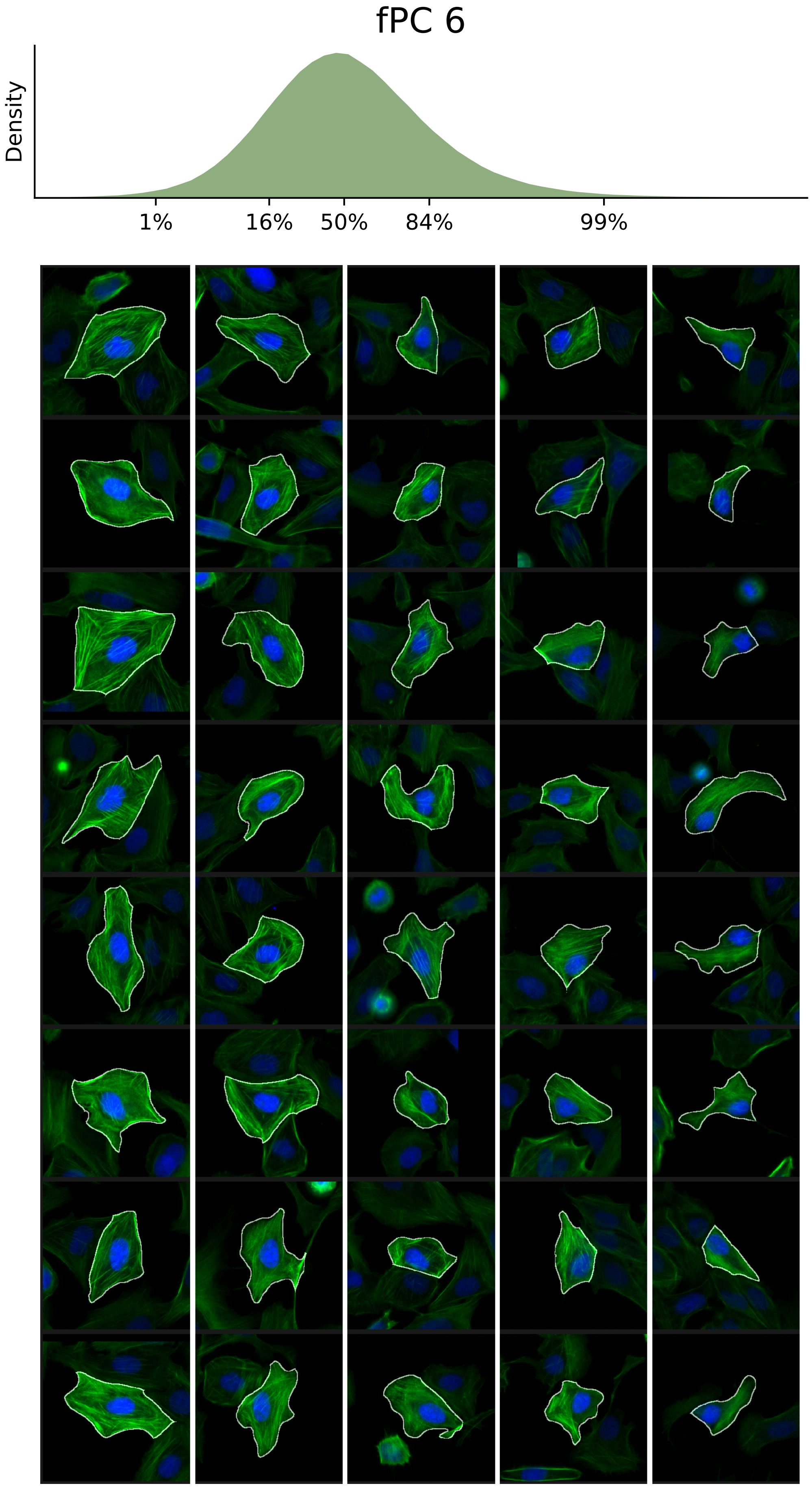

### VIEWS_fPC7.pdf

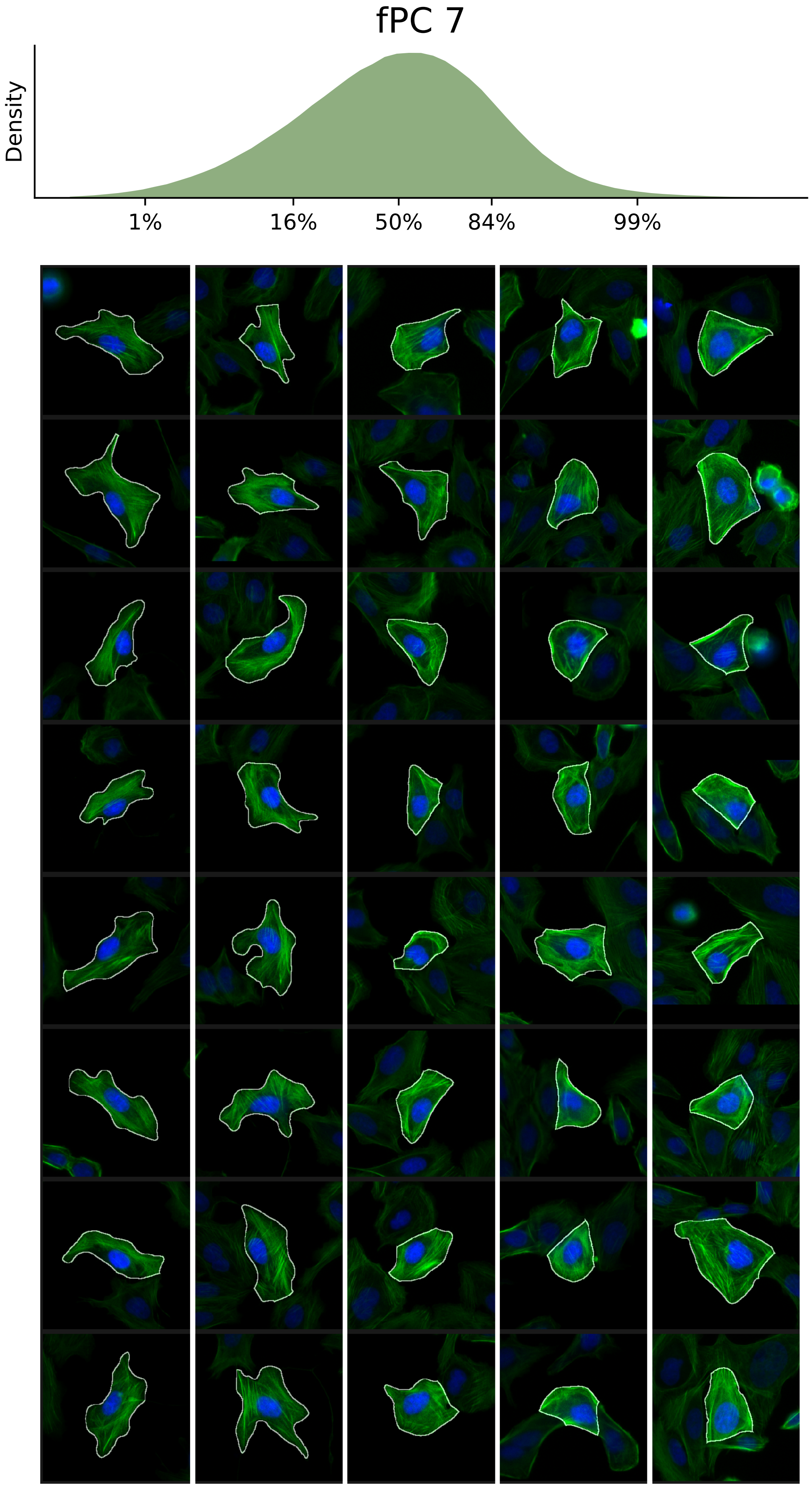

### VIEWS_fPC8.pdf

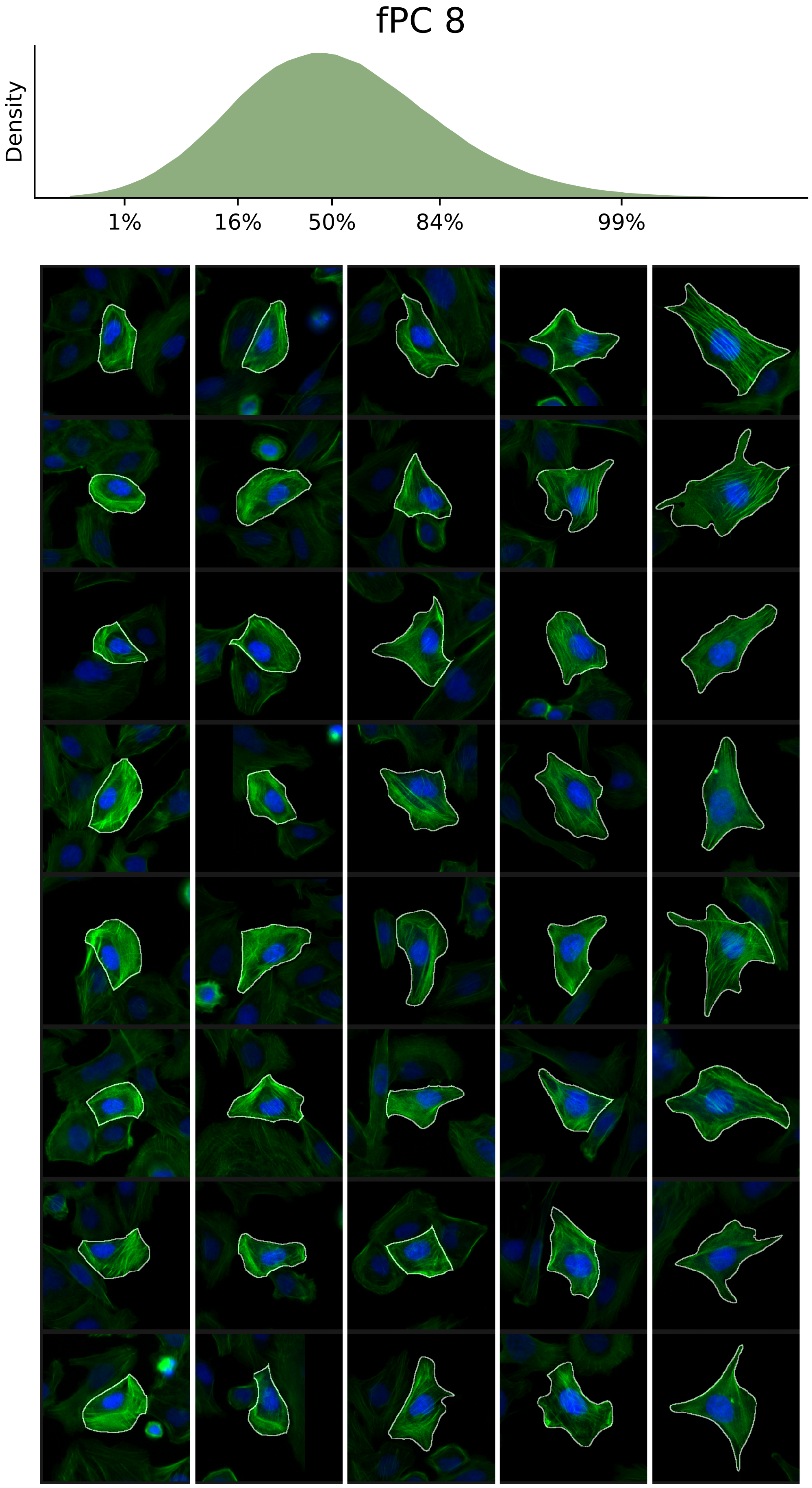

### VIEWS_fPC9.pdf

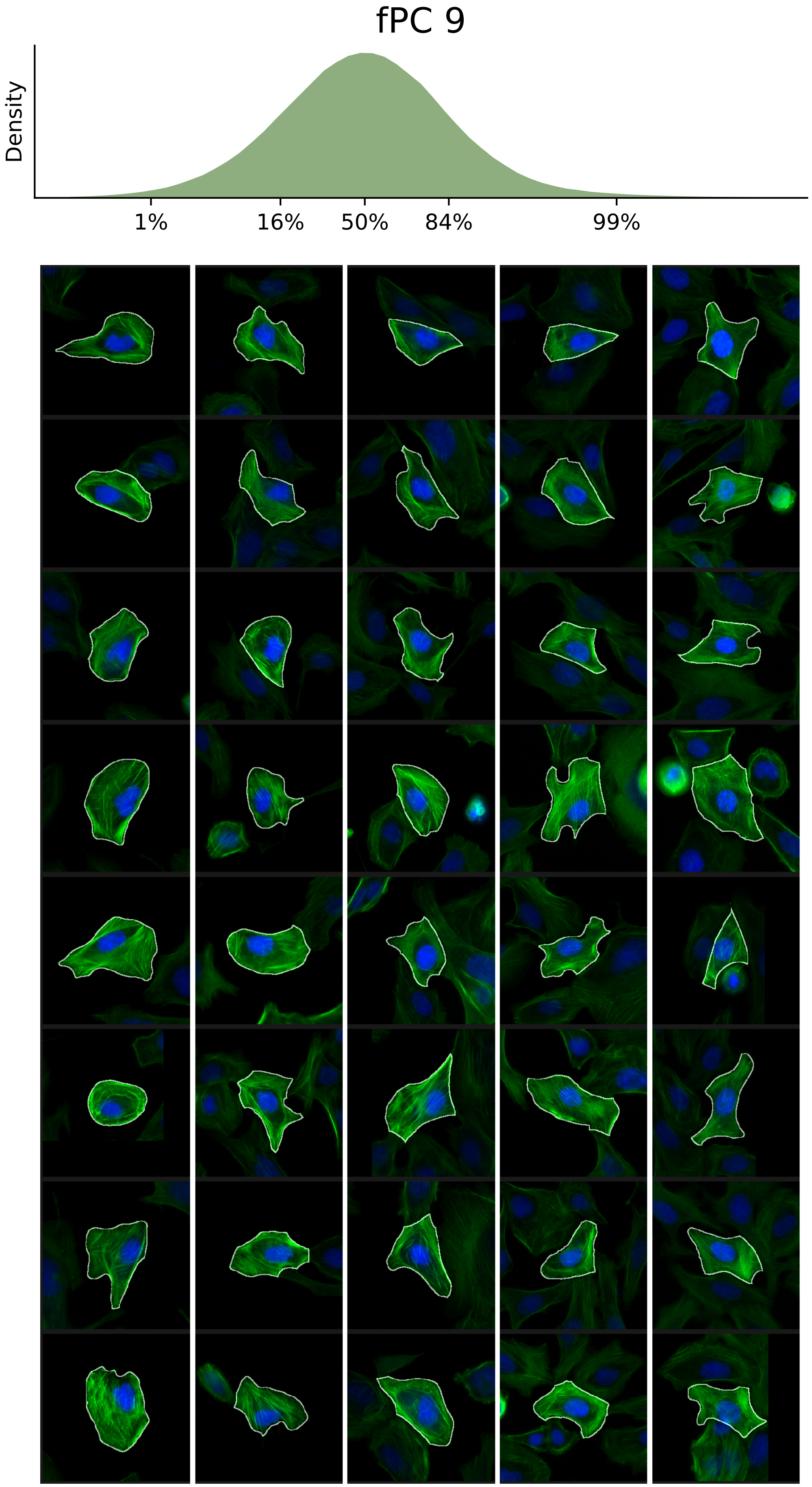

### VIEWS_fPC10.pdf

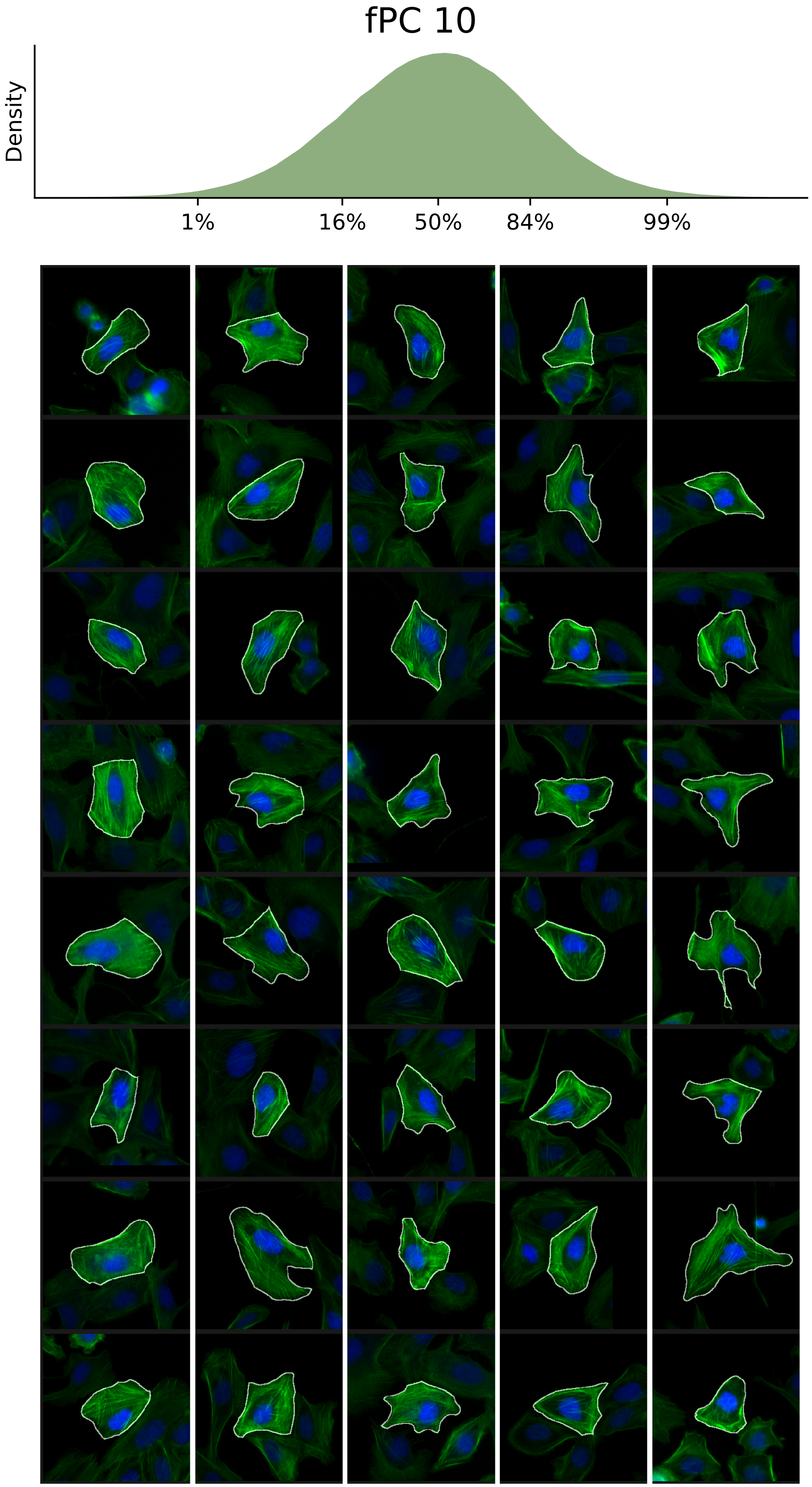

### VIEWS_fPC11.pdf

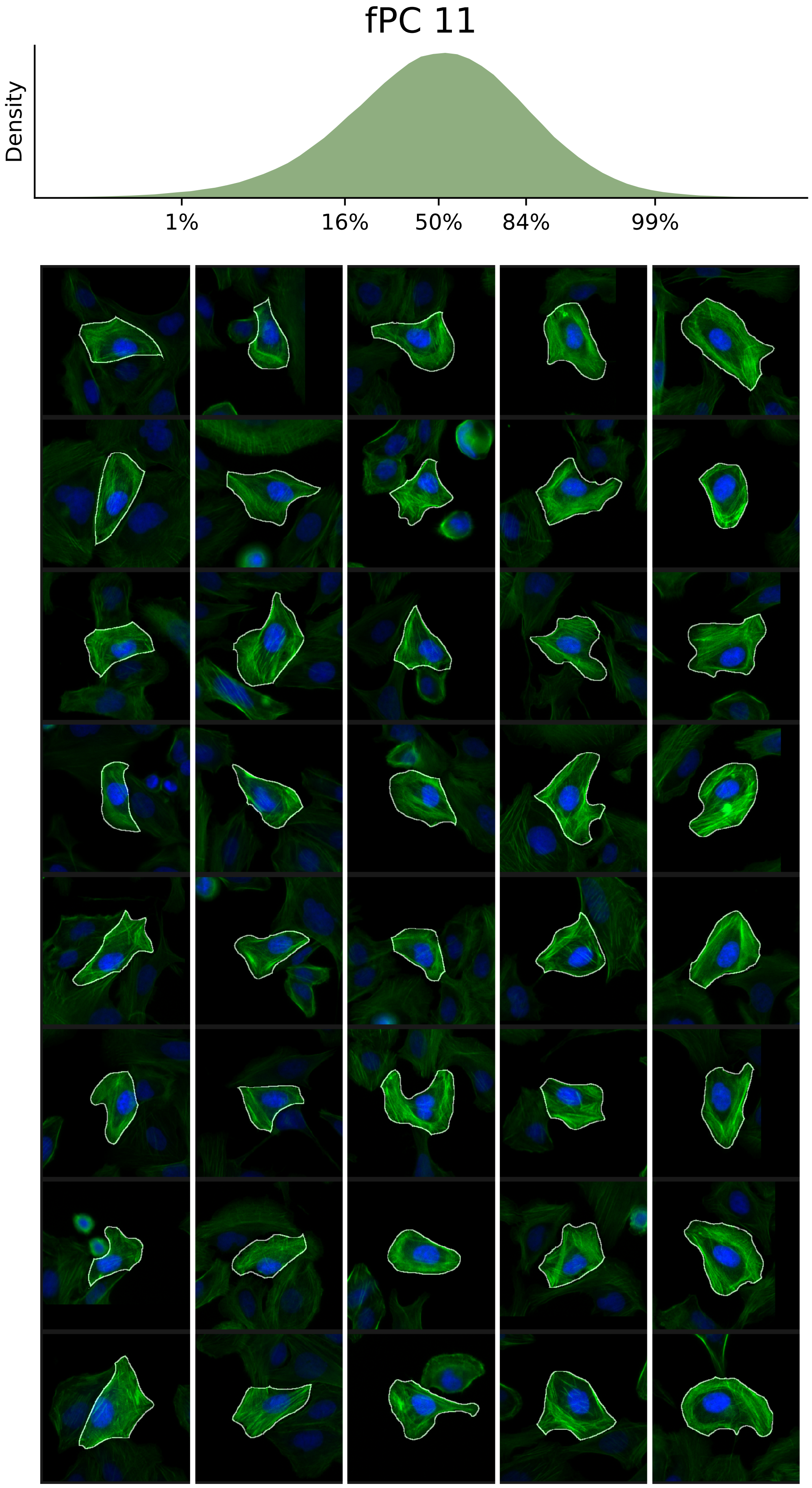

### VIEWS_fPC12.pdf

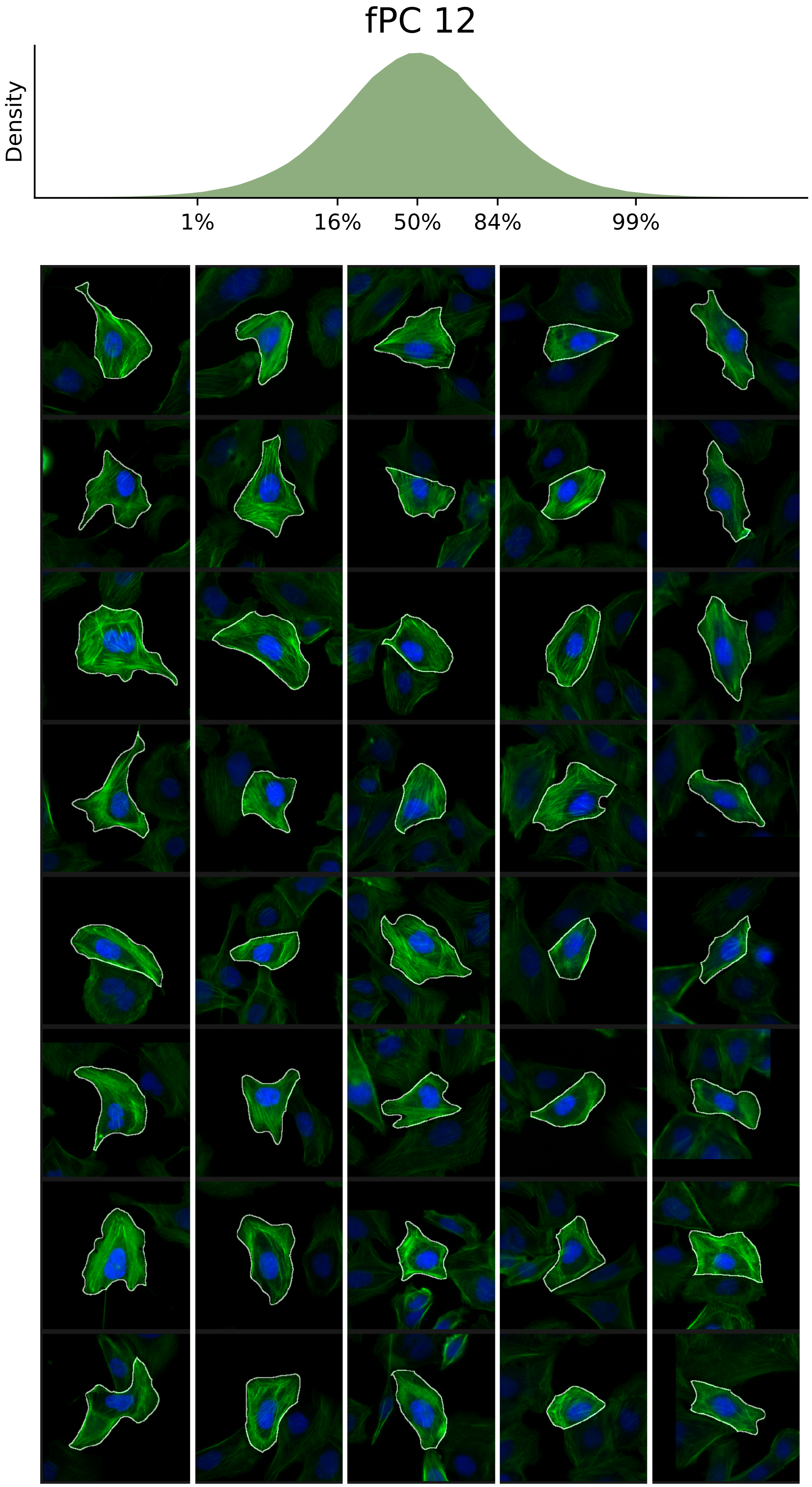

### VIEWS_fPC13.pdf

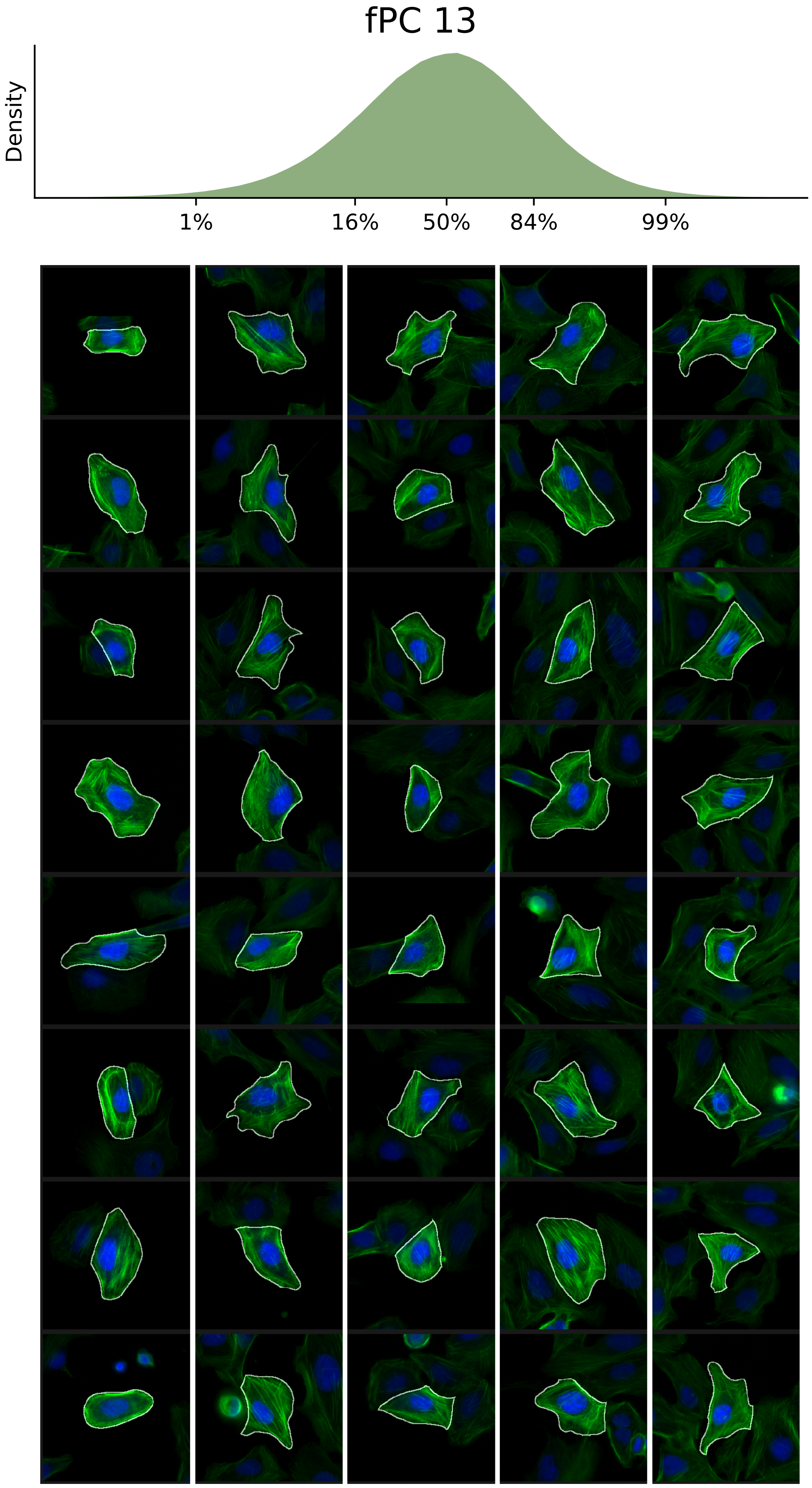

### VIEWS_fPC14.pdf

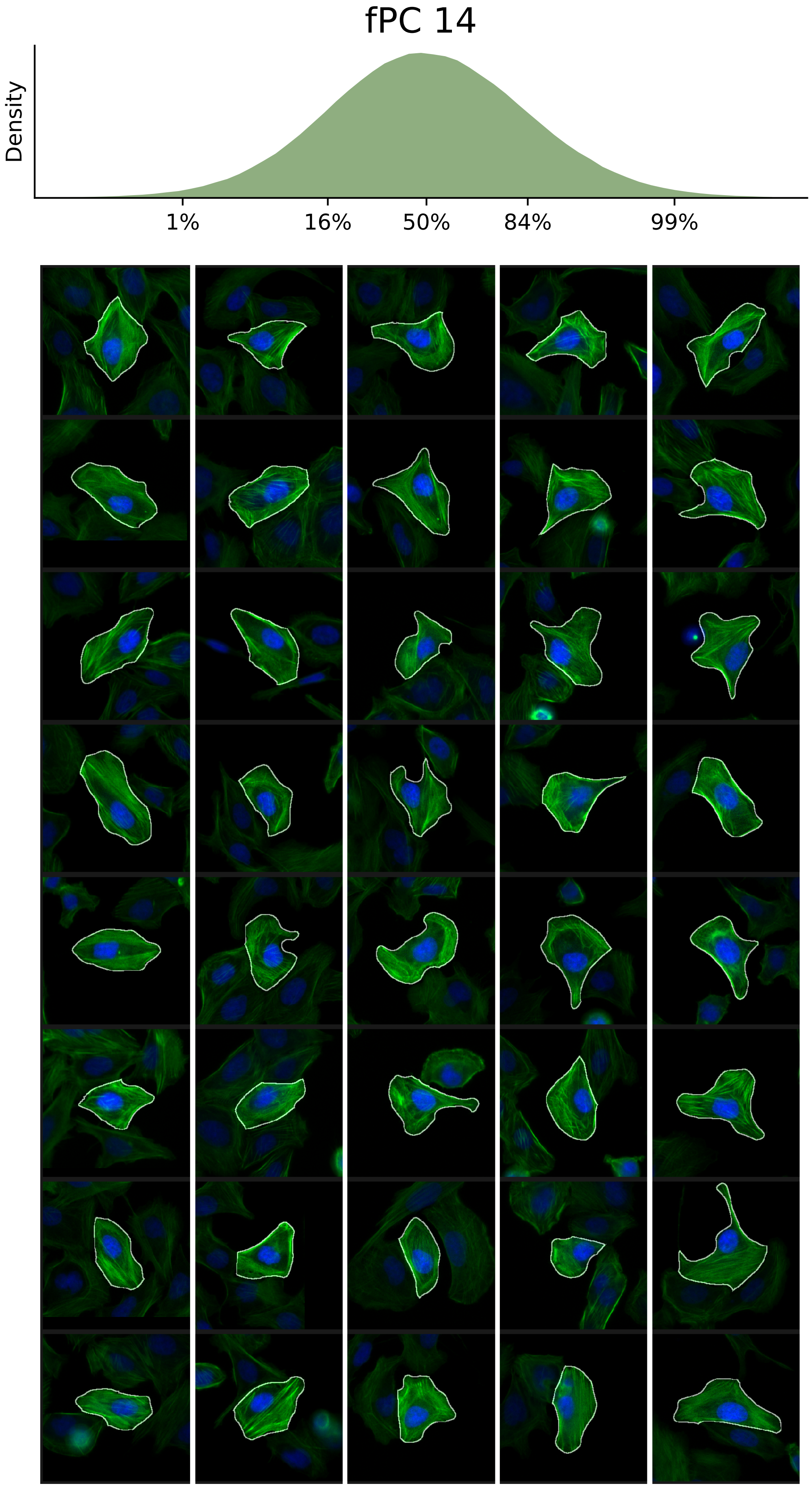

### VIEWS_fPC15.pdf

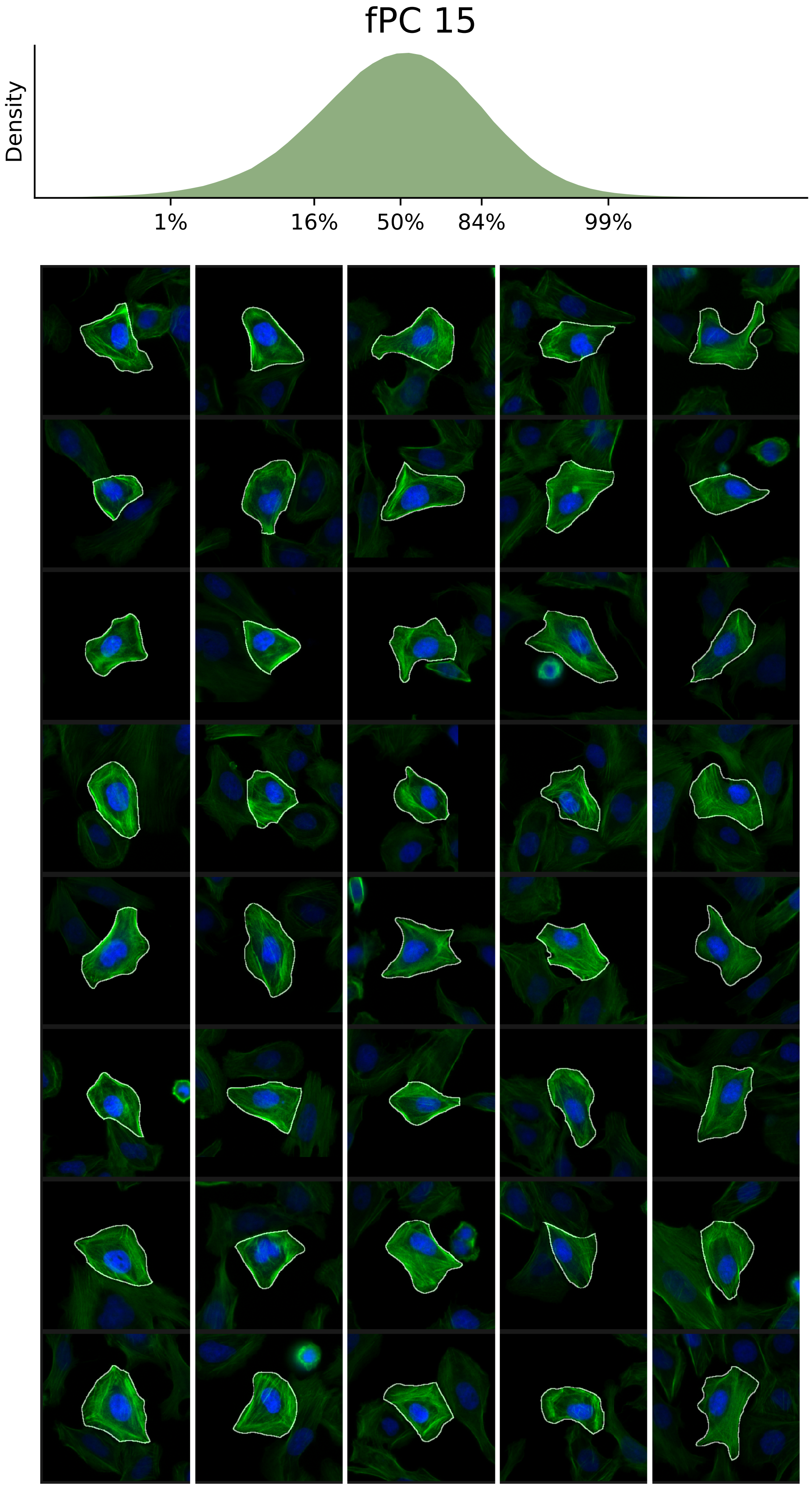

### VIEWS_fPC16.pdf

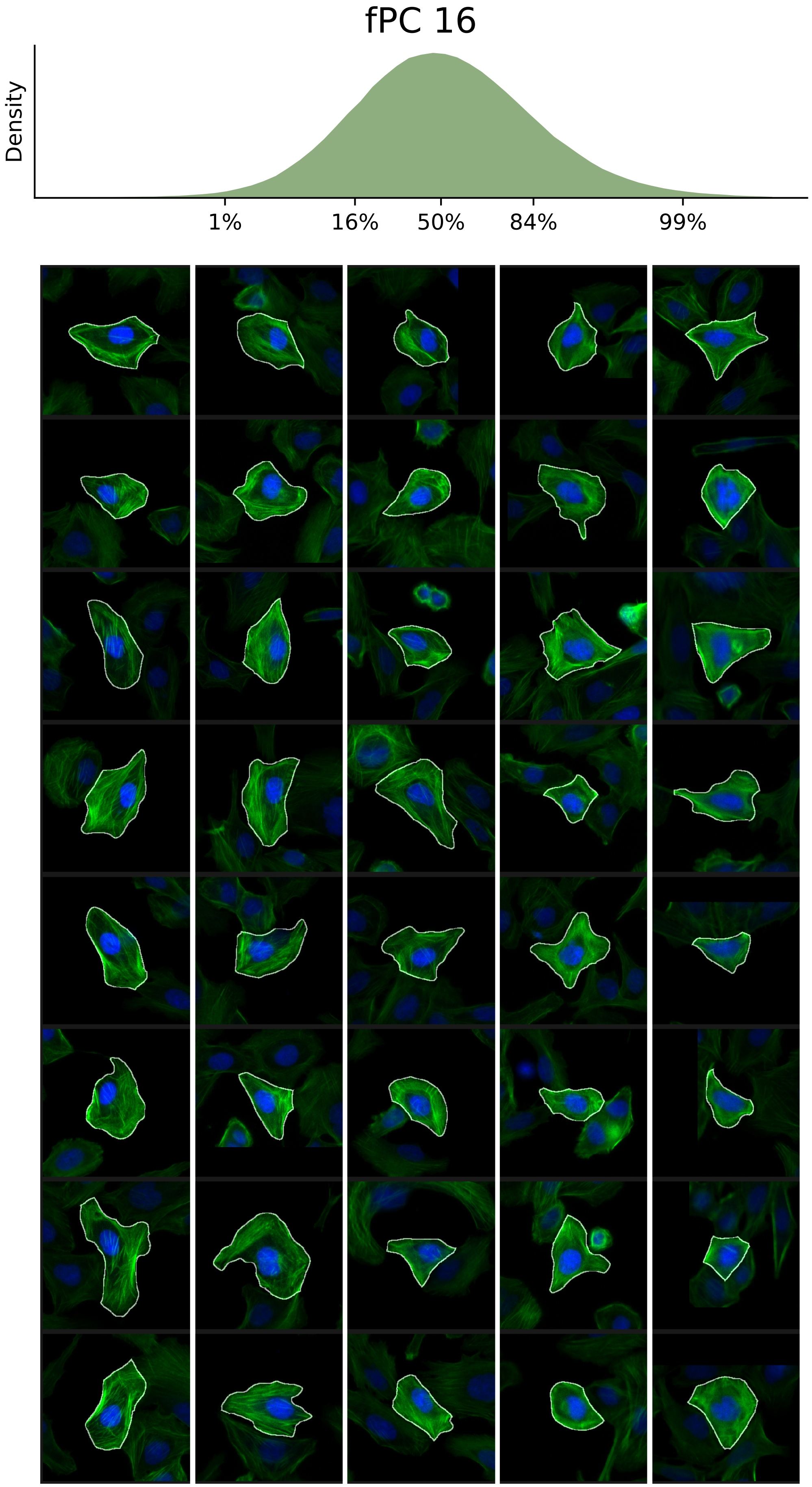

### VIEWS_fPC17.pdf

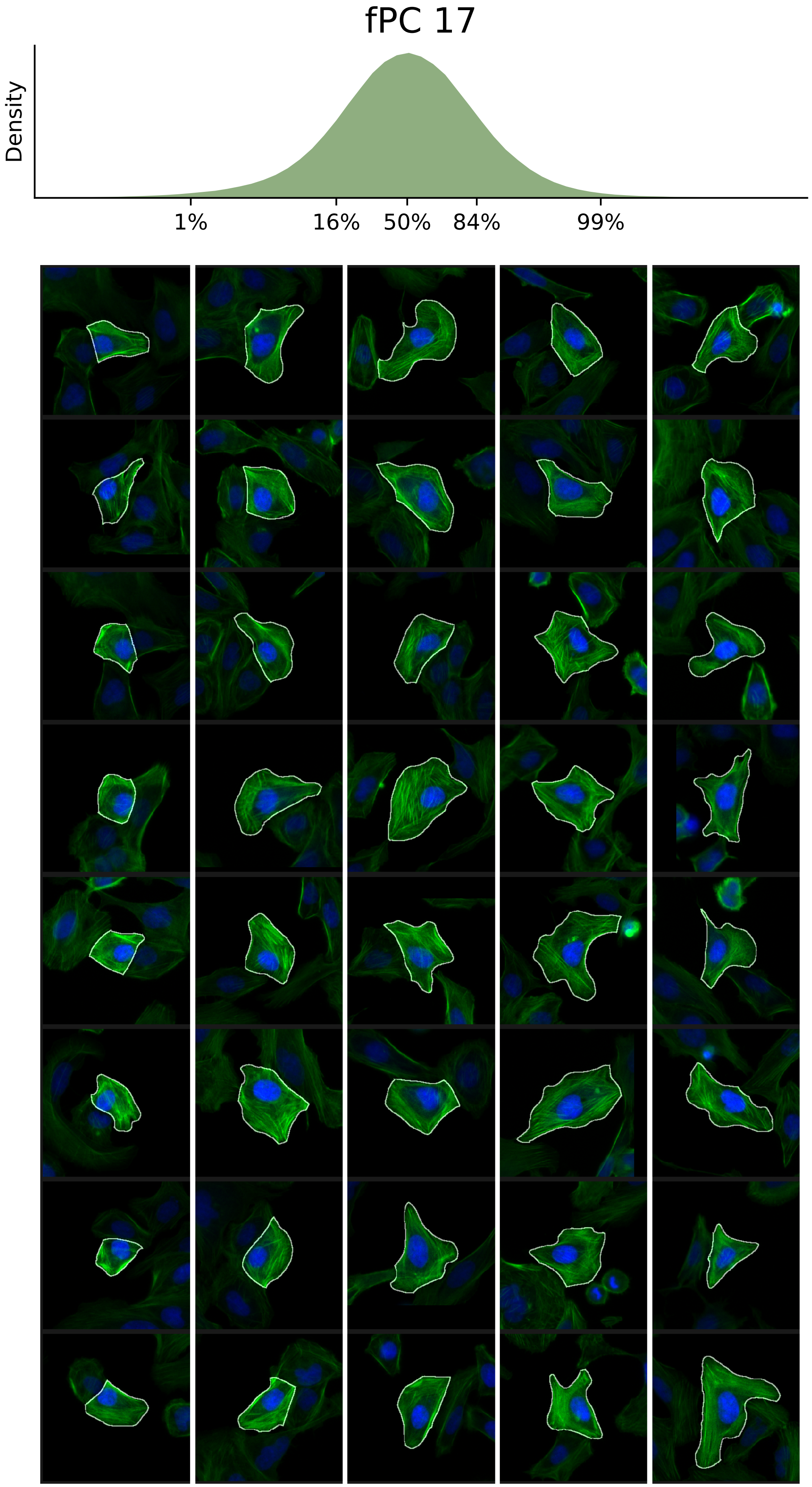

### VIEWS_fPC18.pdf

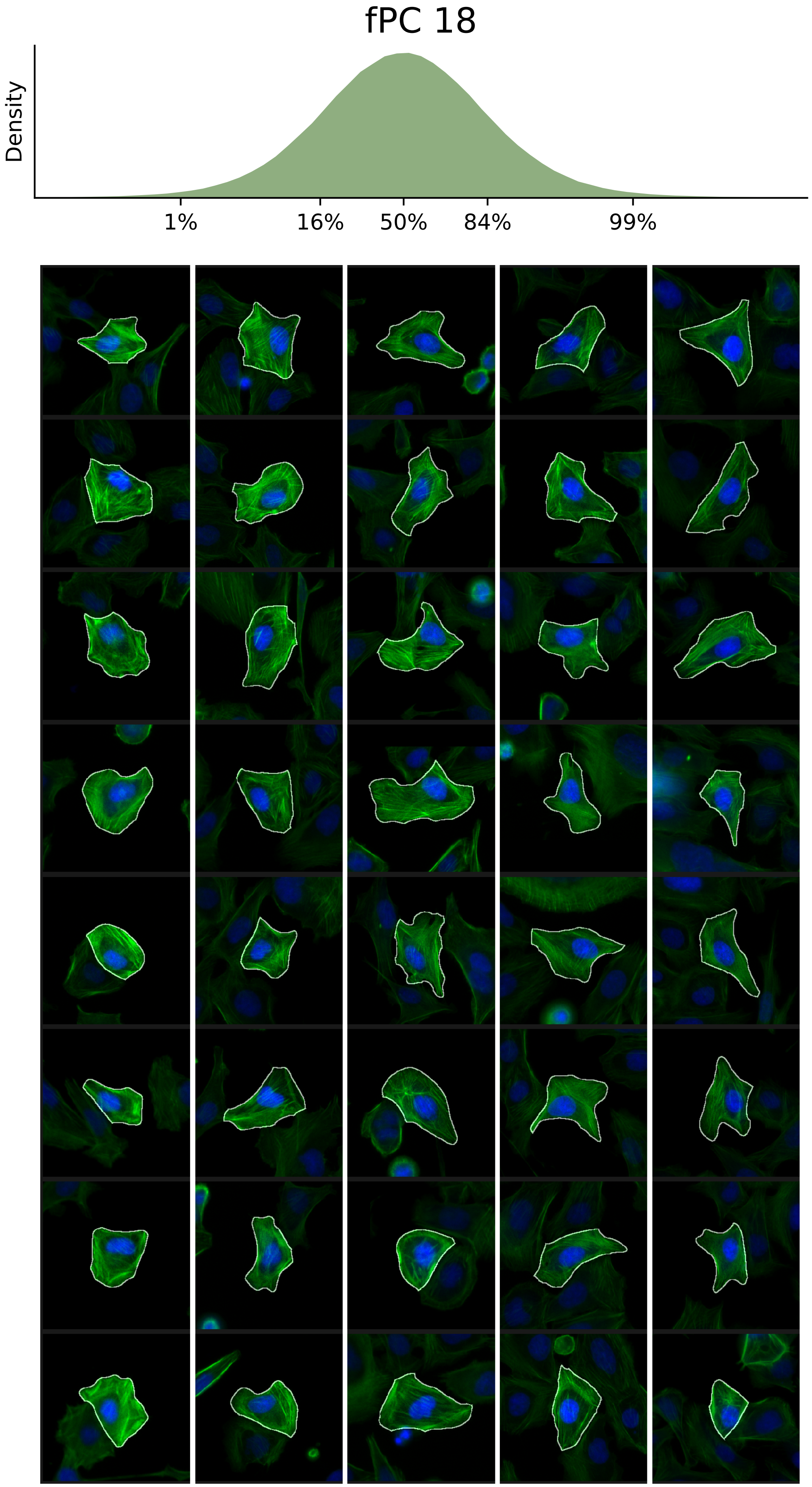

### VIEWS_fPC19.pdf

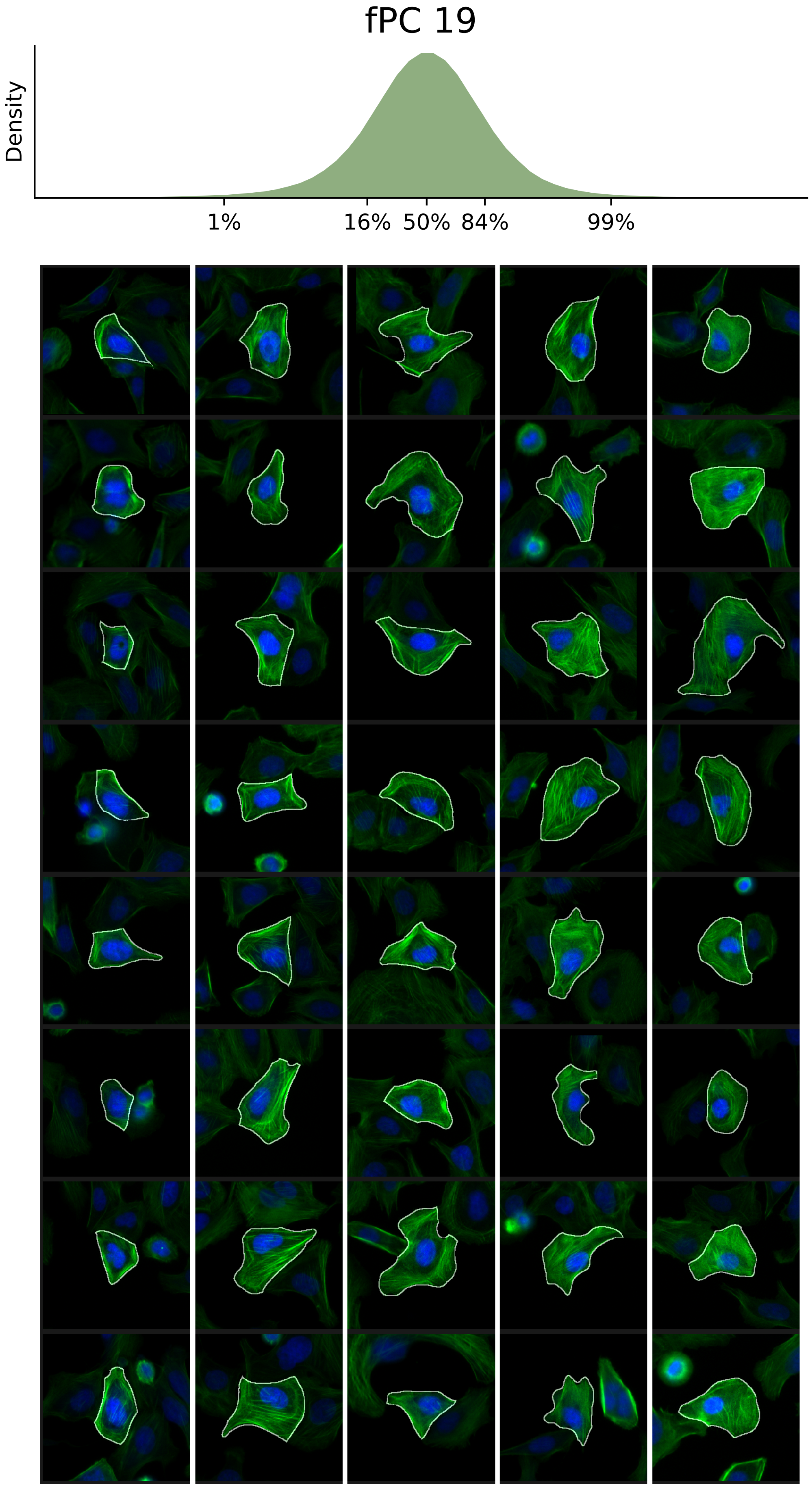

### VIEWS_fPC20.pdf

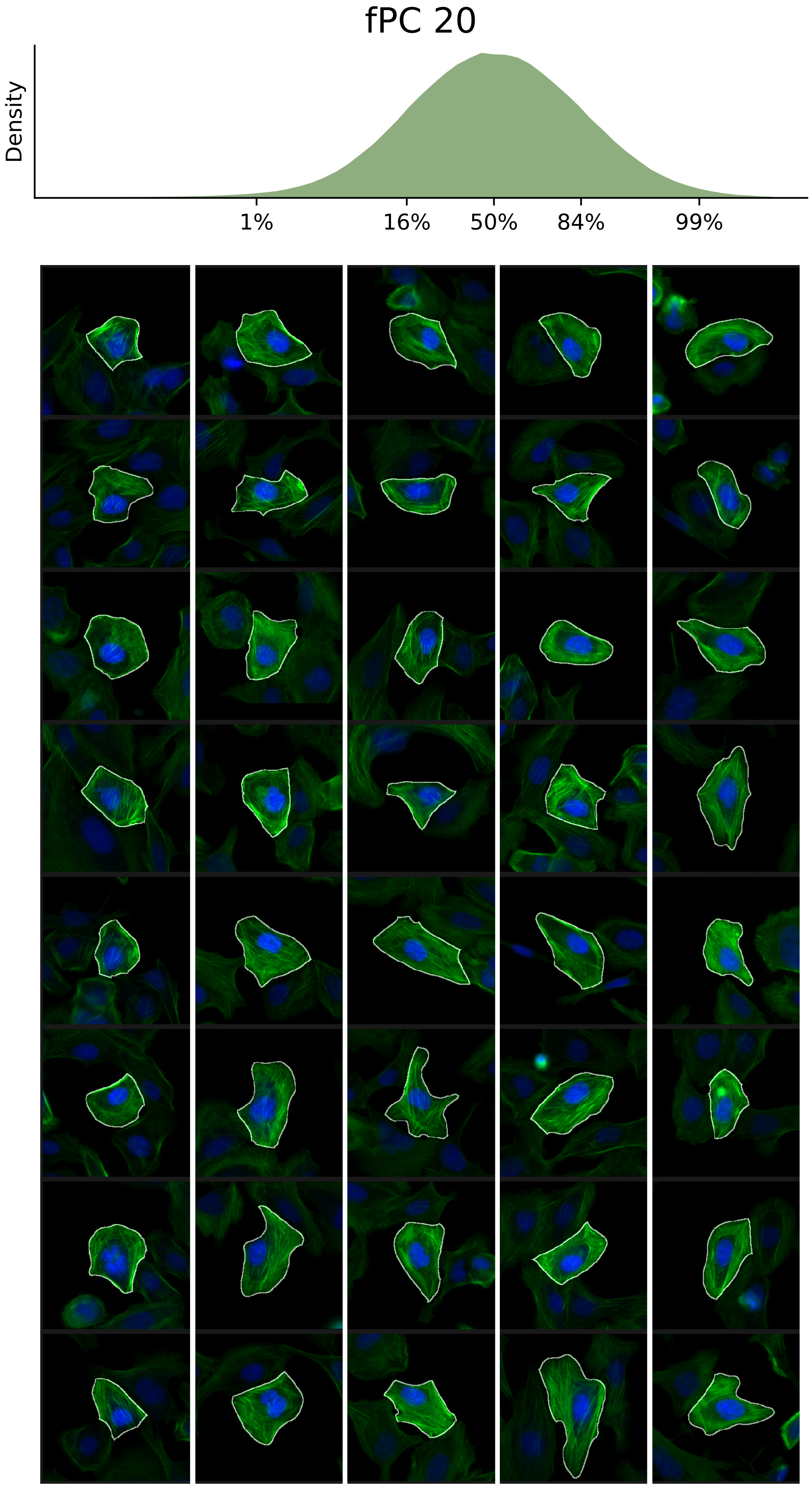

### VIEWS_fPC21.pdf

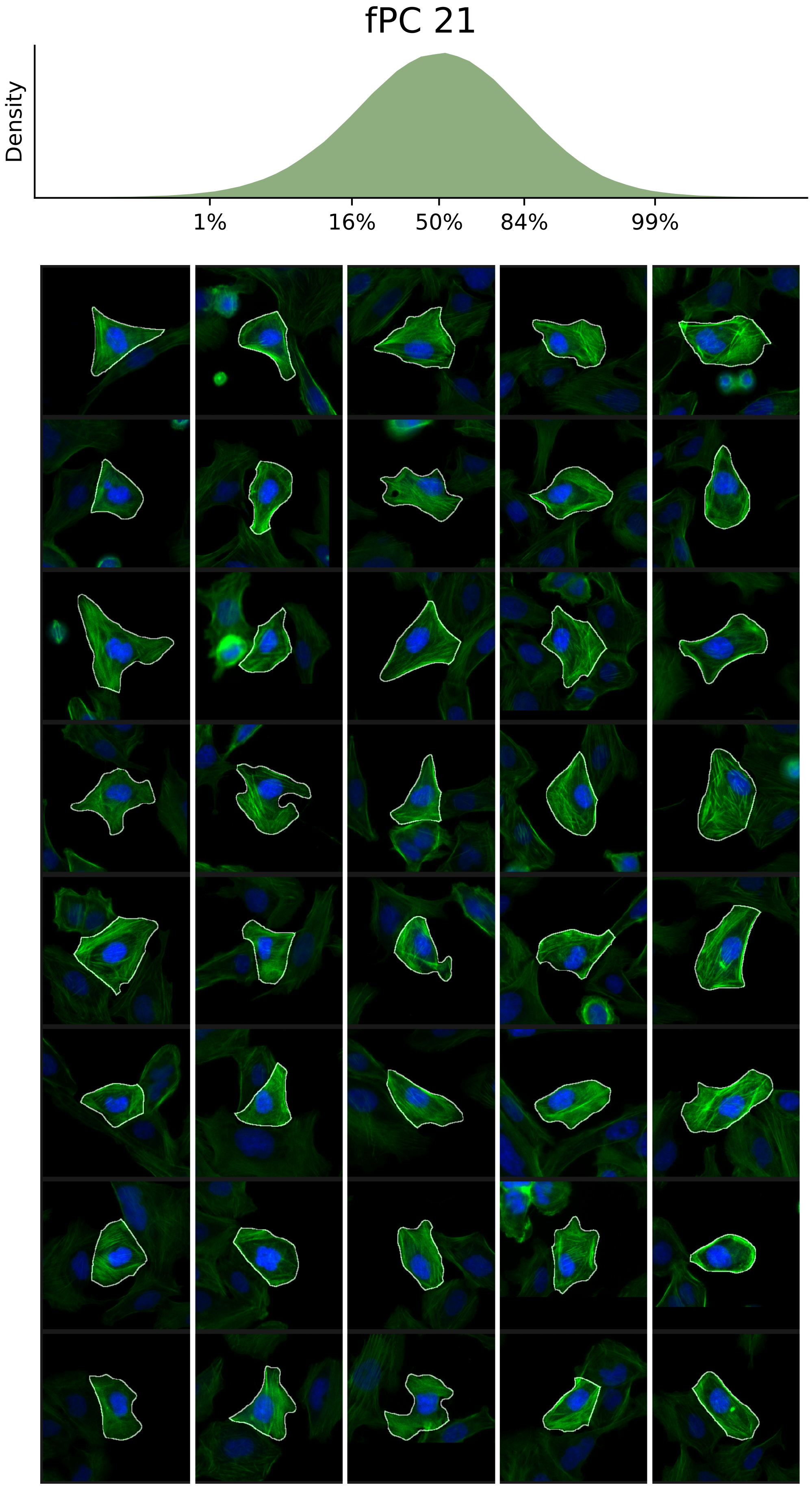

### VIEWS_fPC22.pdf

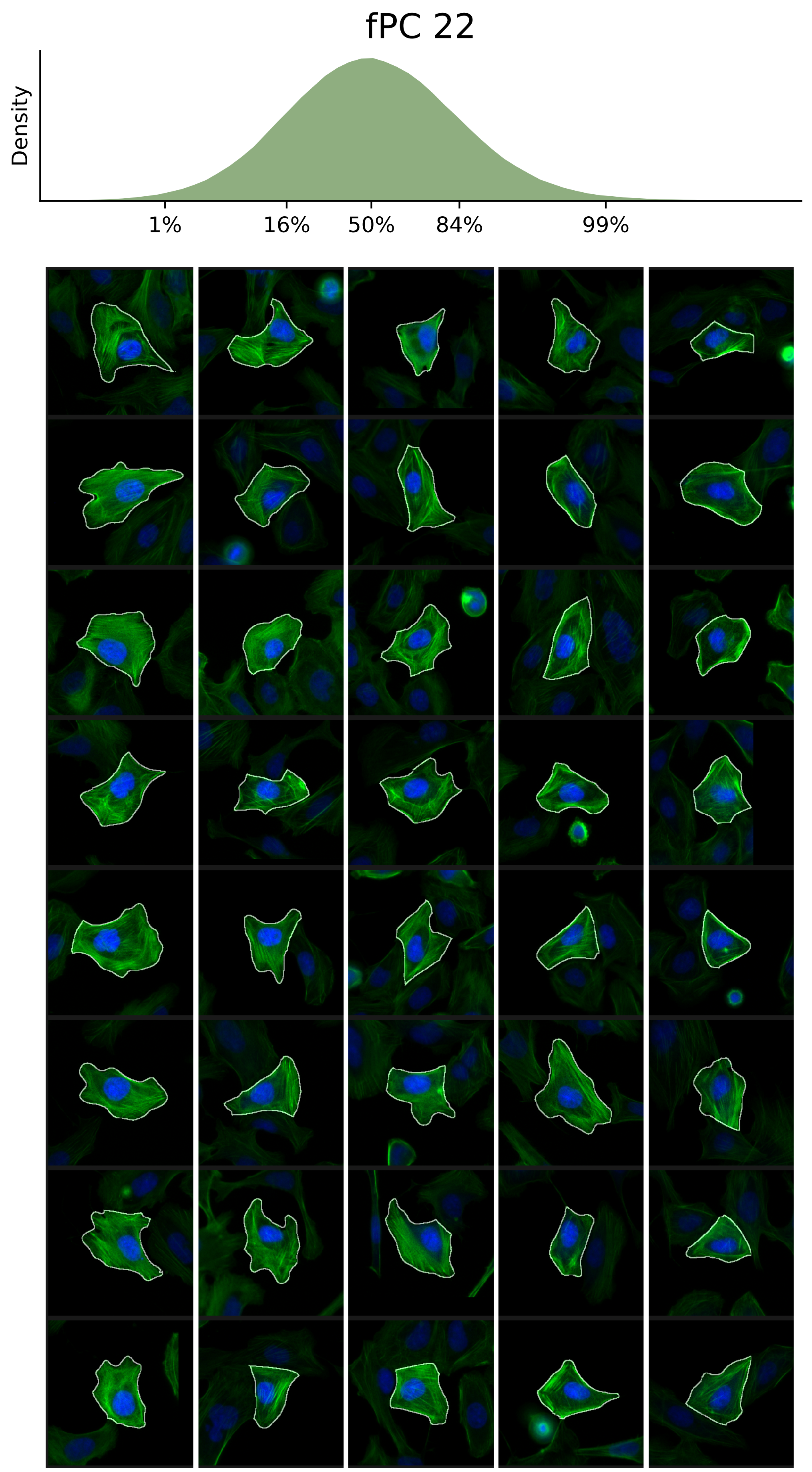

### VIEWS_fPC23.pdf

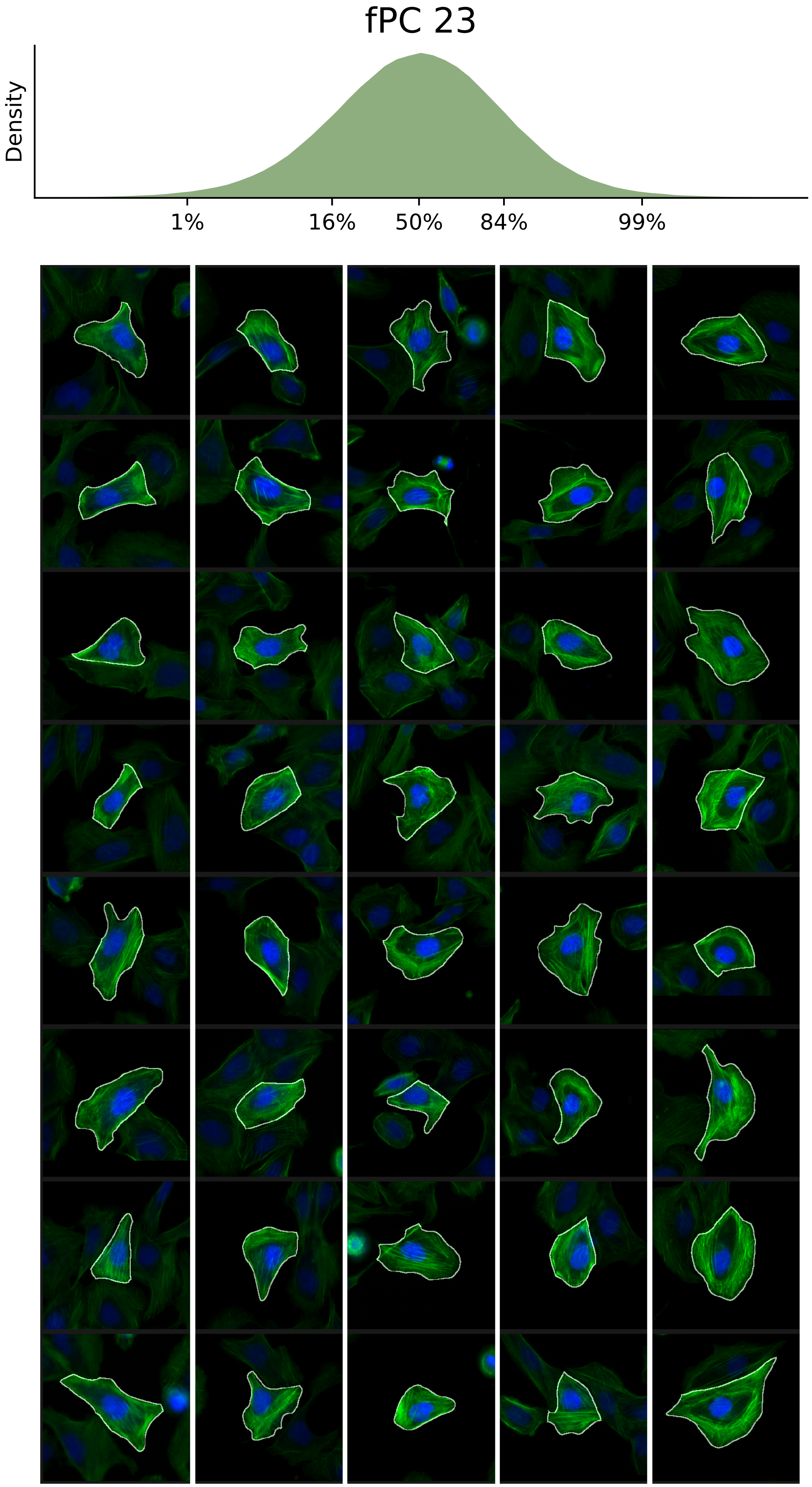

### VIEWS_fPC24.pdf

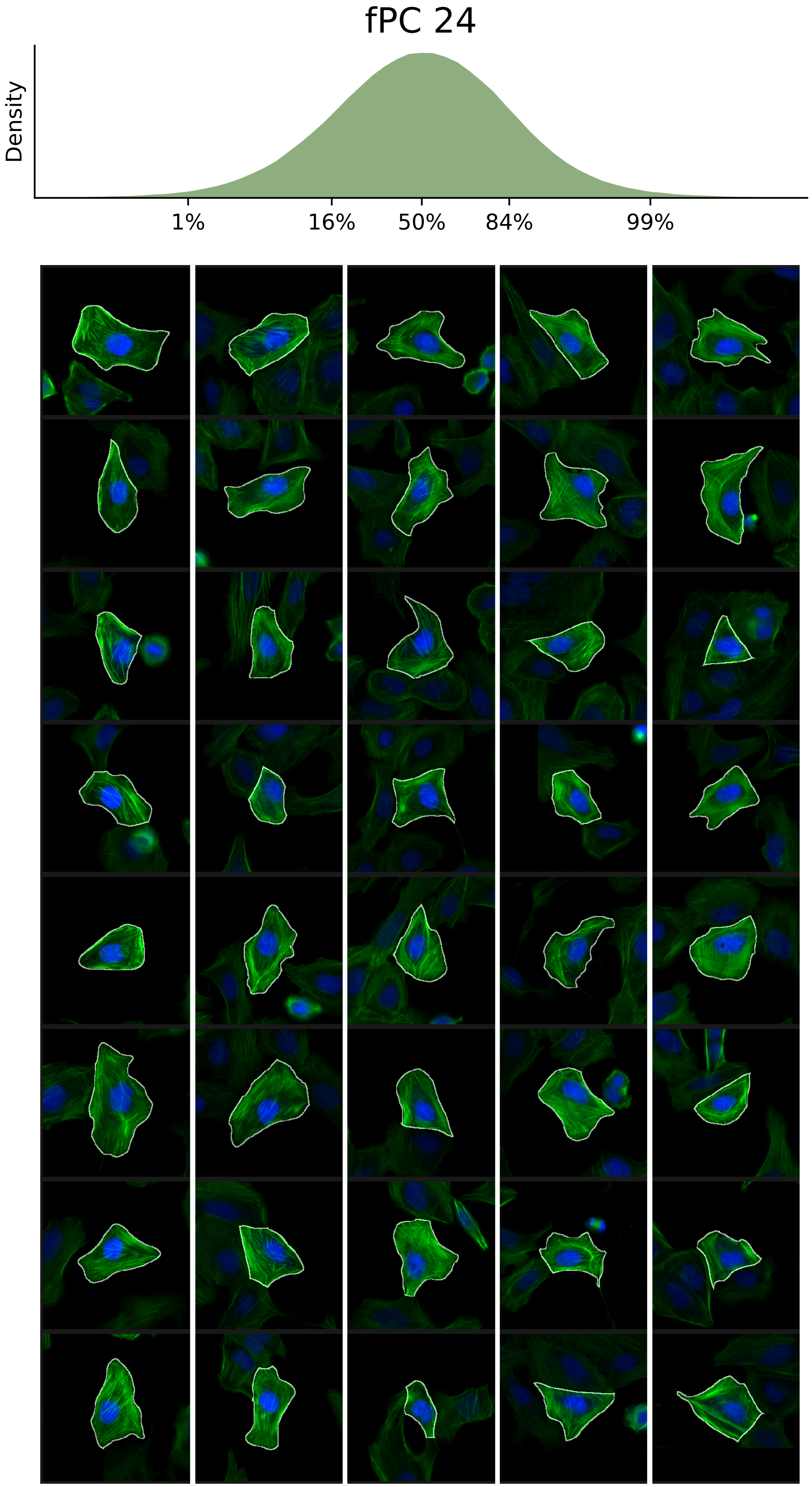

### VIEWS_fPC25.pdf

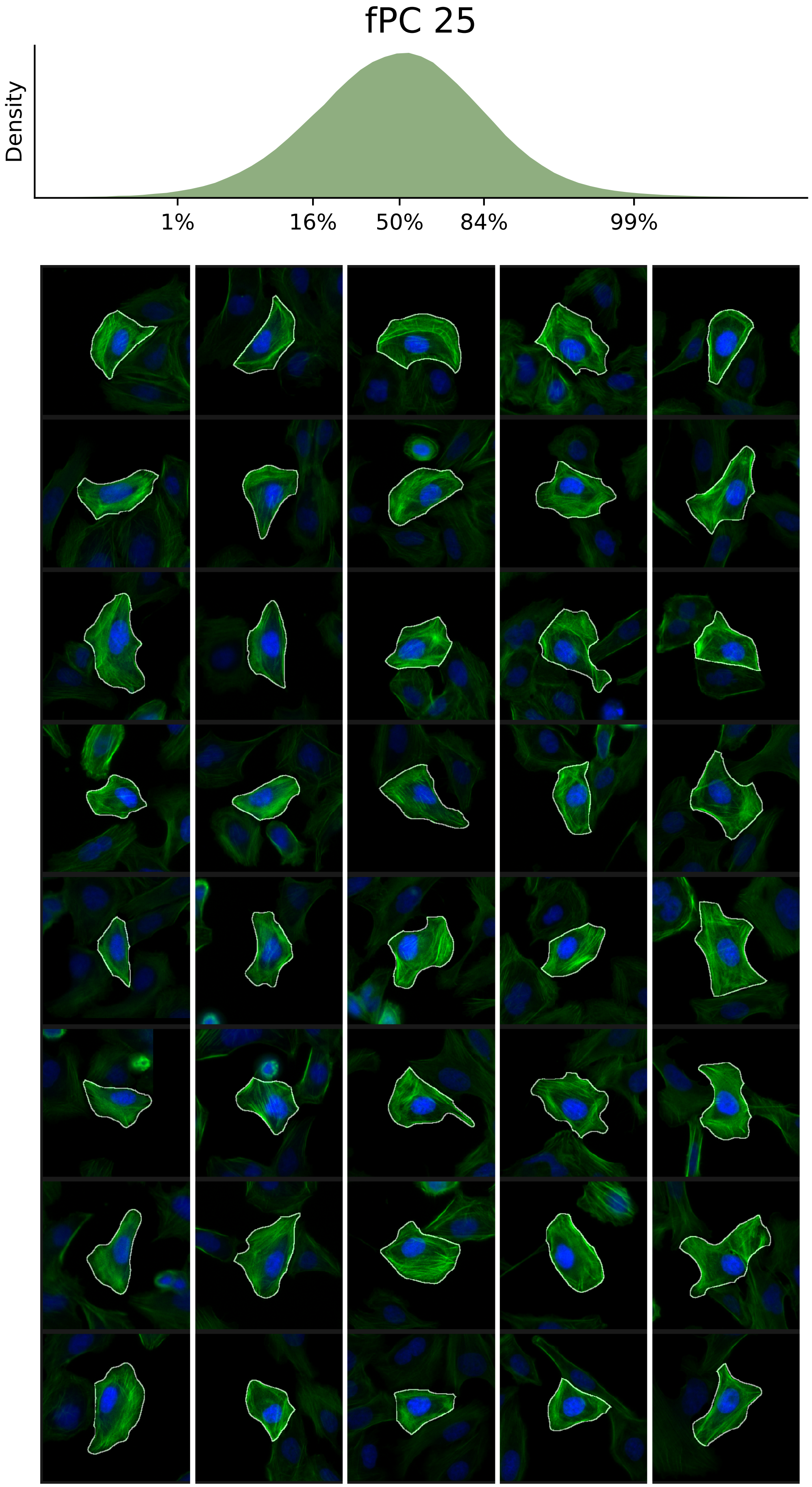

### VIEWS_fPC-LD-verified_ACTB_rep1.pdf

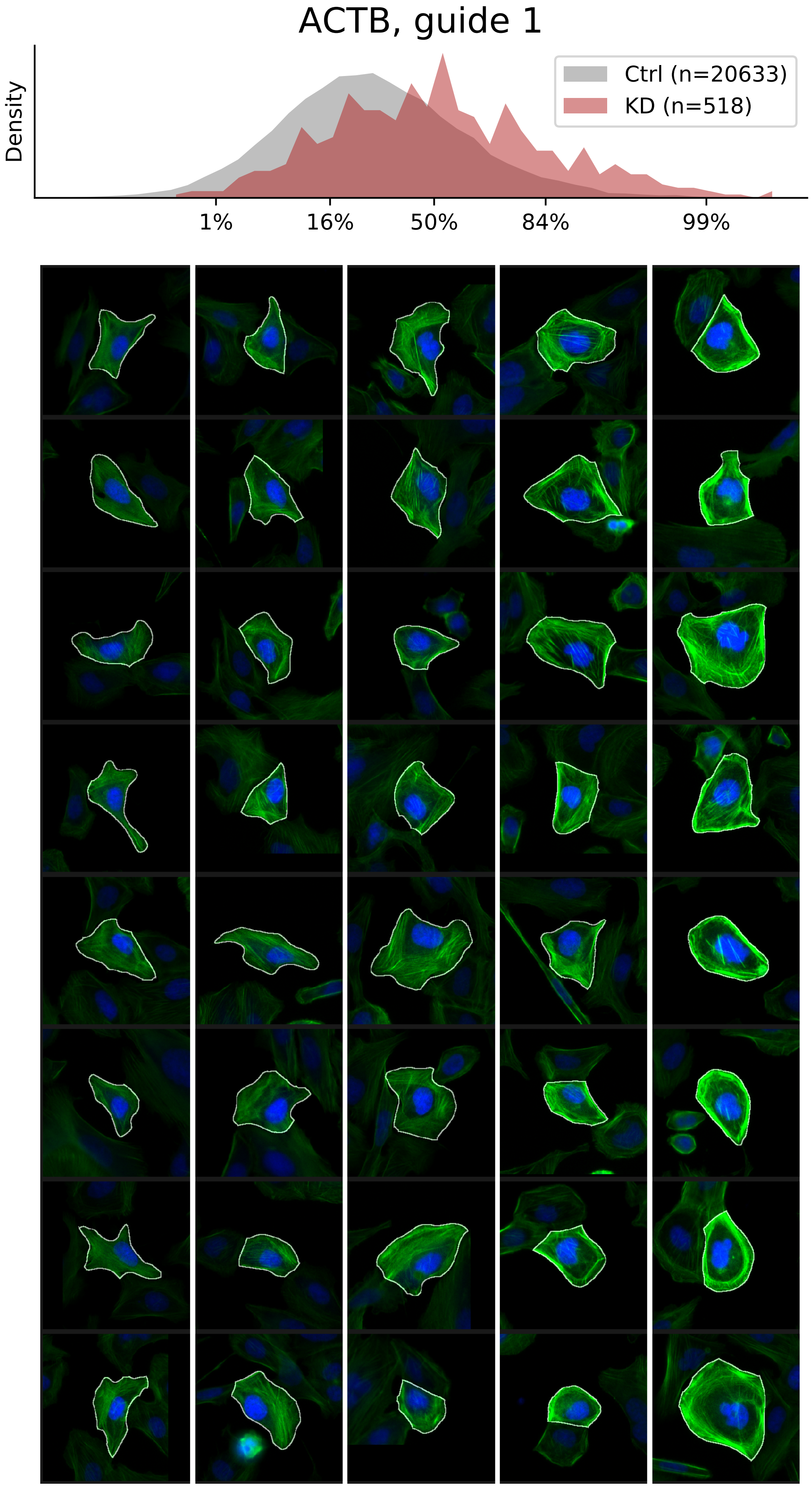

### VIEWS_fPC-LD-verified_BAIAP2L1_rep1.pdf

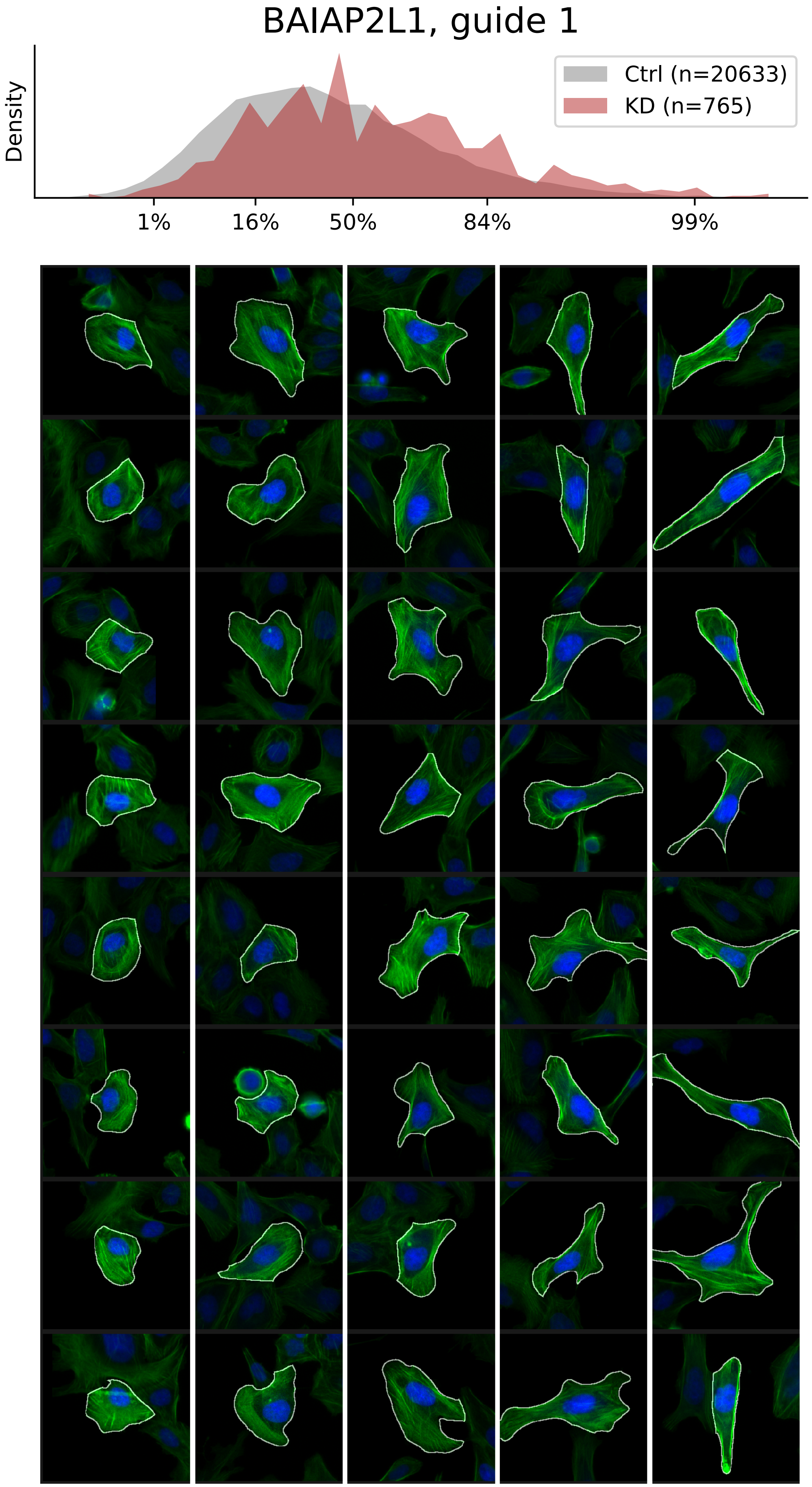

### VIEWS_fPC-LD-verified_MESDC1_rep3.pdf

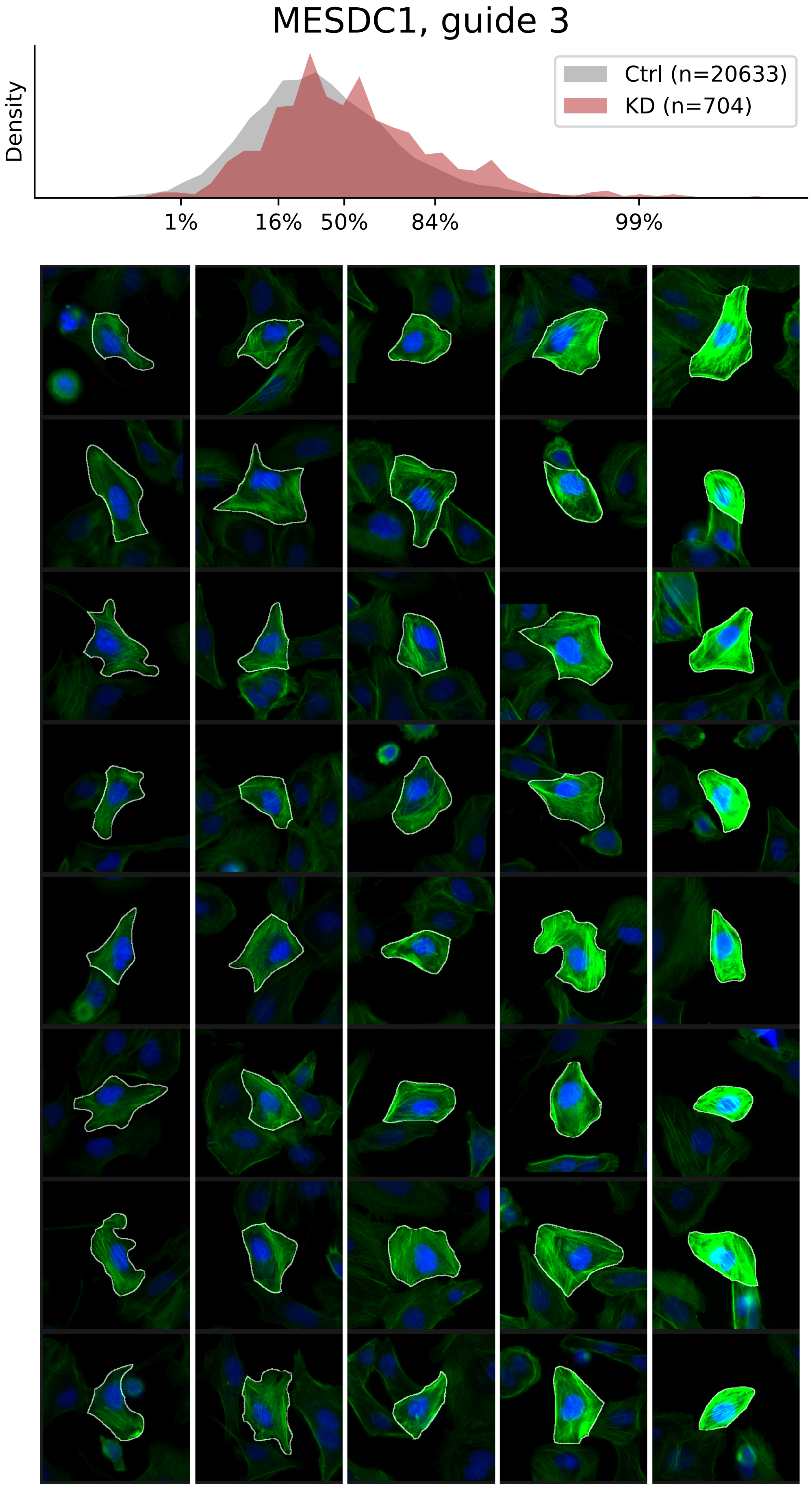

### VIEWS_fPC-LD-verified_PLEC_rep3.pdf

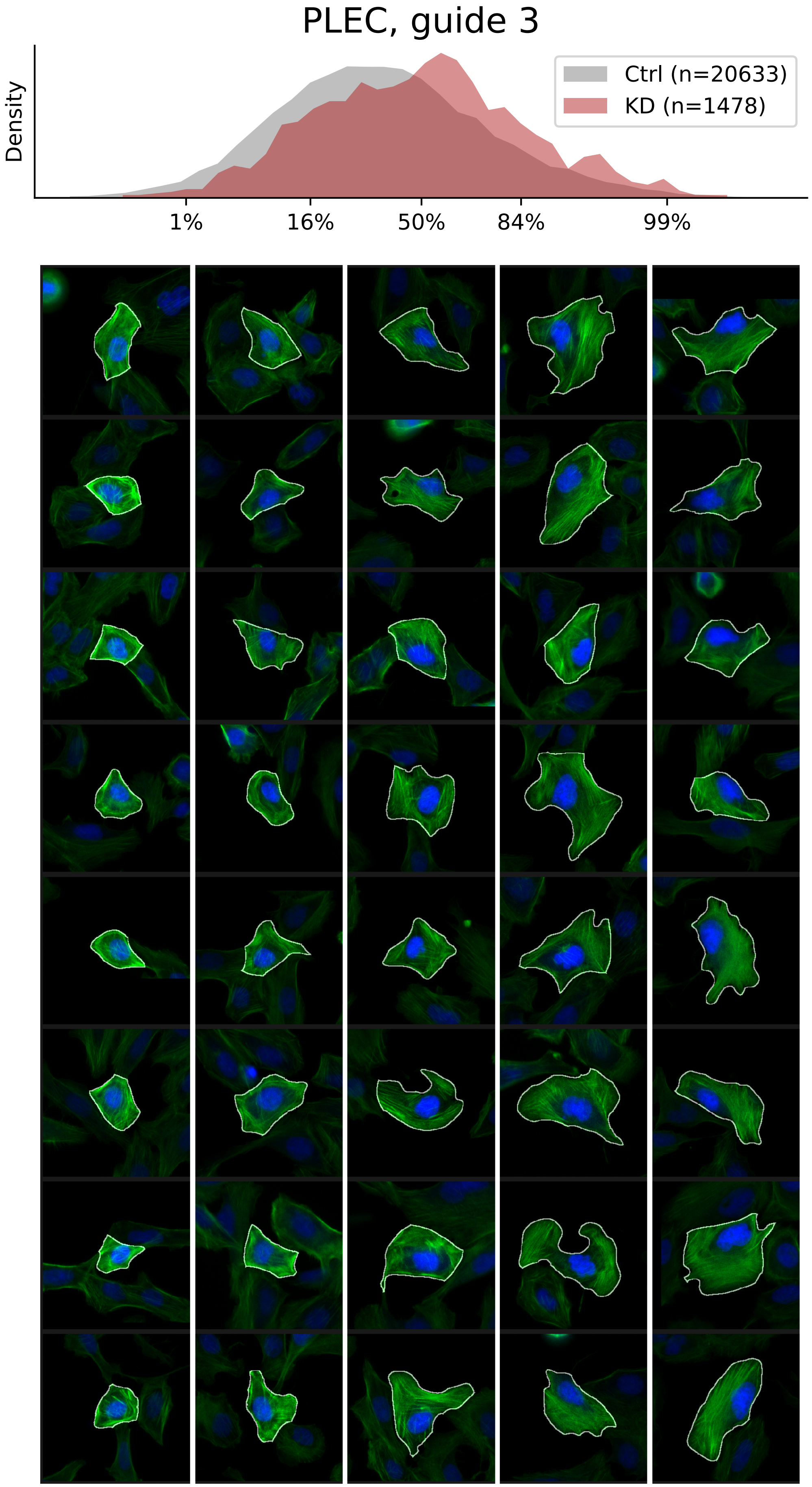

### VIEWS_fPC-LD-verified_SPTAN1_rep3.pdf

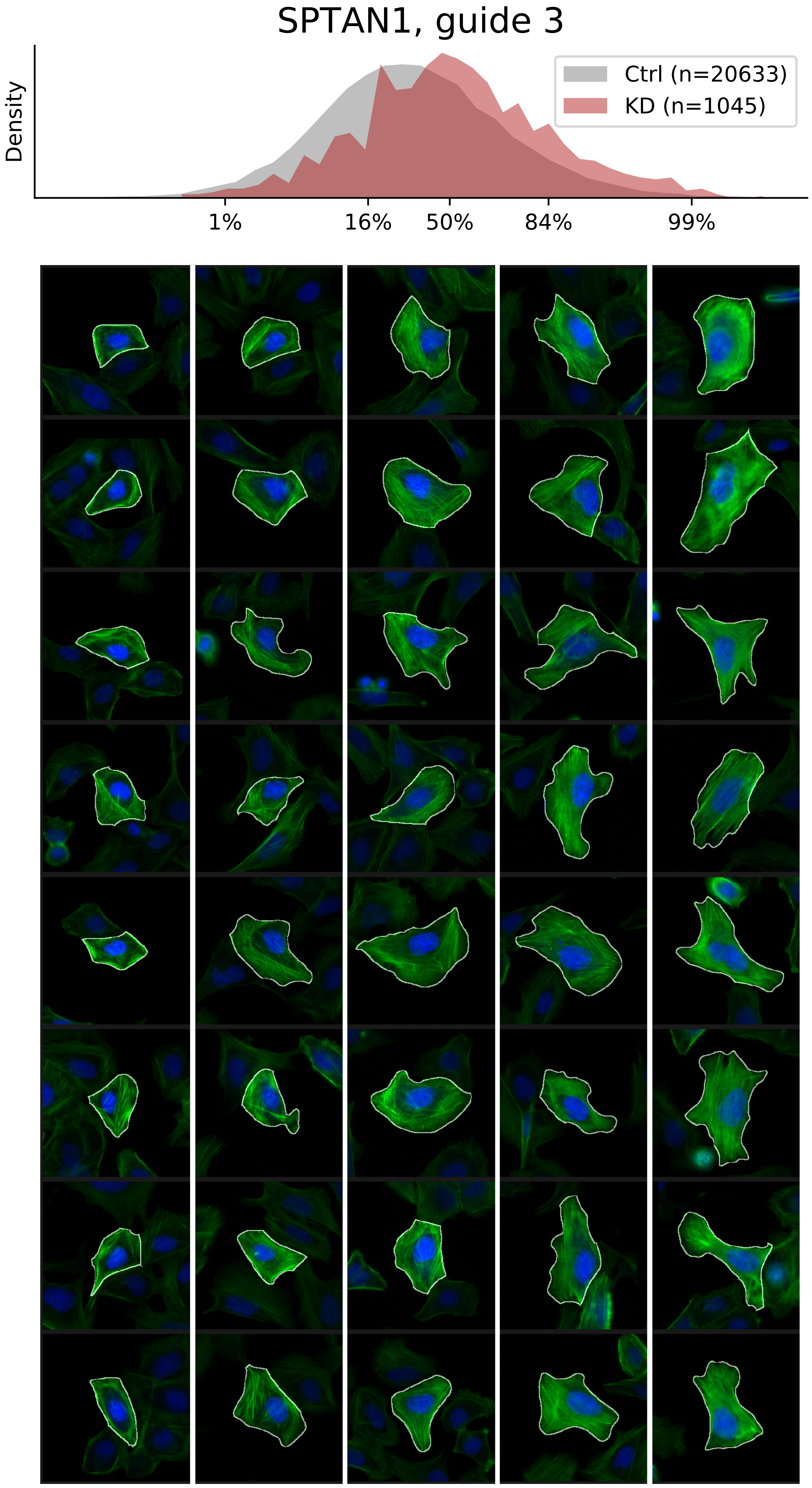

### VIEWS_fPC_ACTR2_rep1.pdf

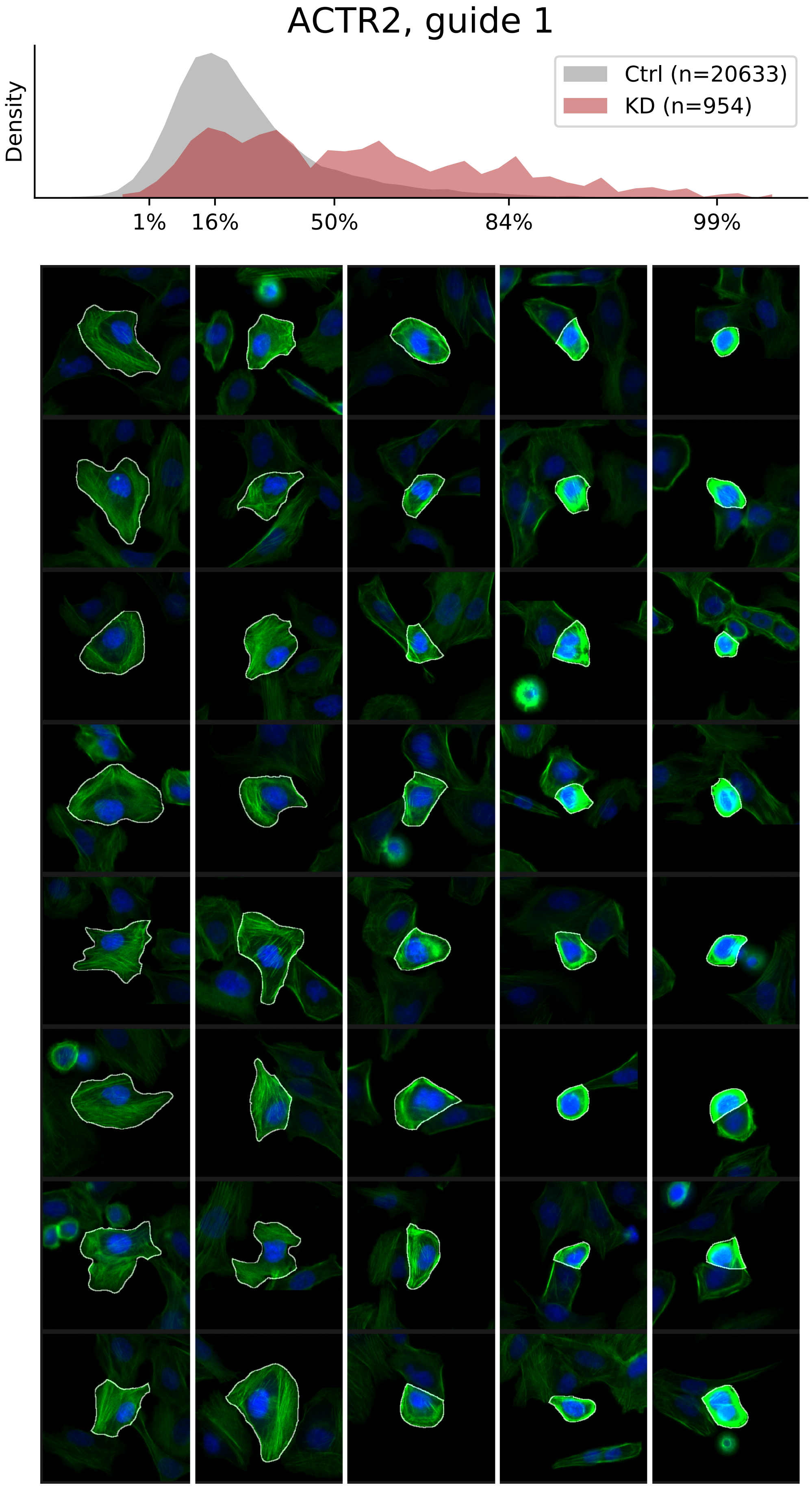

### VIEWS_fPC_FERMT2_rep2.pdf

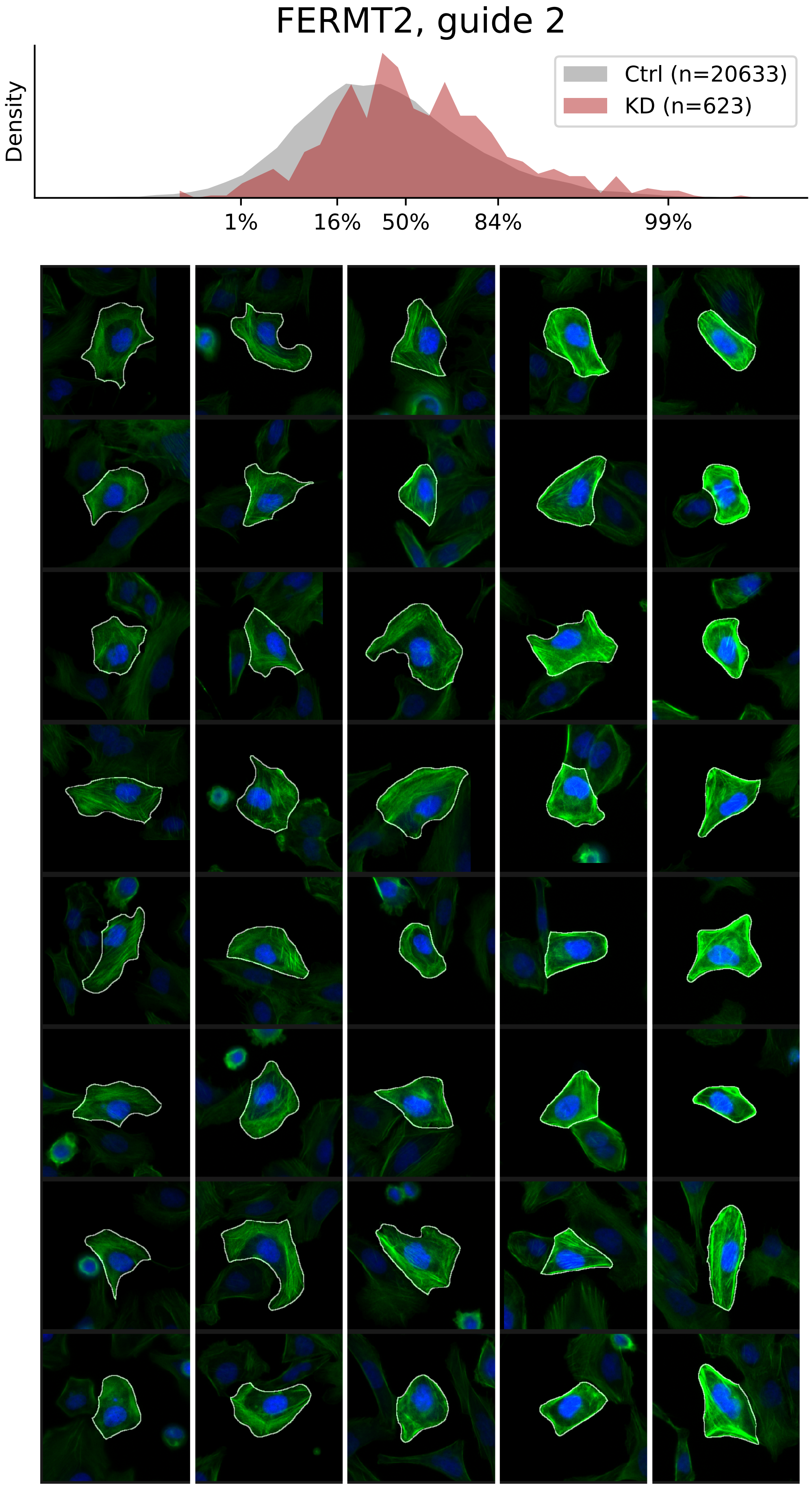

### VIEWS_fPC_FERMT2_rep3.pdf

#### FERMT2, guide 3

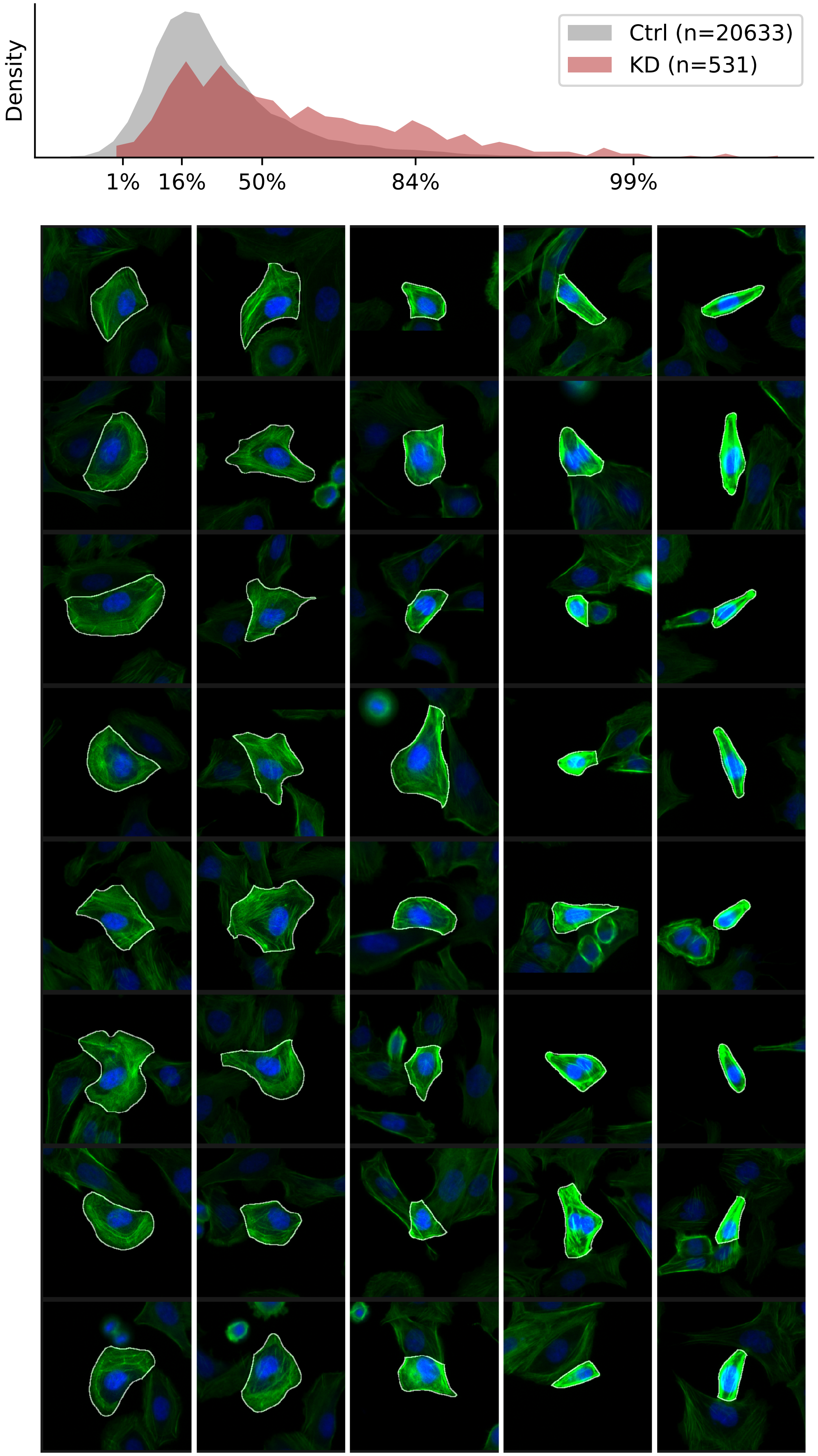

### VIEWS_fPC_GNA12_rep1.pdf

#### GNA12, guide 1

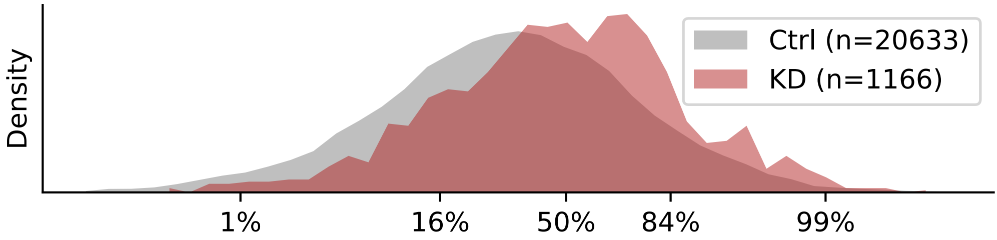


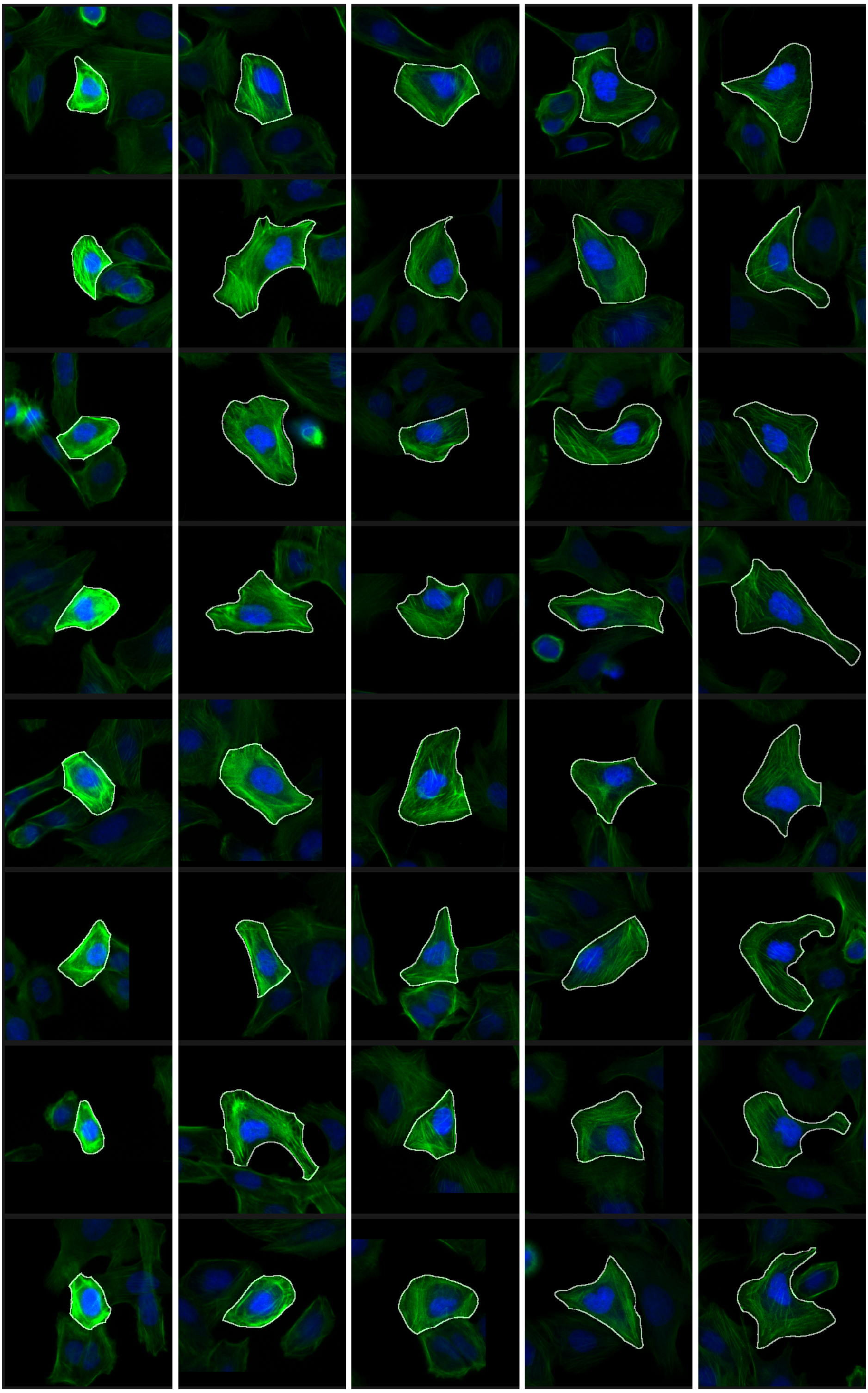

### VIEWS_fPC_ILK_rep1.pdf

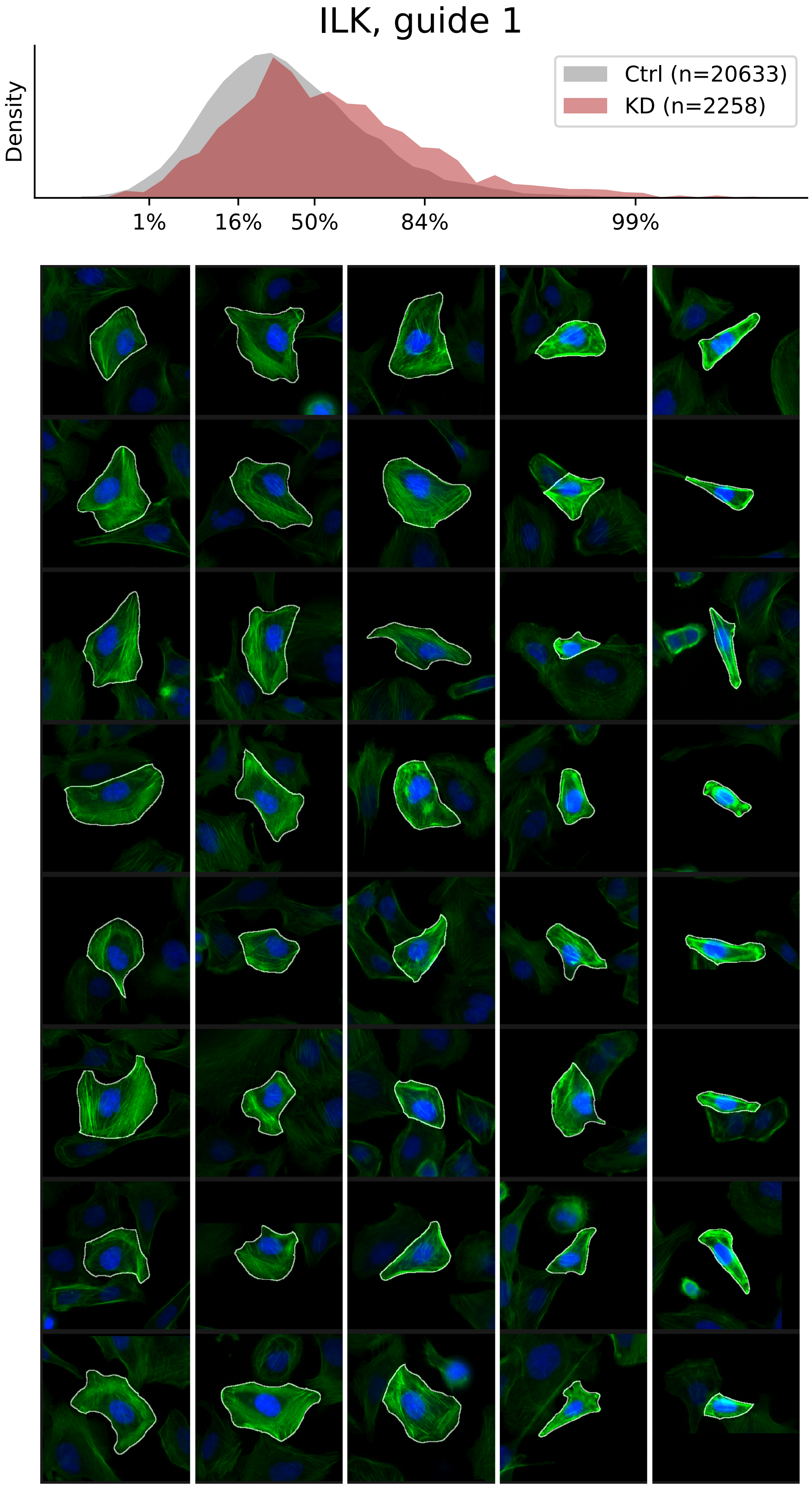

### VIEWS_fPC_ITGB1_rep2.pdf

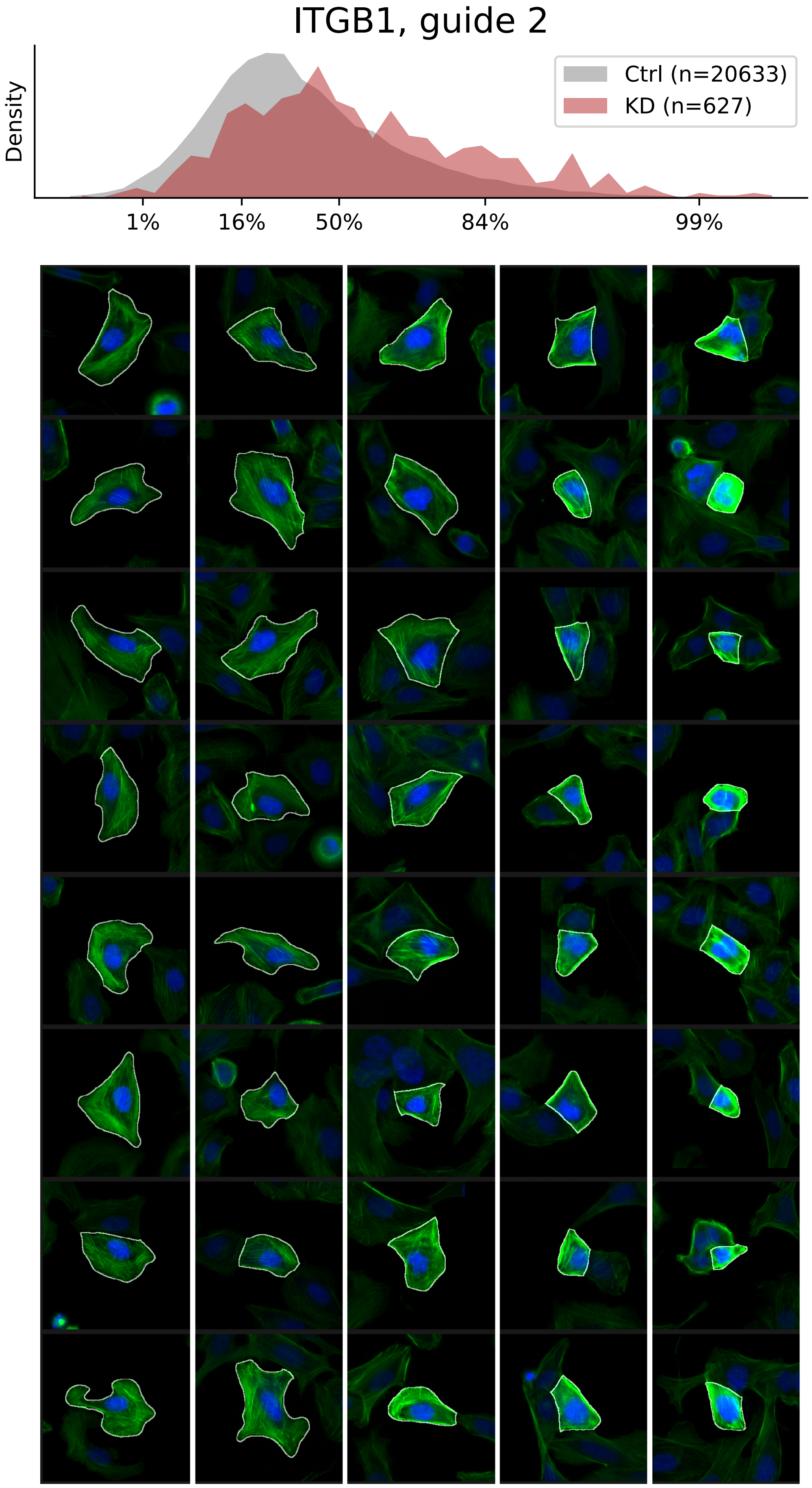

### VIEWS_fPC_ITGB1_rep3.pdf

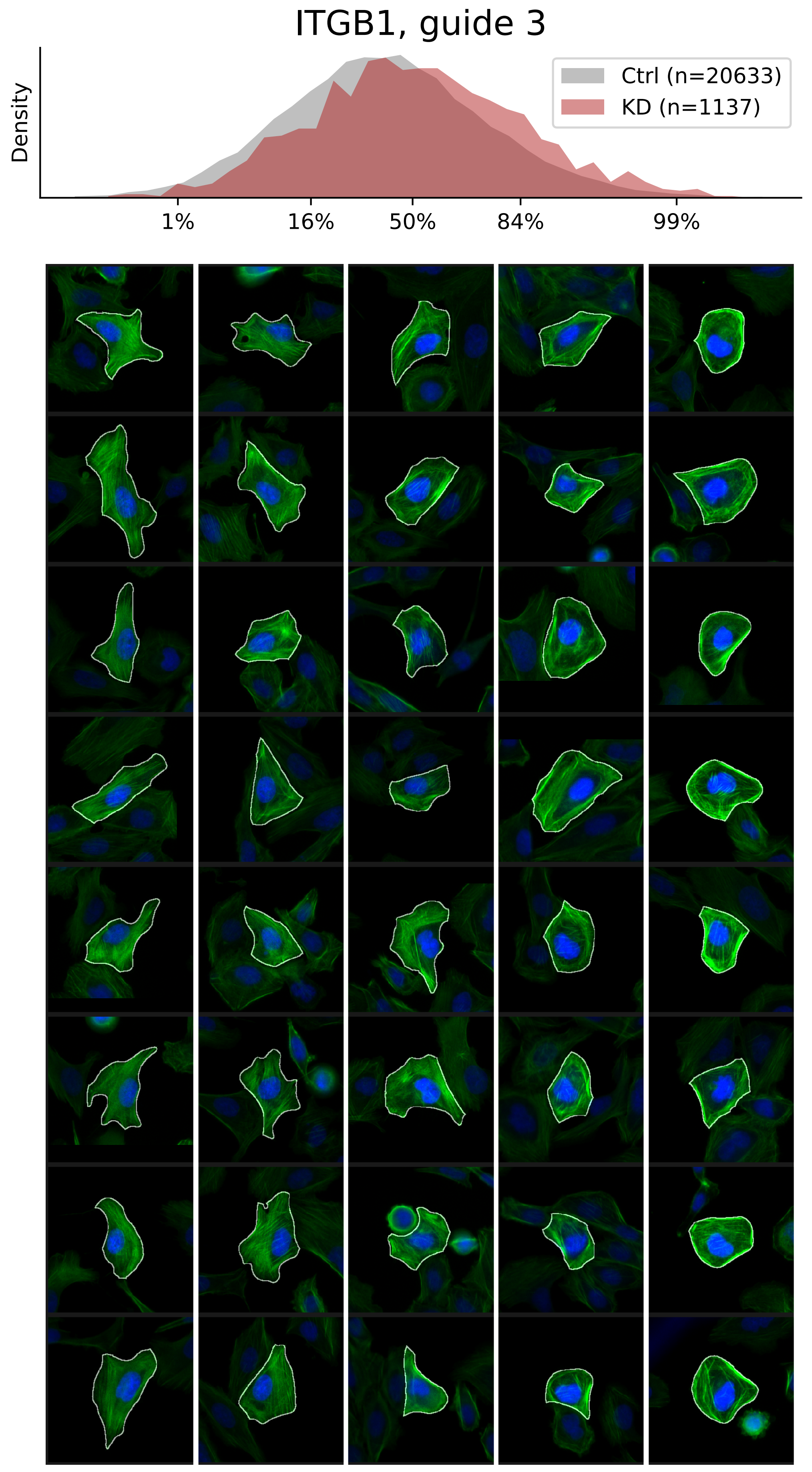

### VIEWS_fPC_LIMK2_rep1.pdf

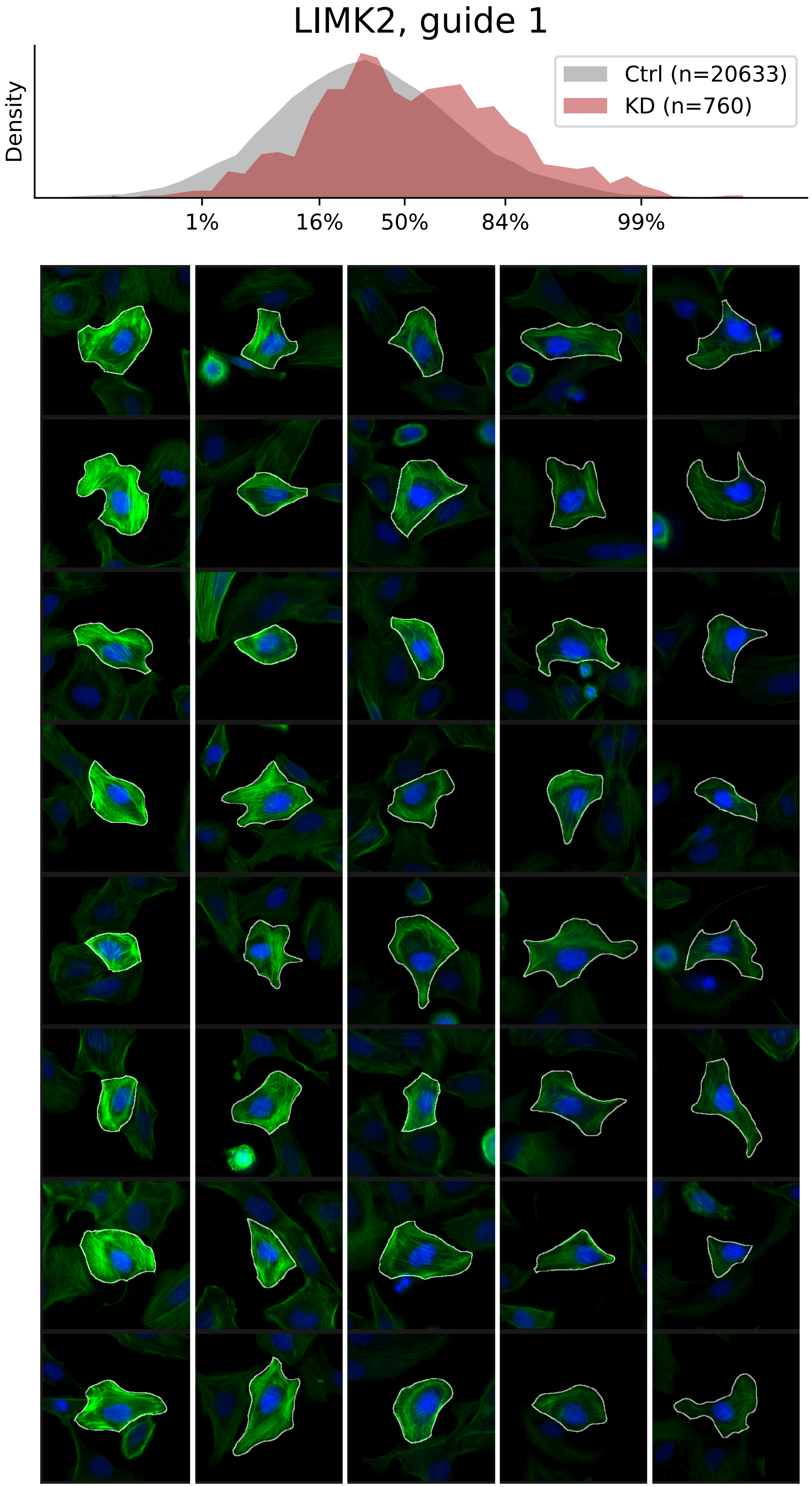

### VIEWS_fPC_LIMK2_rep2.pdf

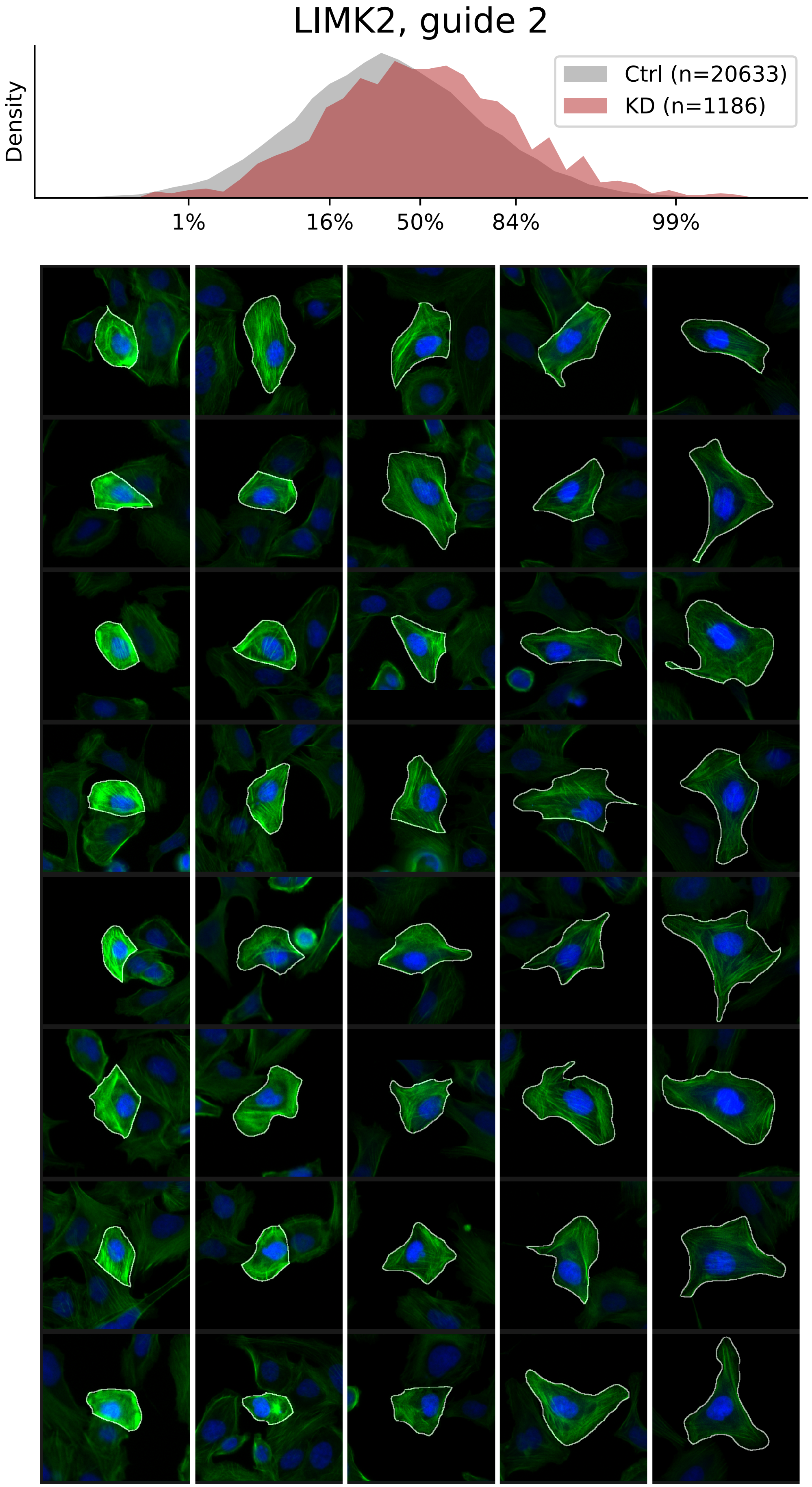

### VIEWS_fPC_LIMK2_rep3.pdf

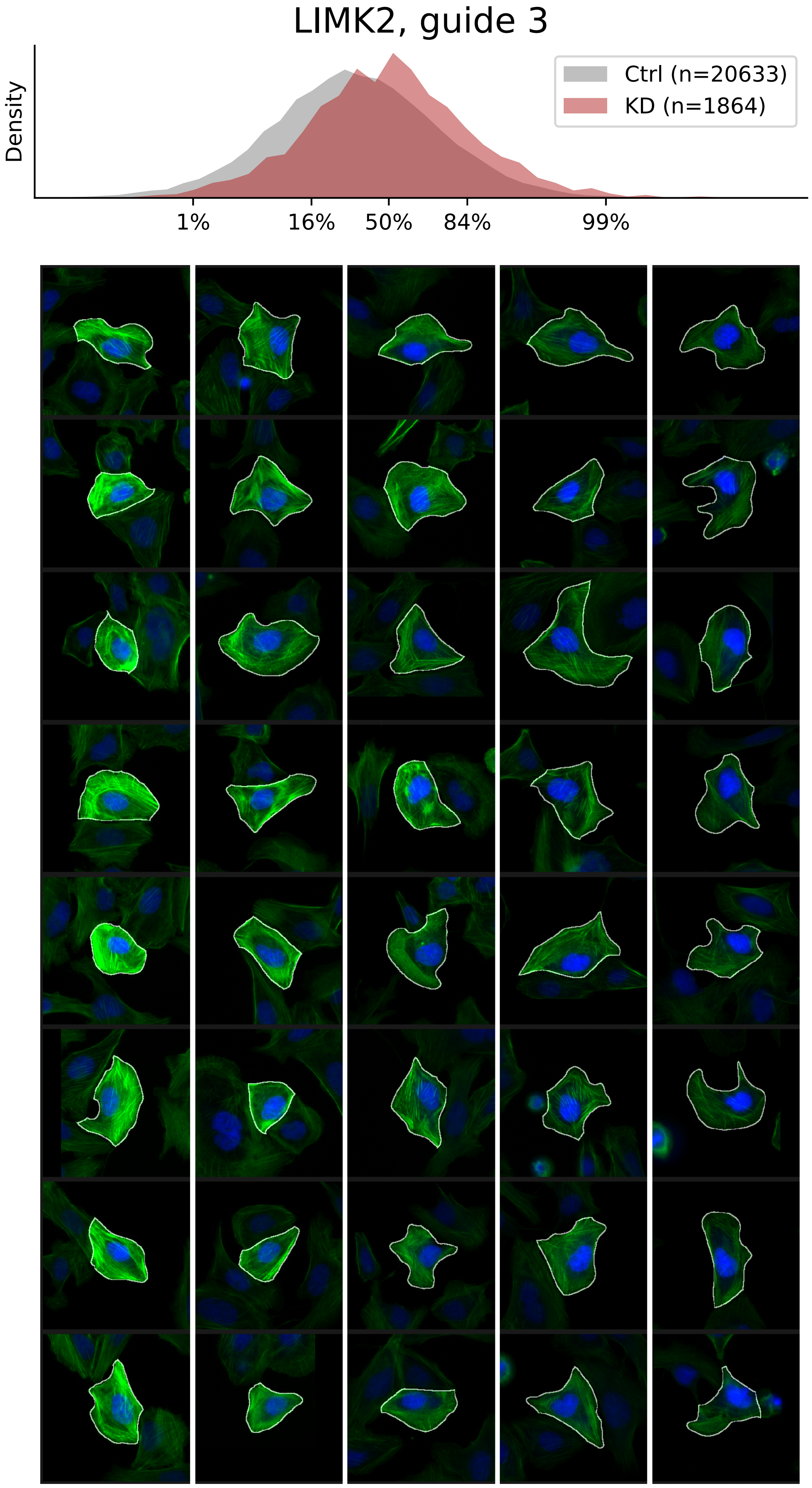

### VIEWS_fPC_MTOR_rep1.pdf

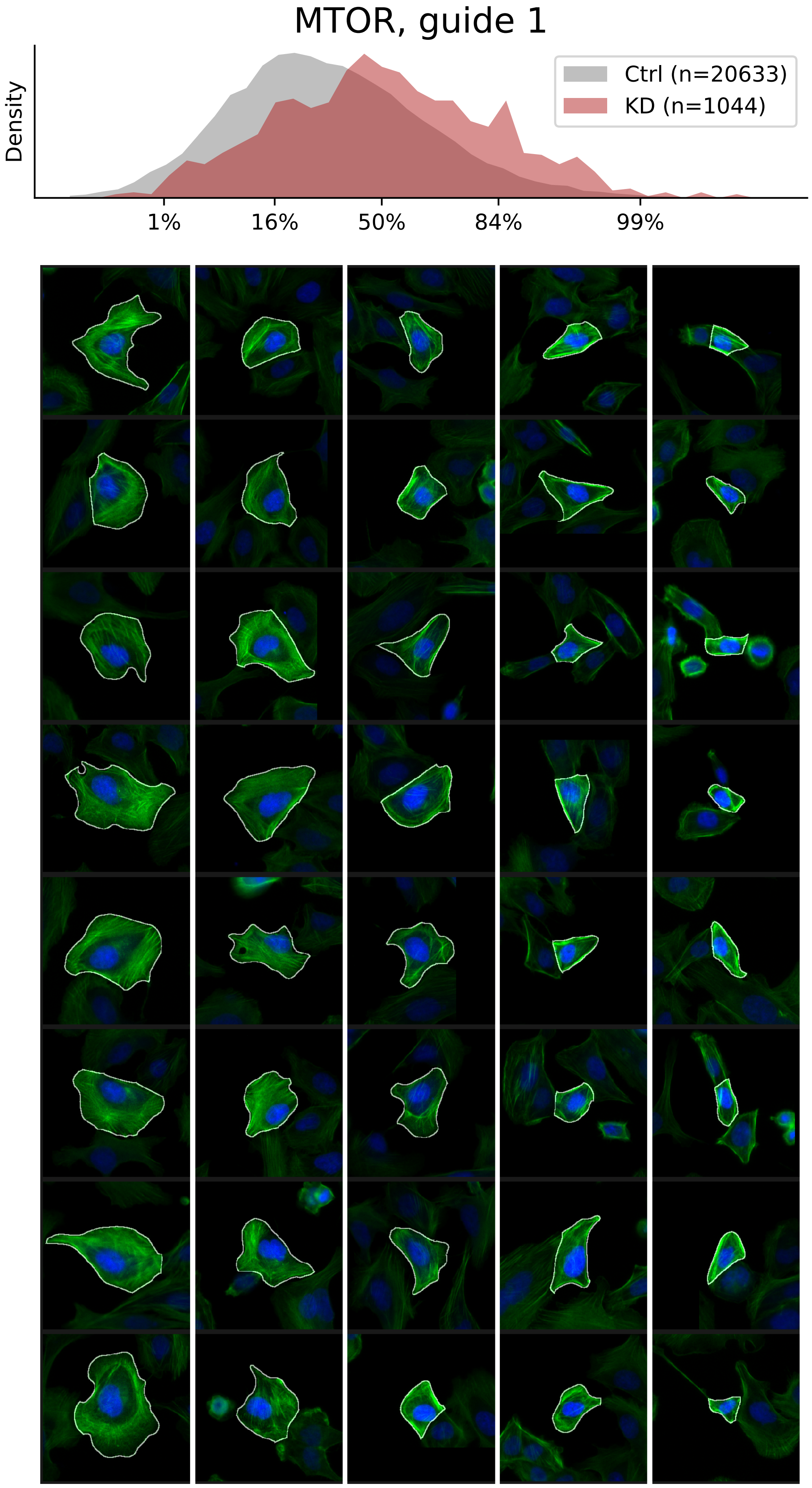

### VIEWS_fPC_MTOR_rep2.pdf

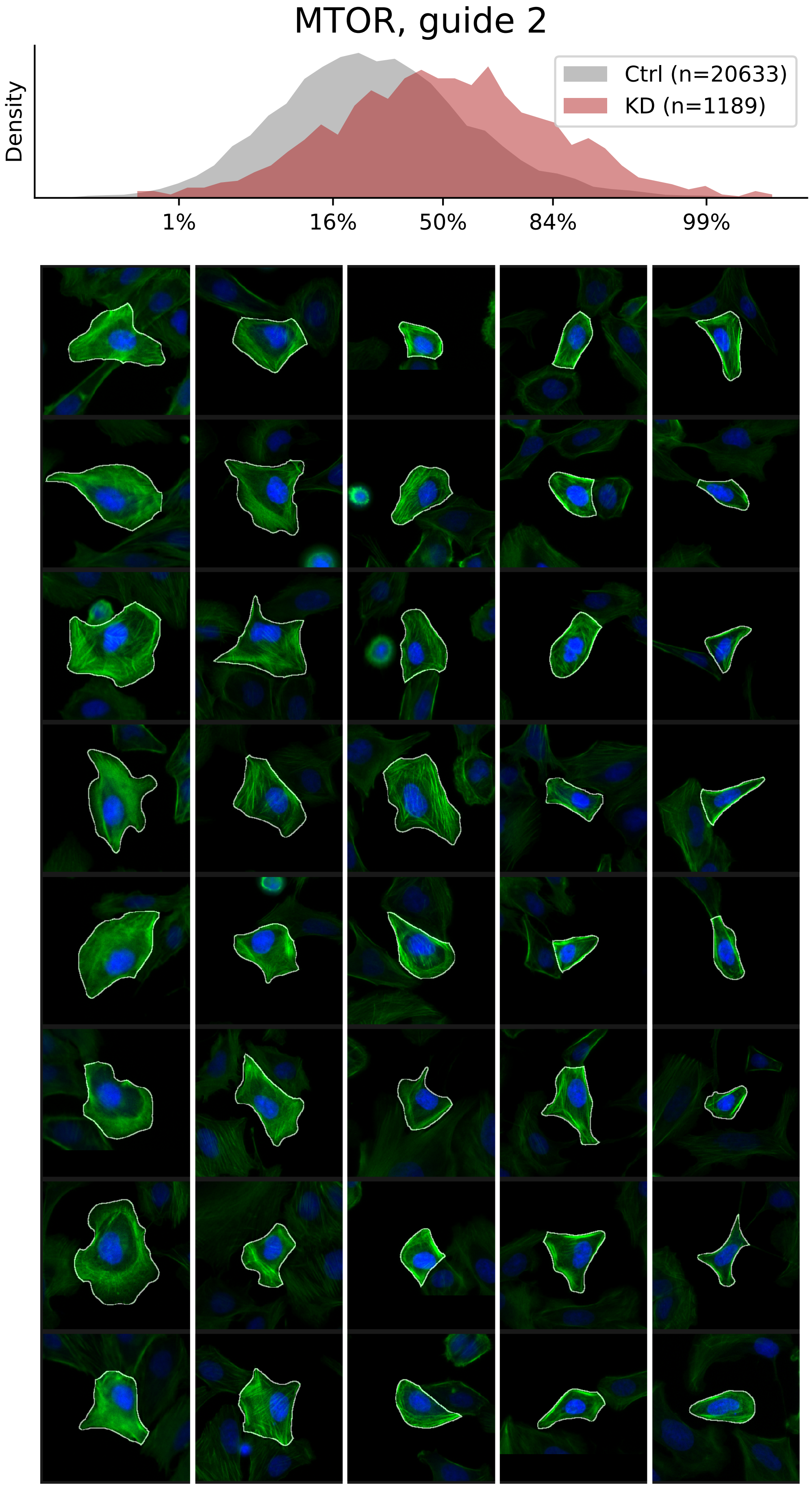

### VIEWS_fPC_MTOR_rep3.pdf

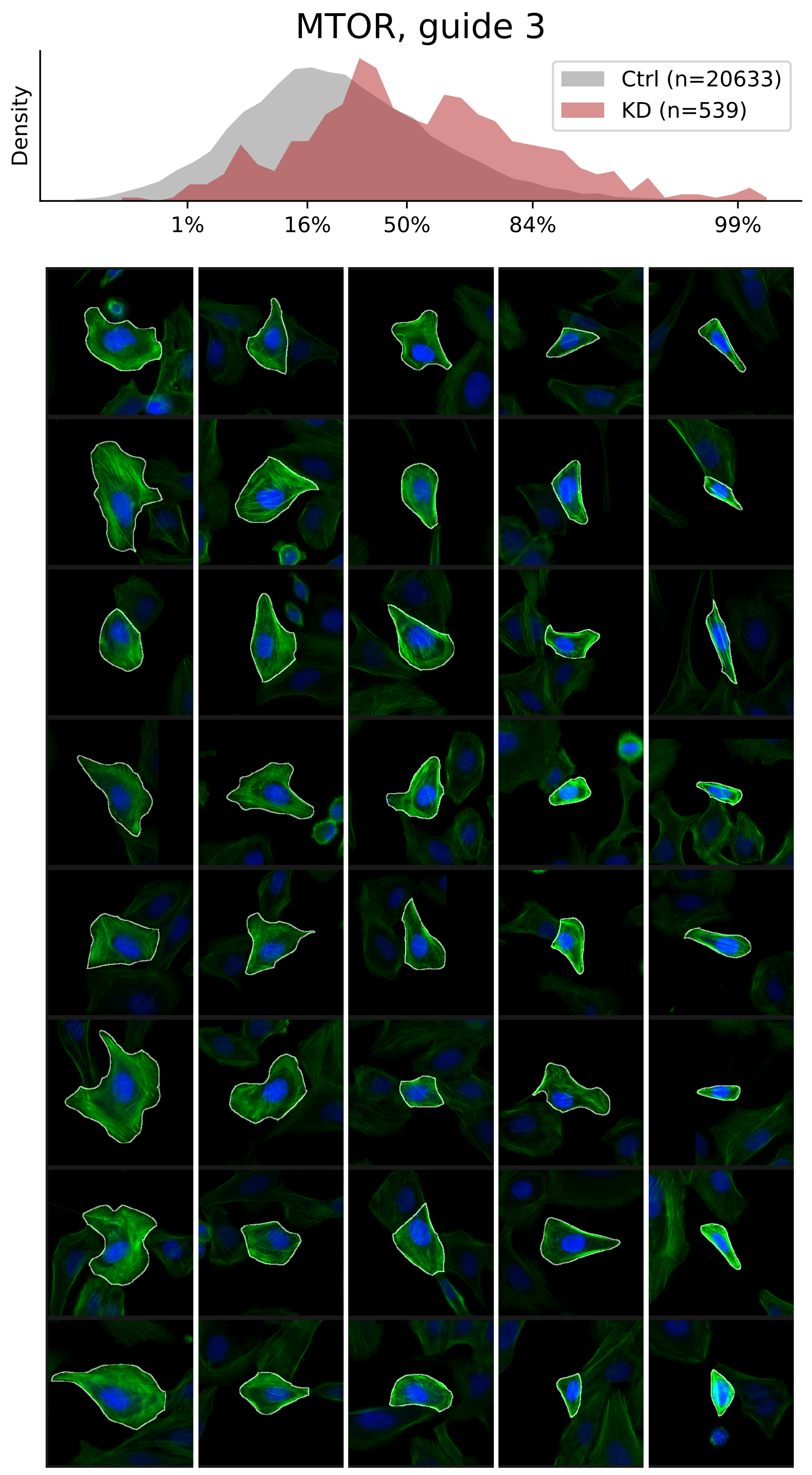

### VIEWS_fPC_MYH9_rep1.pdf

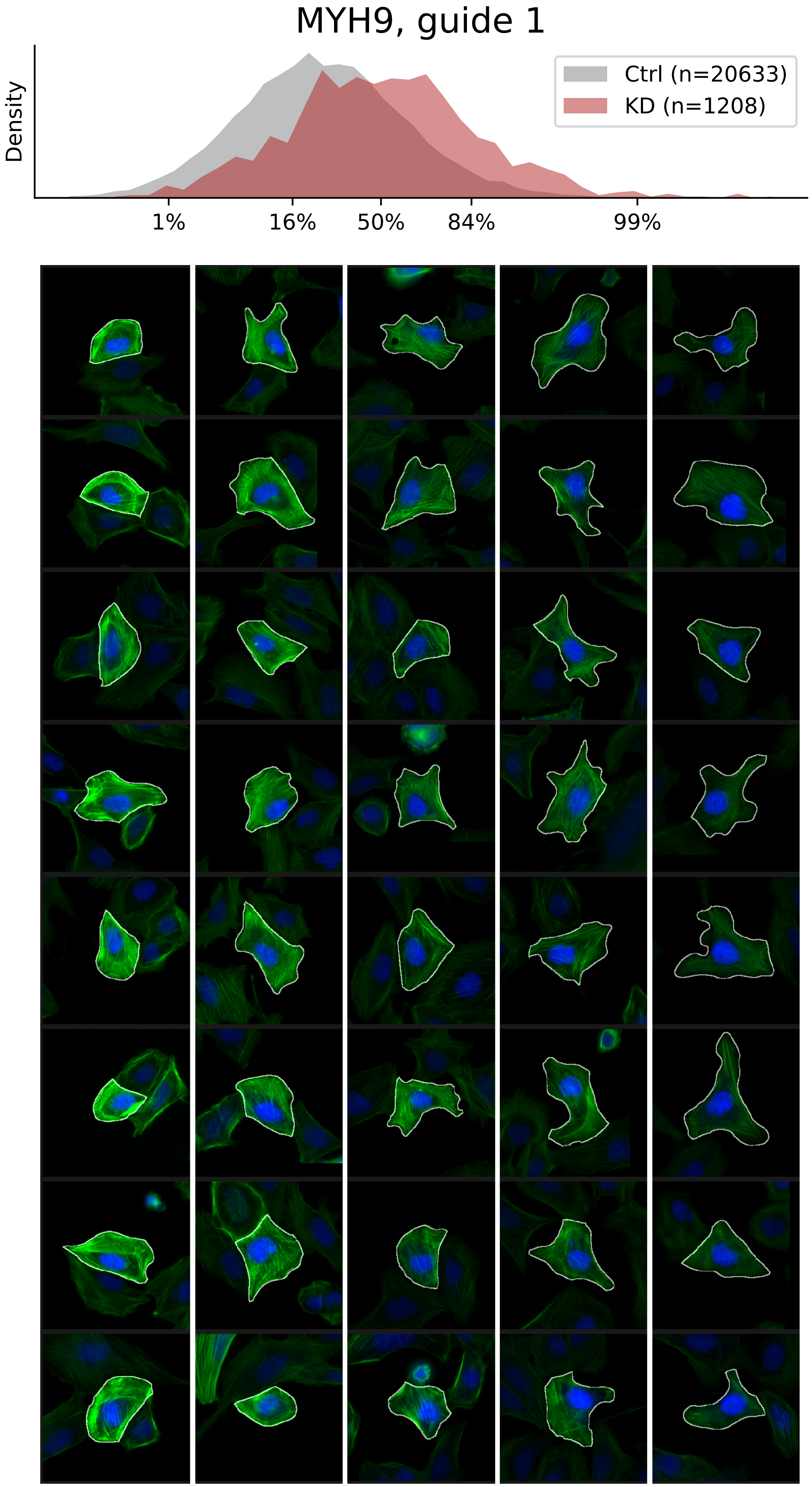

### VIEWS_fPC_MYH9_rep2.pdf

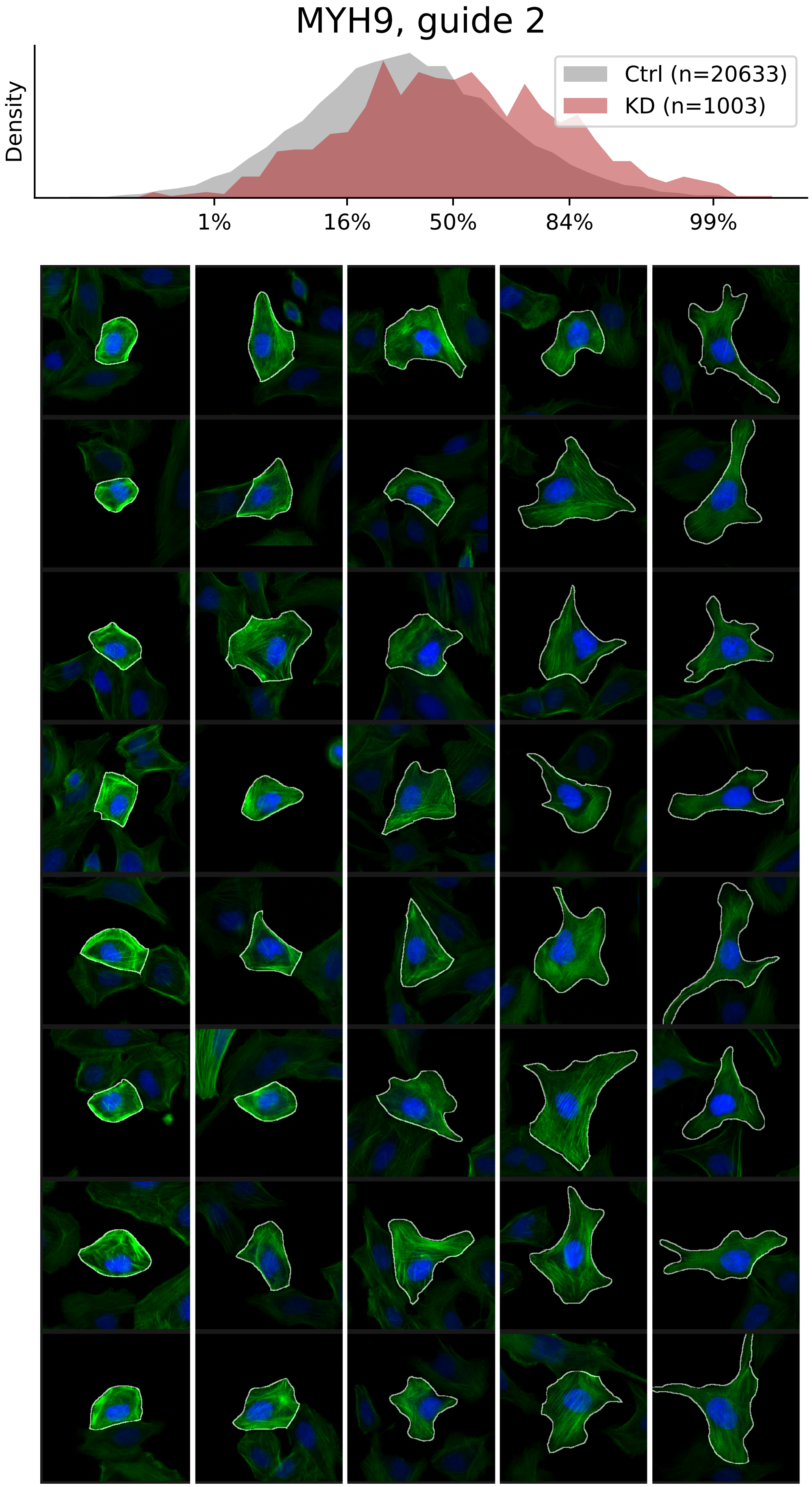

### VIEWS_fPC_MYH9_rep3.pdf

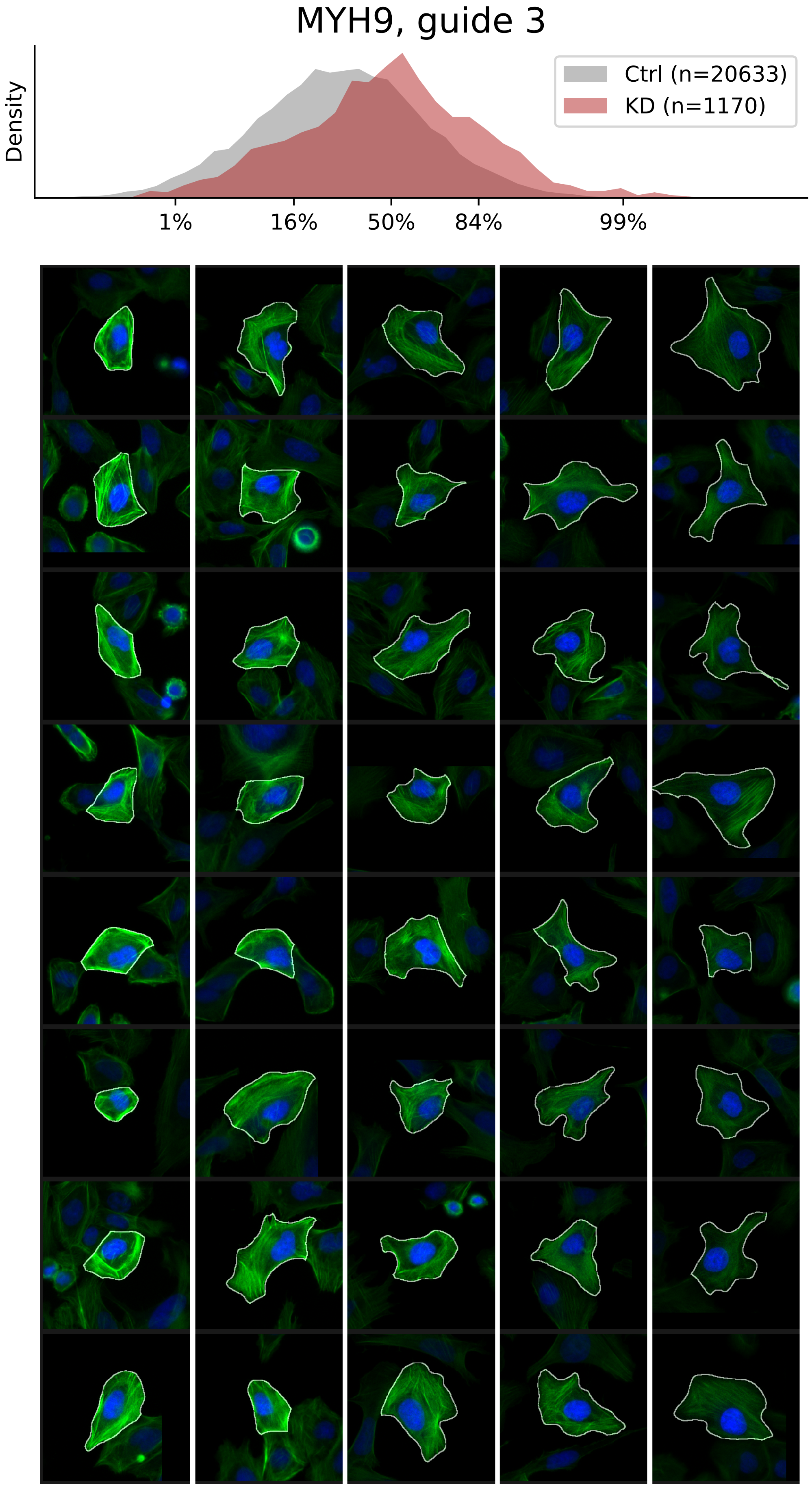

### VIEWS_fPC_MYL6_rep1.pdf

##### MYL6, guide 1

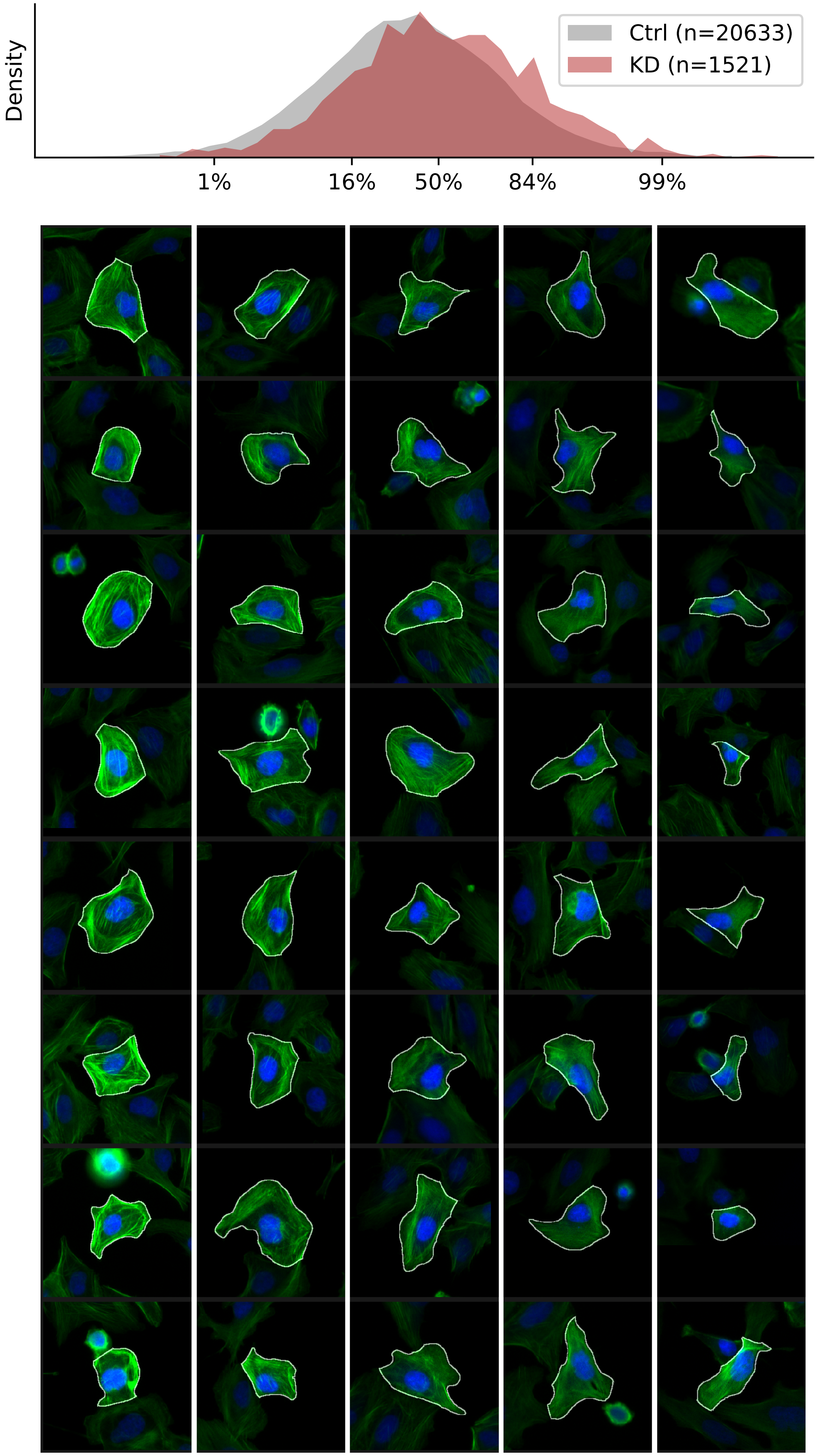

### VIEWS_fPC_MYL6_rep3.pdf

#### MYL6, guide 3

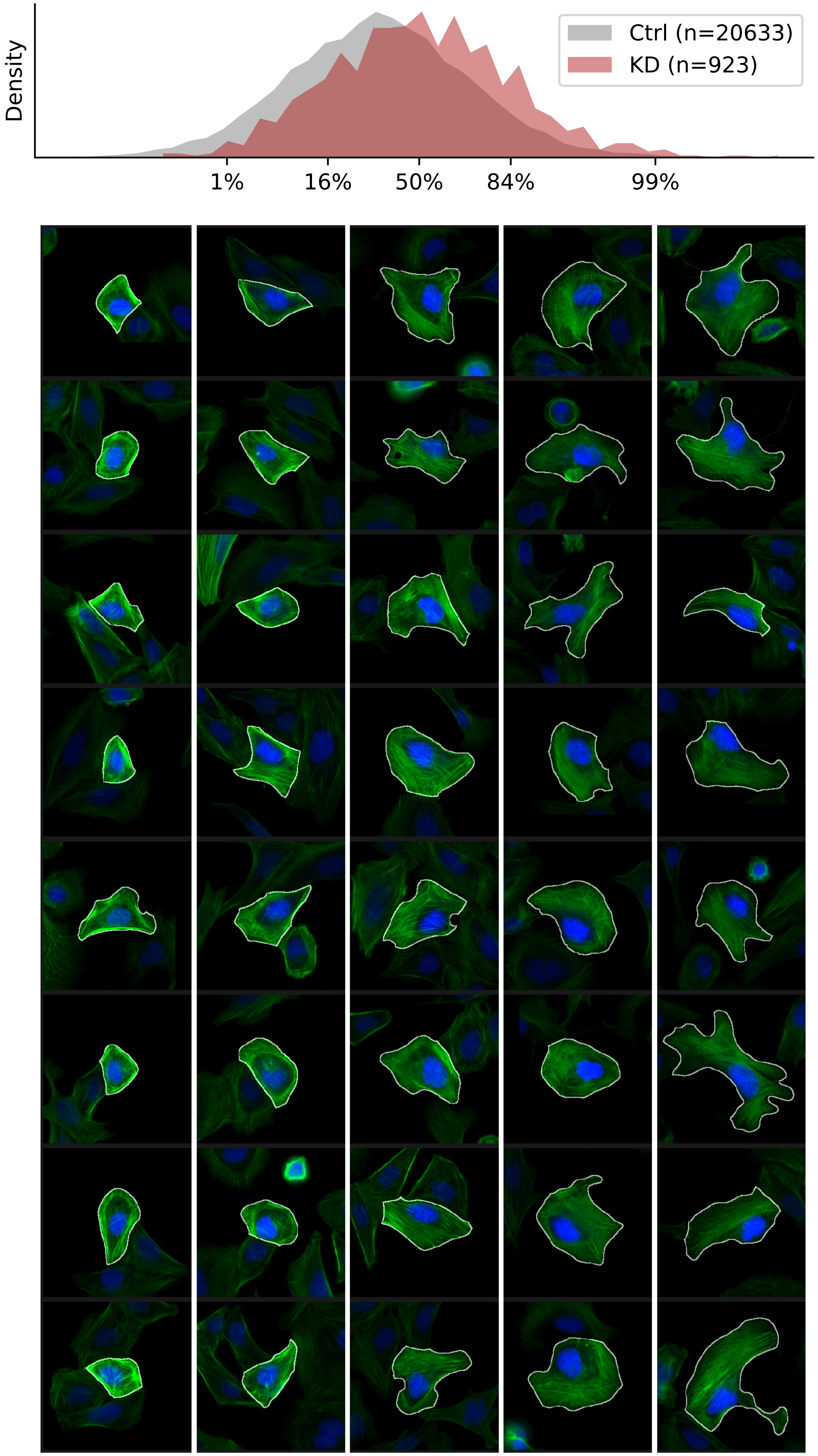

### VIEWS_fPC_PIP5K1A_rep1.pdf

PIP5K1A, guide 1

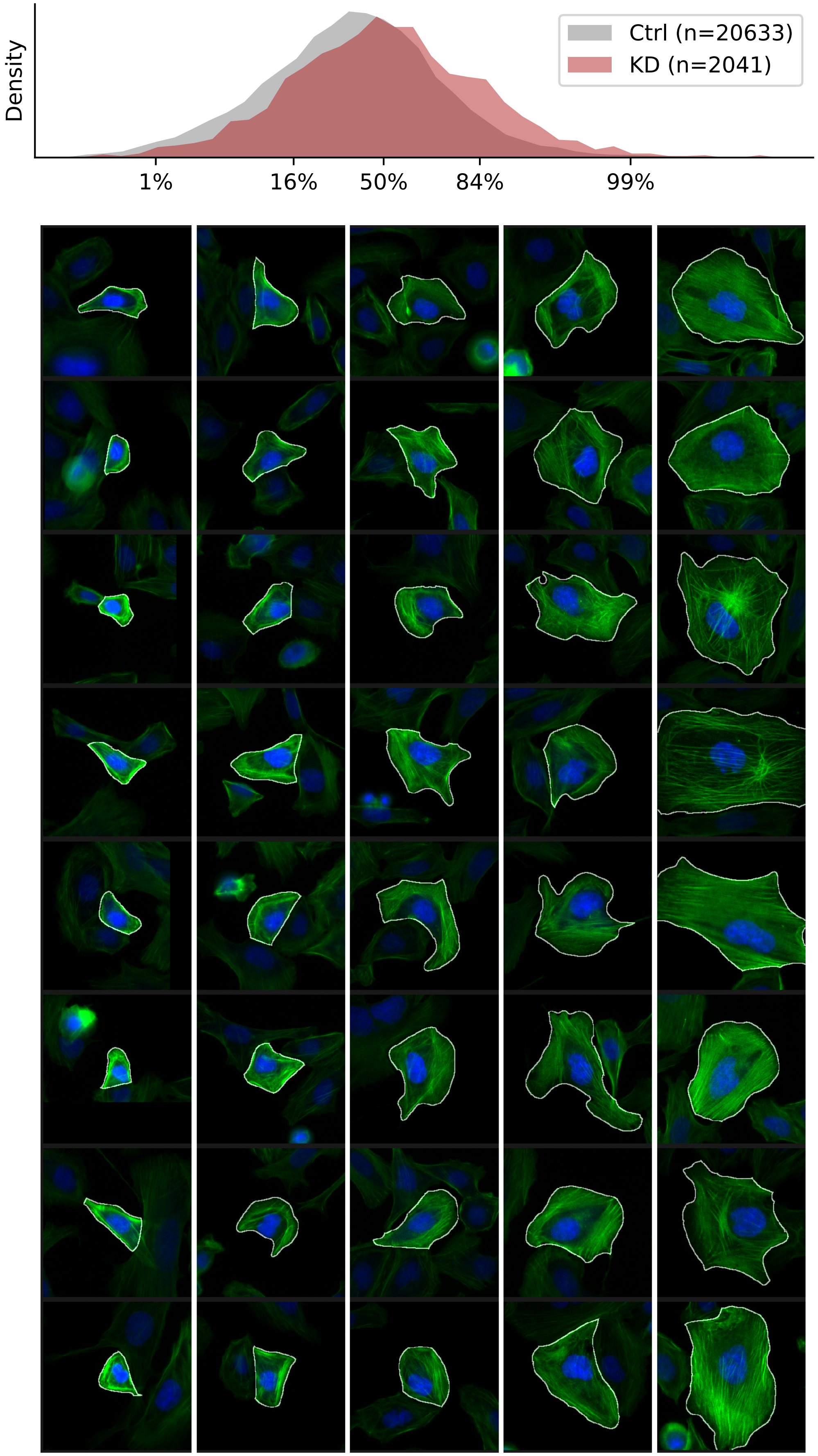

### VIEWS_fPC_PIP5K1A_rep2.pdf

#### PIP5K1A, guide 2

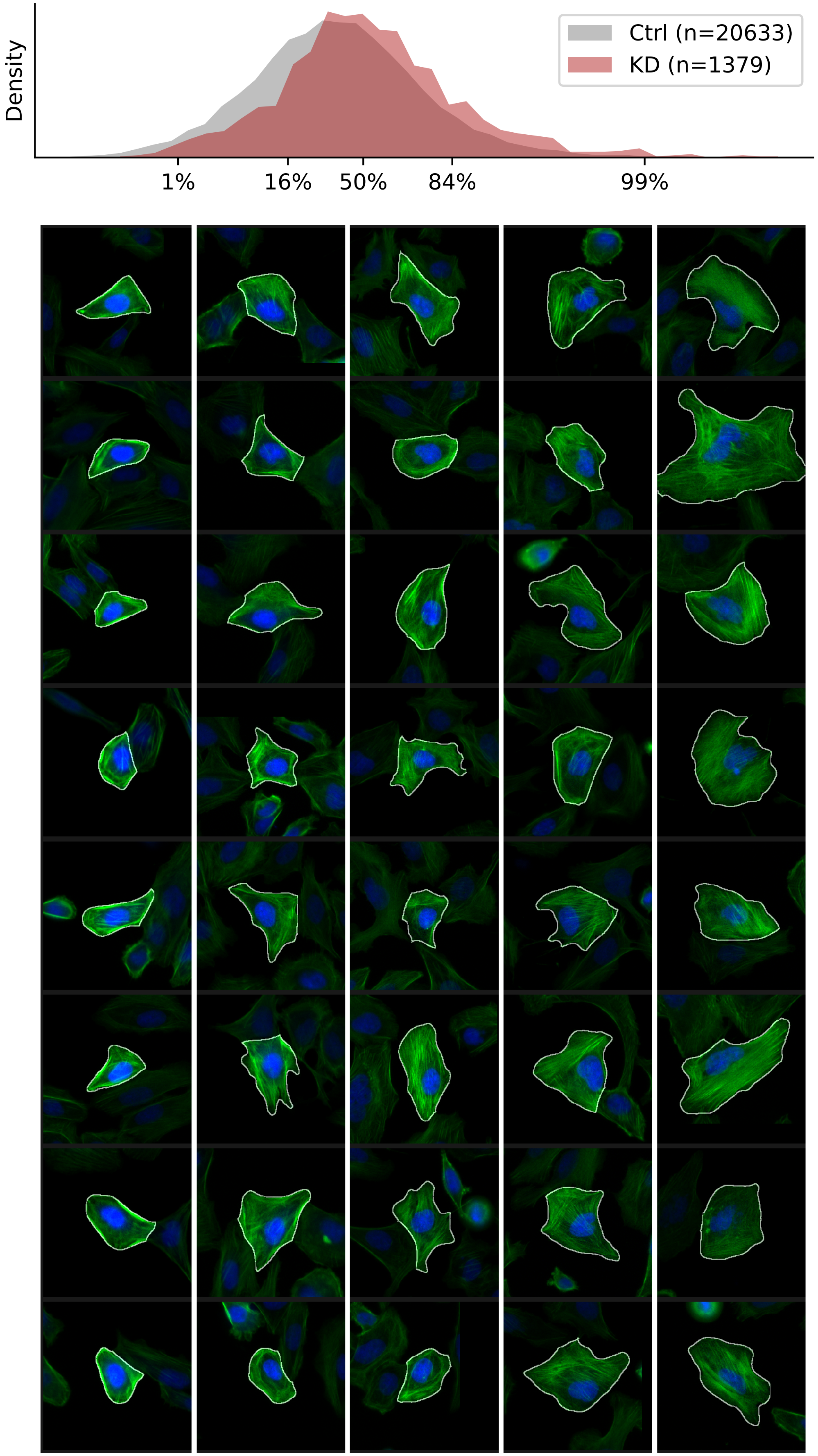

### VIEWS_fPC_PIP5K1A_rep3.pdf

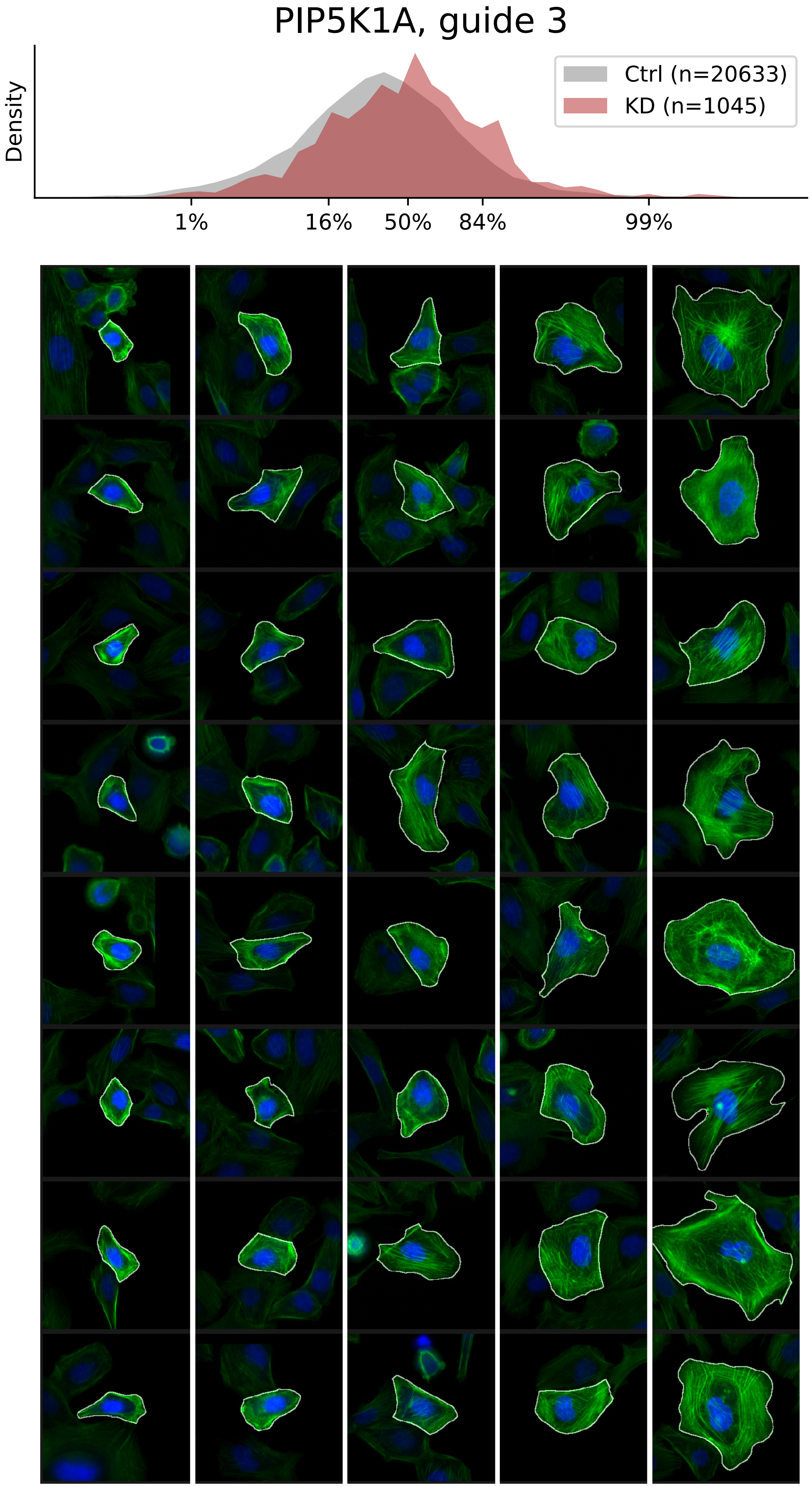

### VIEWS_fPC_PTK2_rep1.pdf

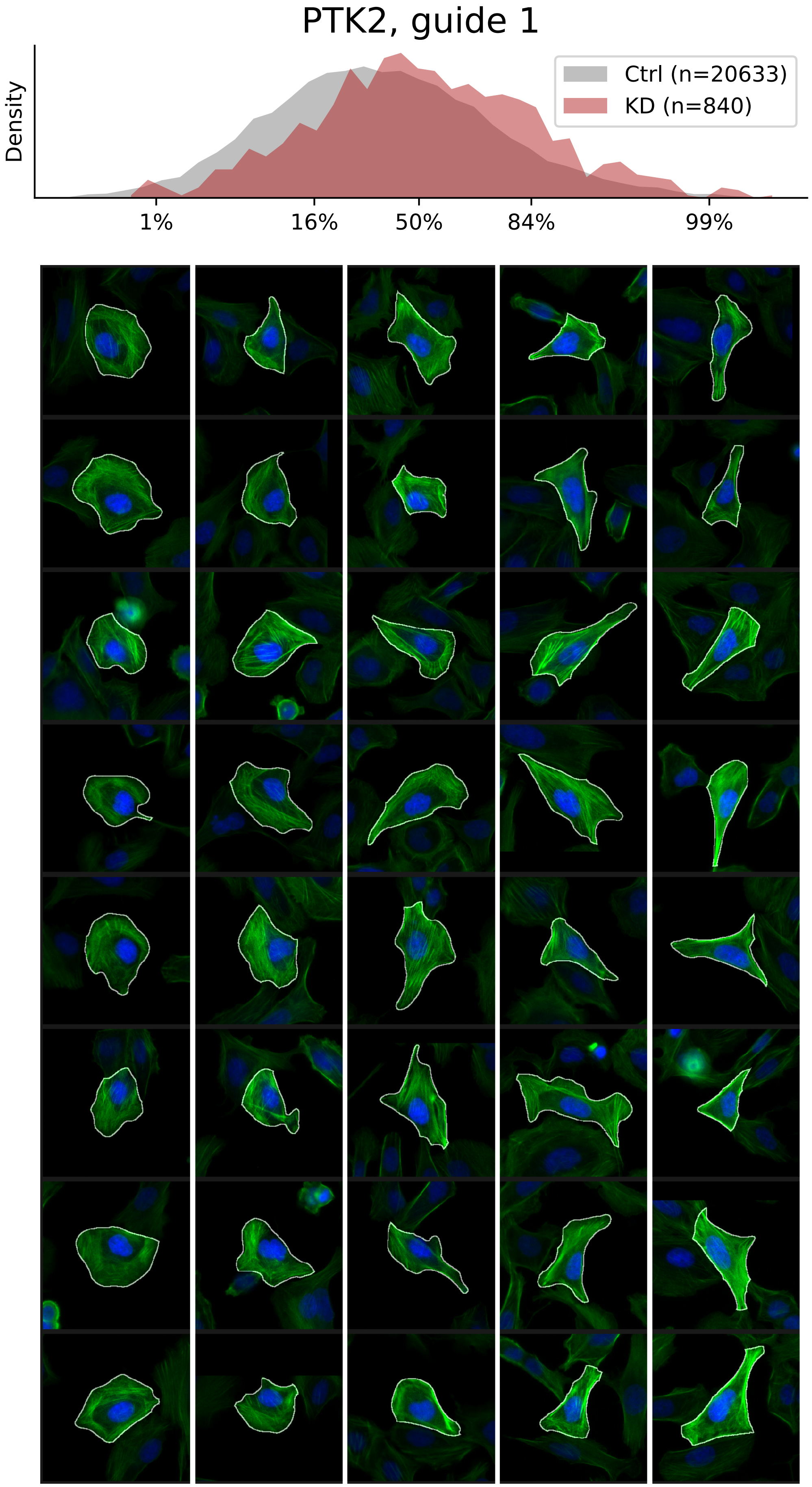

### VIEWS_fPC_PTK2_rep3.pdf

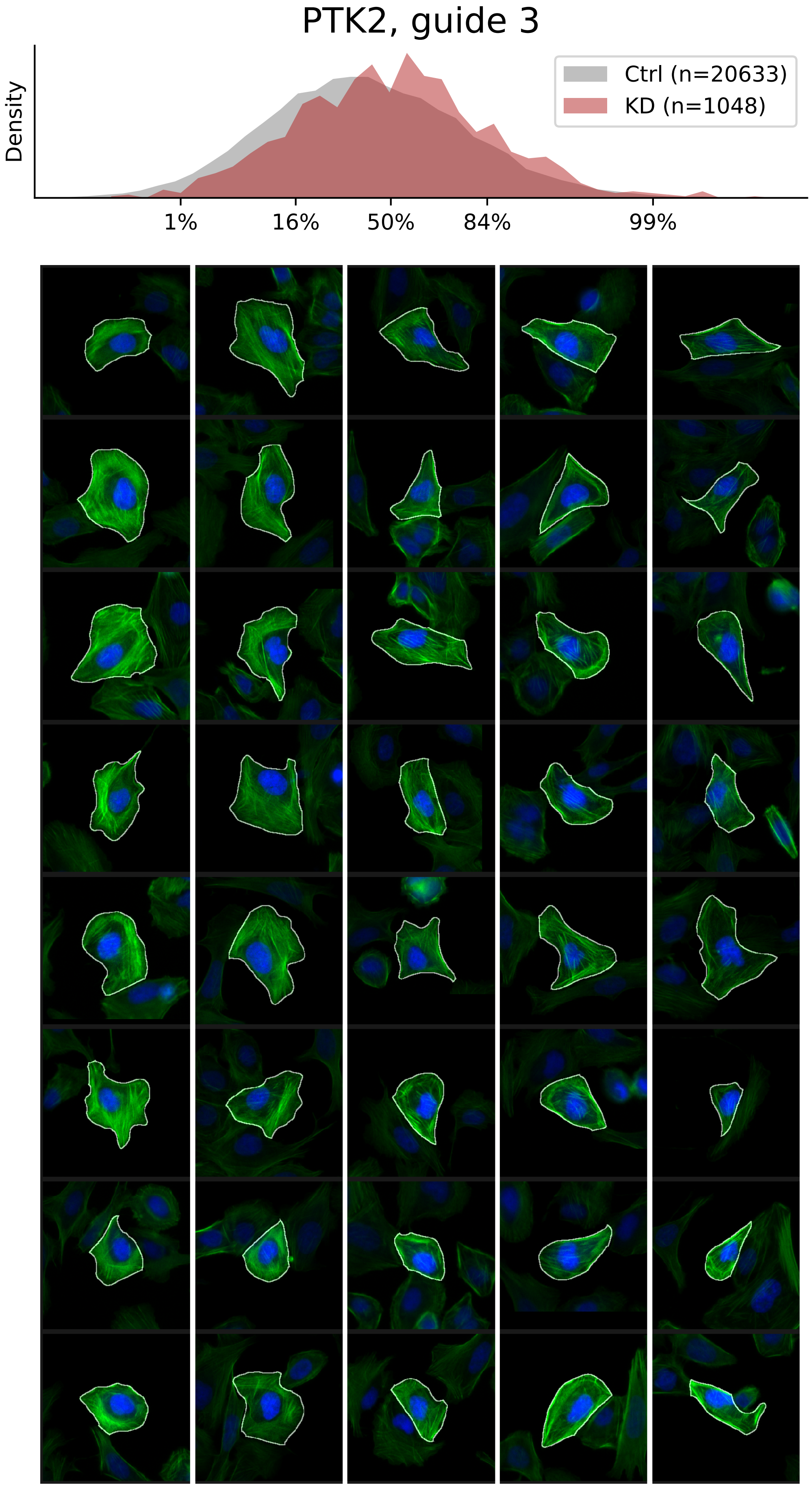

### VIEWS_fPC_SEPT7_rep1.pdf

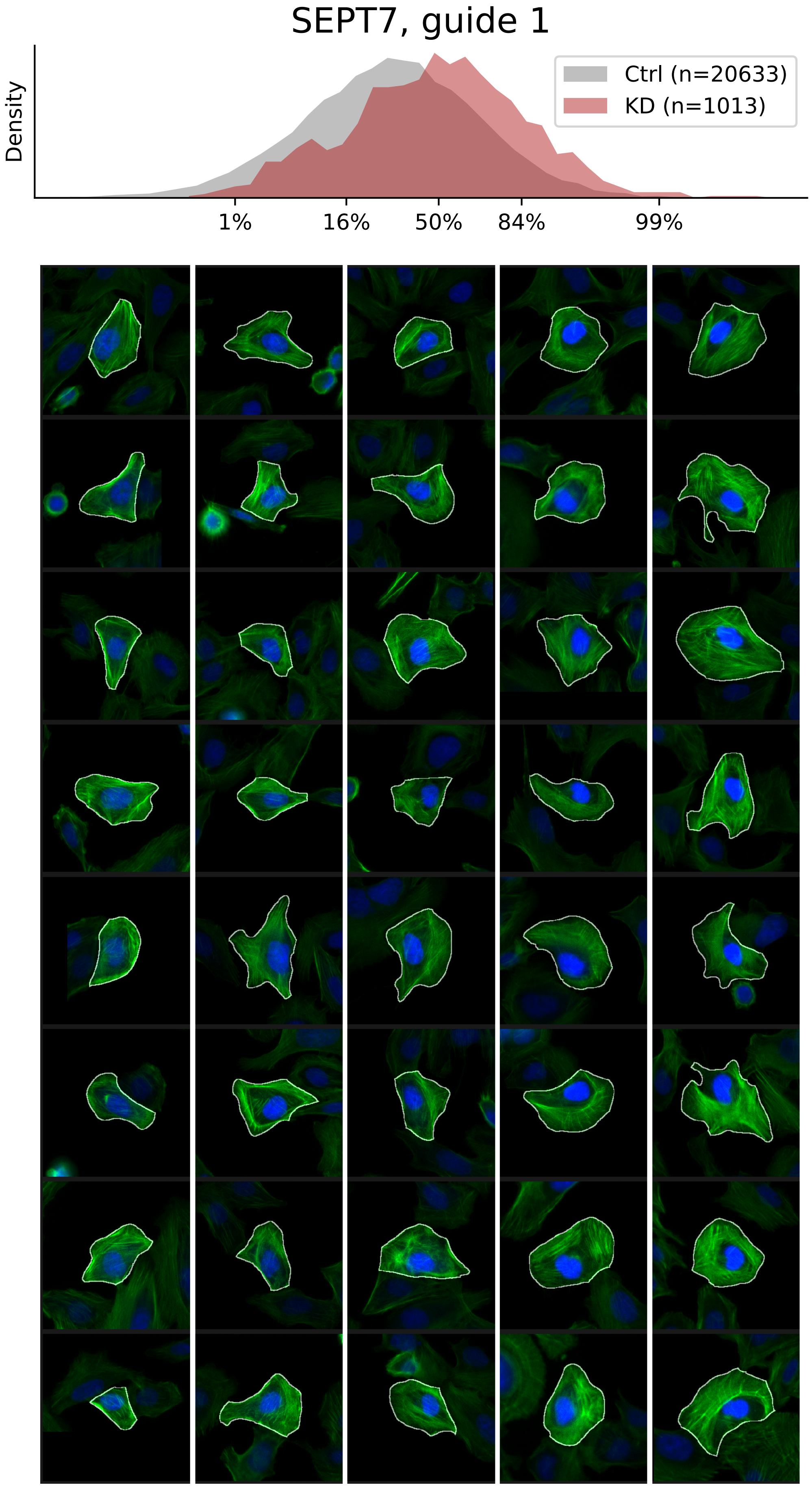

### VIEWS_fPC_SEPT7_rep2.pdf

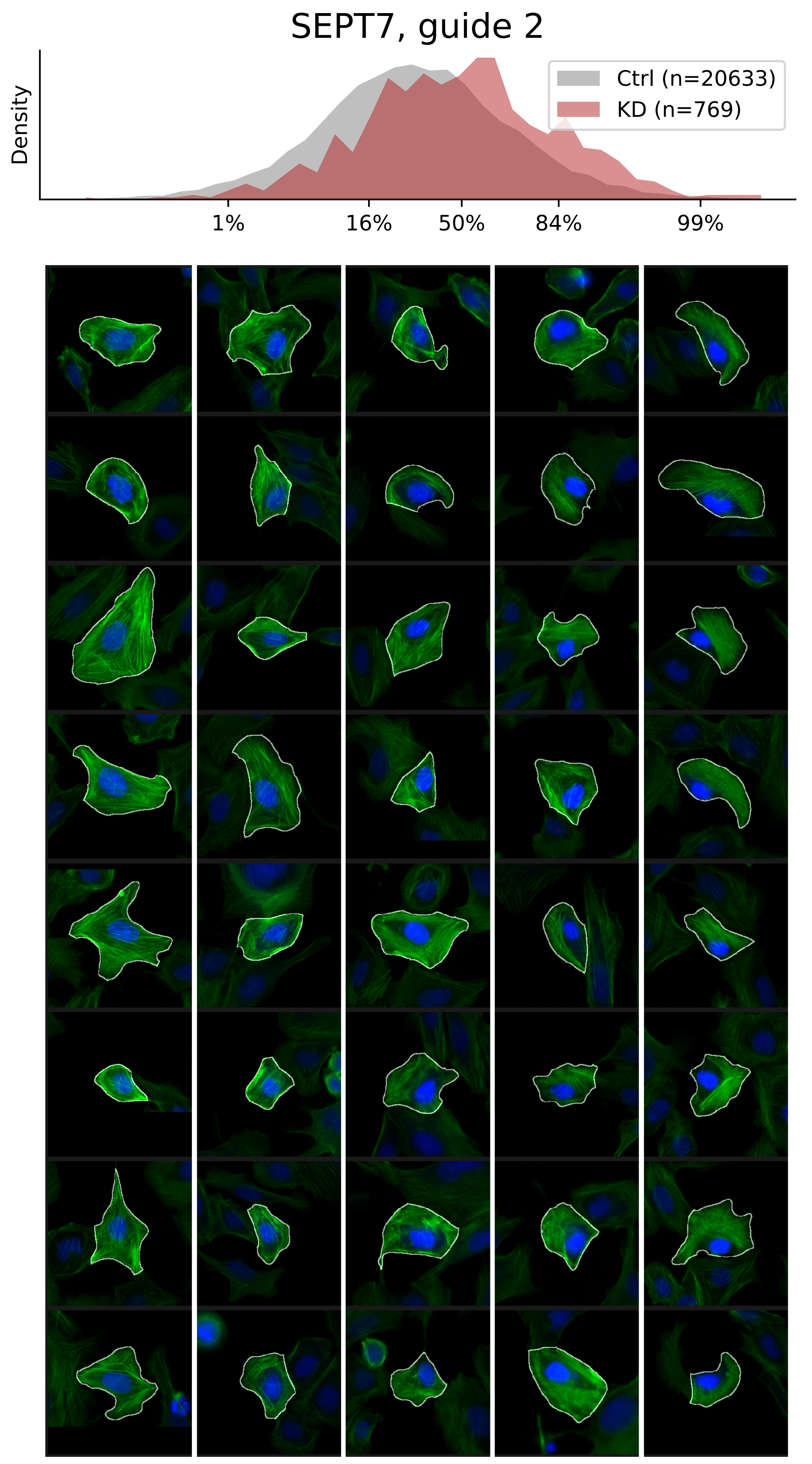

### VIEWS_fPC_SEPT7_rep3.pdf

#### SEPT7, guide 3

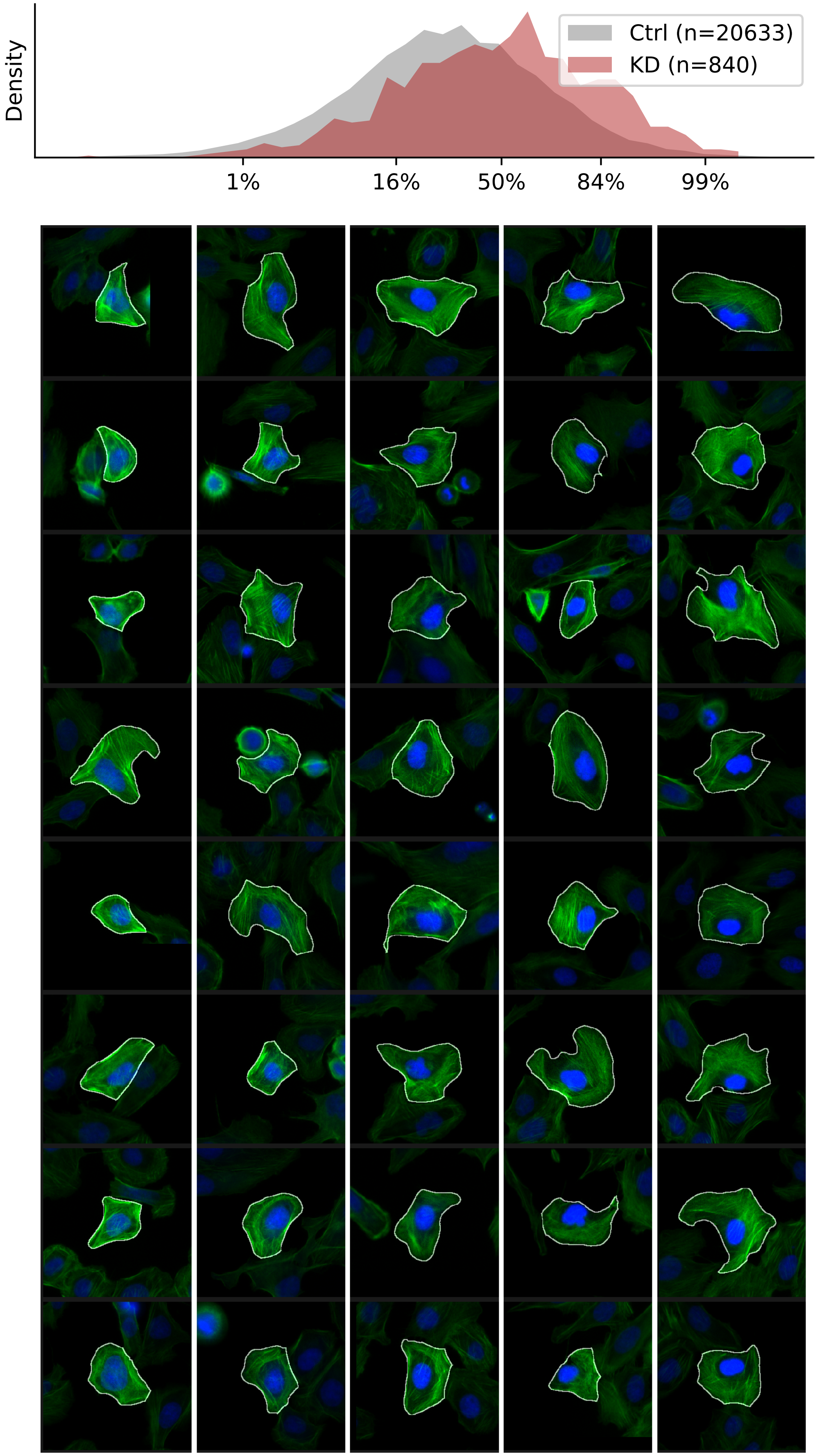

### VIEWS_fPC_TLN1_rep1.pdf

#### TLN1, guide 1

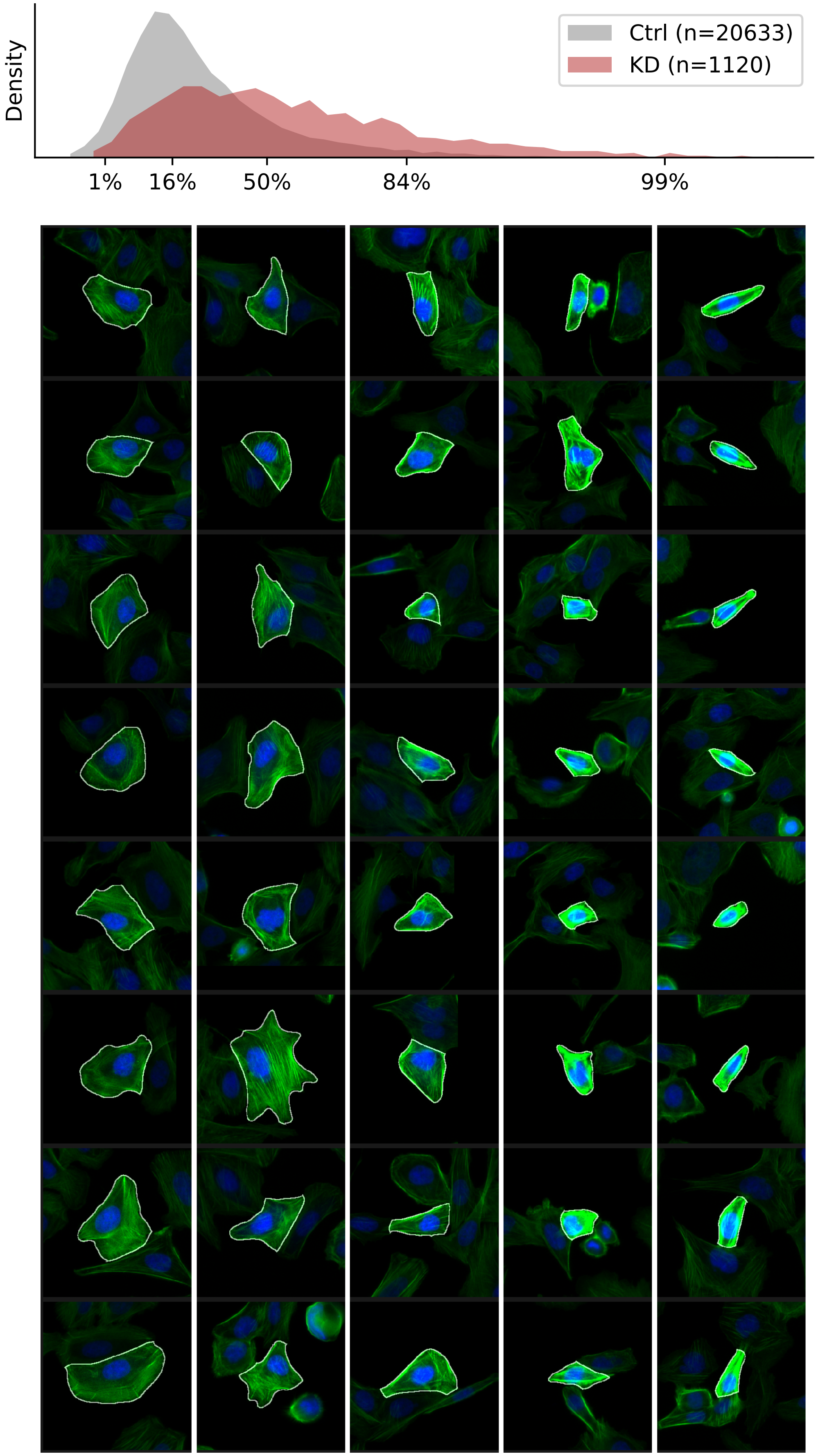

### VIEWS_fPC_TLN1_rep2.pdf

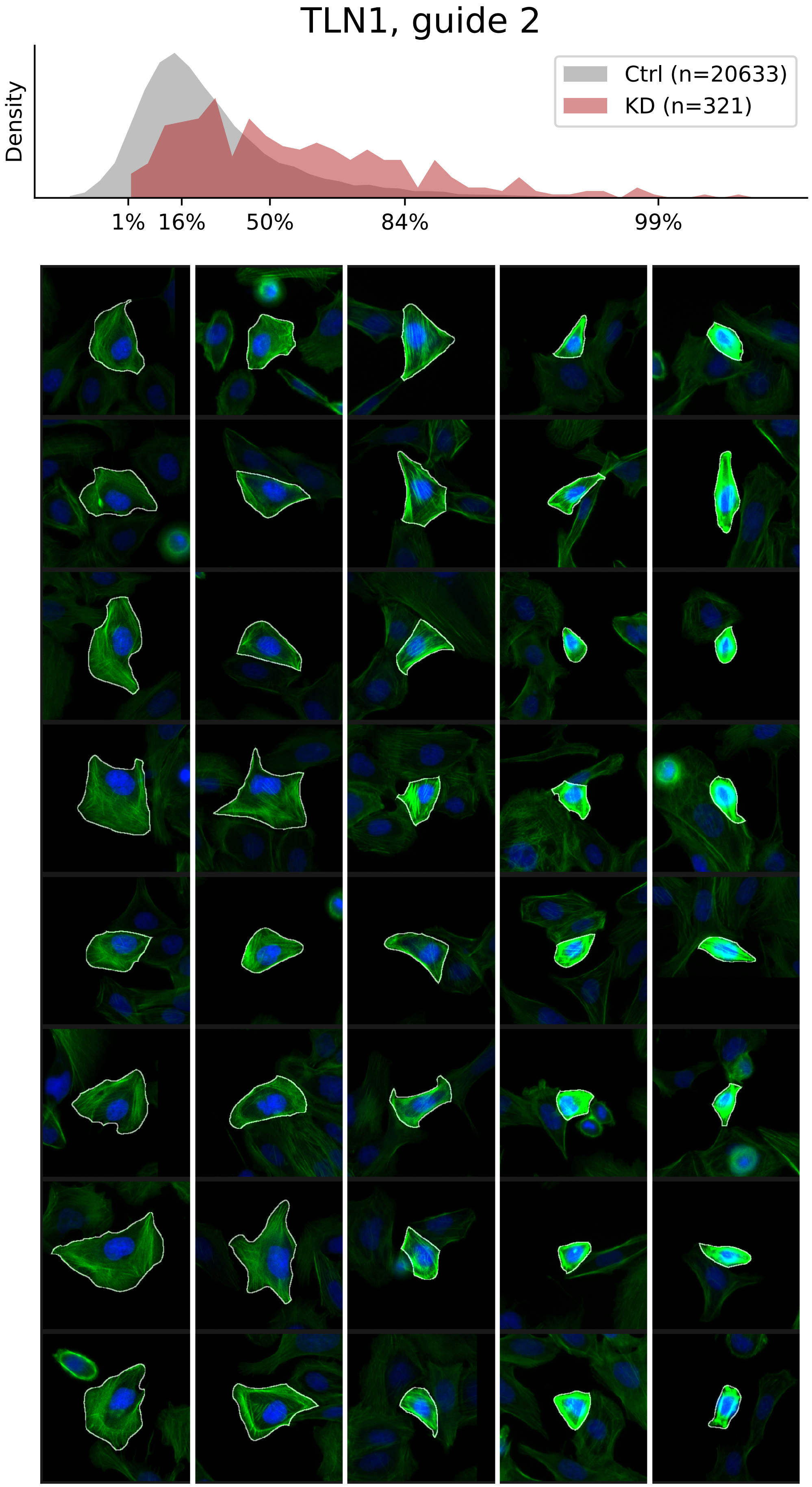

### VIEWS_fPC_TLN1_rep3.pdf

#### TLN1, guide 3

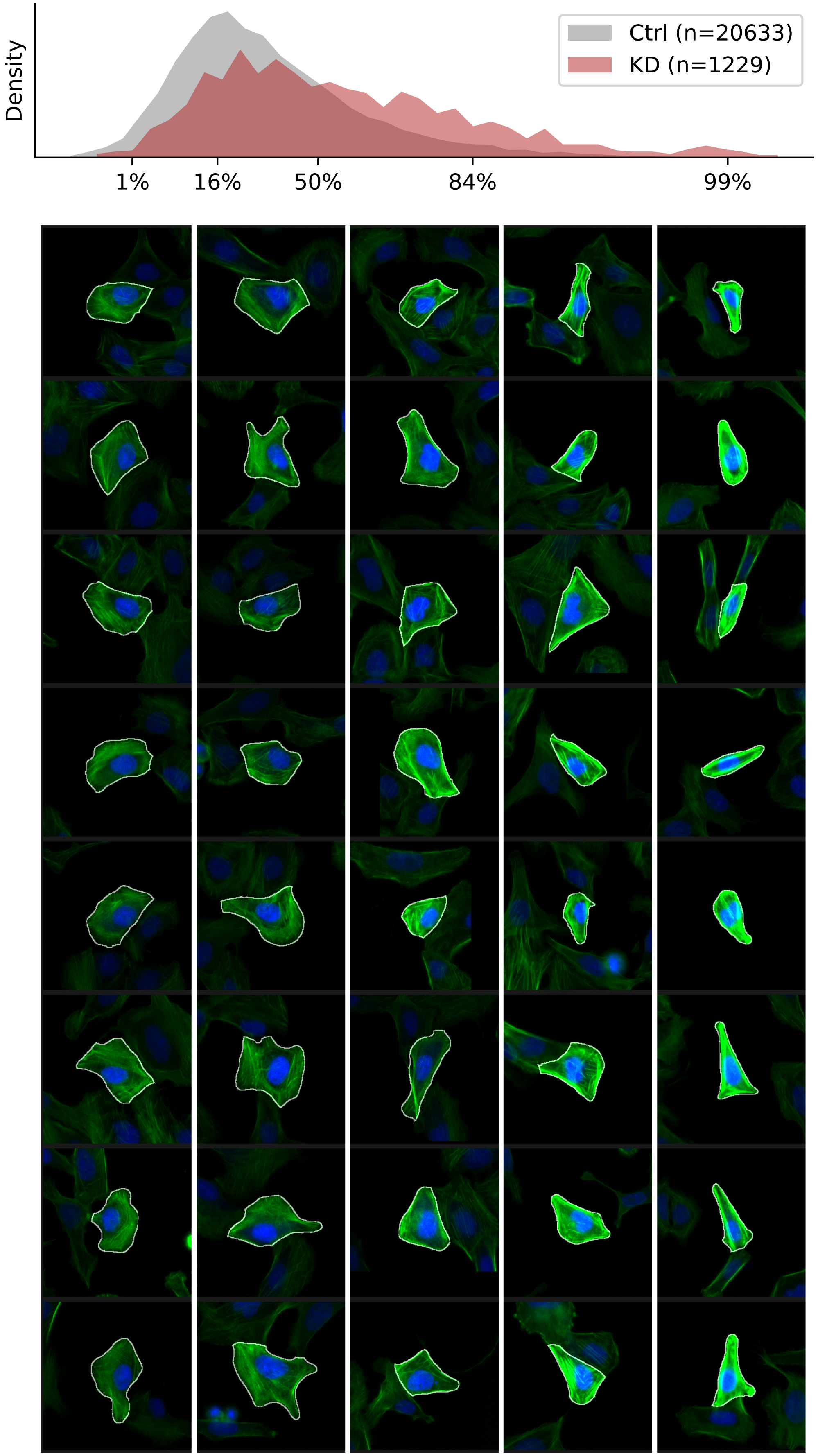

### VIEWS_fPC_TPM3_rep2.pdf

#### TPM3, guide 2

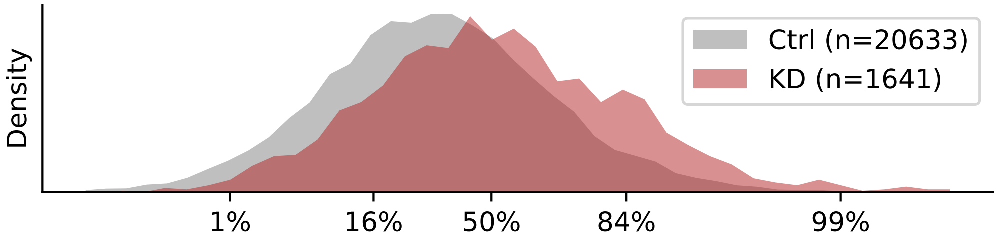


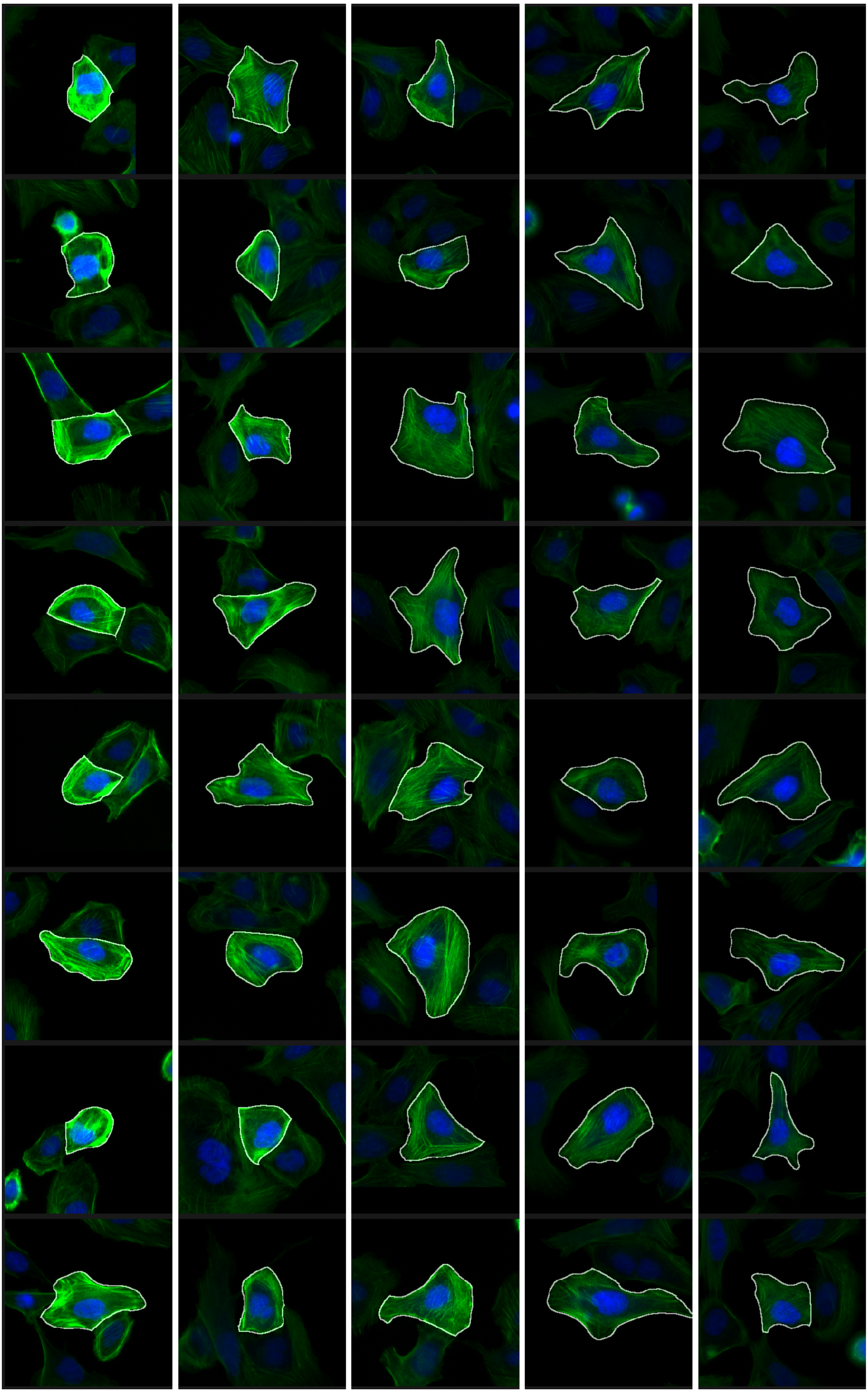

### VIEWS_fPC_VCL_rep1.pdf

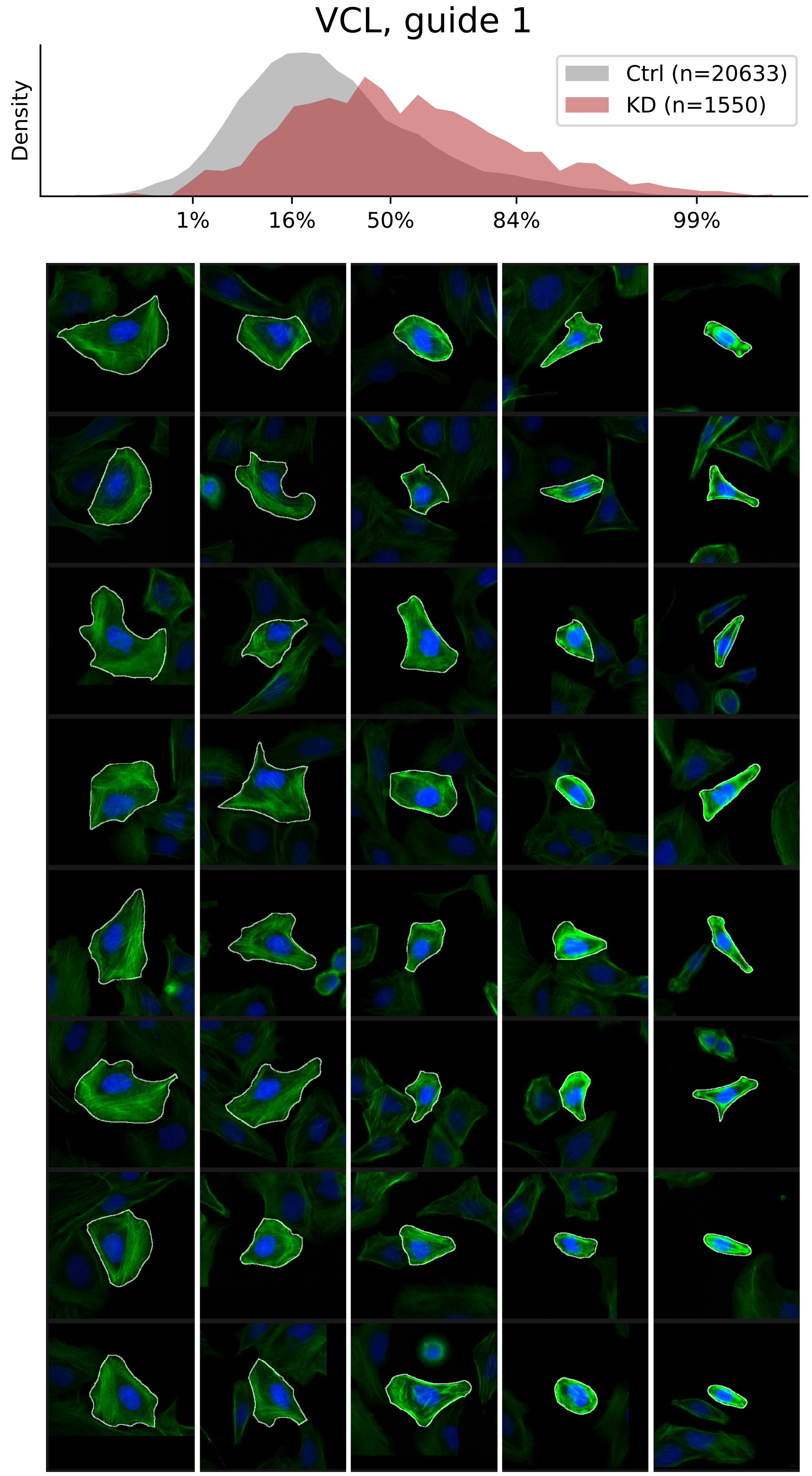

### VIEWS_fPC_VCL_rep2.pdf

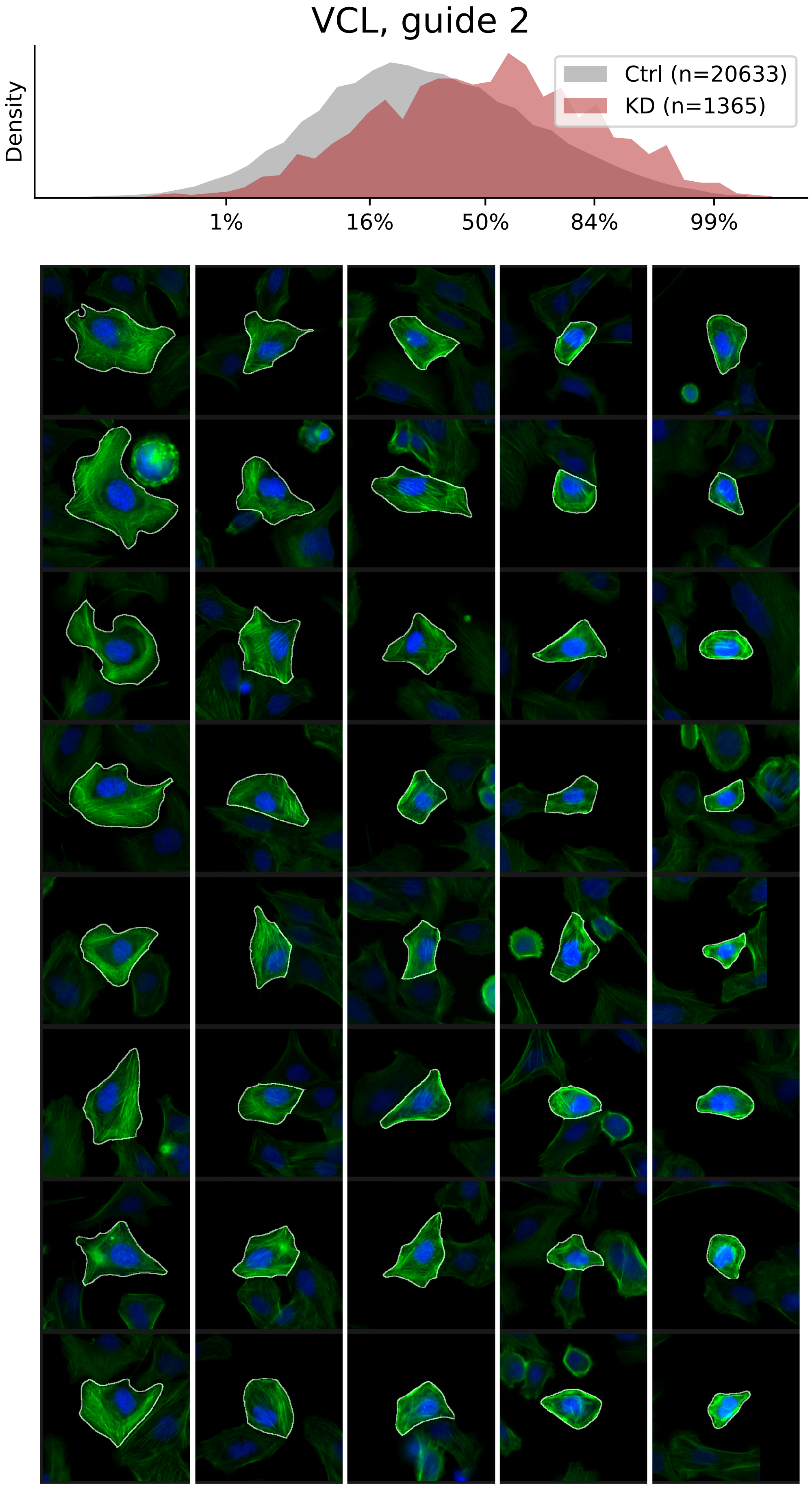

### VIEWS_fPC_VCL_rep3.pdf

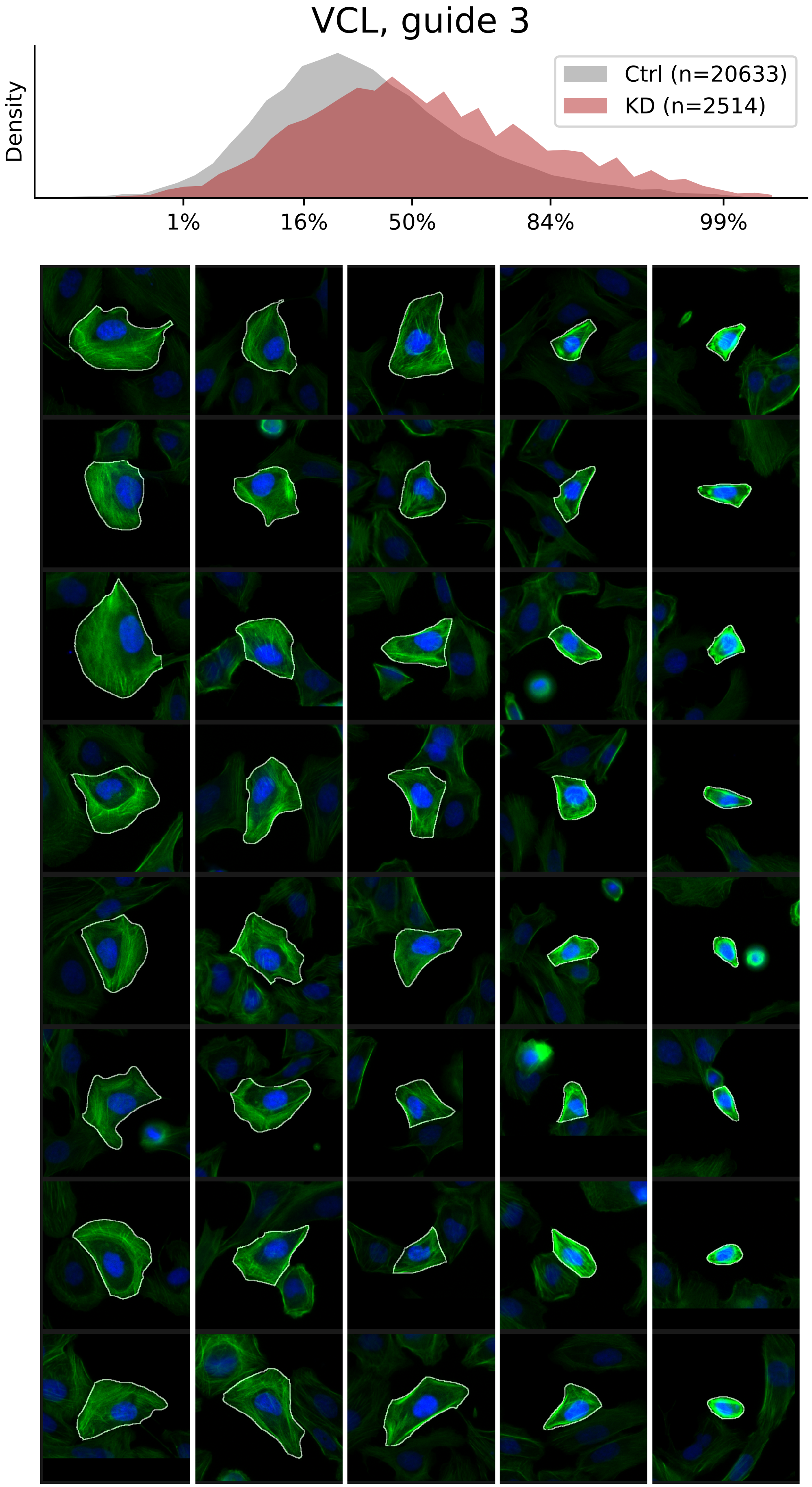

### VIEWS_iAD1.pdf

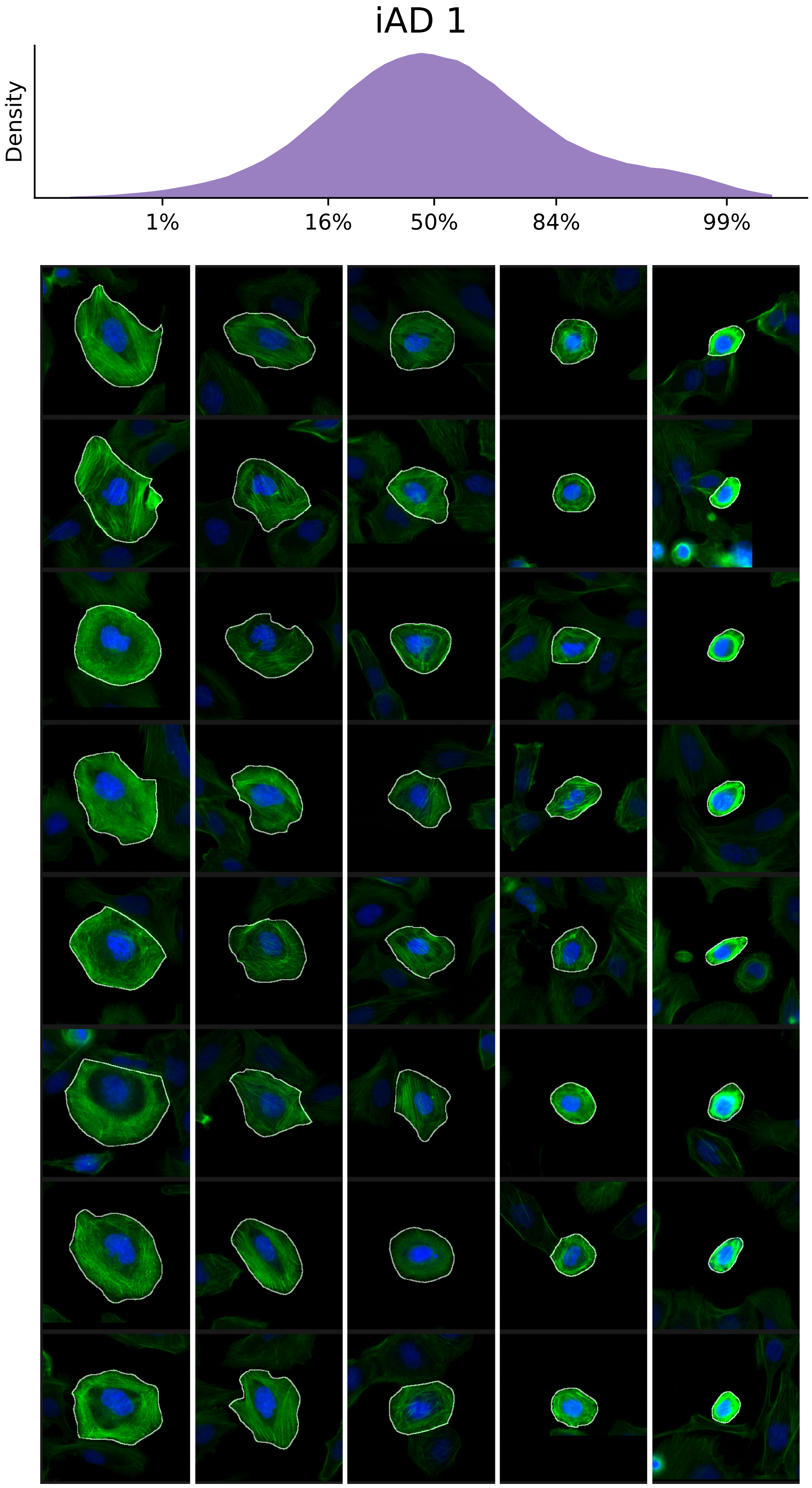

### VIEWS_iAD2.pdf

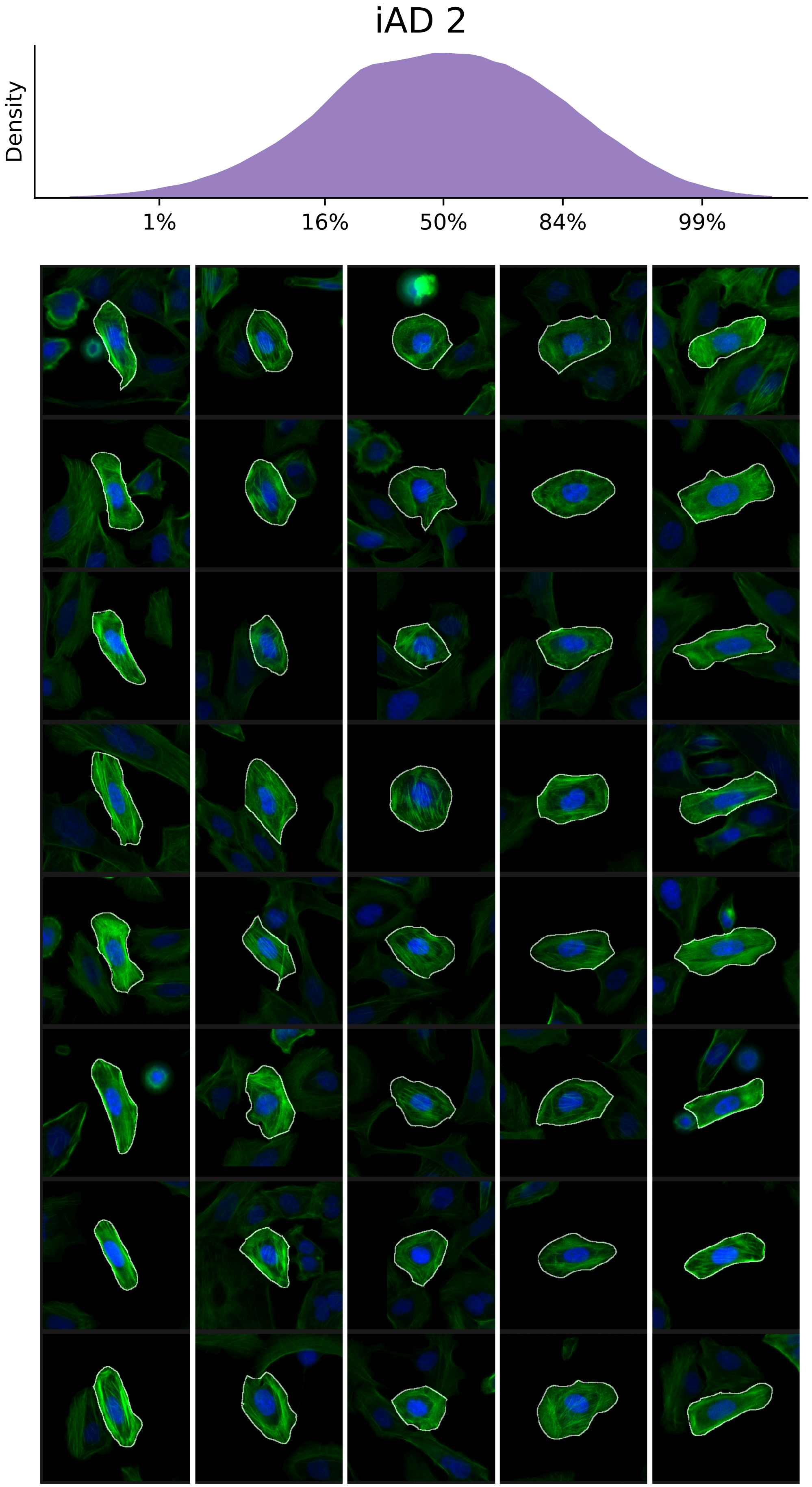

### VIEWS_iAD3.pdf

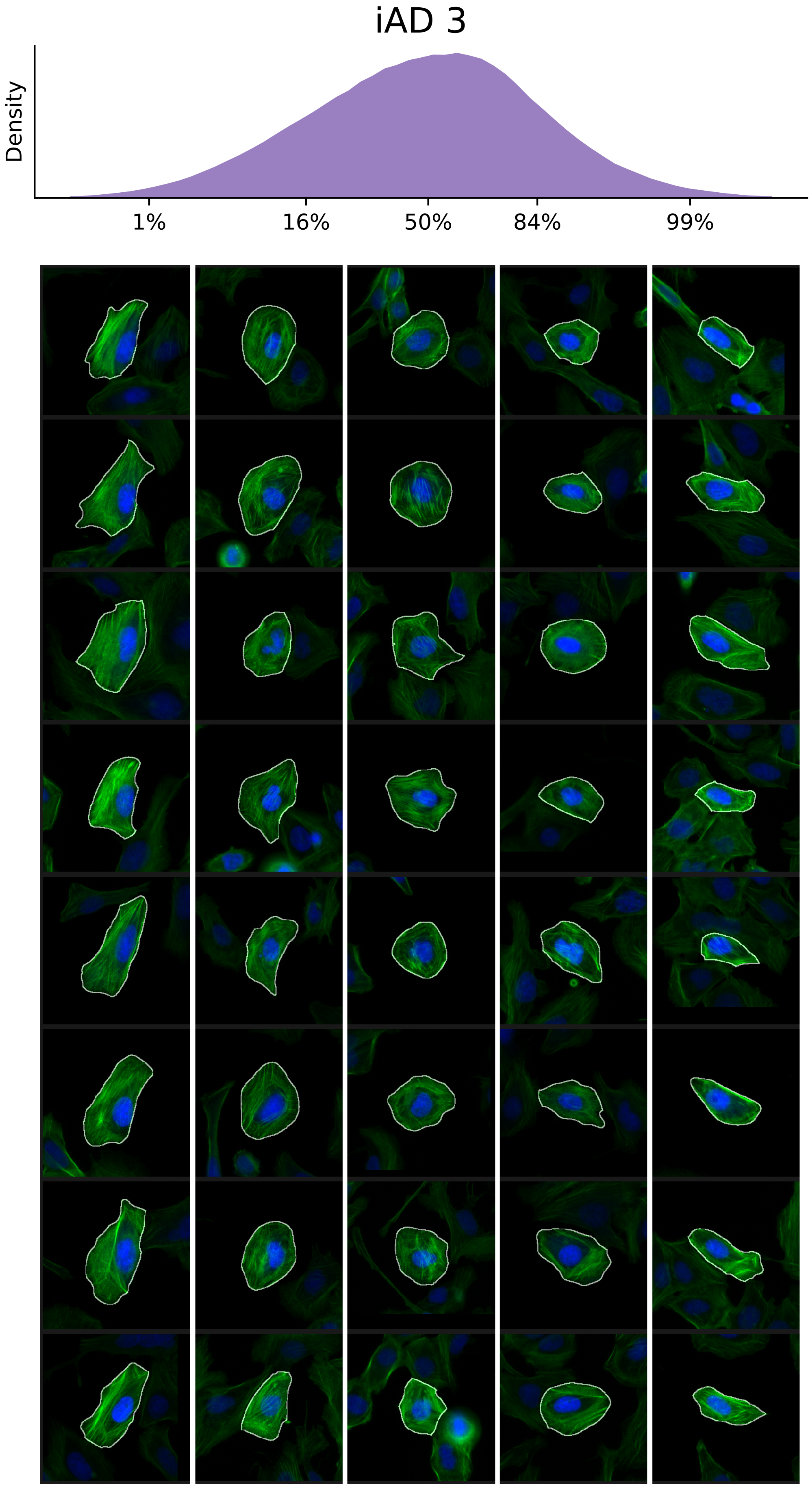

### VIEWS_iAD4.pdf

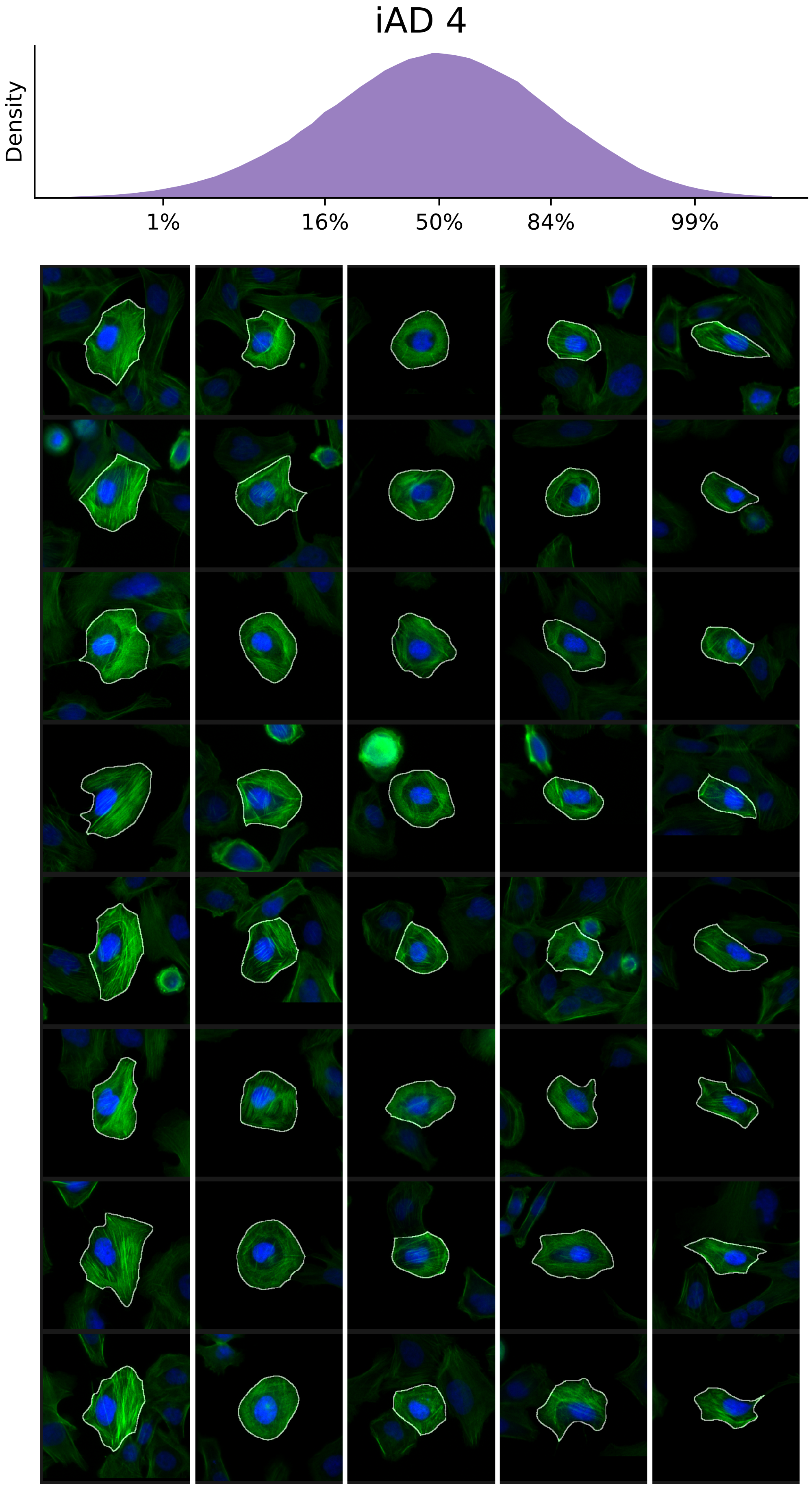

### VIEWS_iAD5.pdf

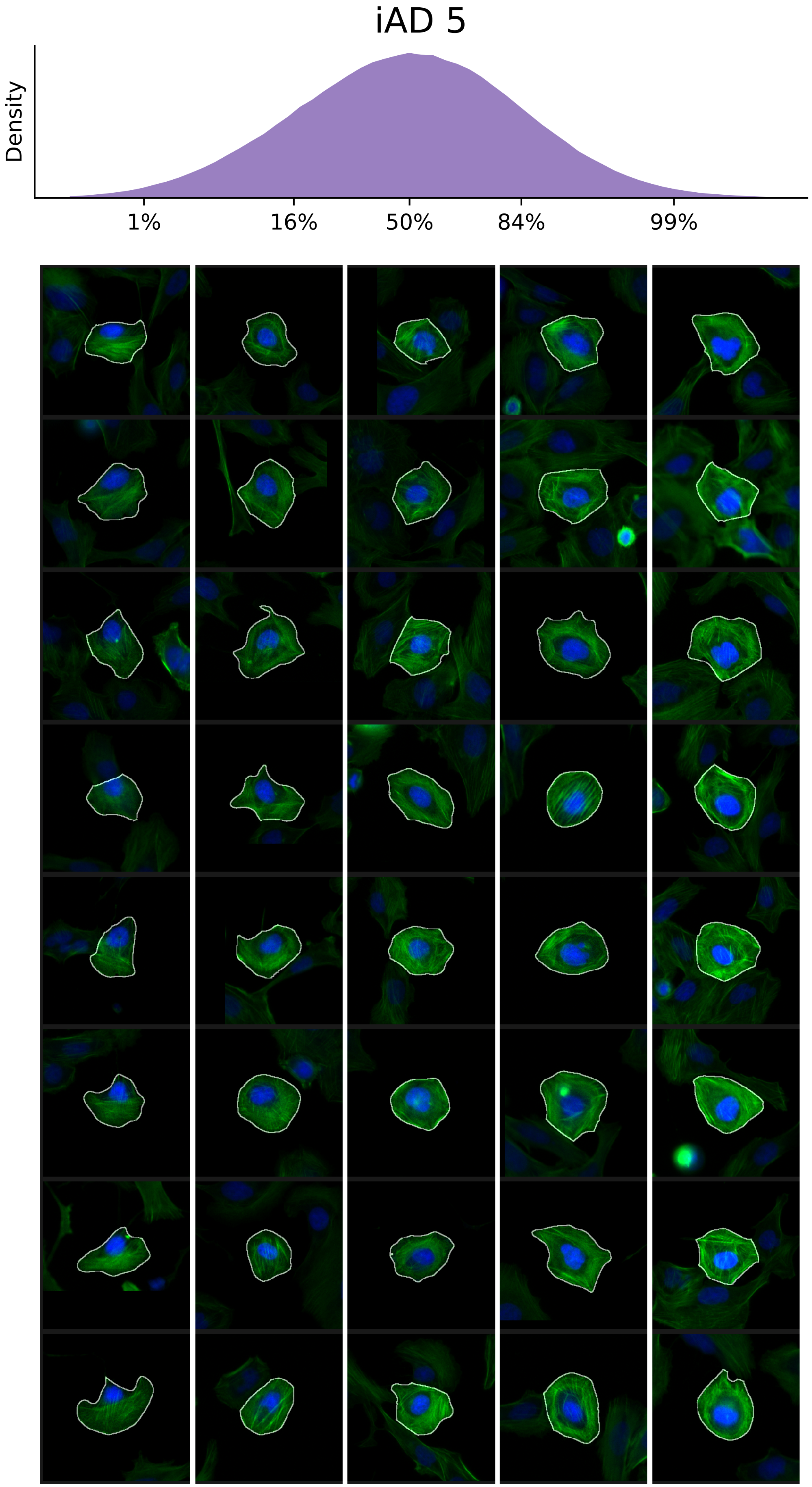

### VIEWS_iAD6.pdf

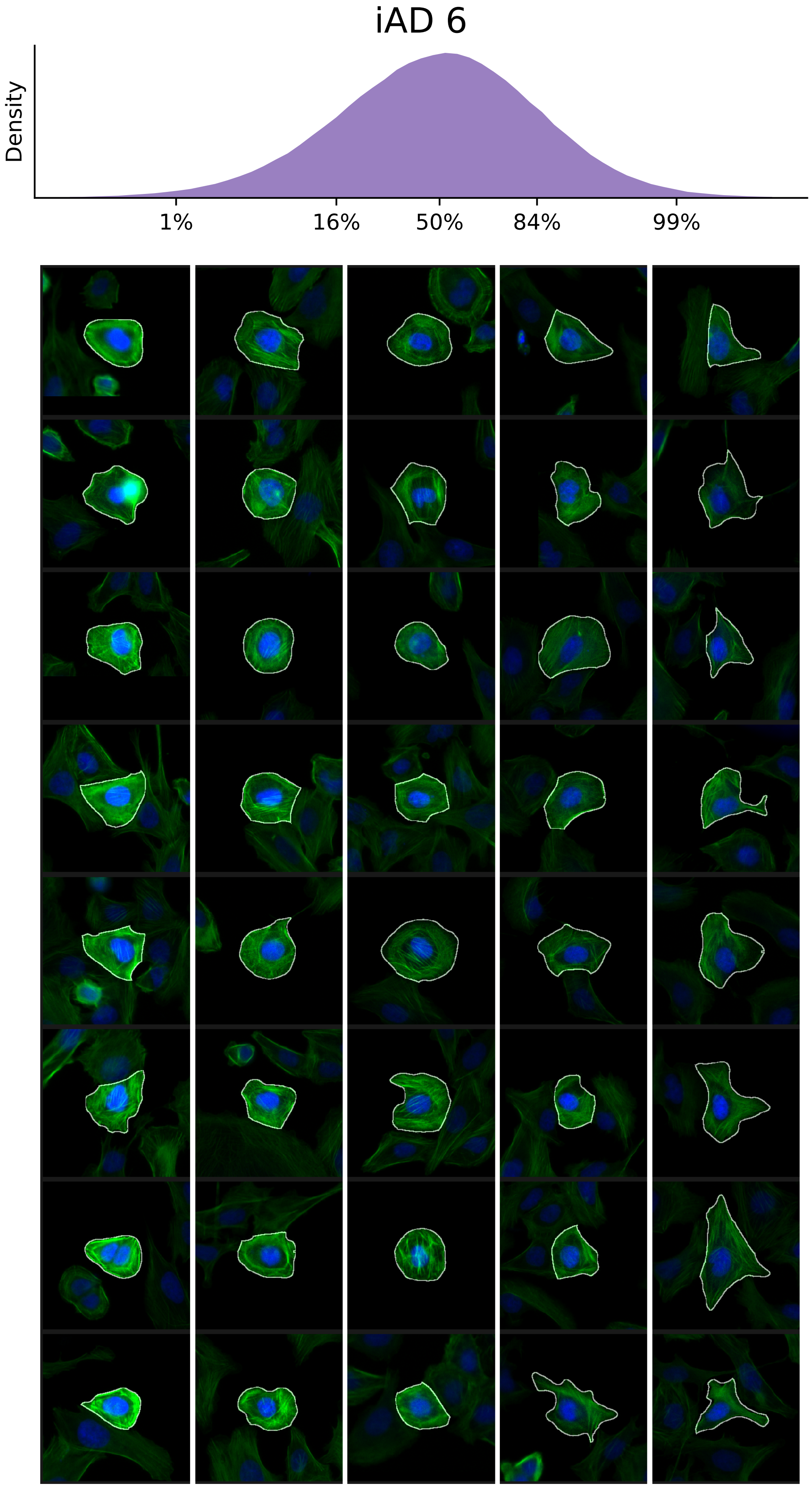

### VIEWS_iAD7.pdf

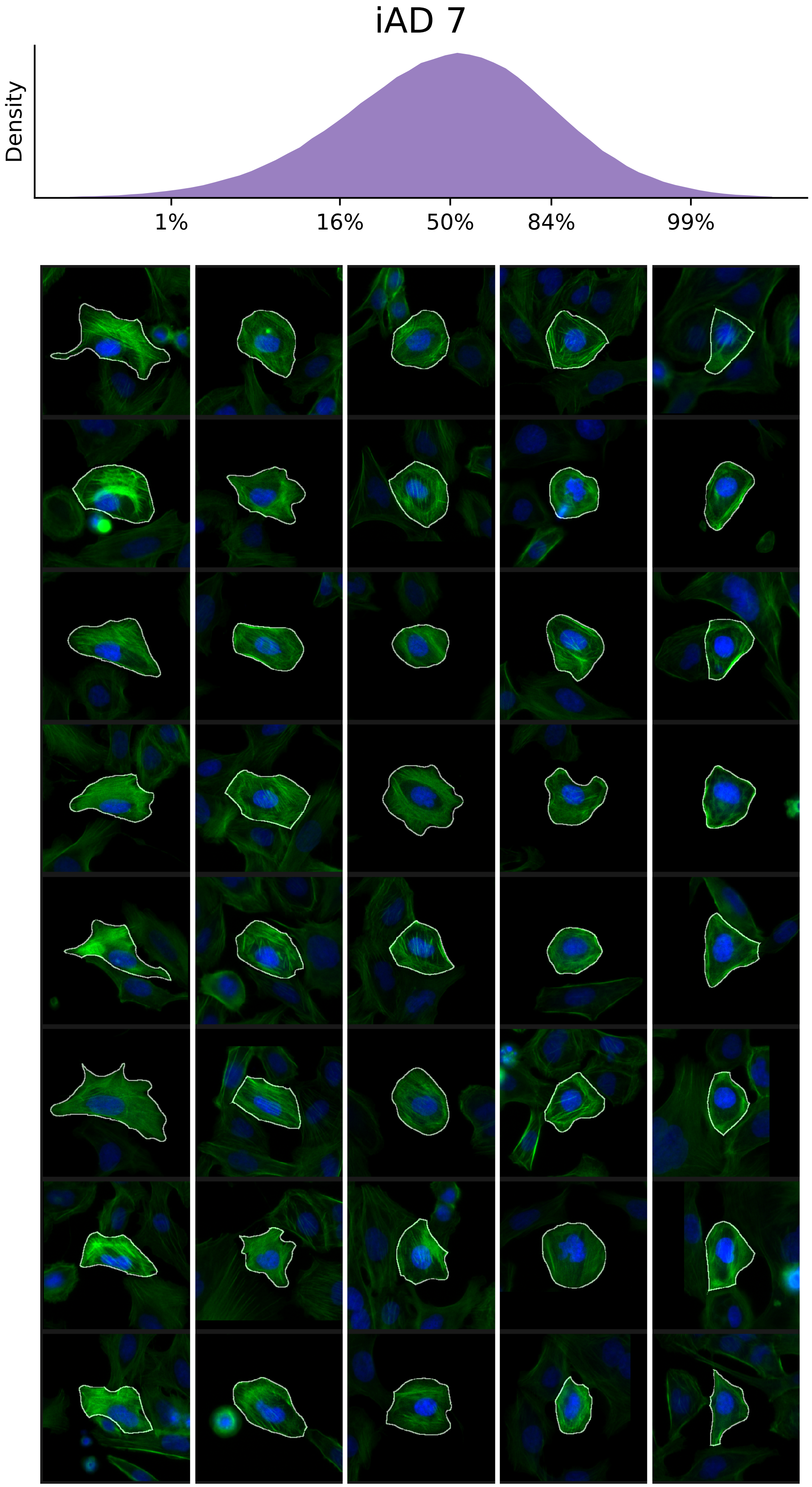

### VIEWS_iAD8.pdf

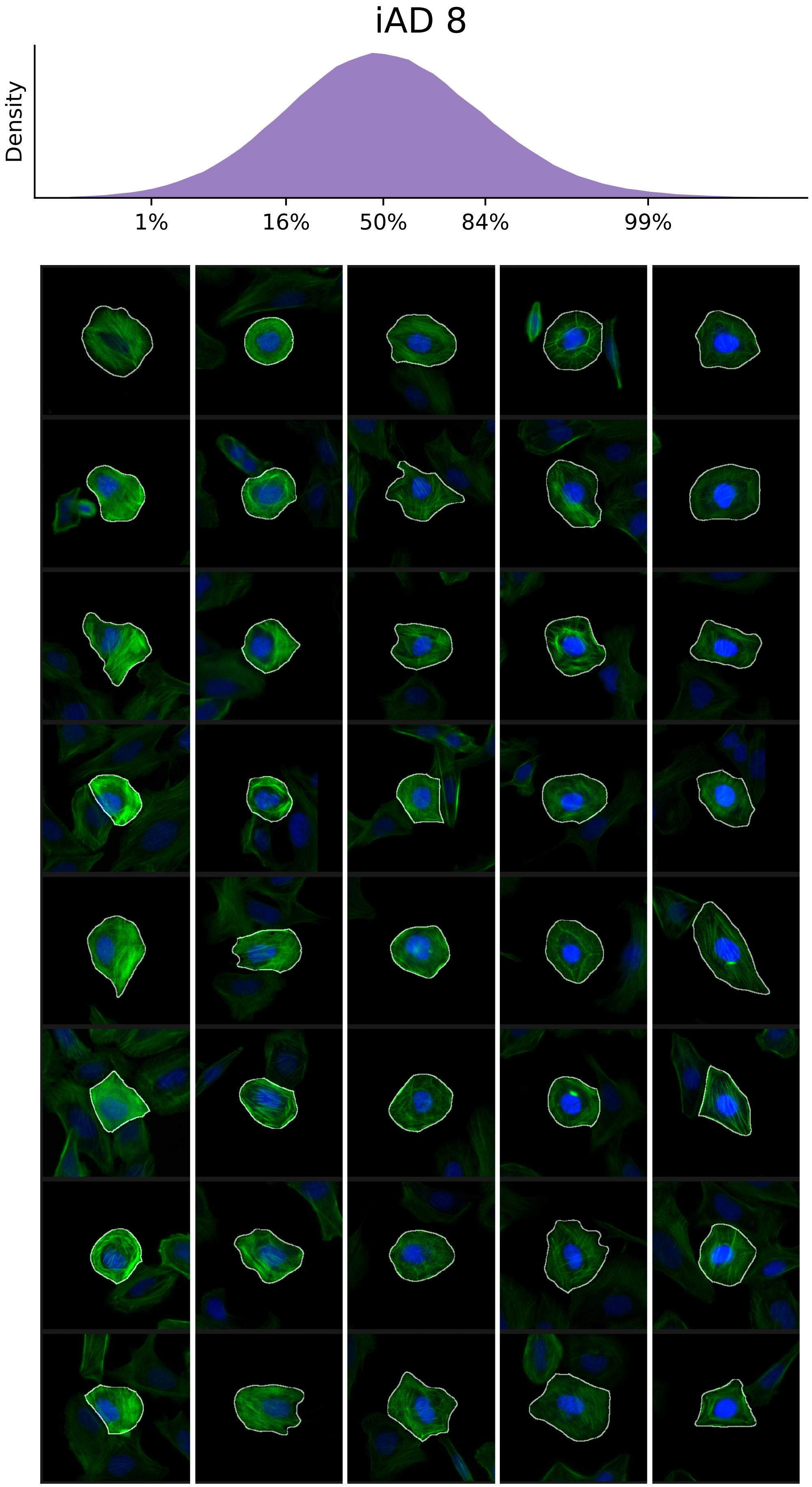

### VIEWS_iAD9.pdf

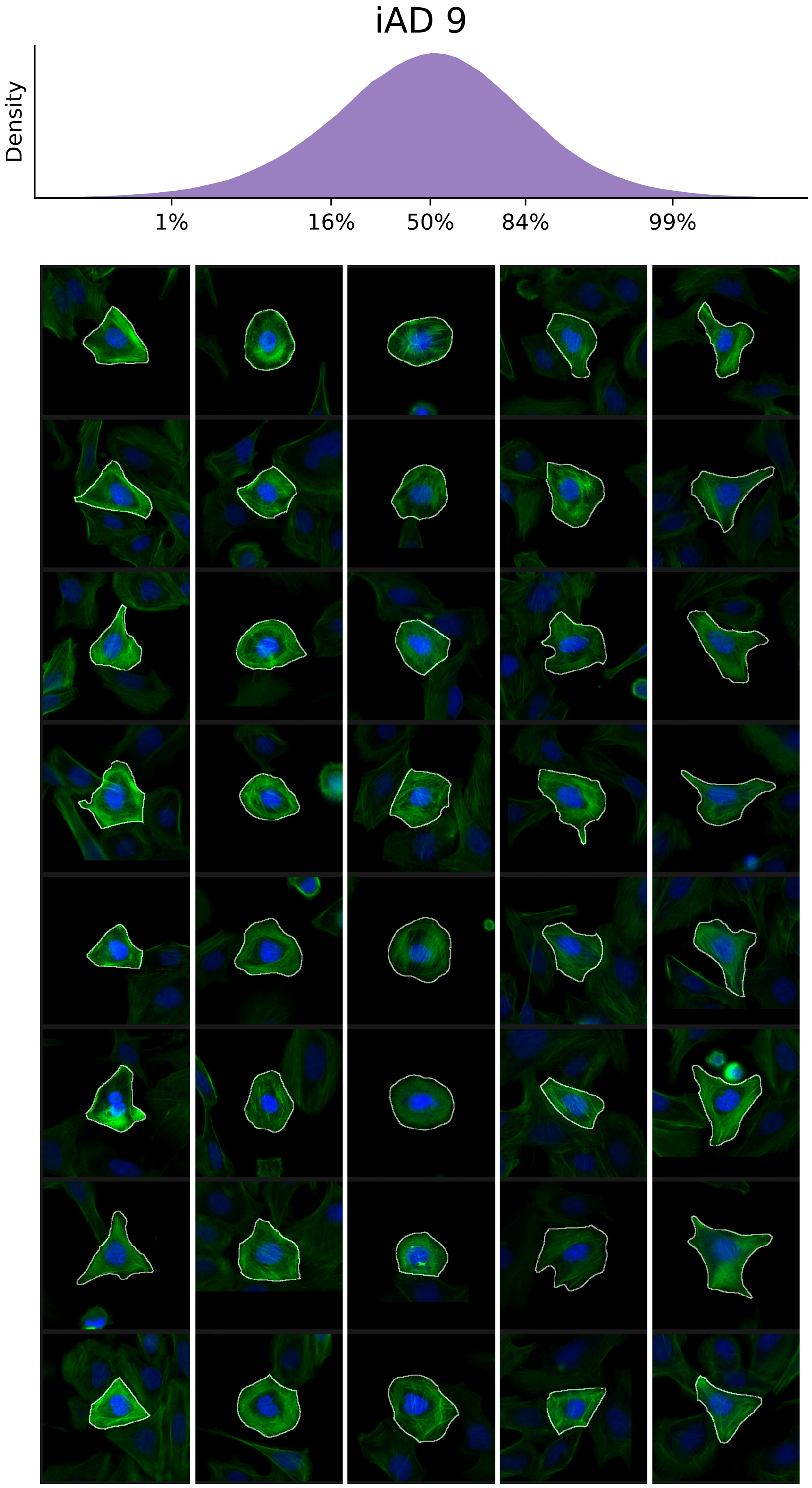

### VIEWS_iAD10.pdf

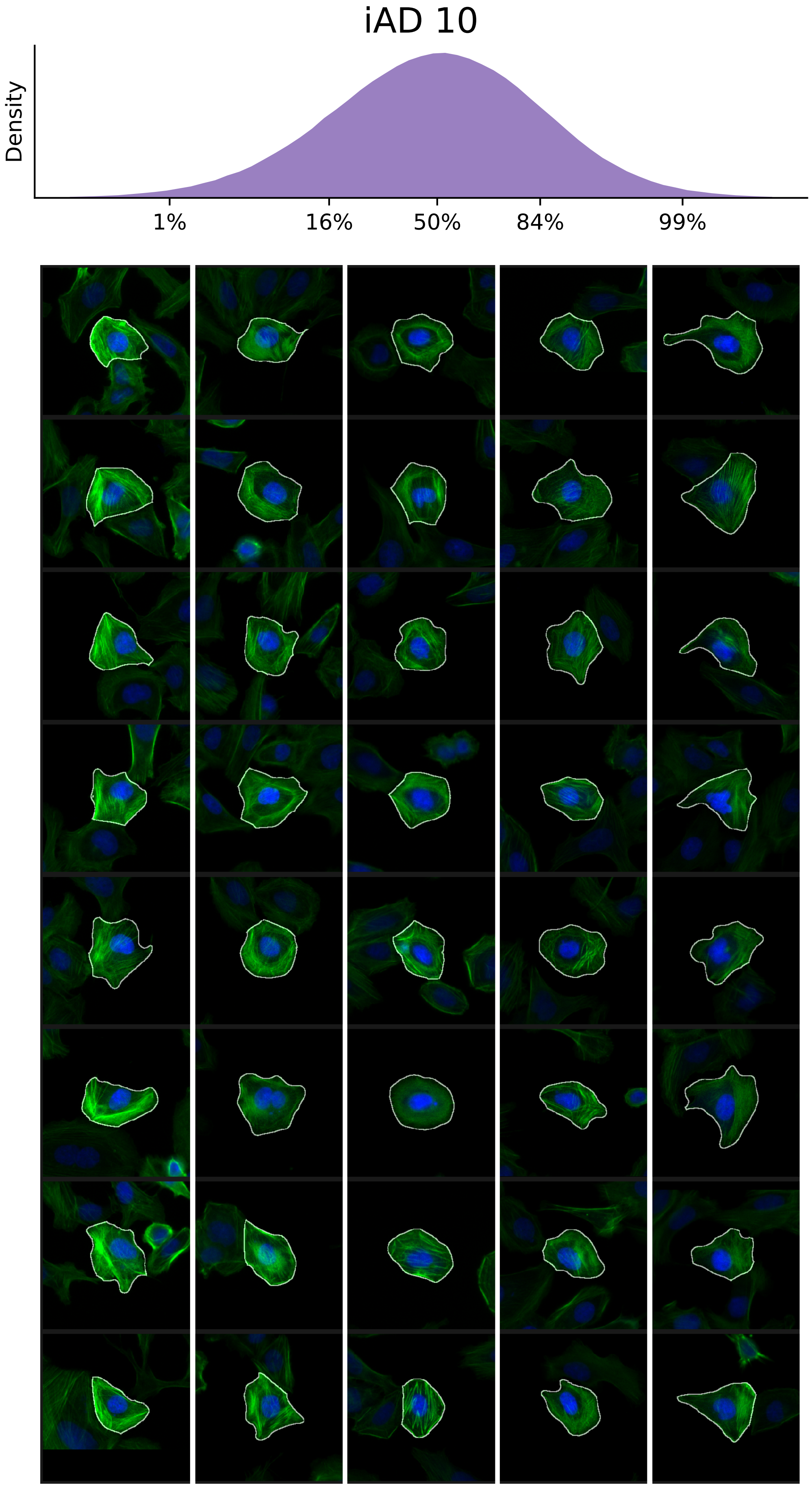

### VIEWS_iAD11.pdf

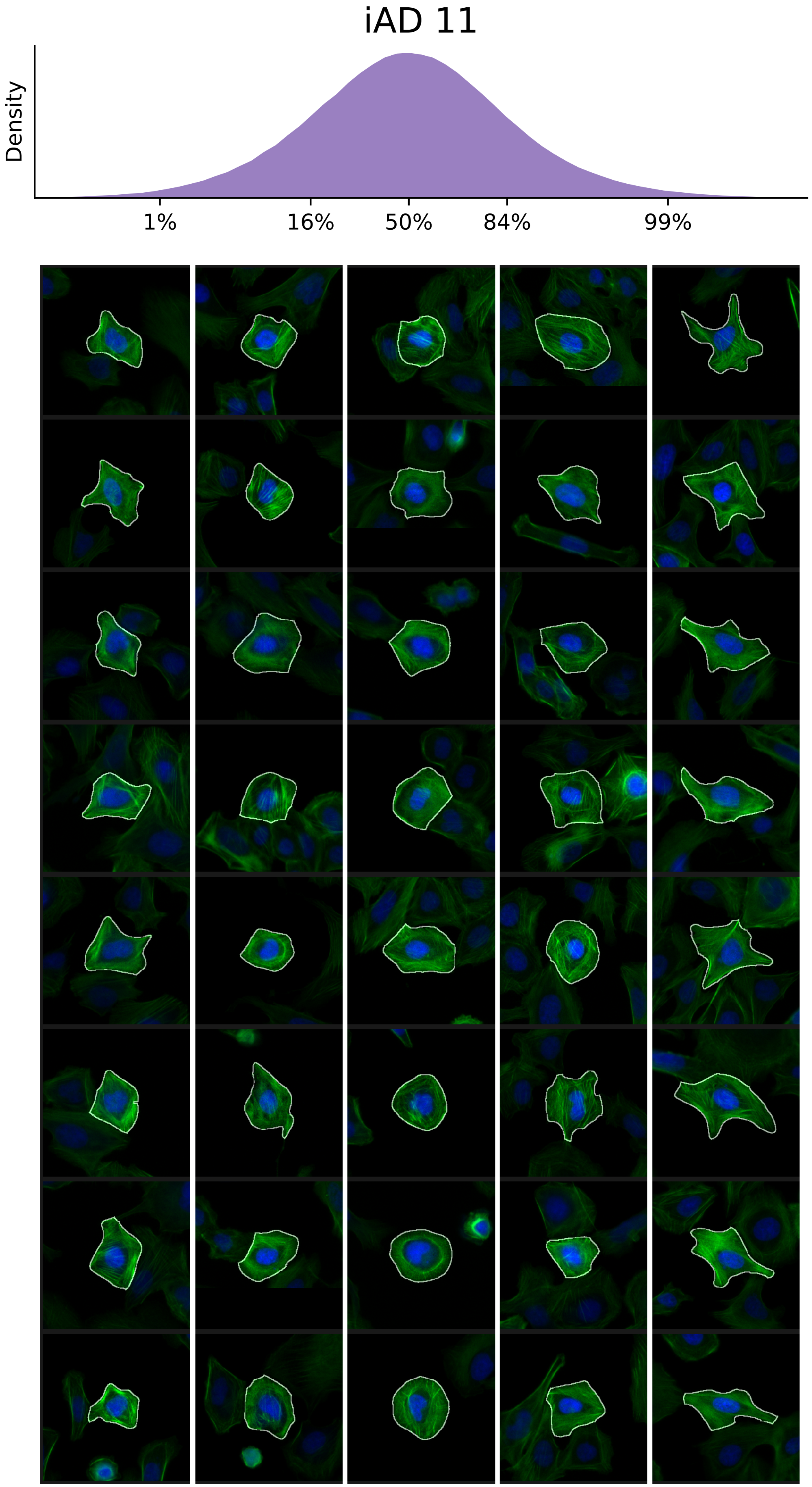

### VIEWS_iAD12.pdf

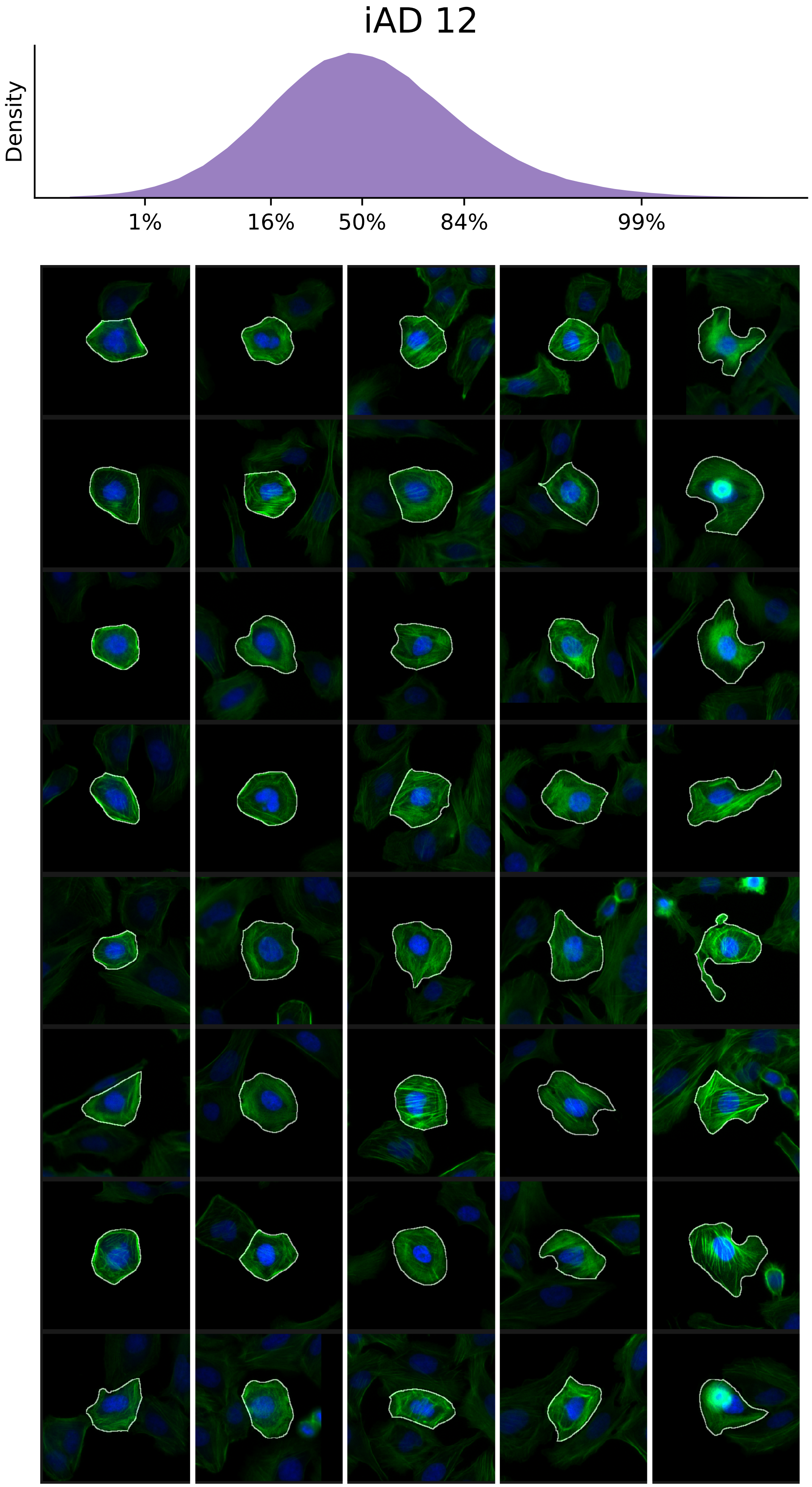

### VIEWS_iAD13.pdf

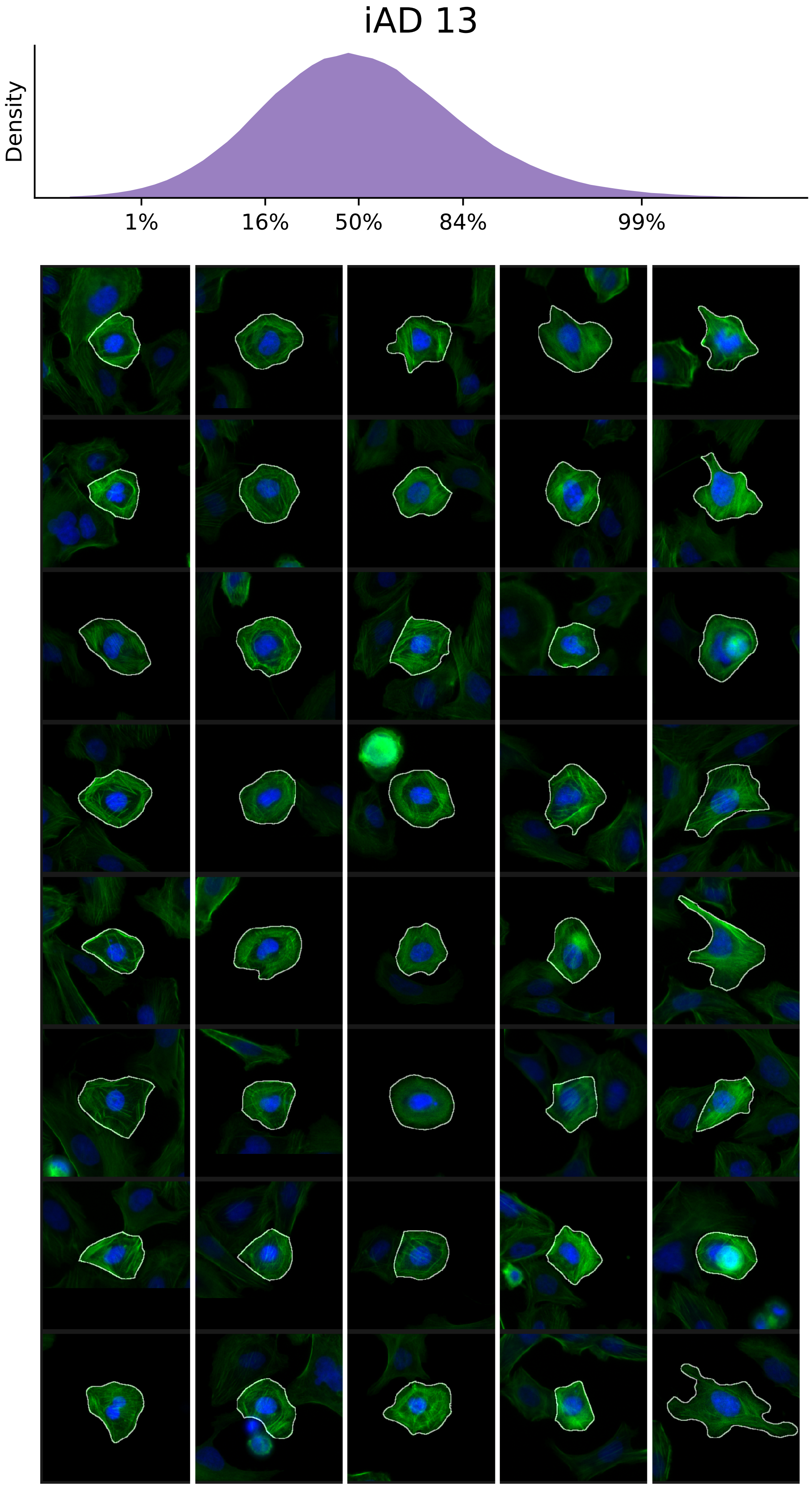

### VIEWS_iAD14.pdf

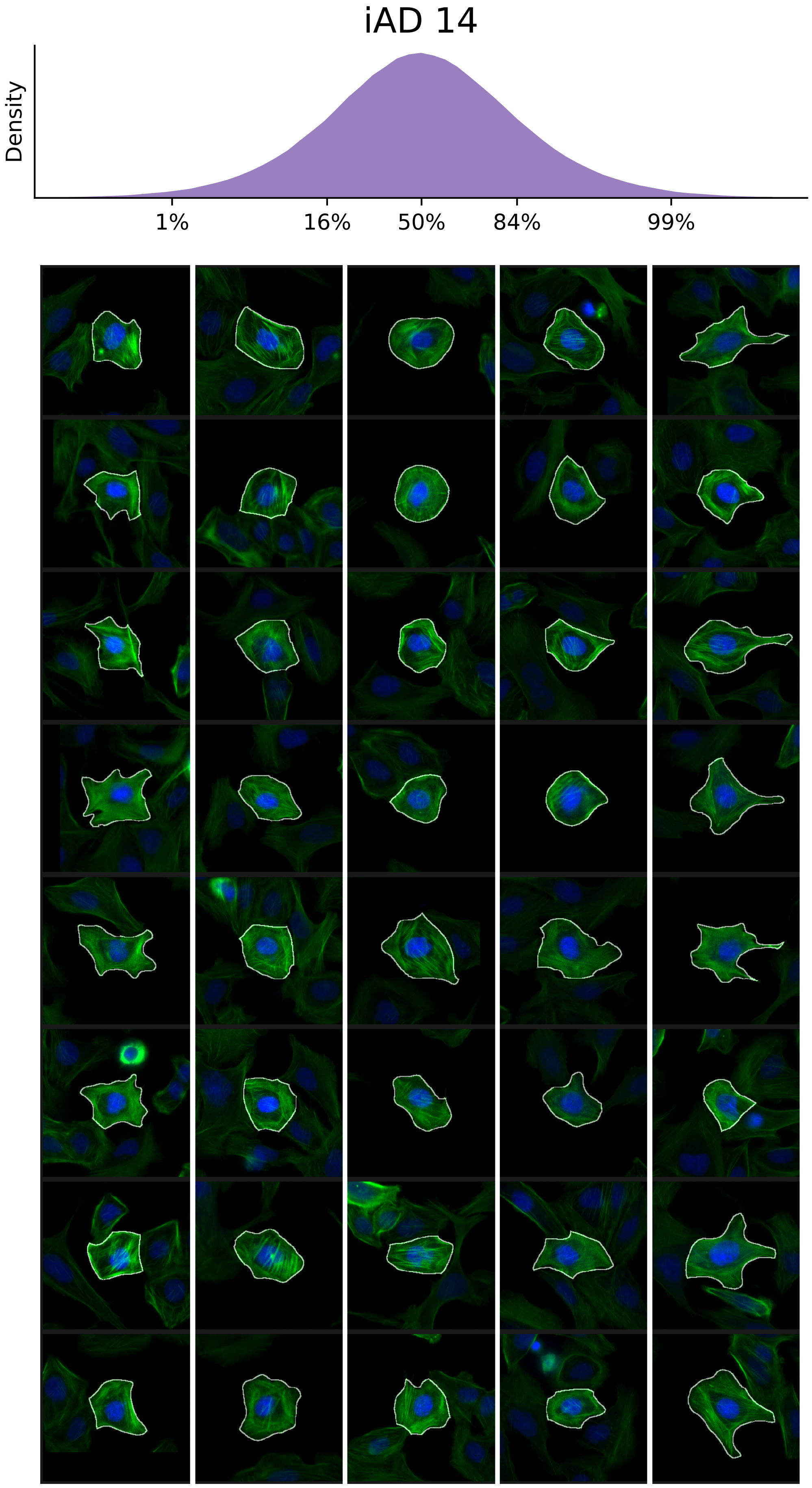

### VIEWS_iAD15.pdf

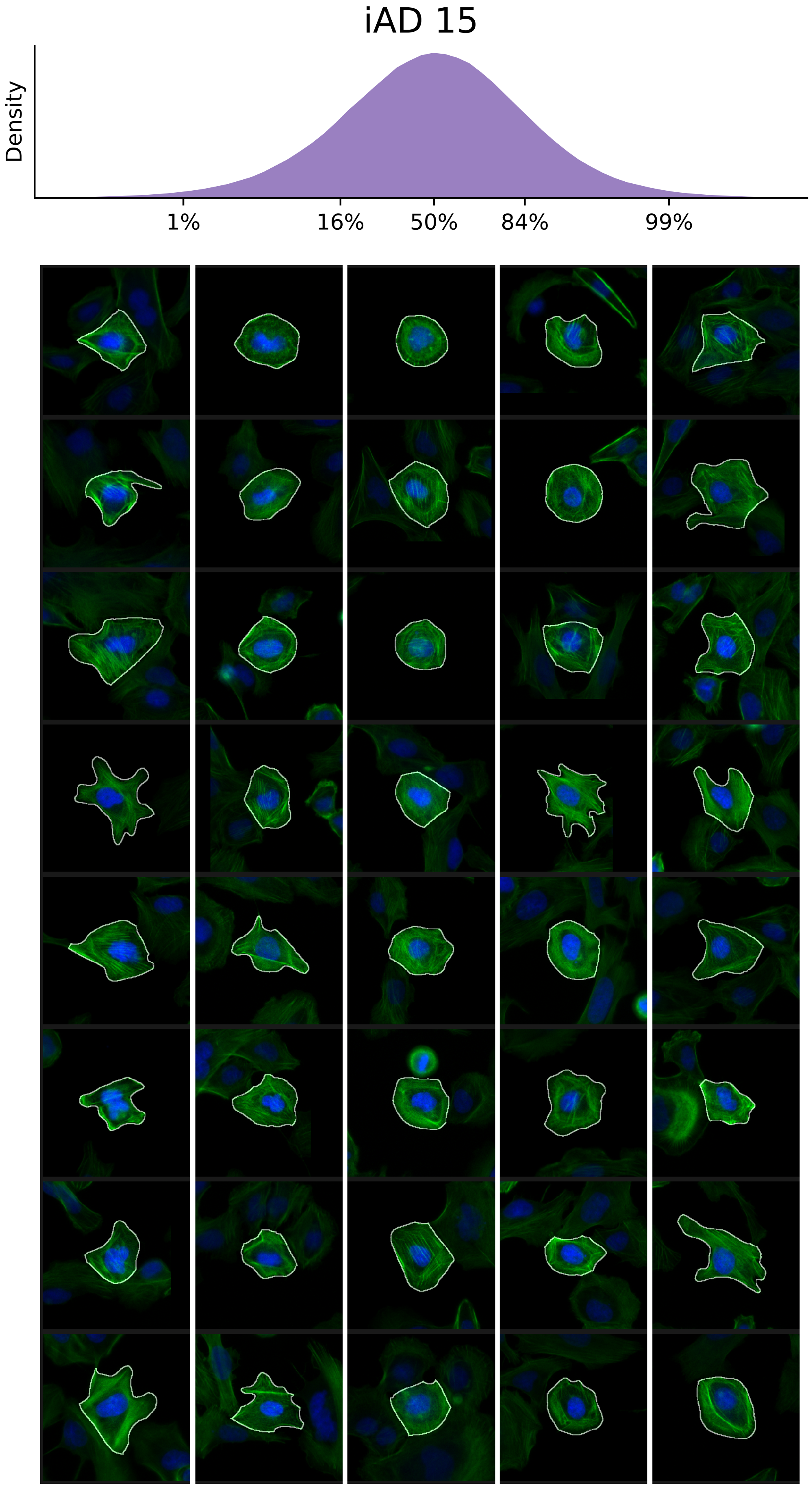

### VIEWS_iAD-LD-verified_ARHGEF7_rep2.pdf

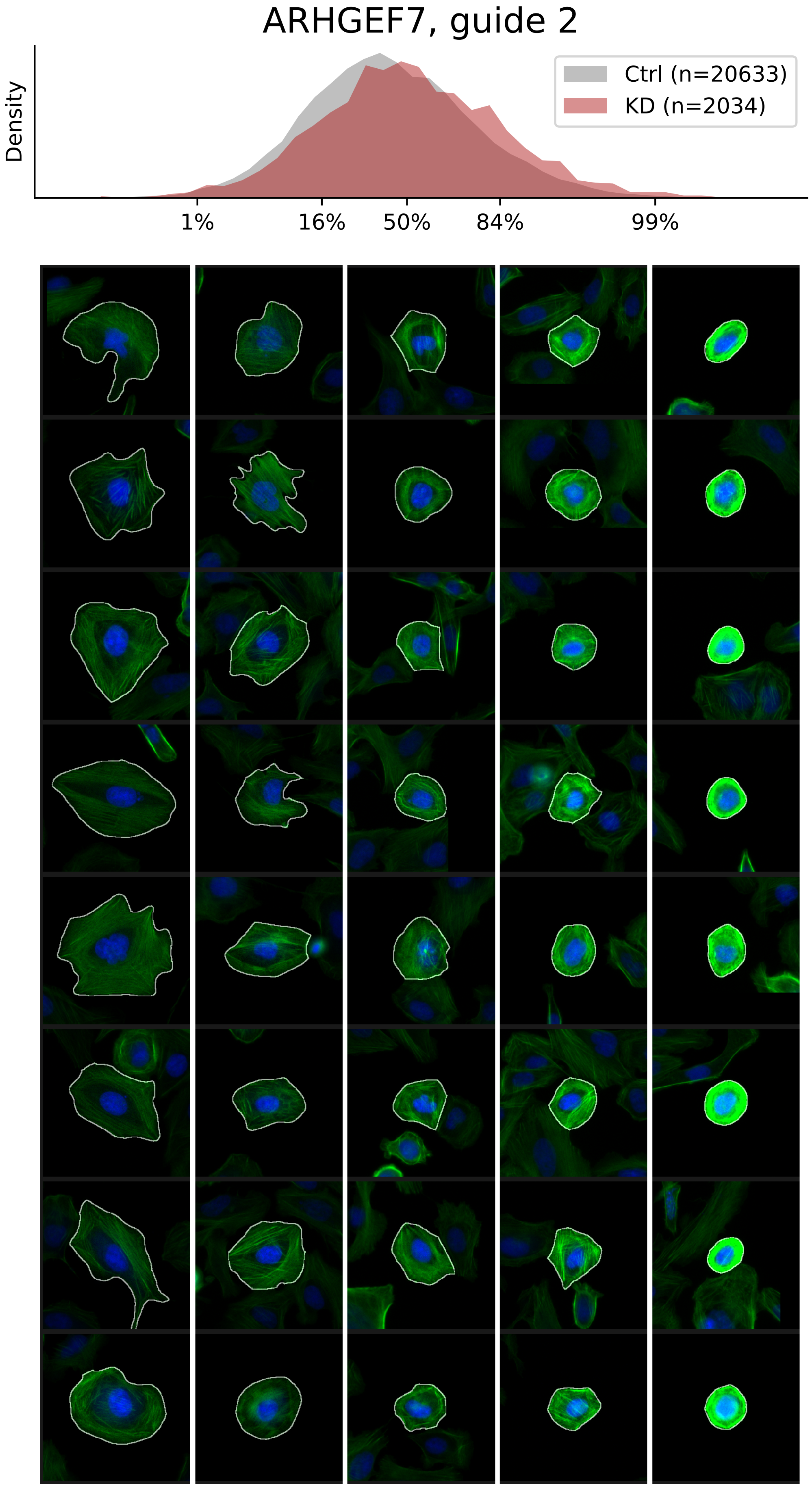

### VIEWS_iAD-LD-verified_ARPC3_rep1.pdf

#### ARPC3, guide 1

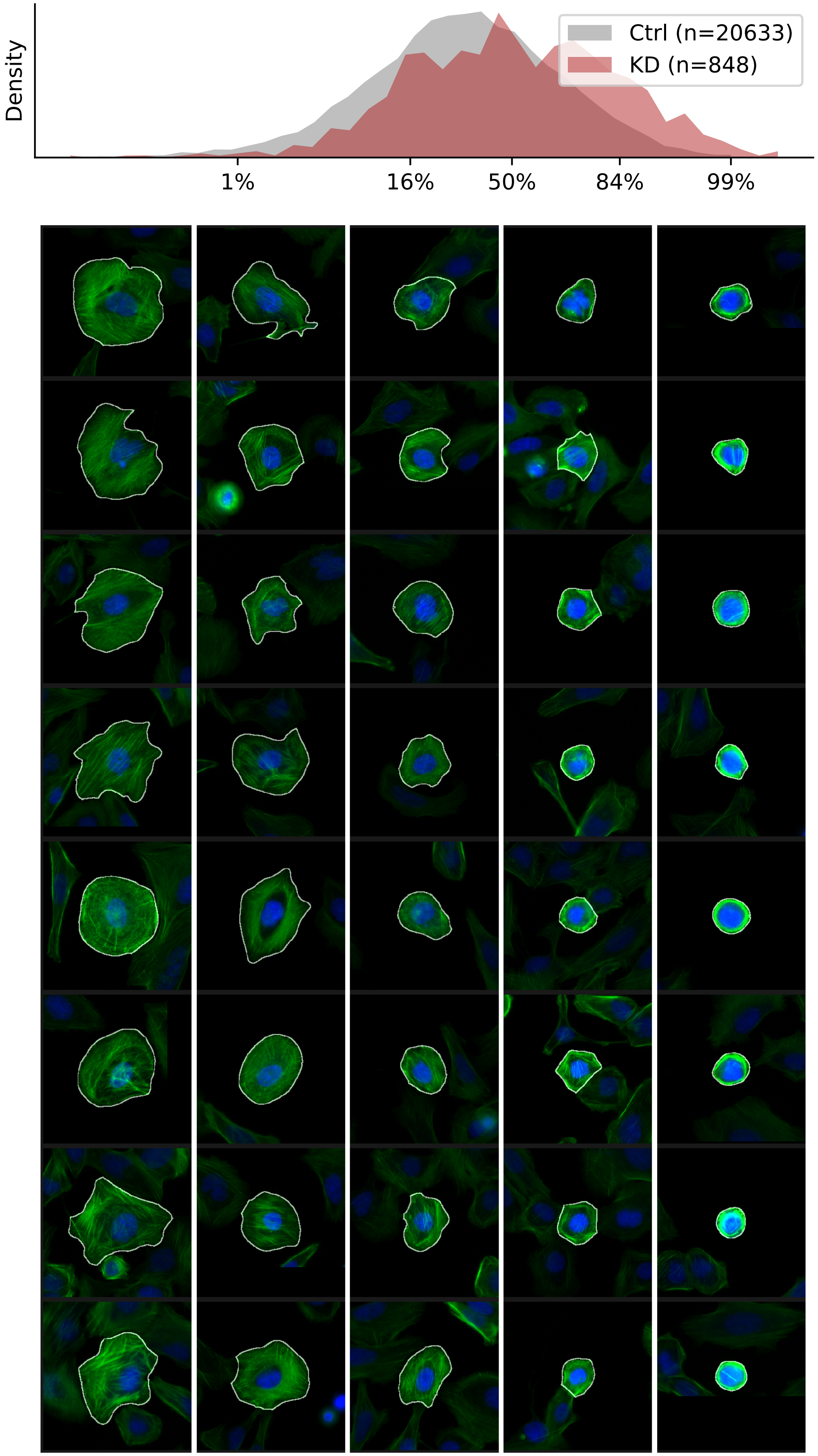

### VIEWS_iAD-LD-verified_CAP1_rep2.pdf

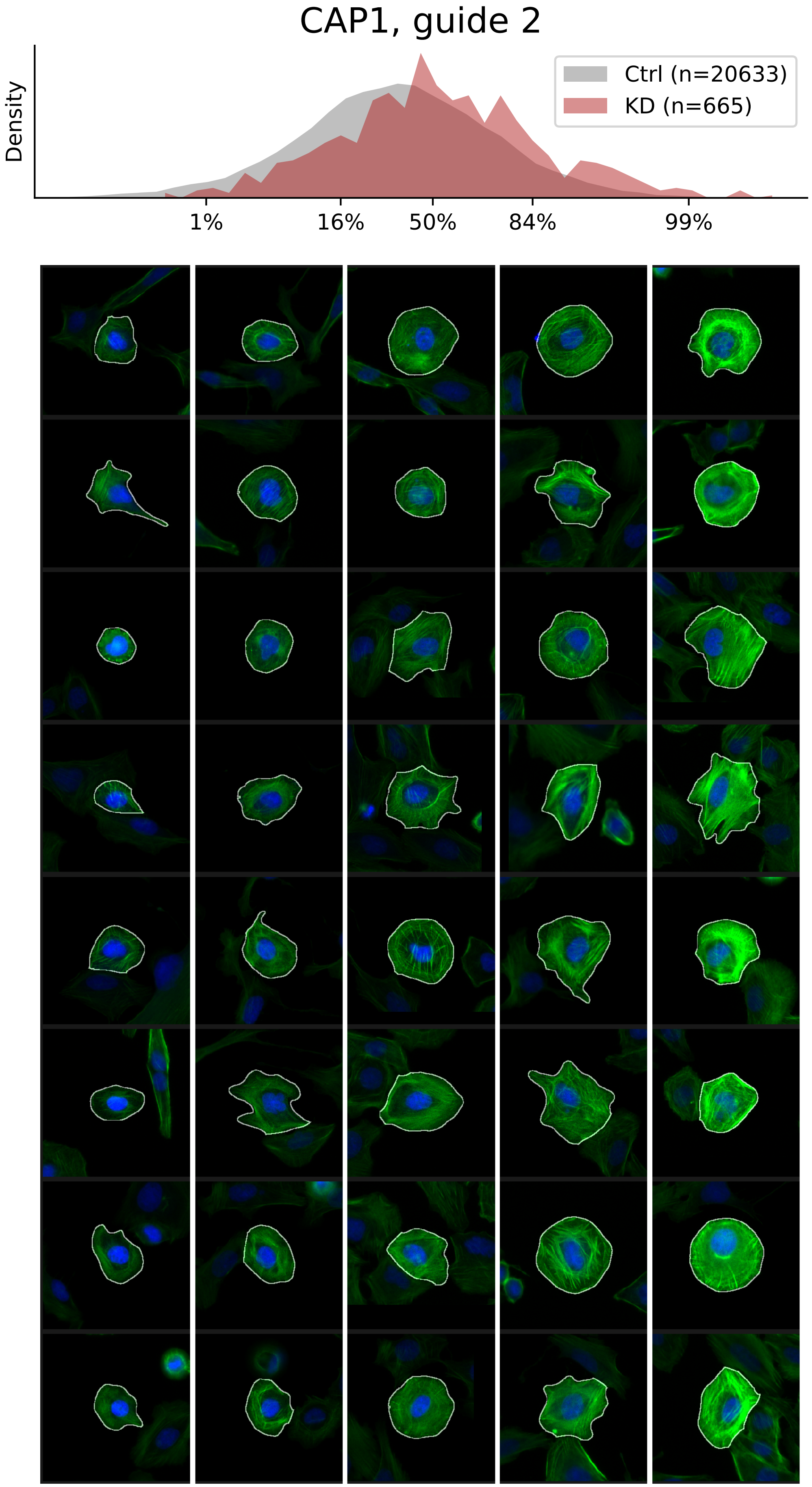

### VIEWS_iAD-LD-verified_MAEA_rep2.pdf

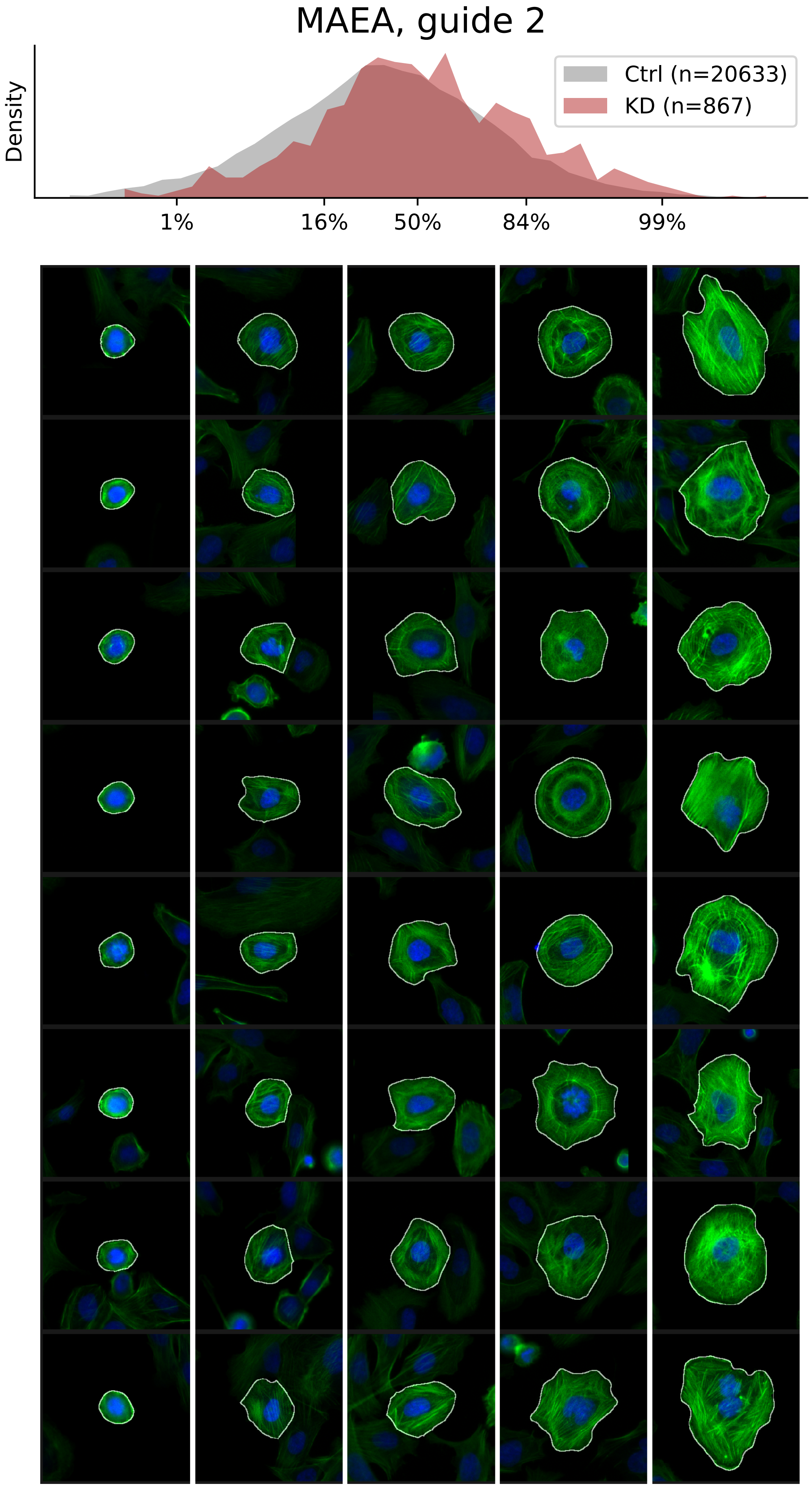

### VIEWS_iAD-LD-verified_ROCK2_rep1.pdf

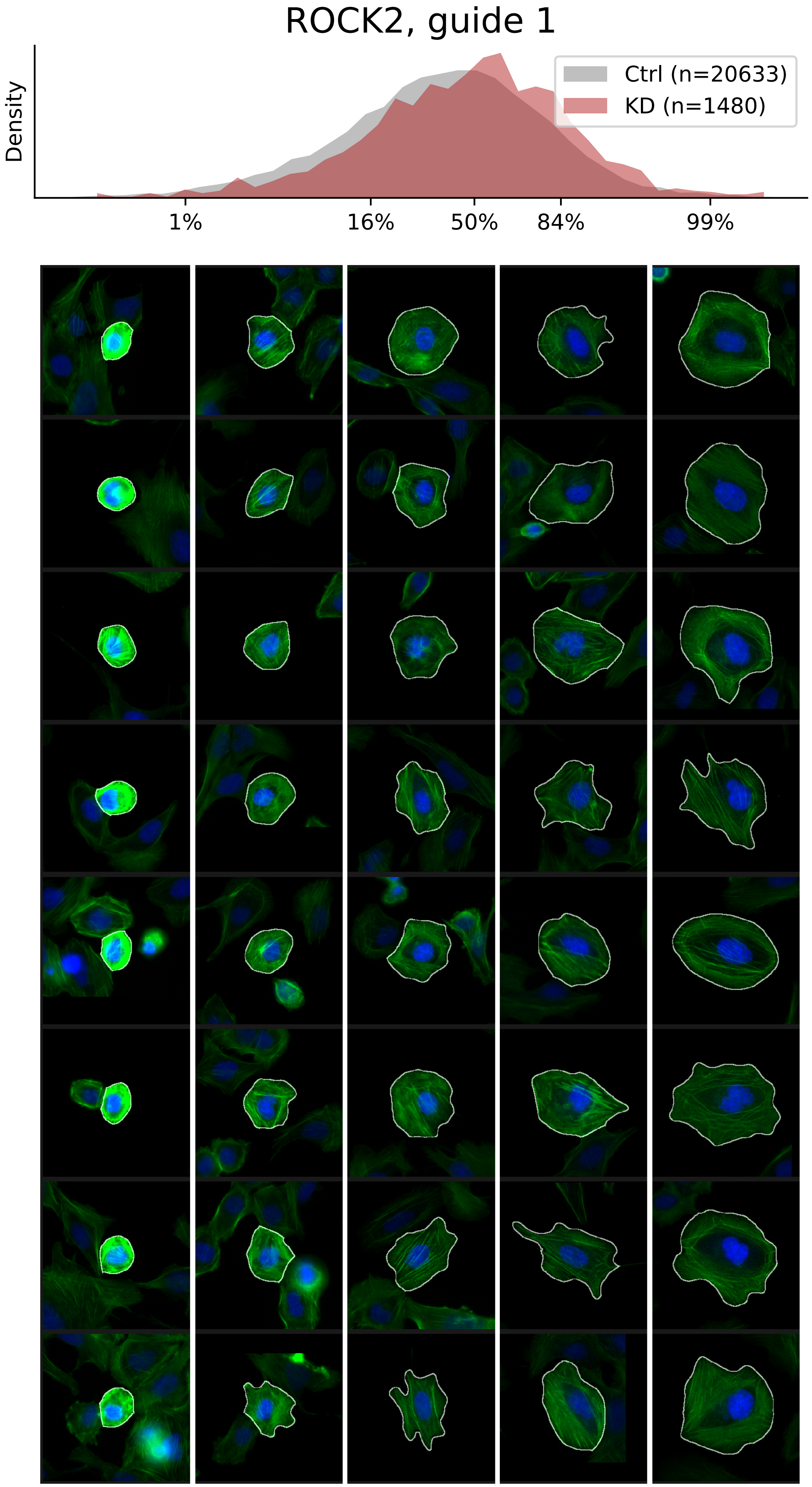

### VIEWS_iAD_ANLN_rep2.pdf

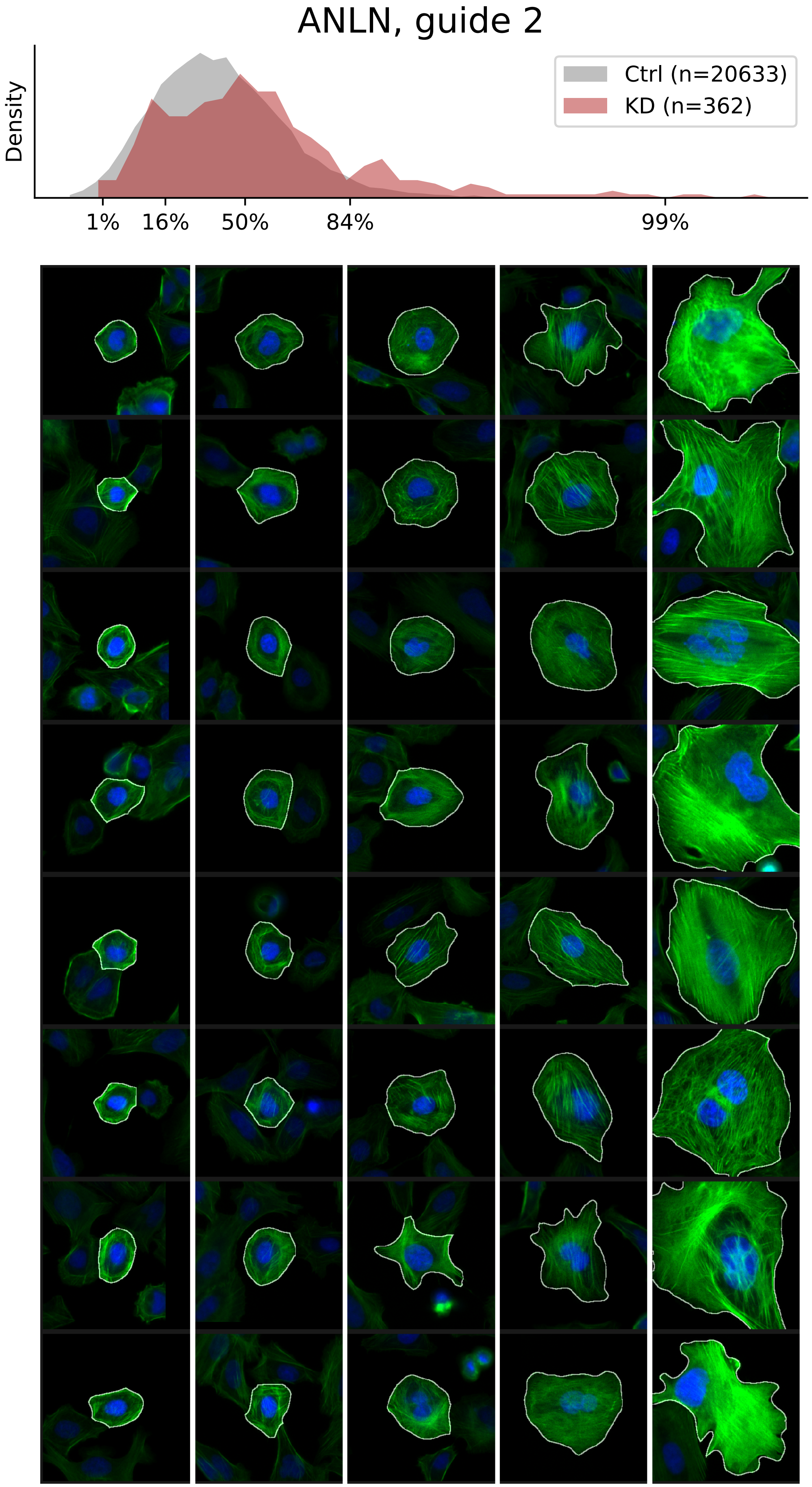

### VIEWS_iAD_ANLN_rep3.pdf

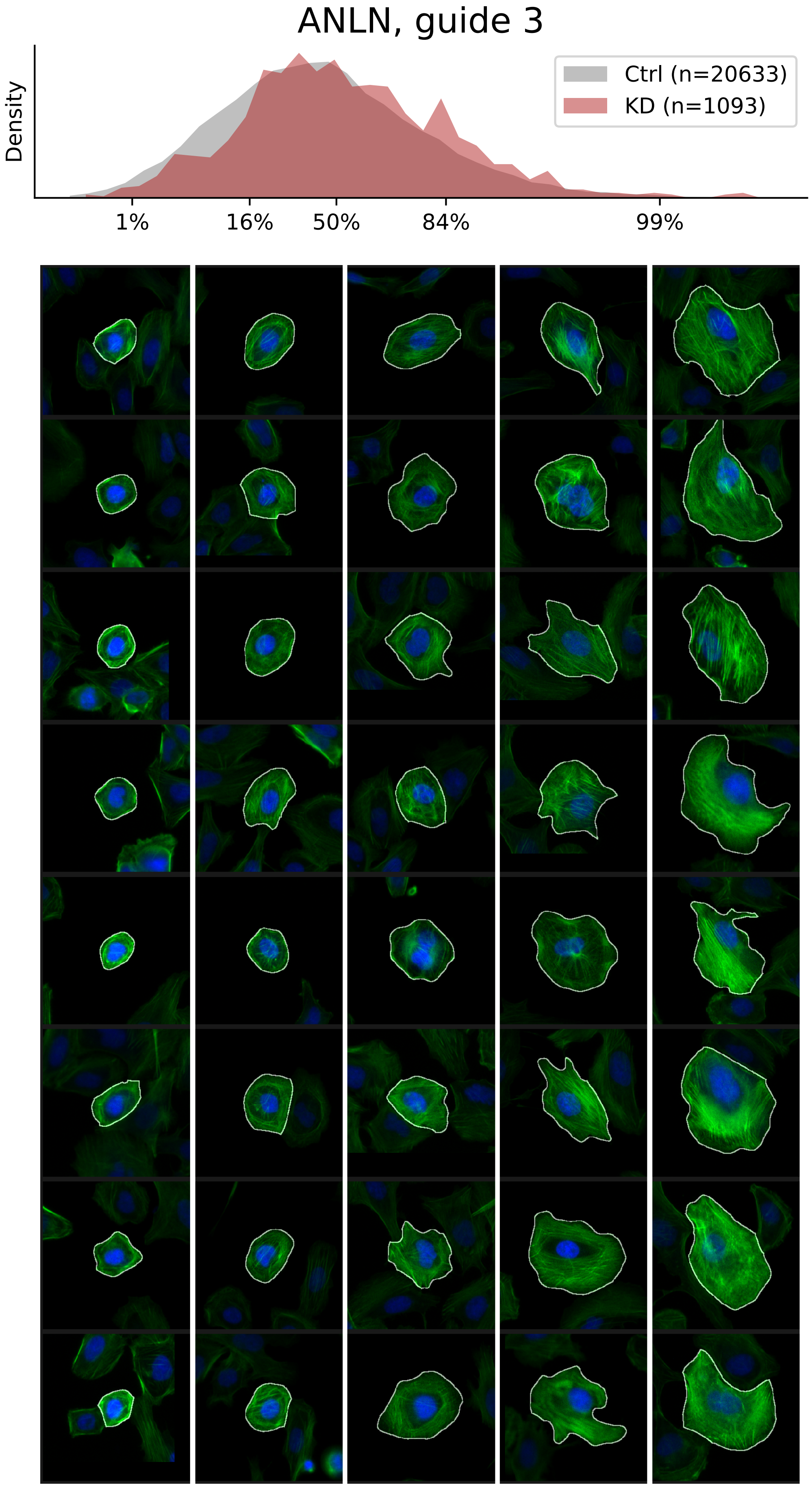

### VIEWS_iAD_BRK1_rep2.pdf

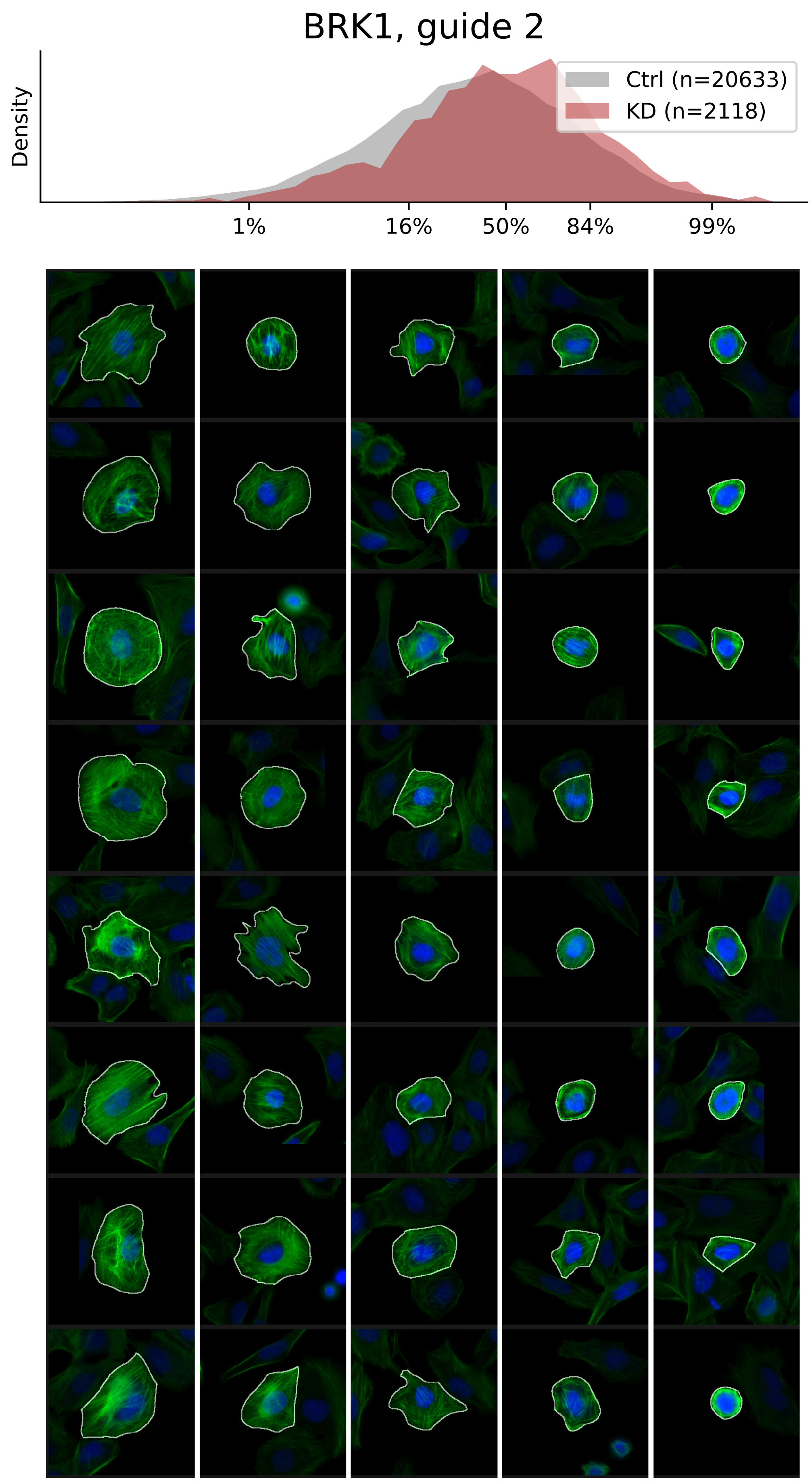

### VIEWS_iAD_BRK1_rep3.pdf

#### BRK1, guide 3

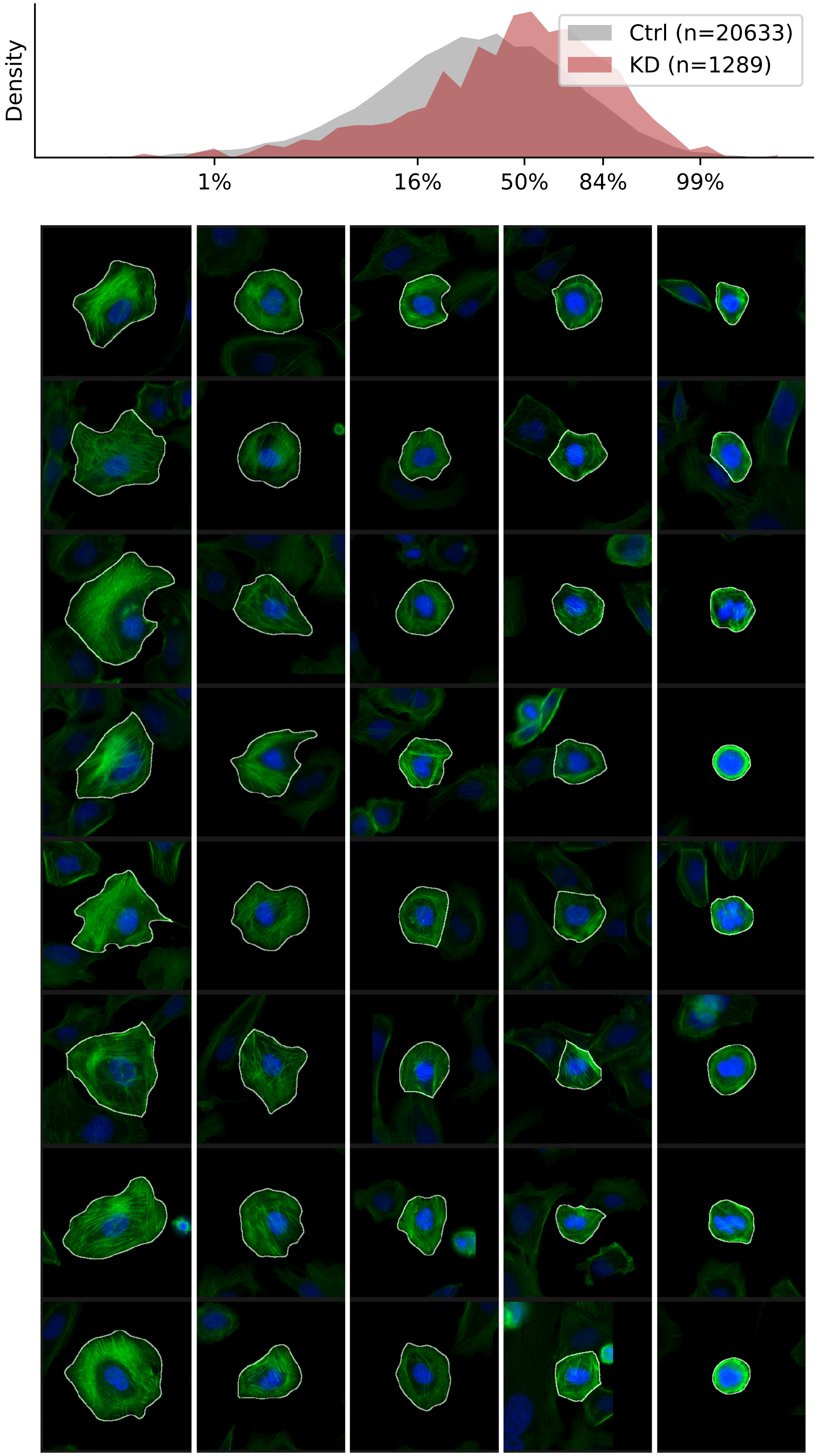

### VIEWS_iAD_CAPZA1_rep2.pdf

#### CAPZA1, guide 2

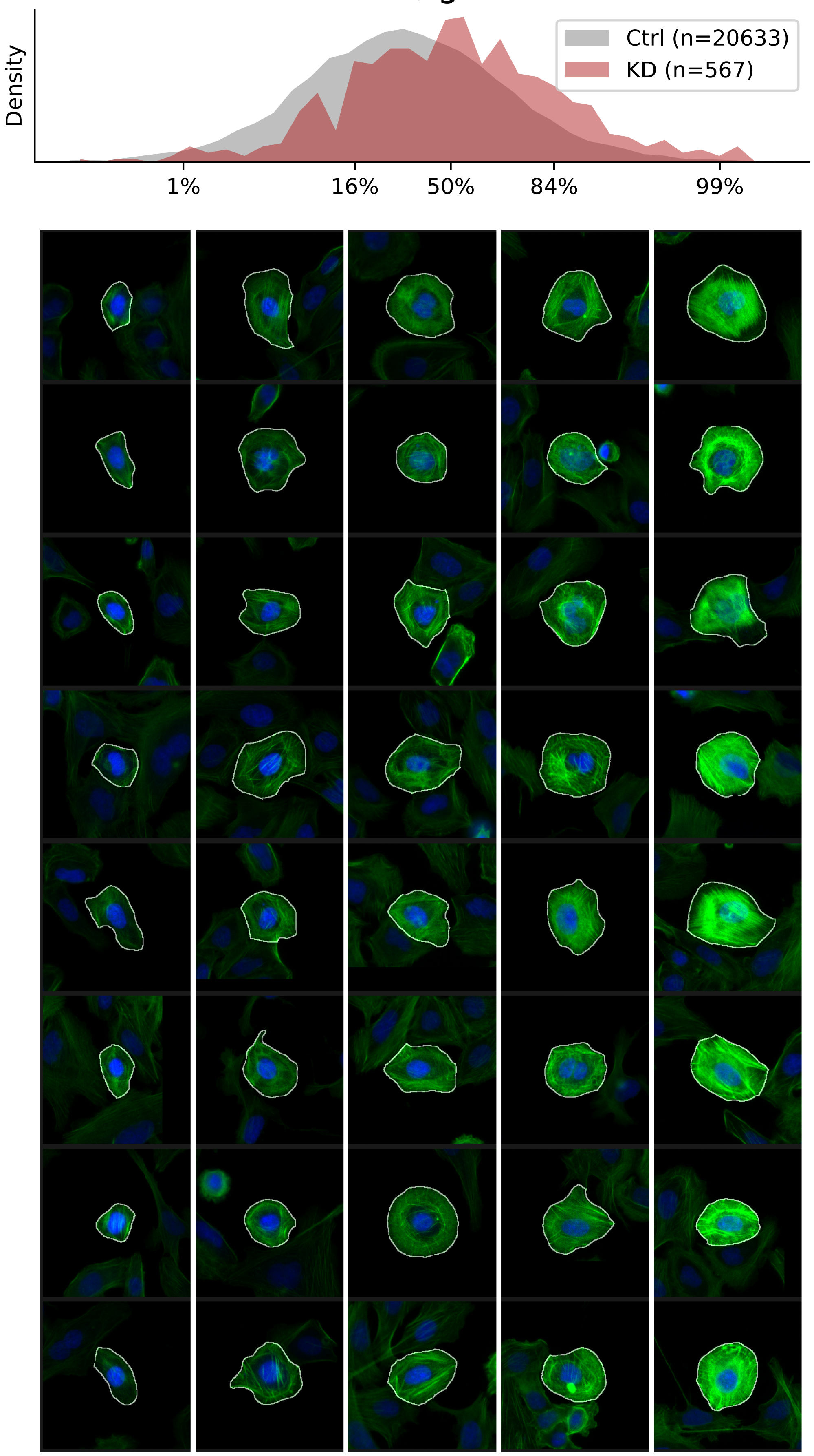

### VIEWS_iAD_CAPZA1_rep3.pdf

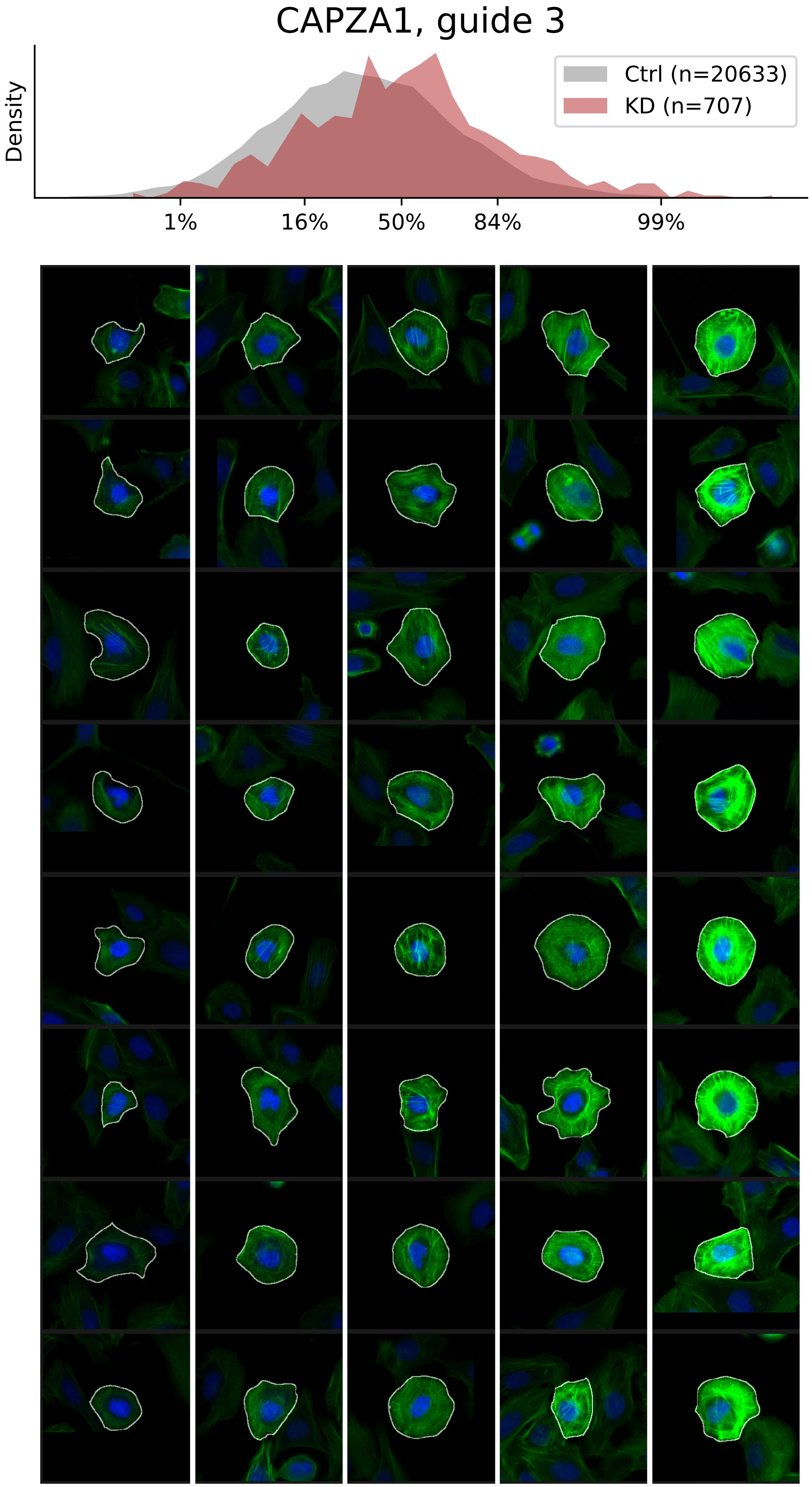

### VIEWS_iAD_CAPZB_rep2.pdf

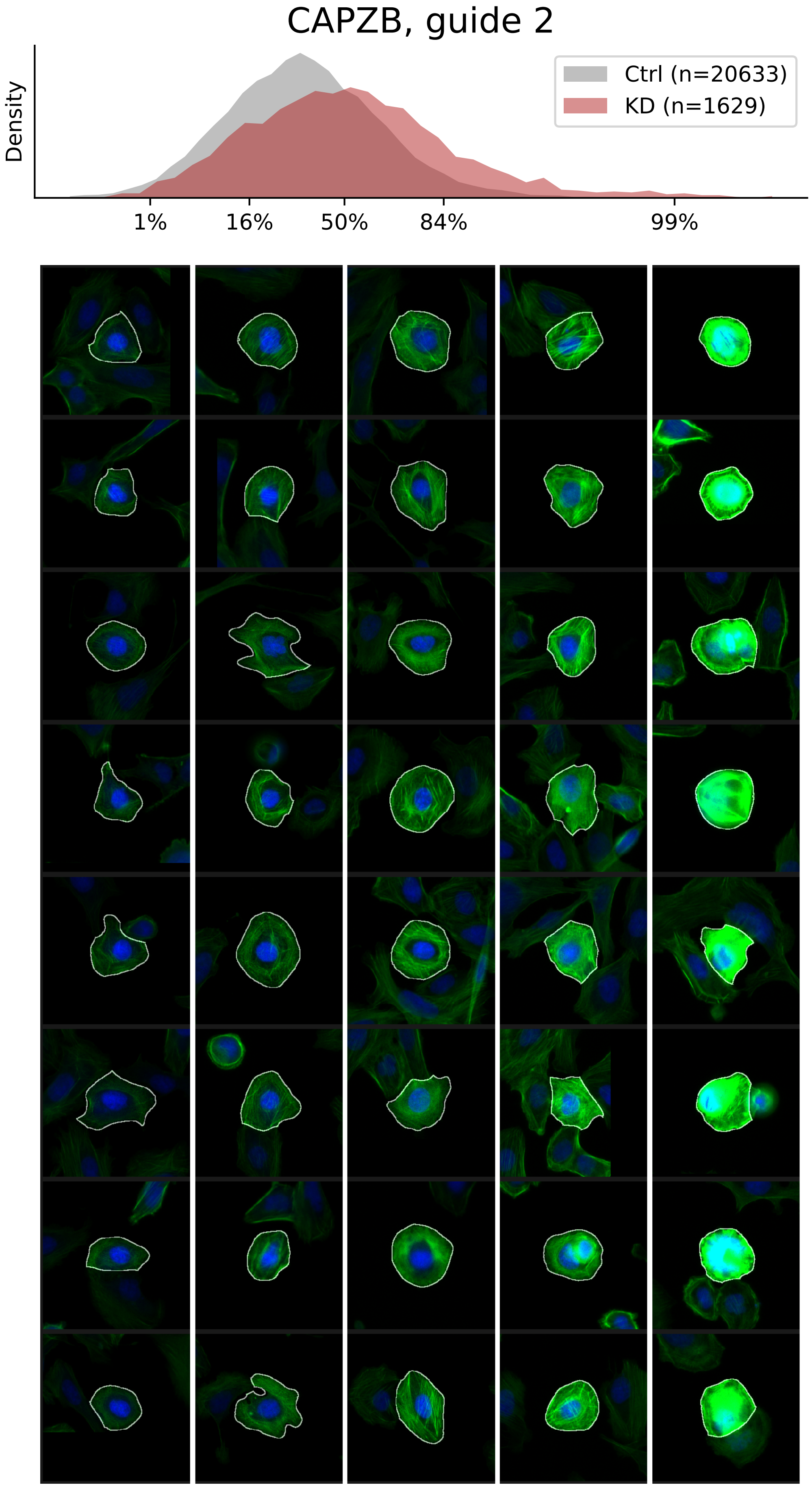

### VIEWS_iAD_CAPZB_rep3.pdf

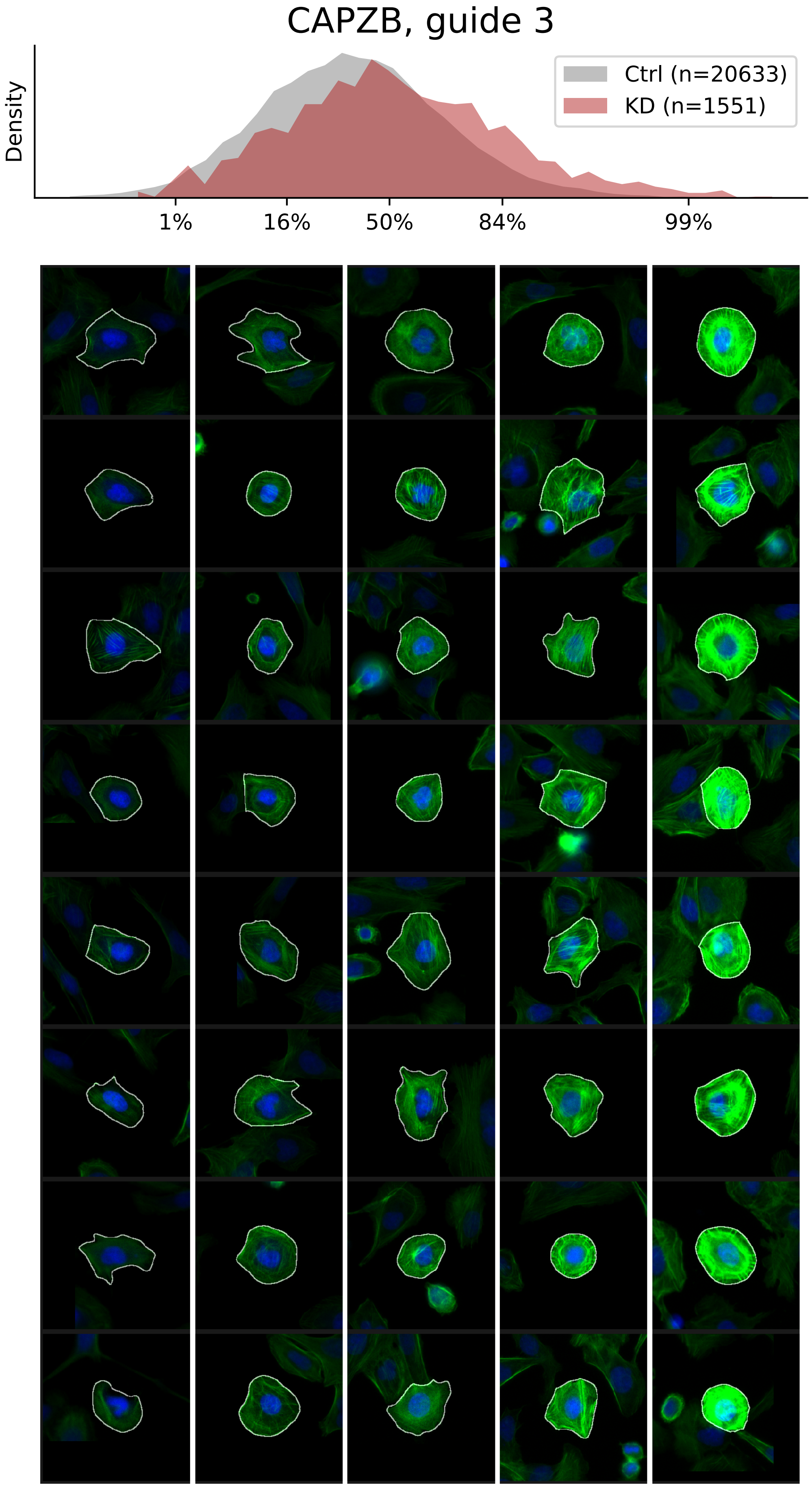

### VIEWS_iAD_FLII_rep2.pdf

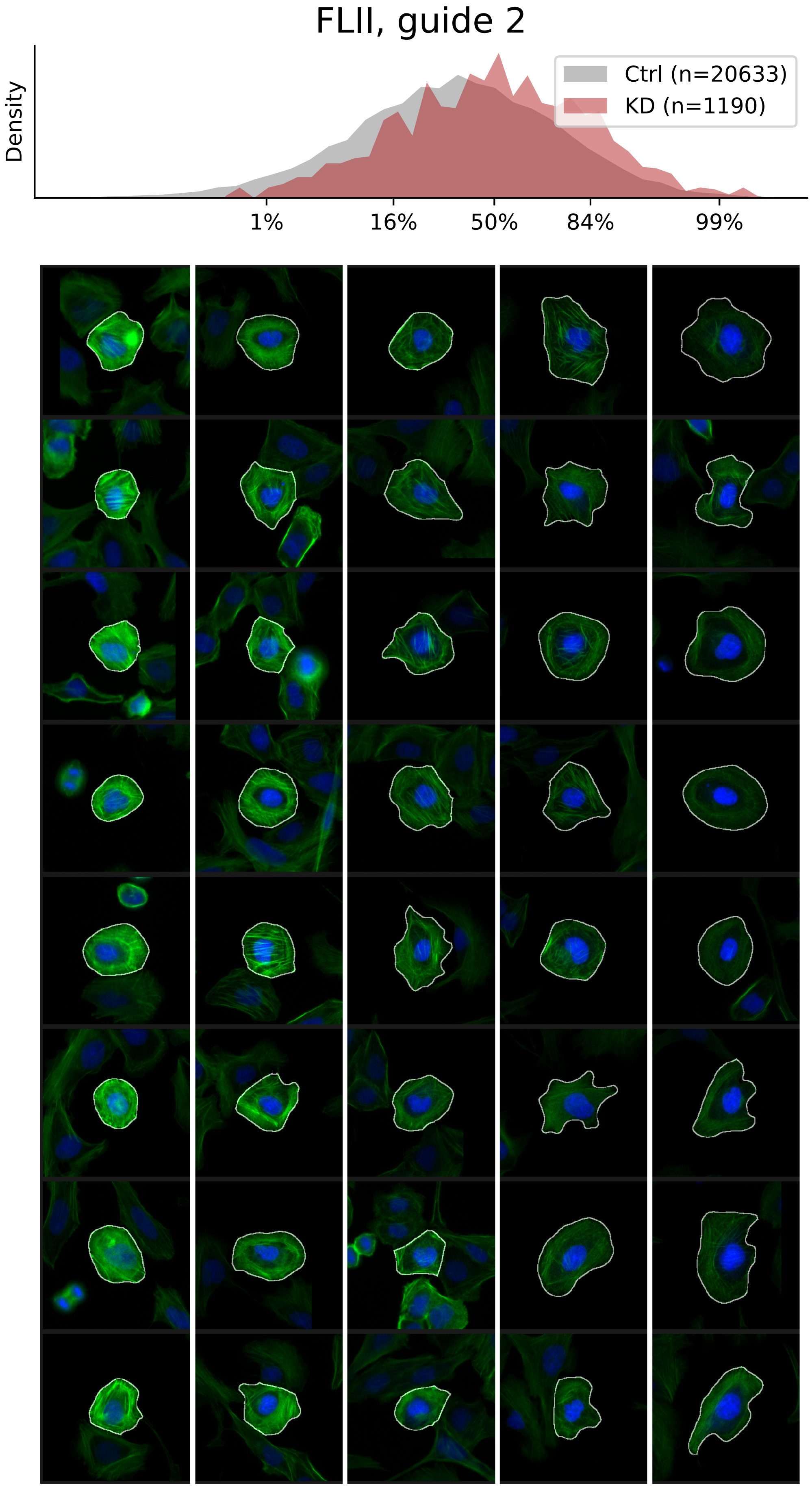

### VIEWS_iAD_FLII_rep3.pdf

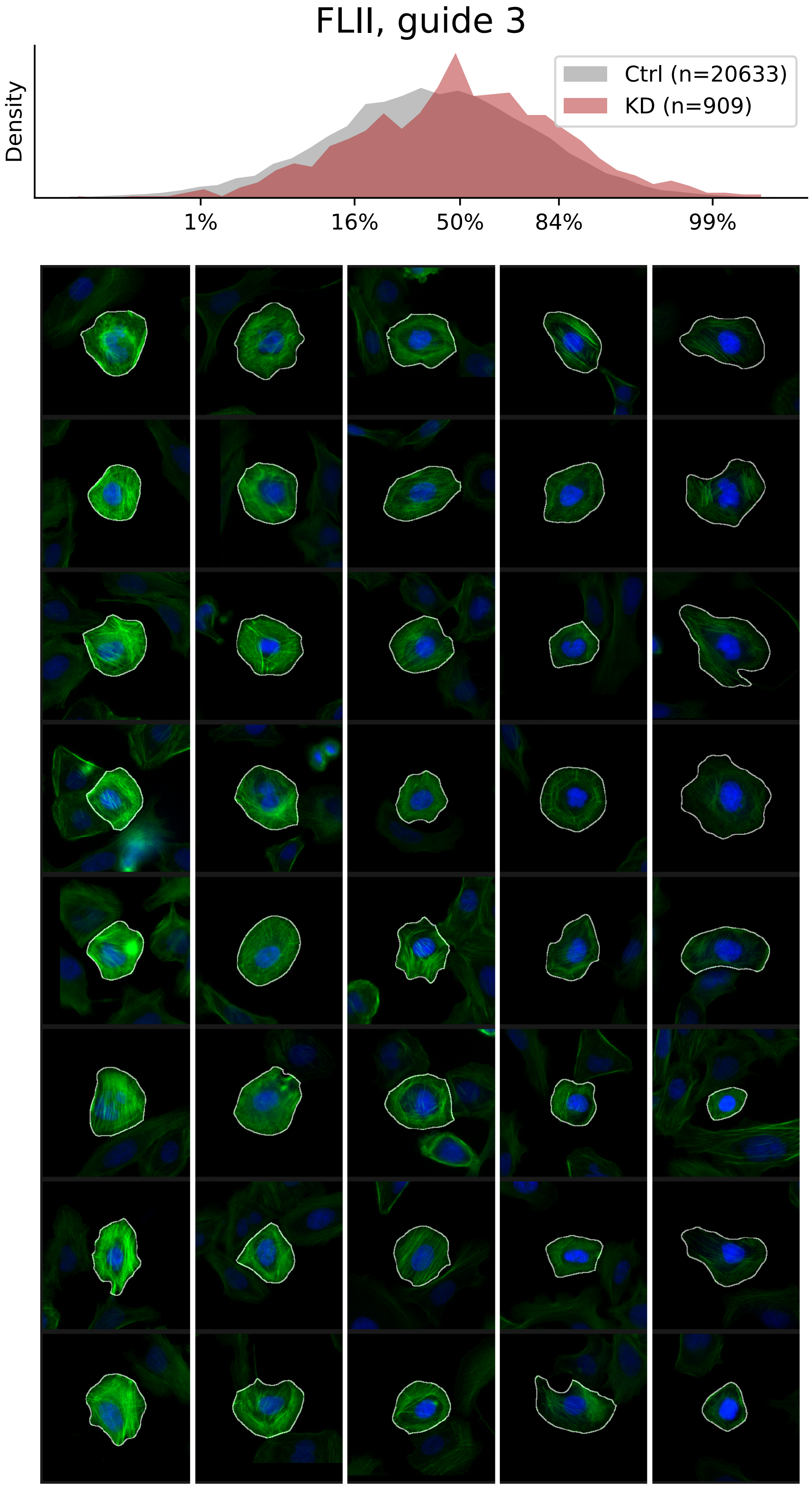

### VIEWS_iAD_ILK_rep1.pdf

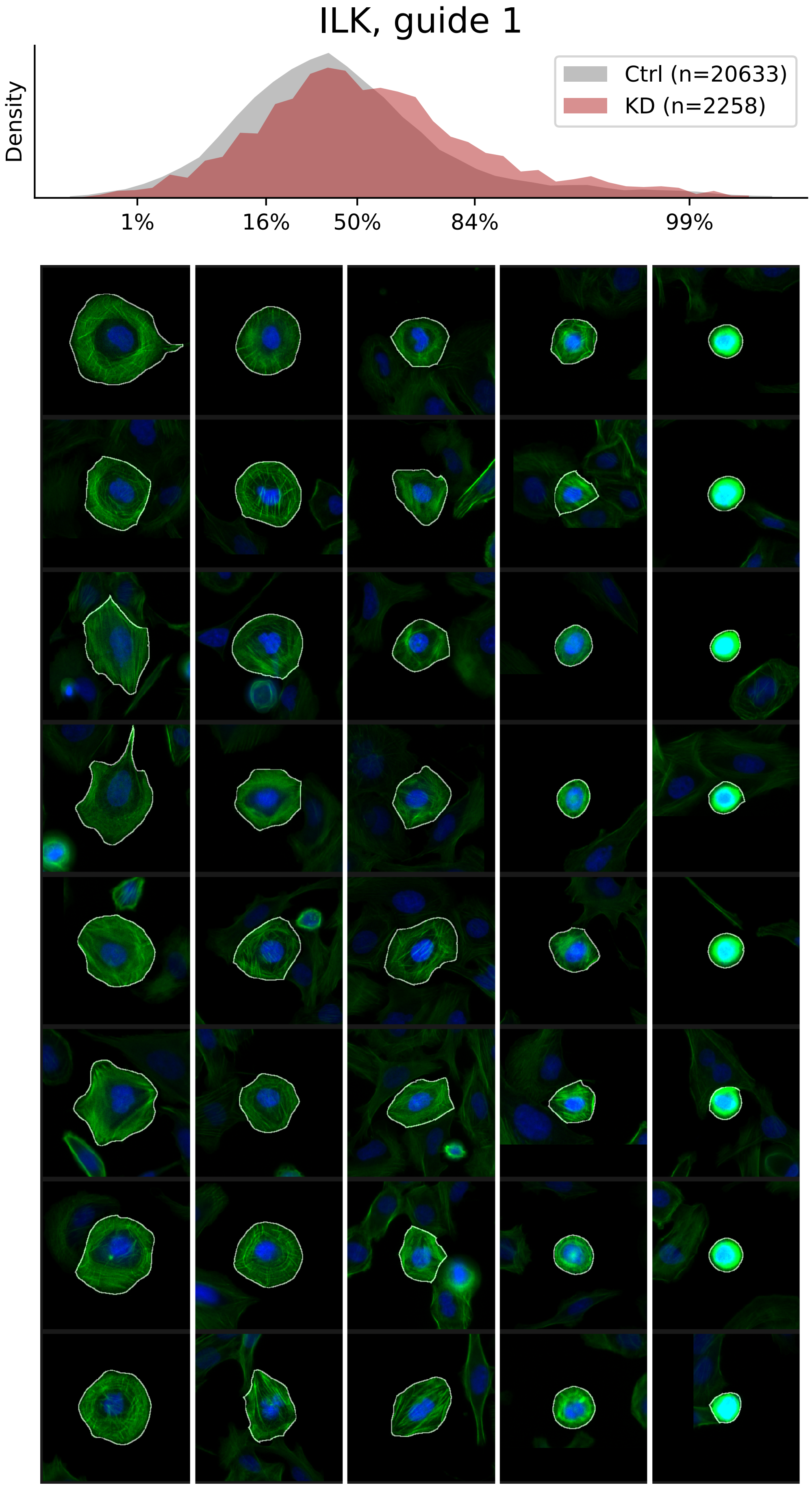

### VIEWS_iAD_SETD3_rep1.pdf

#### SETD3, guide 1

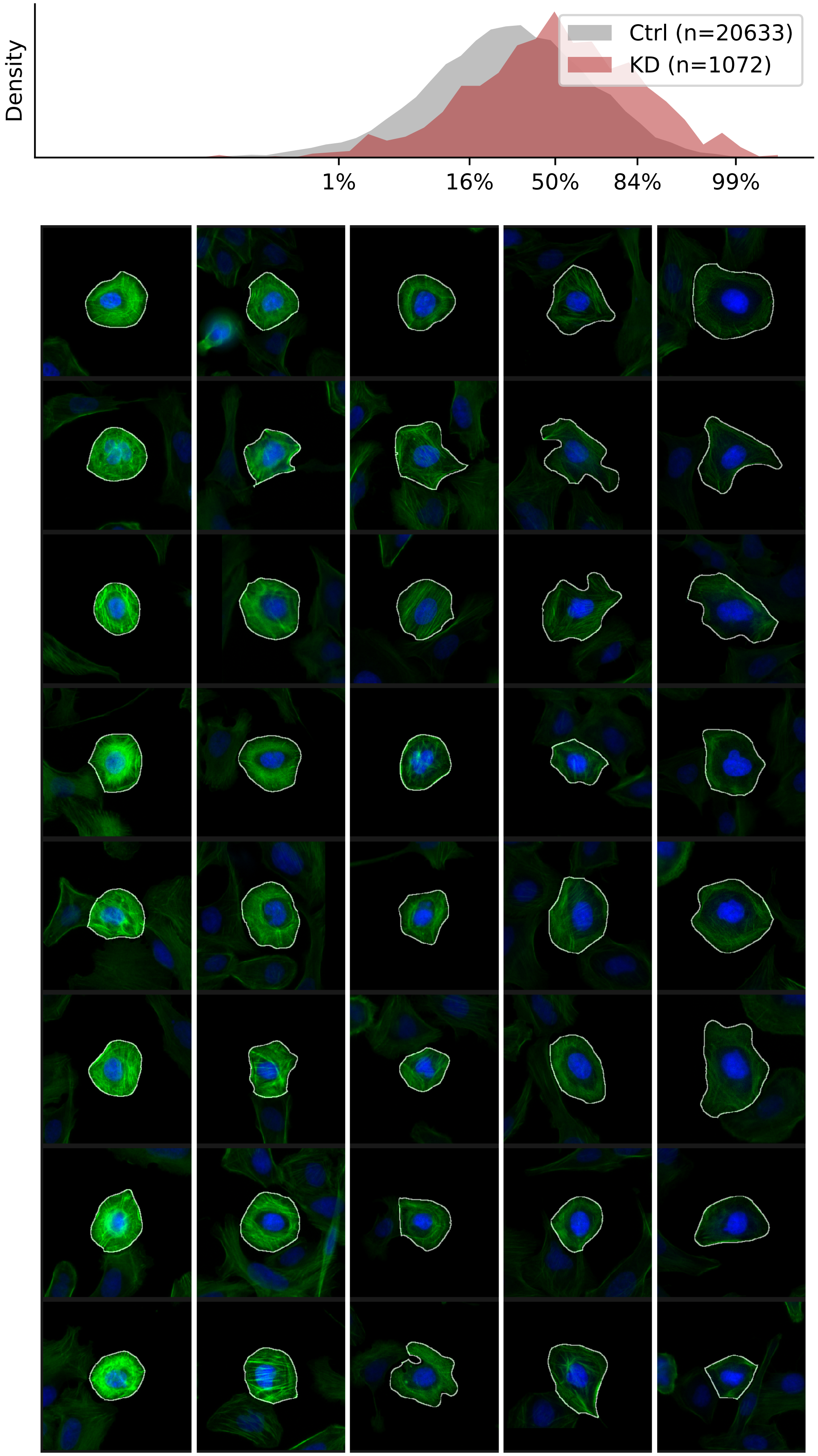

### VIEWS_iAD_SETD3_rep2.pdf

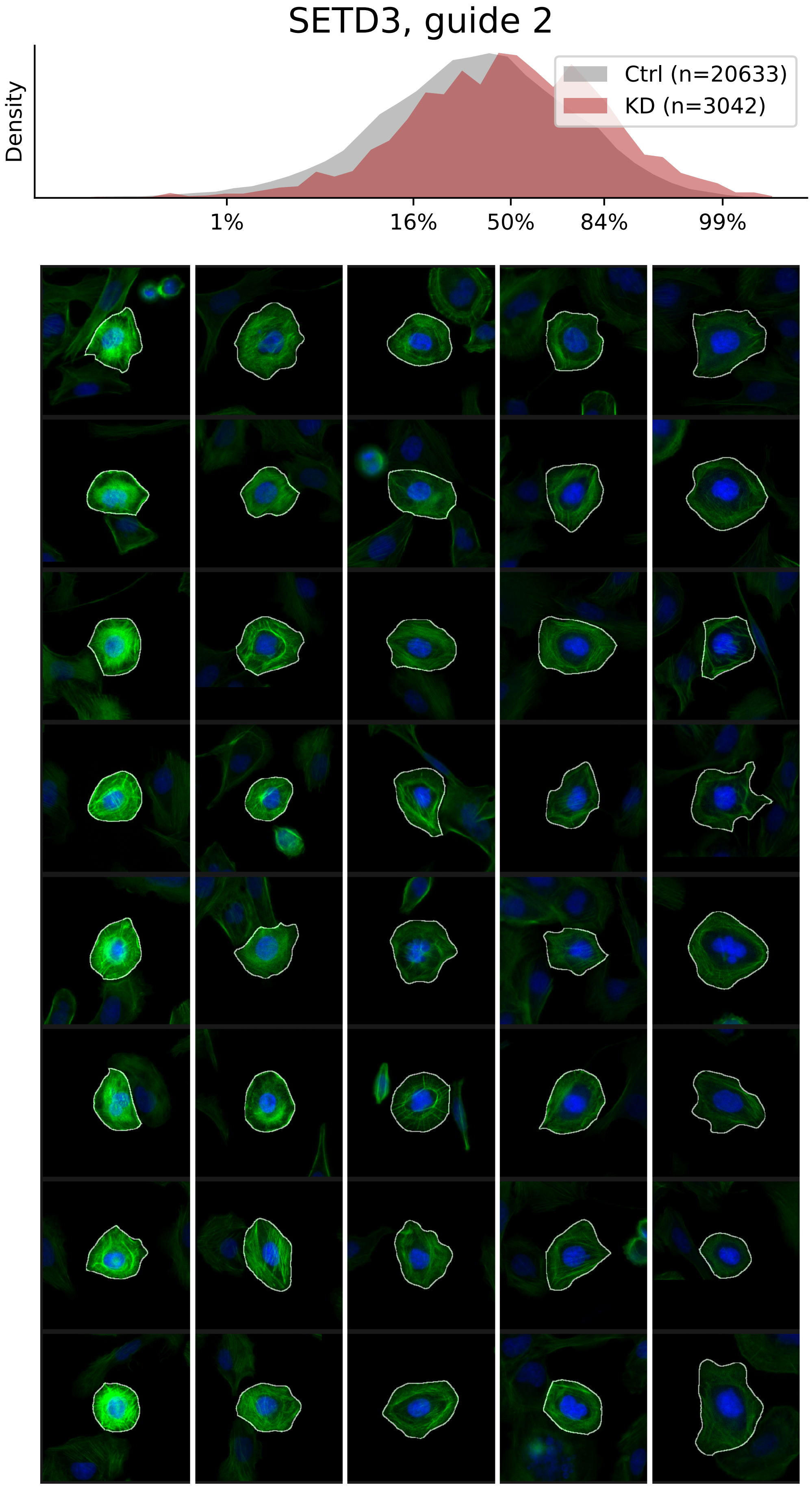

### VIEWS_iAD_SETD3_rep3.pdf

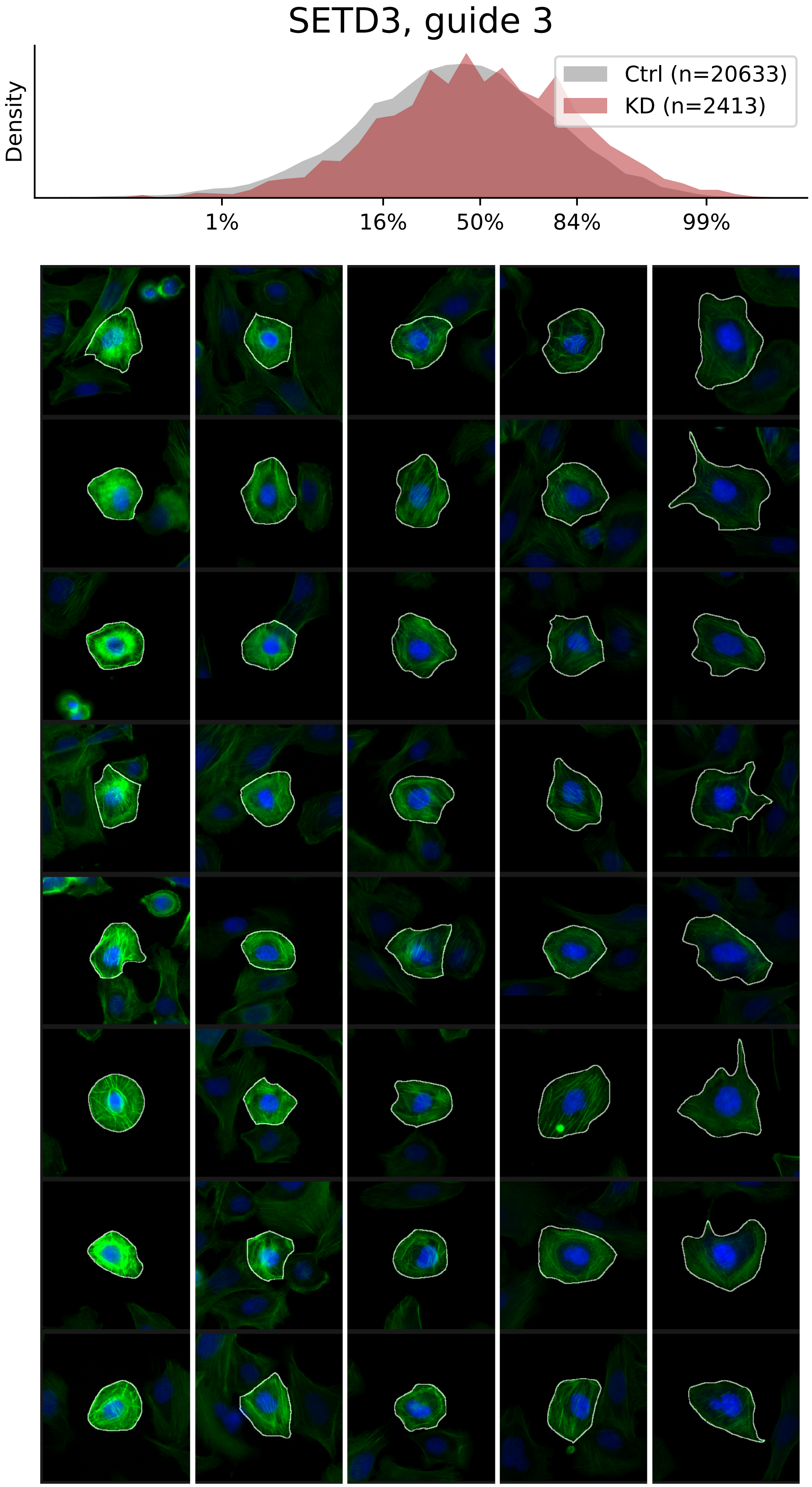

### VIEWS_iAD_VCL_rep1.pdf

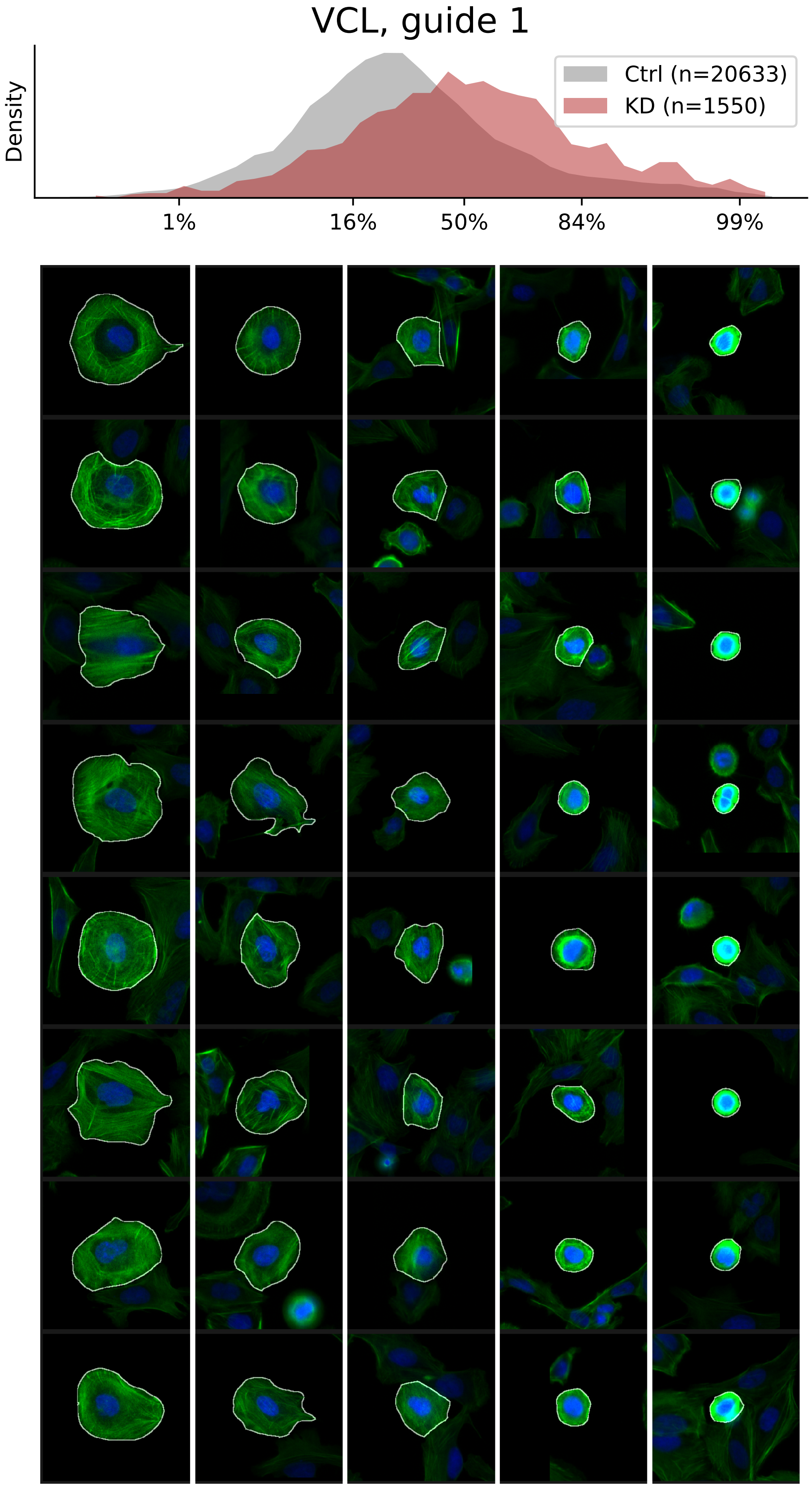

### VIEWS_iAD_VCL_rep2.pdf

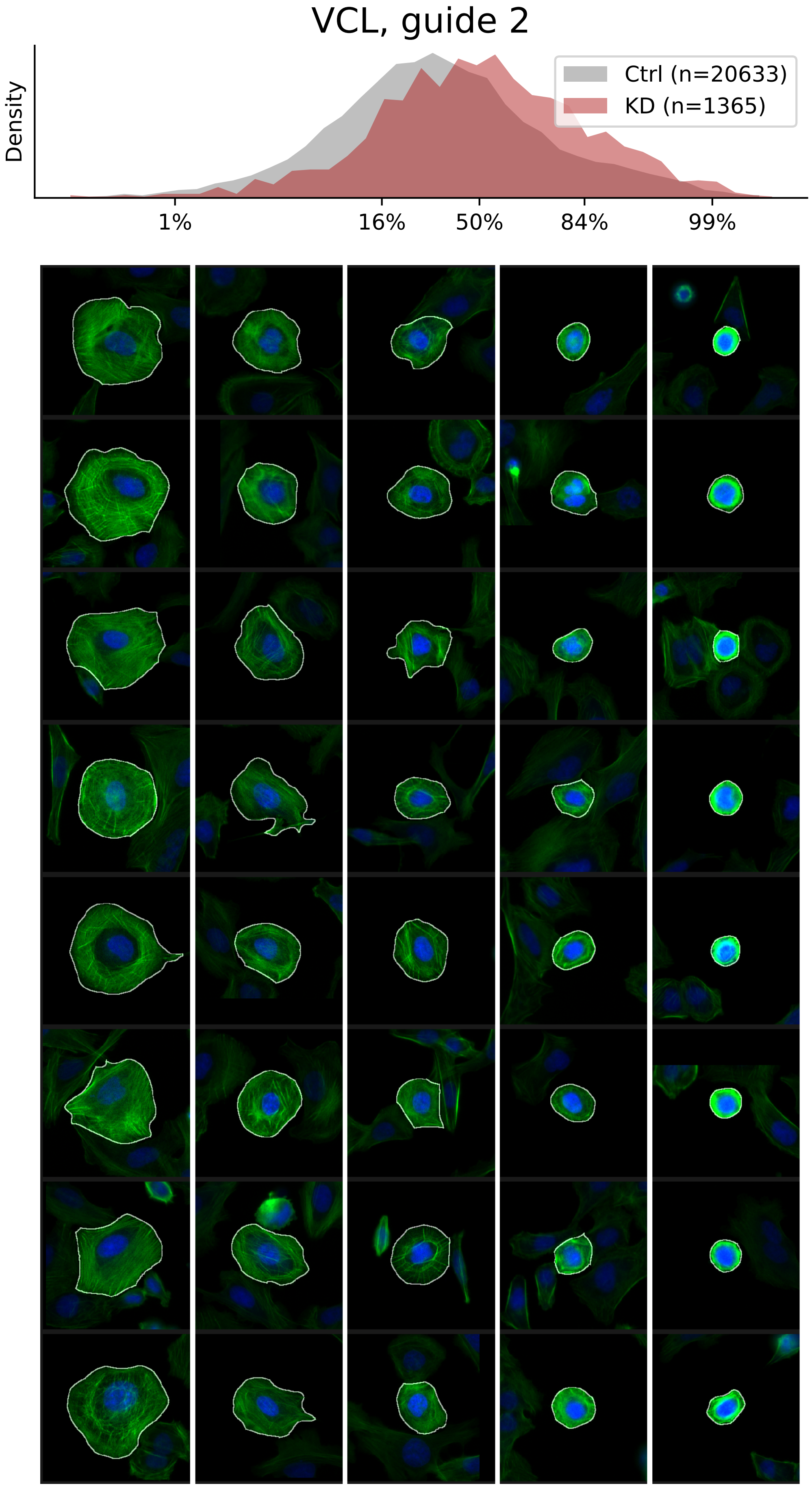

### VIEWS_iAD_VCL_rep3.pdf

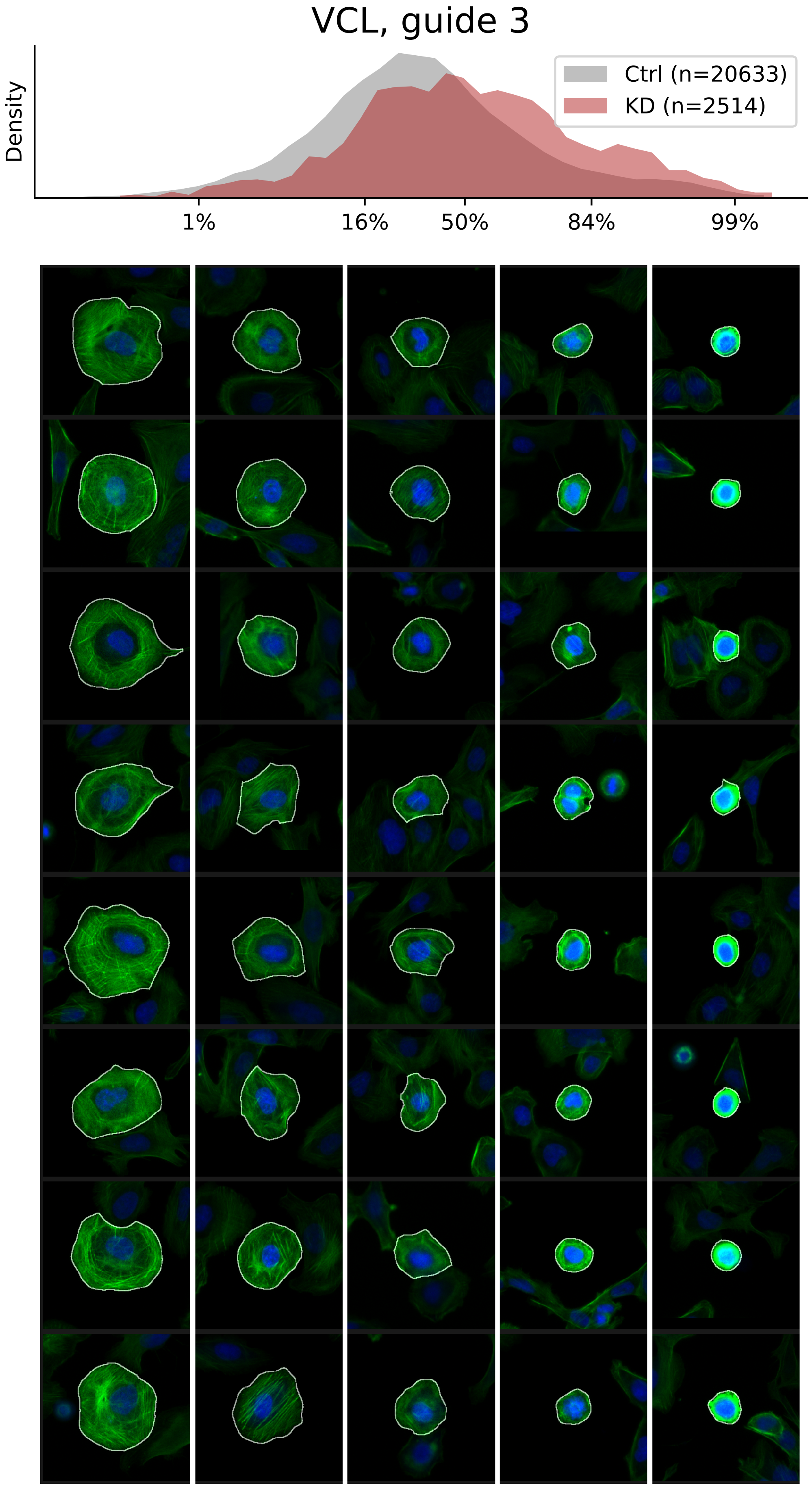
