## Supplementary figures and images for "Mapping variation in the morphological landscape of human cells with optical pooled CRISPRi screening"

### VIEWS_fPC-LD-verified_ACTR3_rep3.pdf

#### ACTR3, guide 3

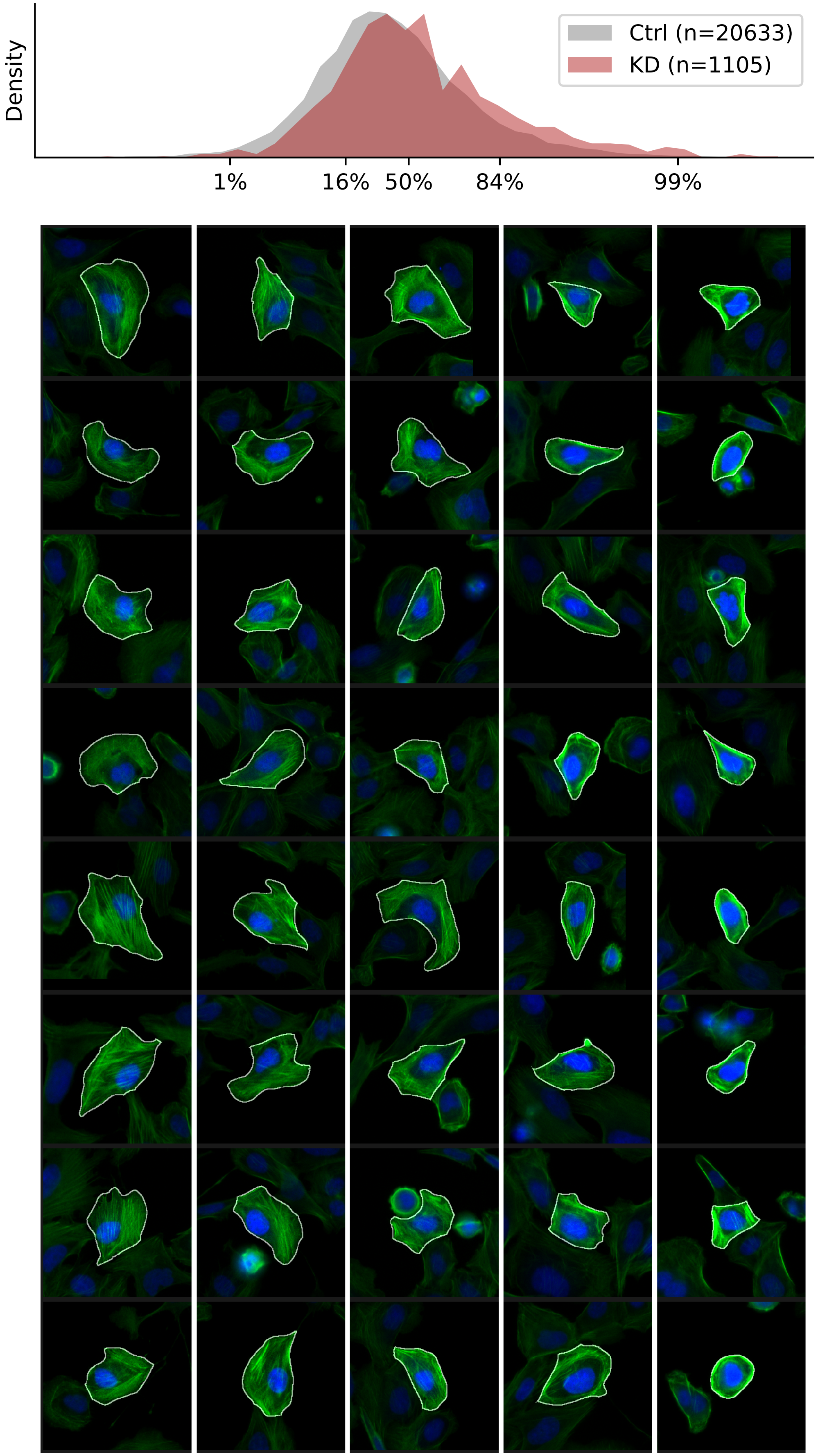

### VIEWS_fPC-LD-verified_LASP1_rep2.pdf

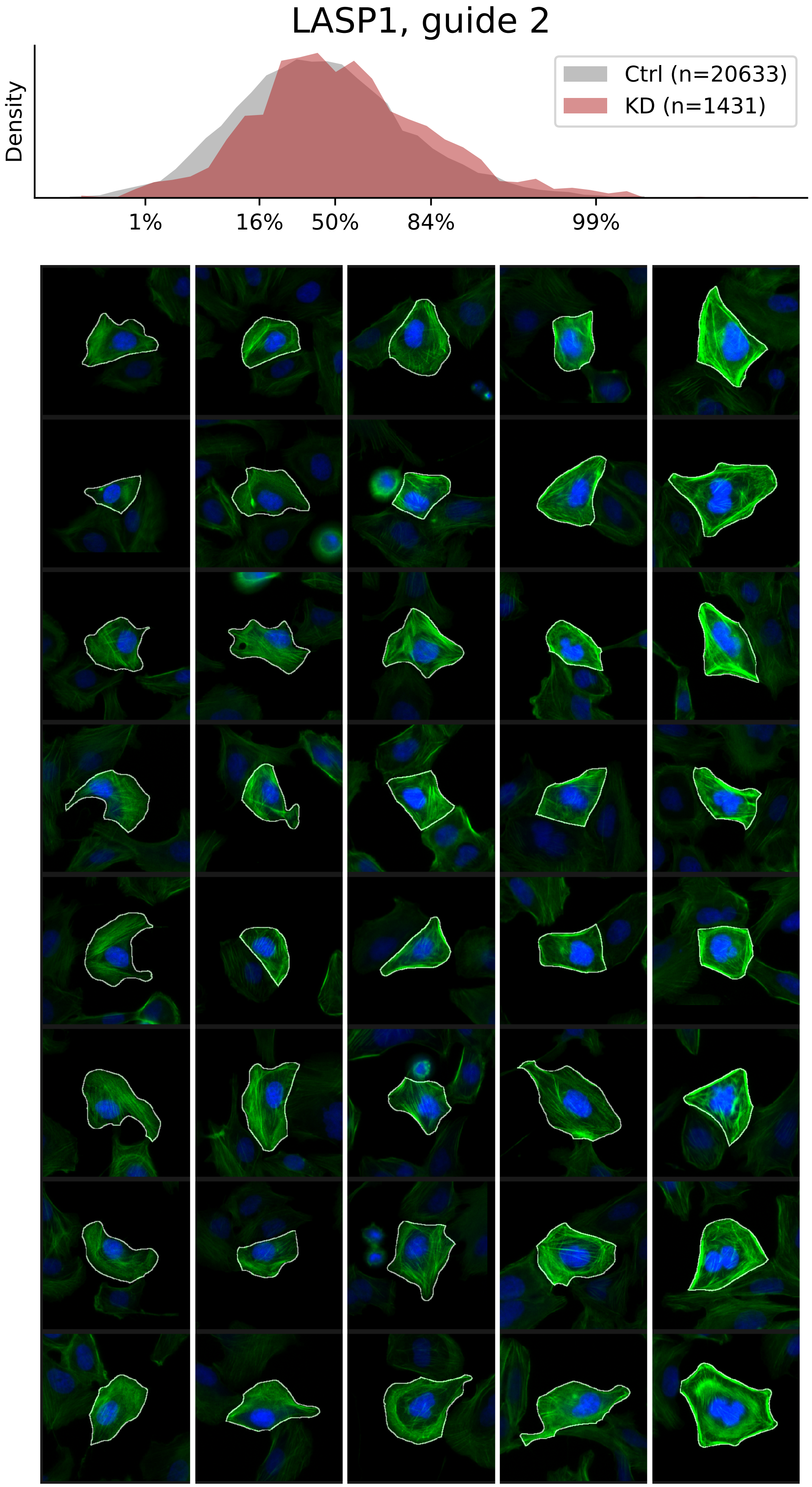

### VIEWS_fPC-LD-verified_MYO9B_rep1.pdf

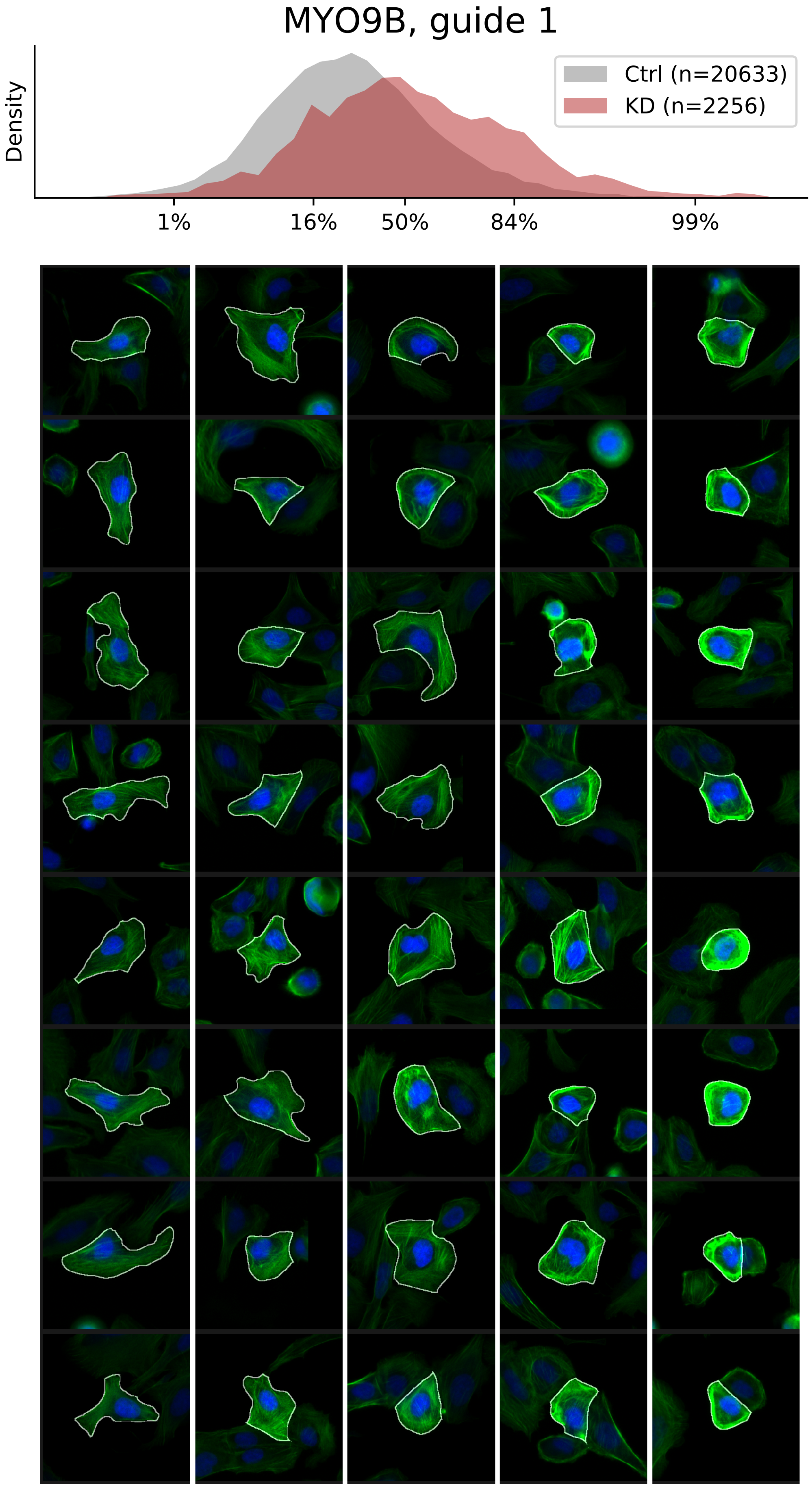

### VIEWS_fPC-LD-verified_NCKAP1_rep2.pdf

#### NCKAP1, guide 2

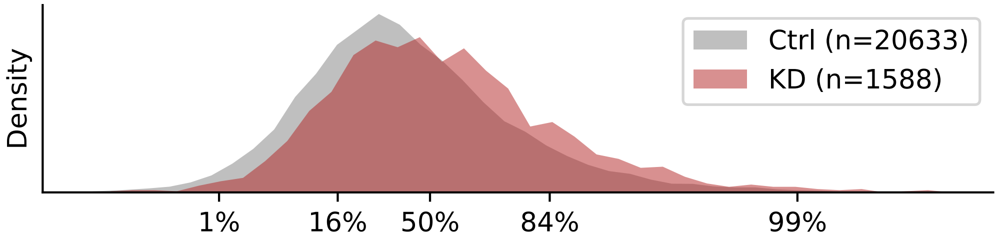


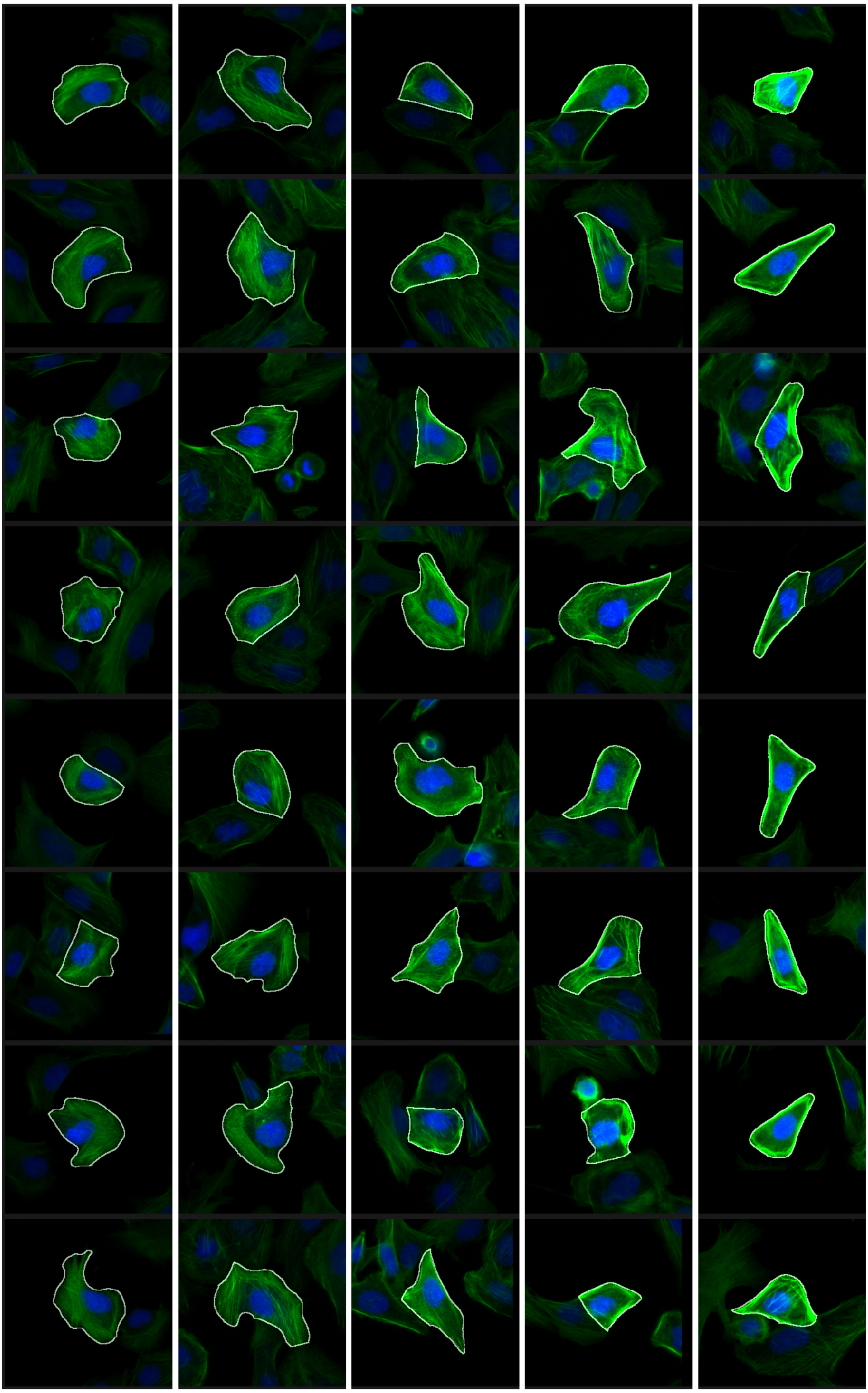

### VIEWS_fPC-LD-verified_SIPA1L3_rep3.pdf

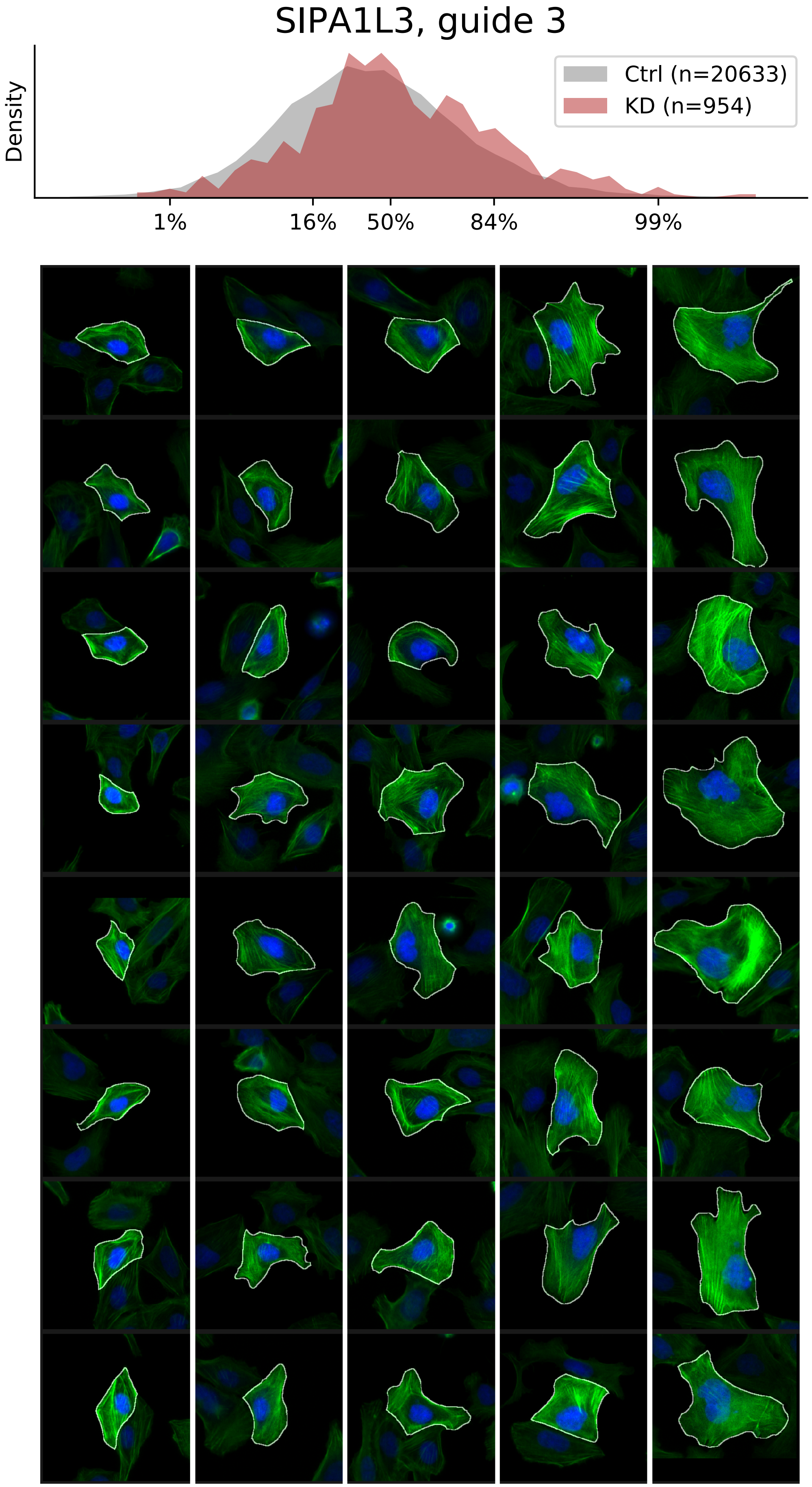

### VIEWS_fPC-LD-verified_TMOD3_rep2.pdf

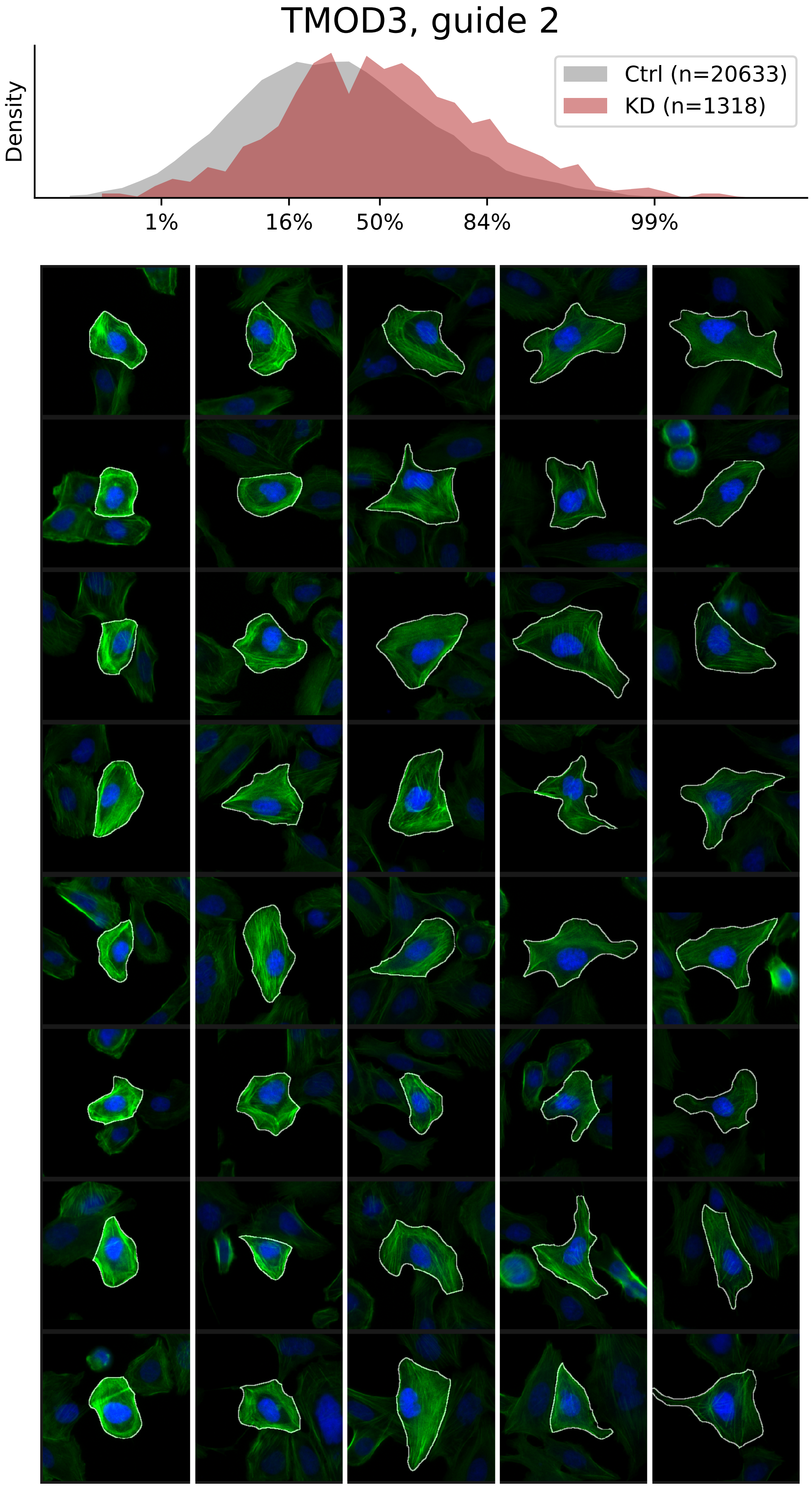

### VIEWS_fPC-LD-verified_TUBA1B_rep3.pdf

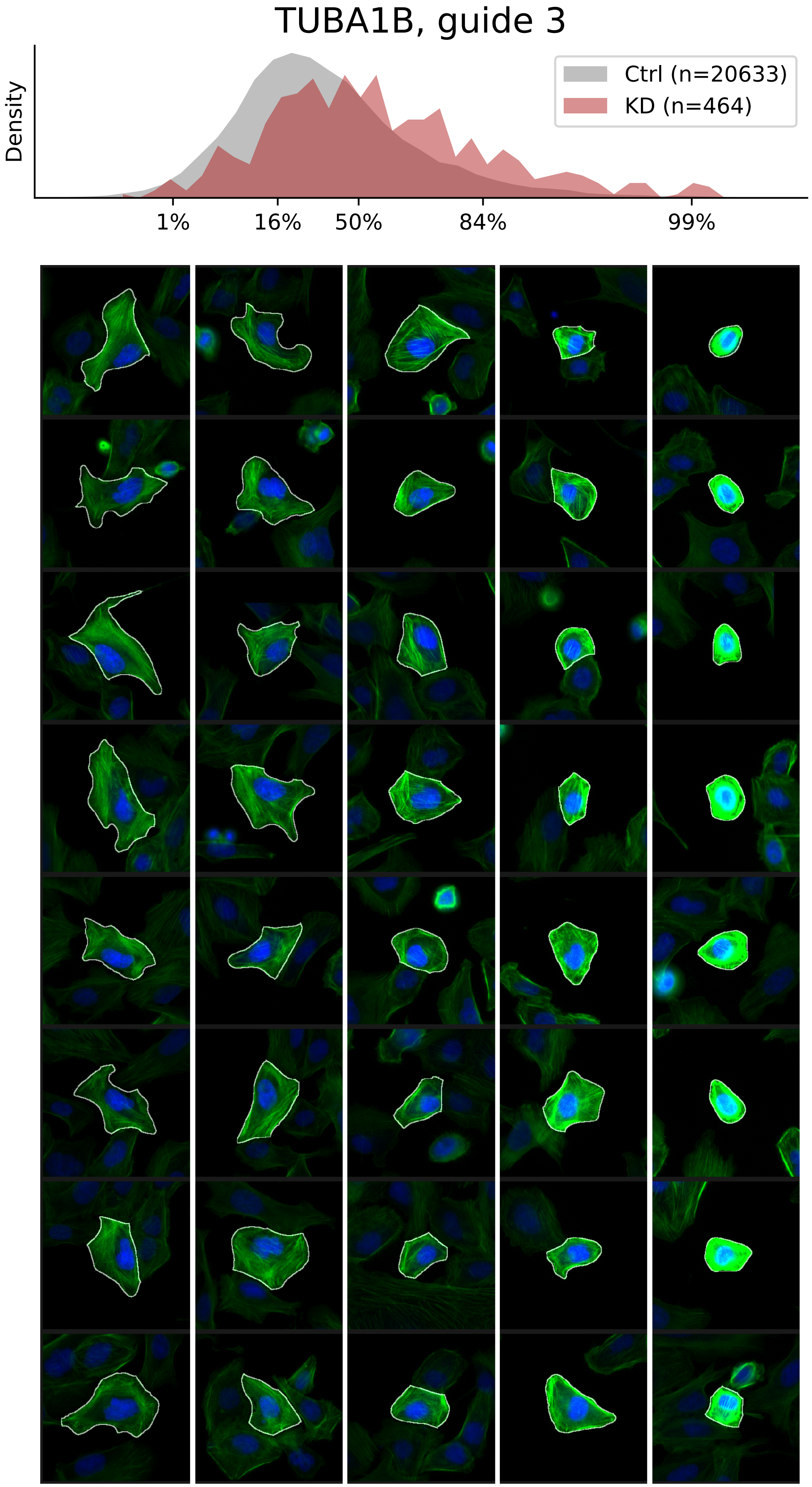

### VIEWS_fPC_ACTR2_rep3.pdf

#### ACTR2, guide 3

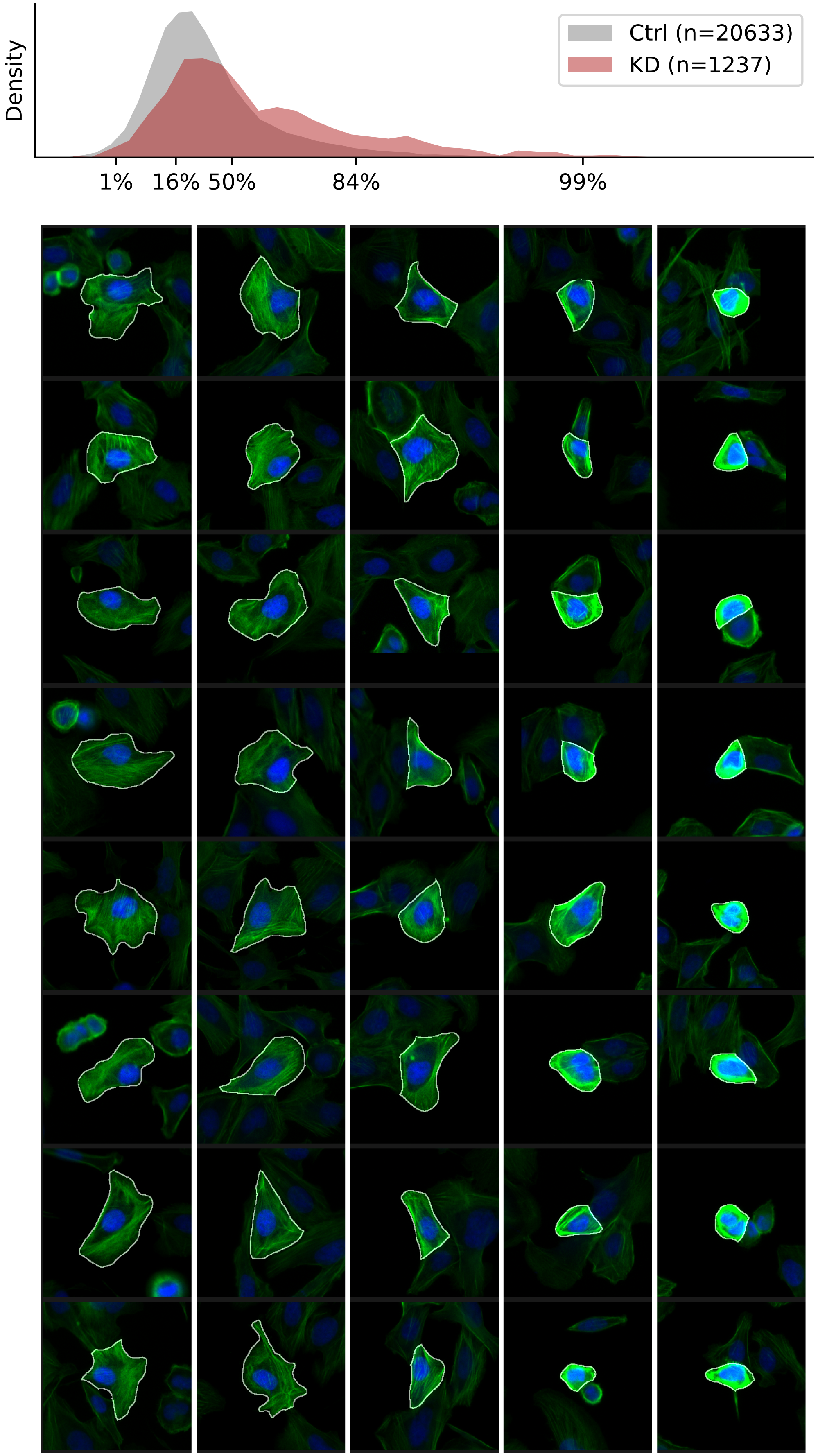

### VIEWS_fPC_ANLN_rep2.pdf

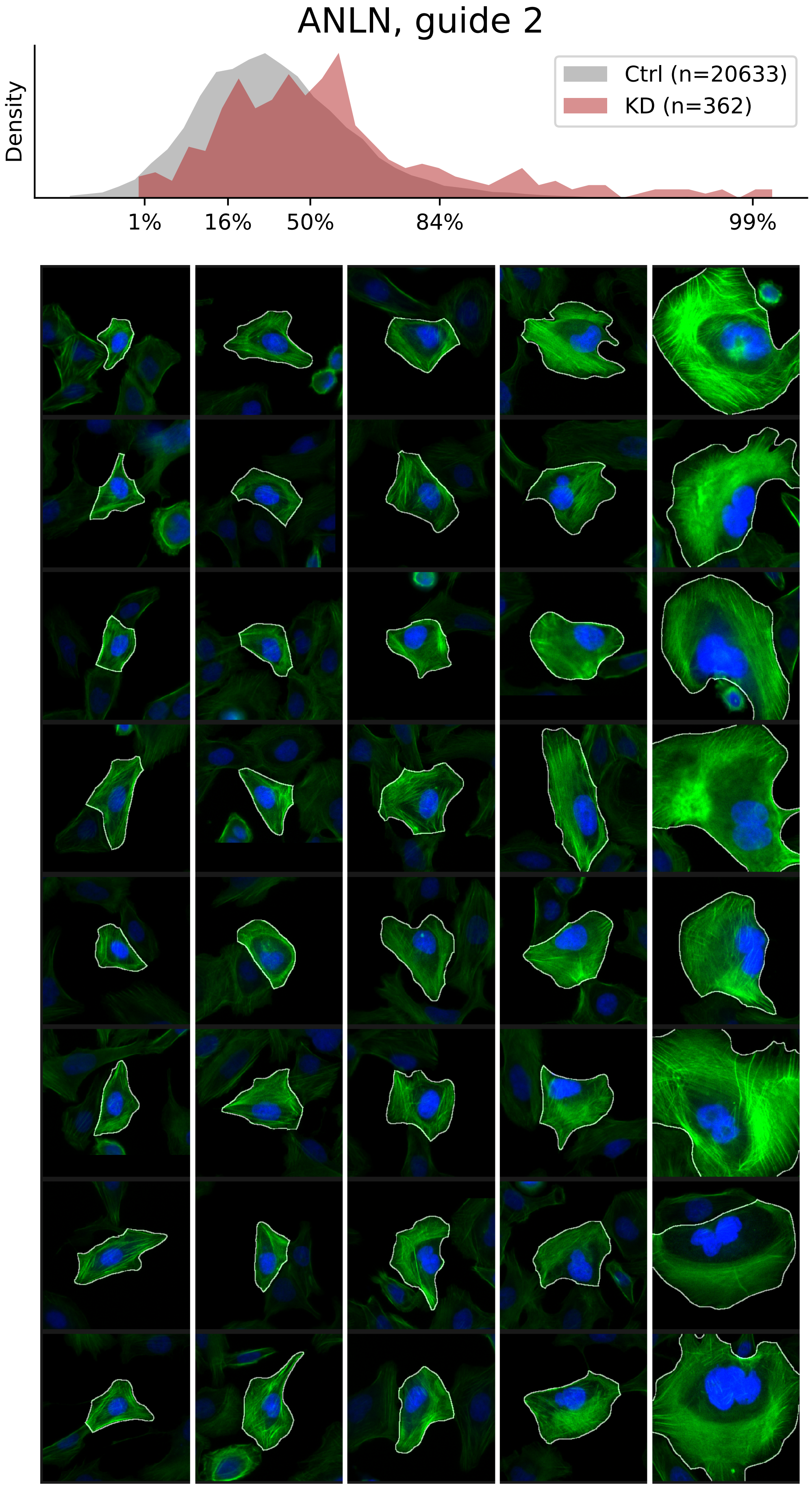

### VIEWS_fPC_ANLN_rep3.pdf

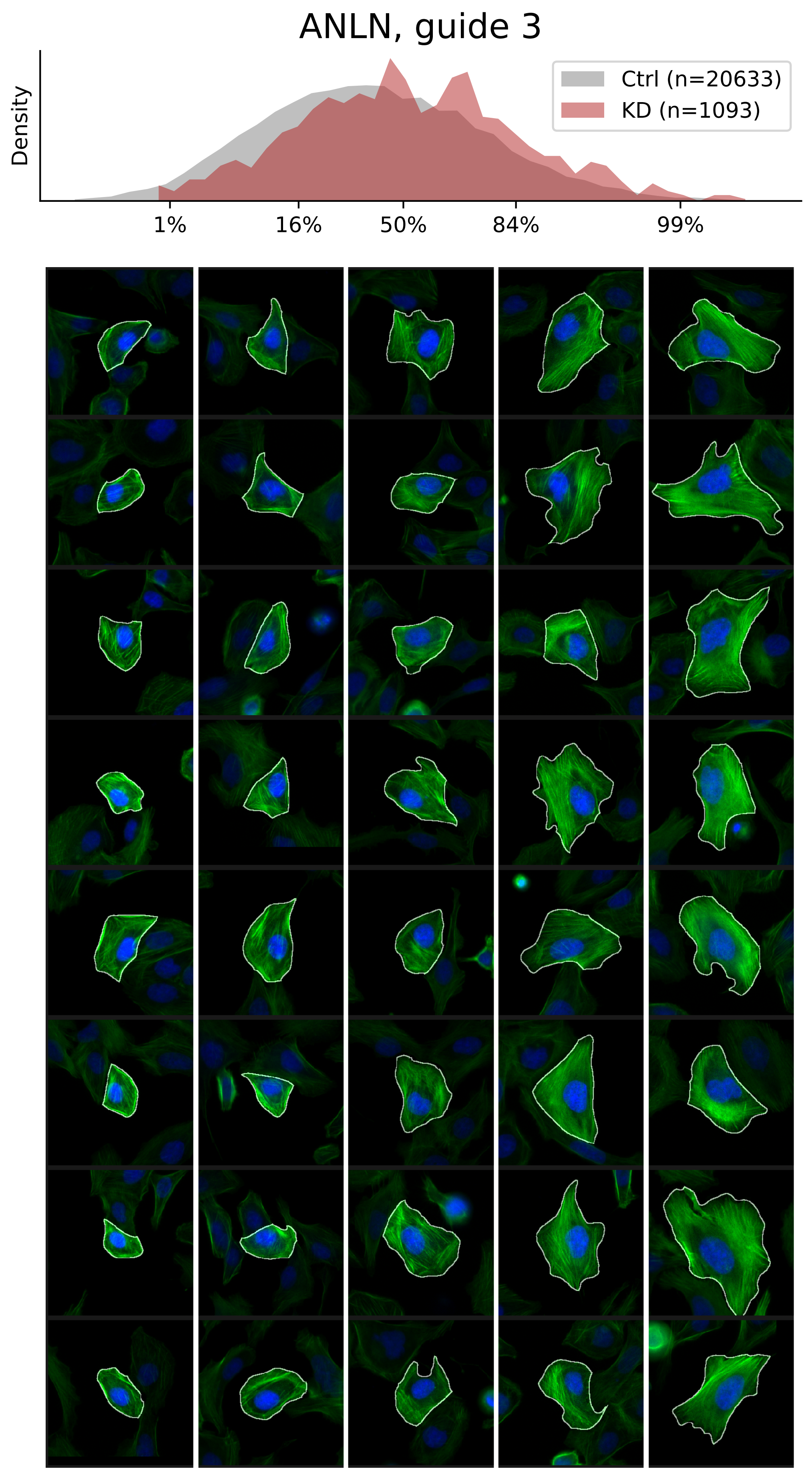

### VIEWS_fPC_ARHGEF7_rep1.pdf

#### ARHGEF7, guide 1

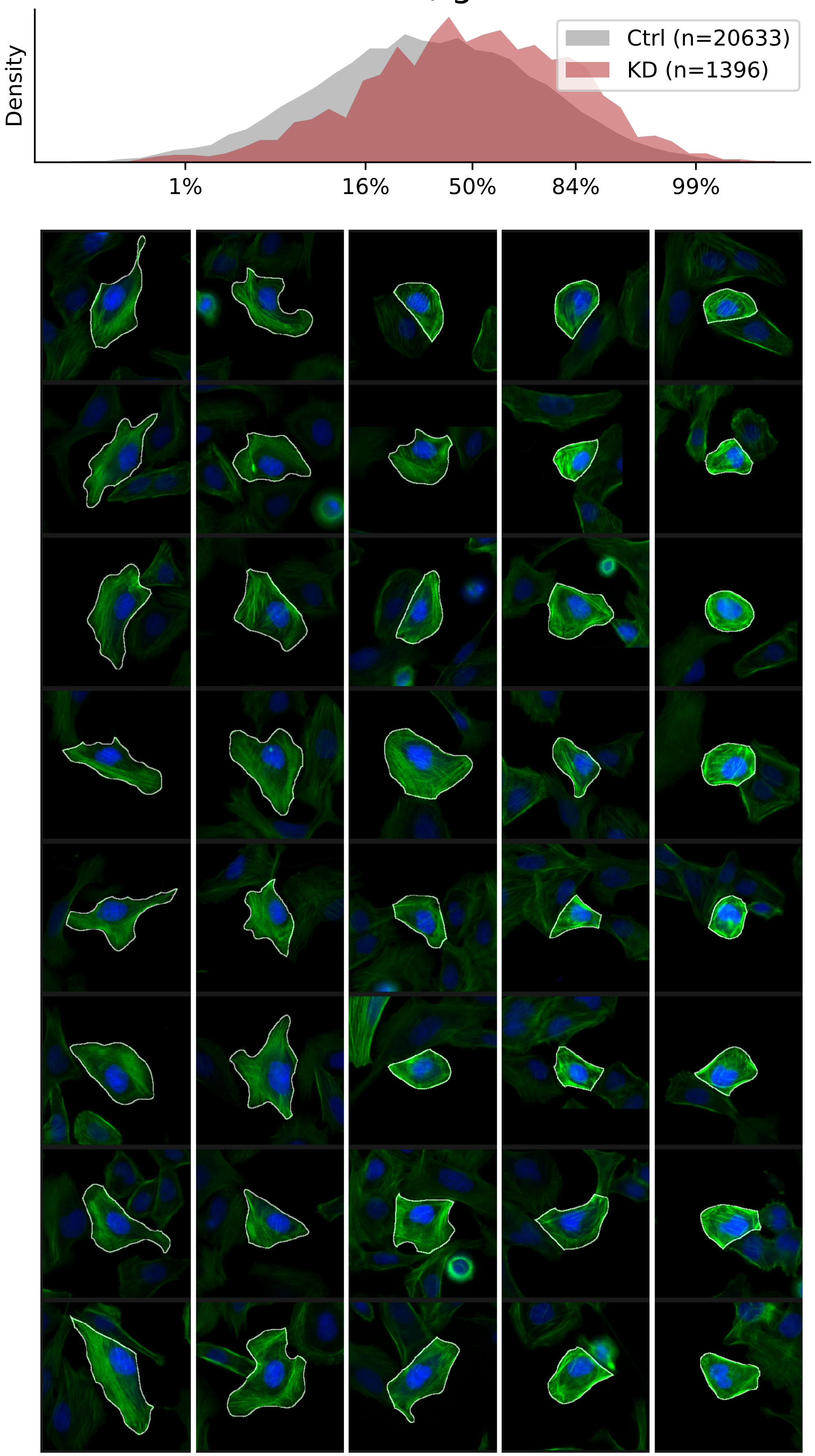

### VIEWS_fPC_ARHGEF7_rep2.pdf

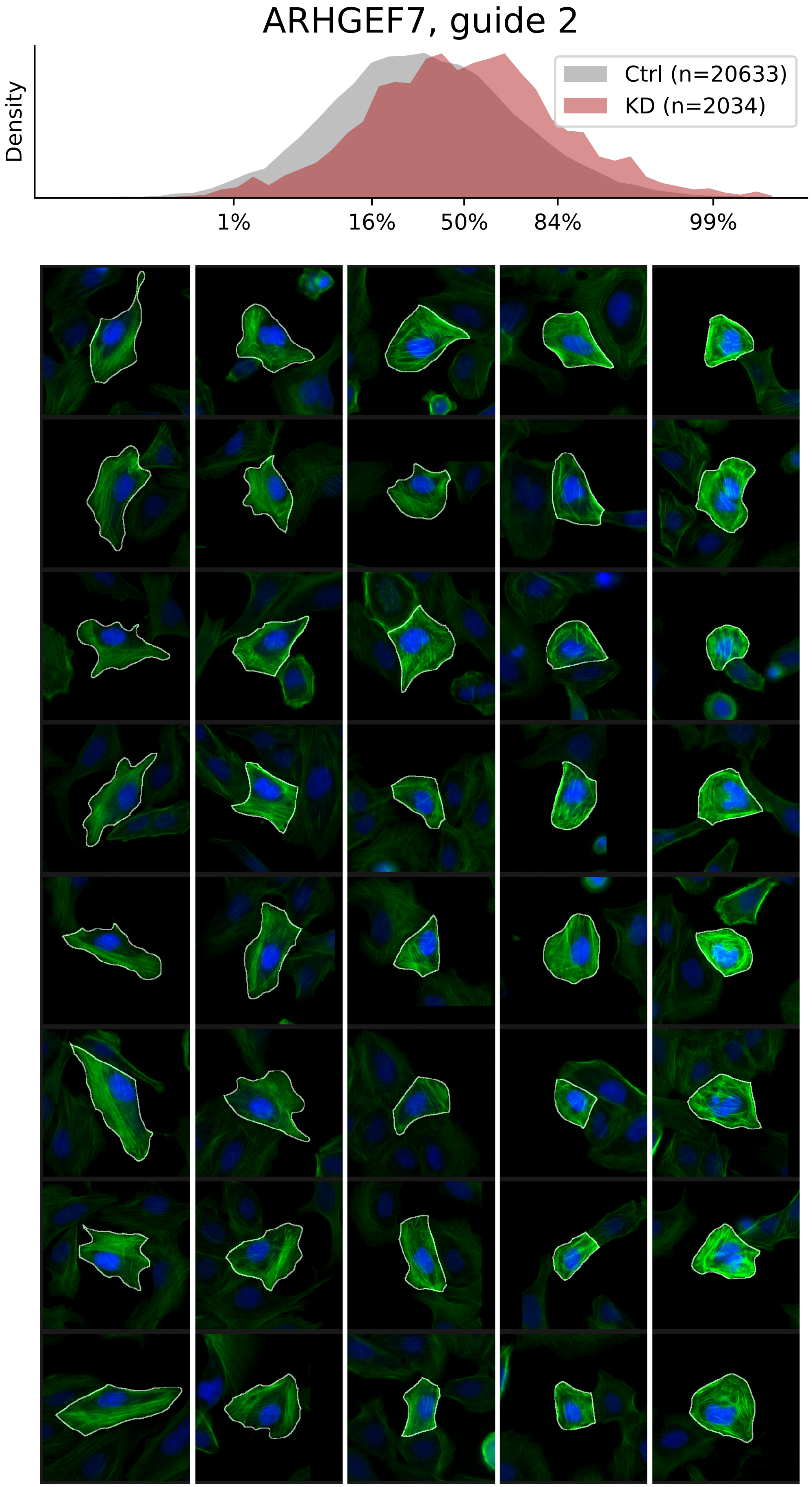

### VIEWS_fPC_ARHGEF7_rep3.pdf

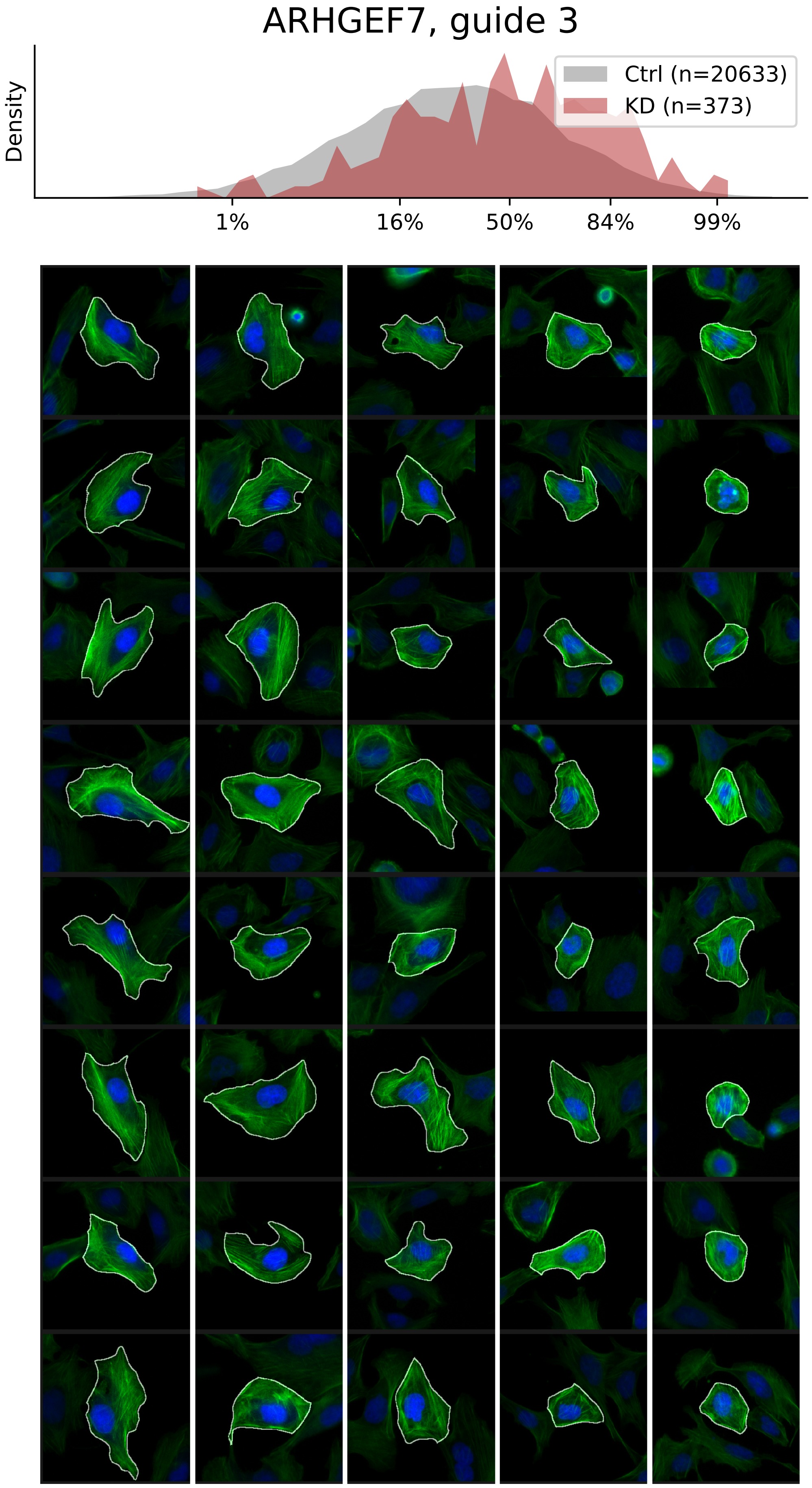

### VIEWS_fPC_ARPC3_rep1.pdf

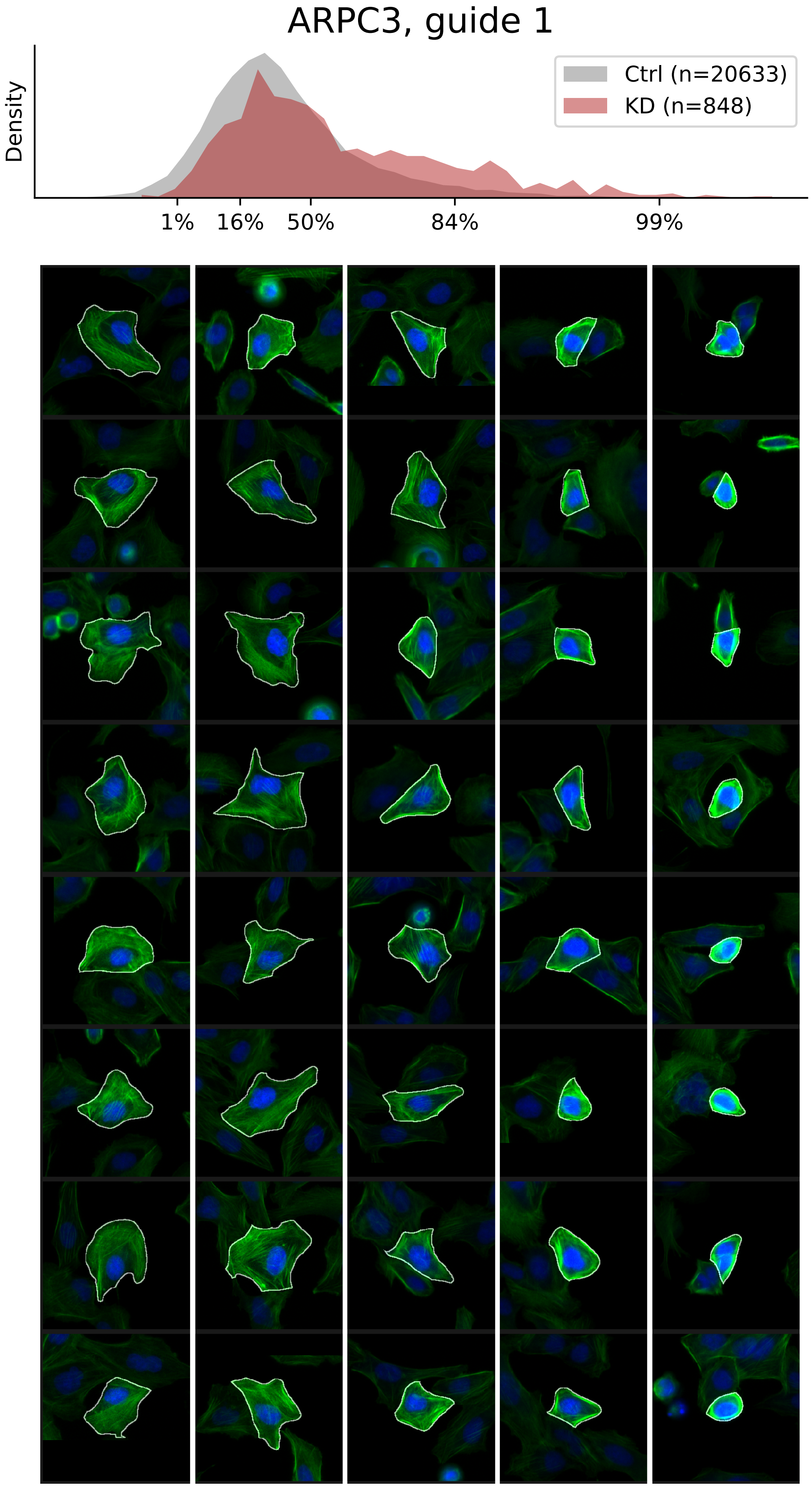

### VIEWS_fPC_ARPC3_rep2.pdf

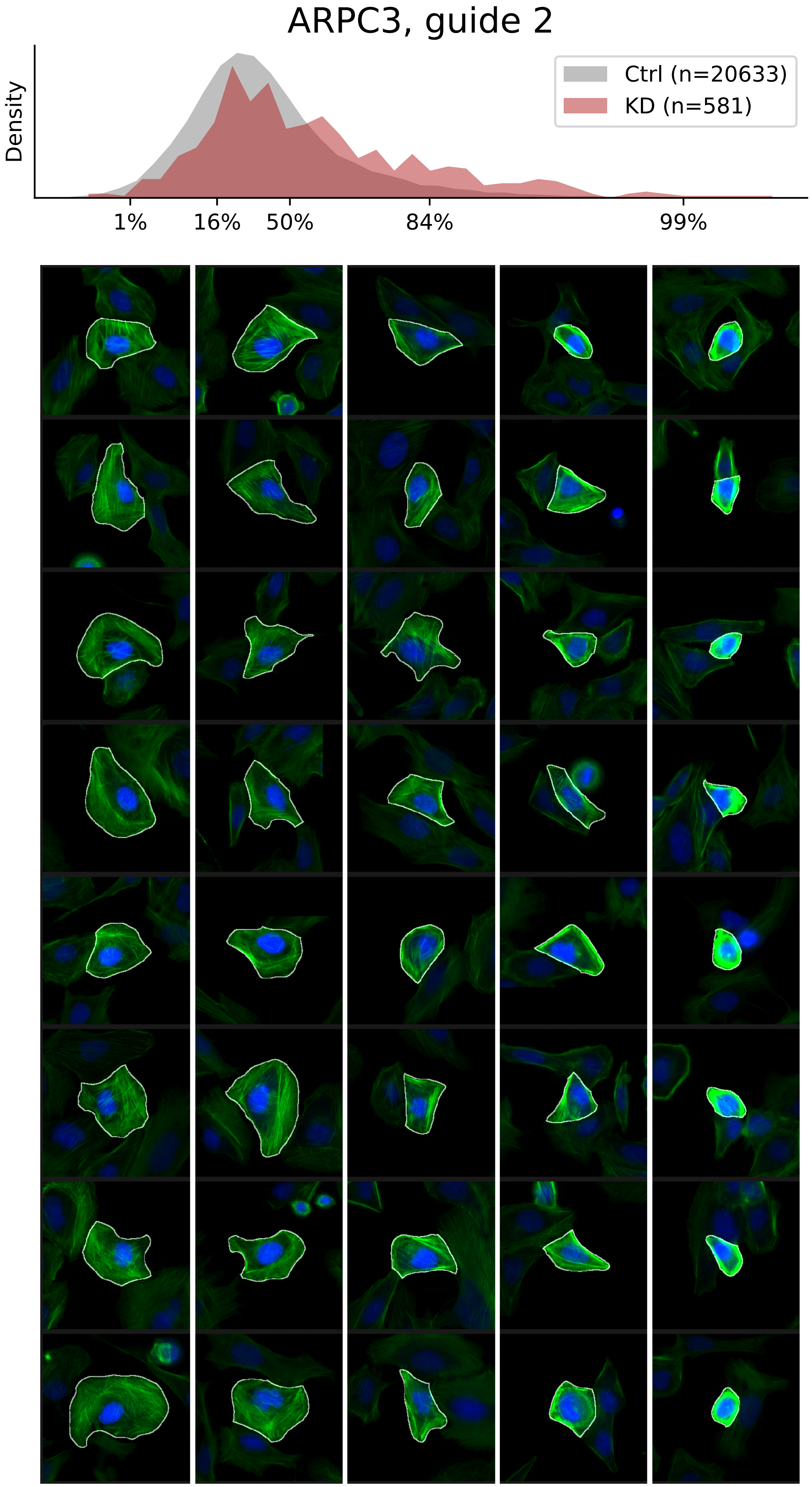

### VIEWS_fPC_BRK1_rep1.pdf

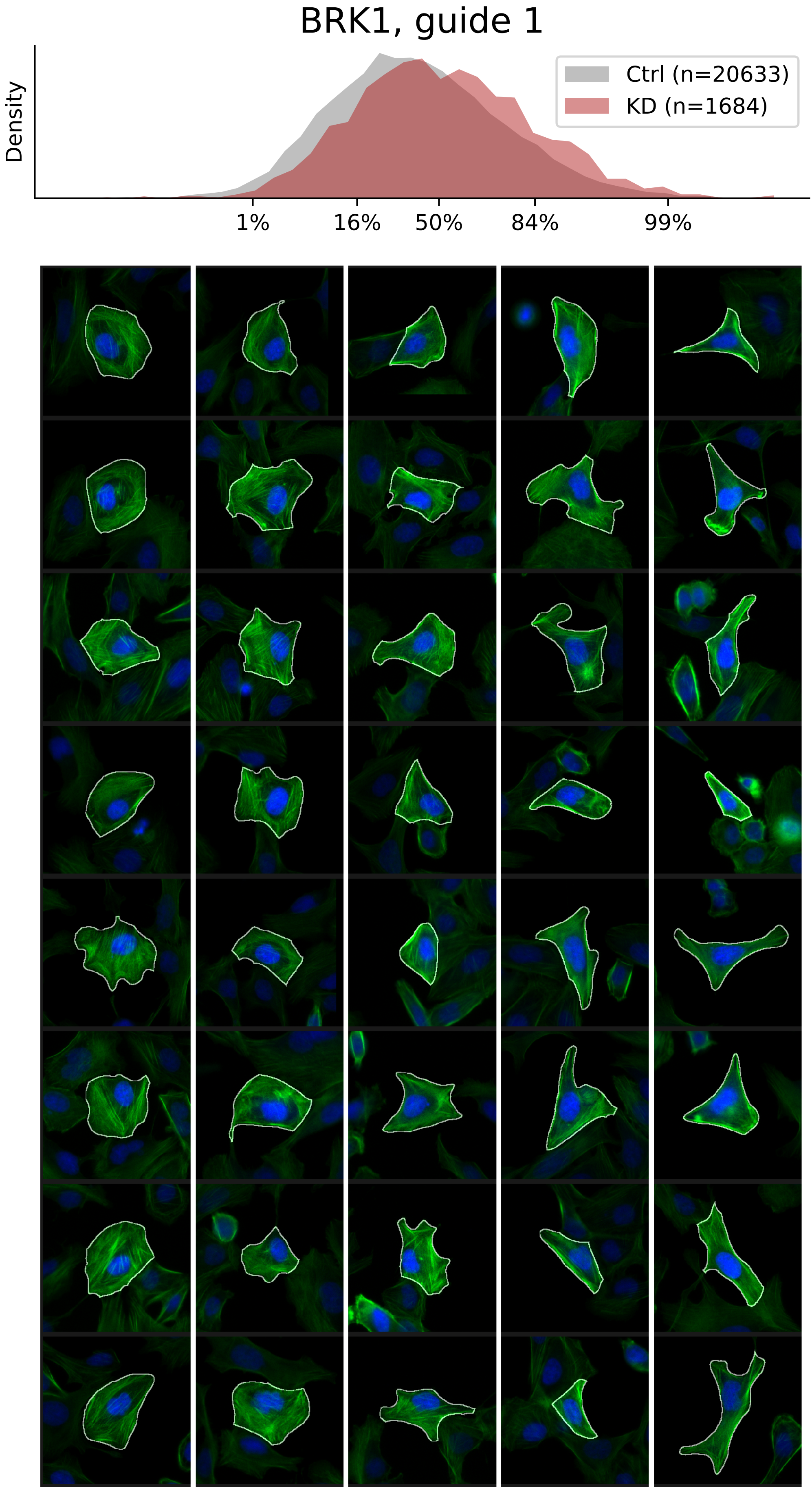

### VIEWS_fPC_BRK1_rep2.pdf

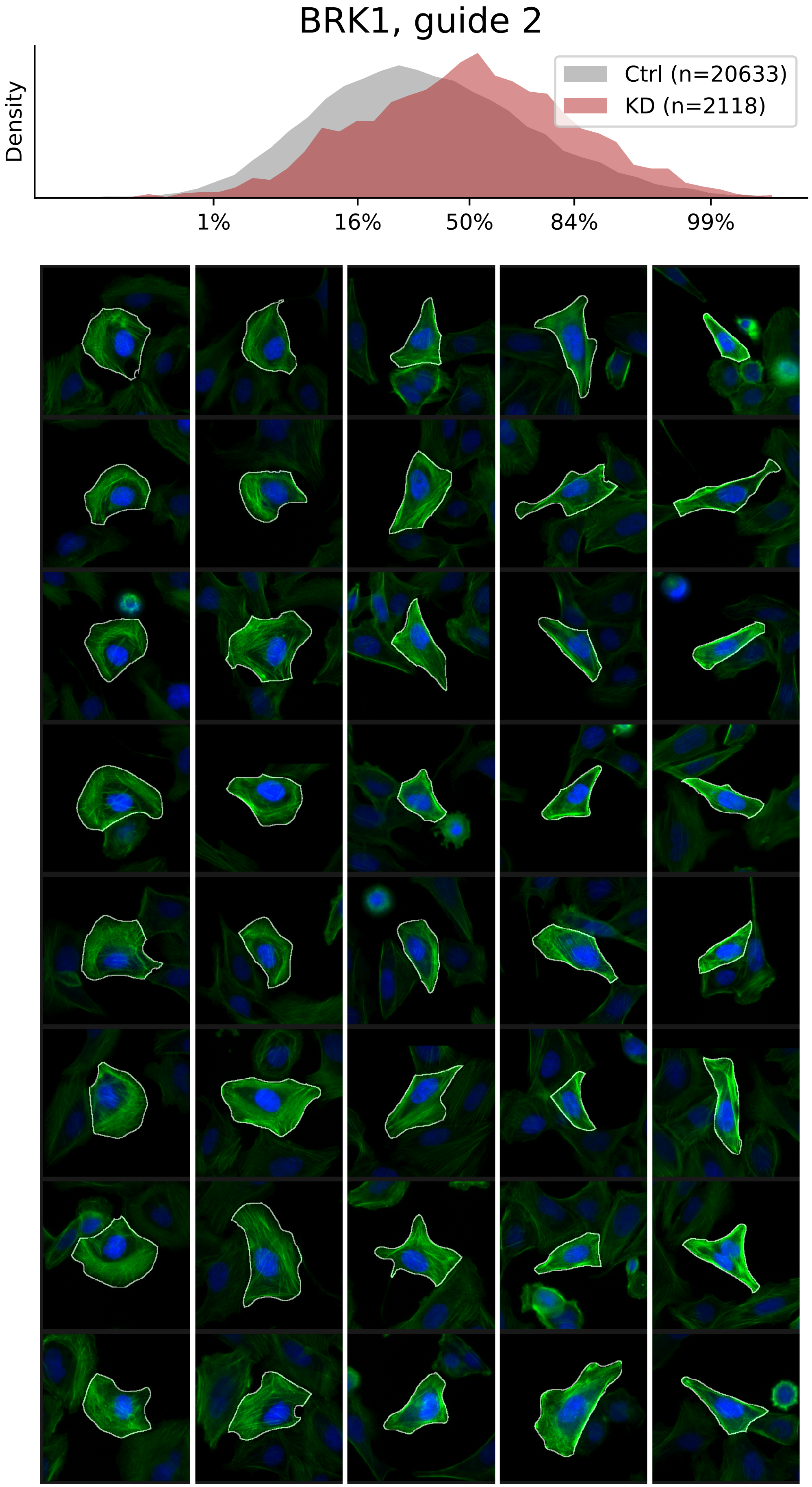

### VIEWS_fPC_BRK1_rep3.pdf

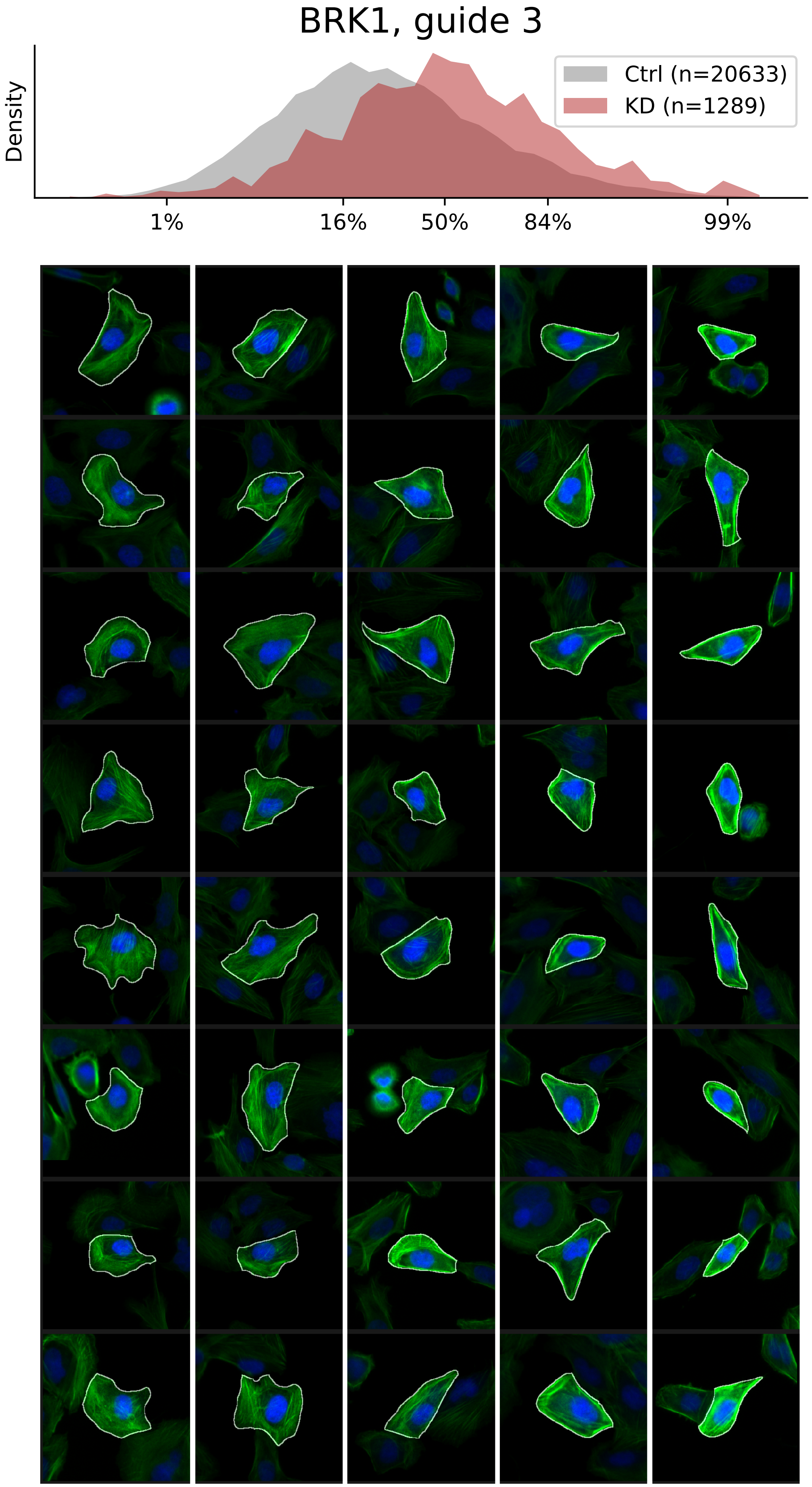

### VIEWS_fPC_CAP1_rep1.pdf

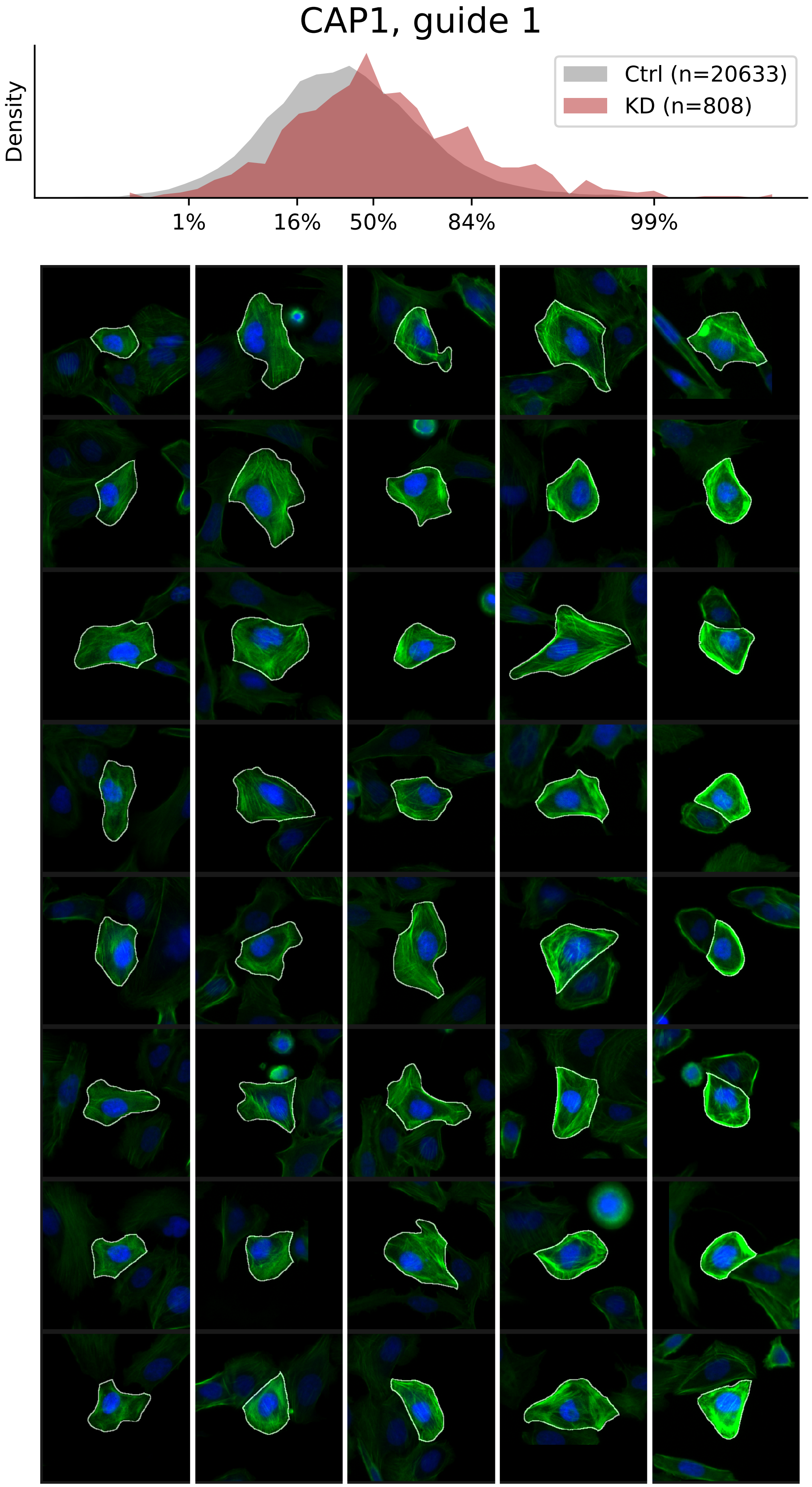

### VIEWS_fPC_CAP1_rep2.pdf

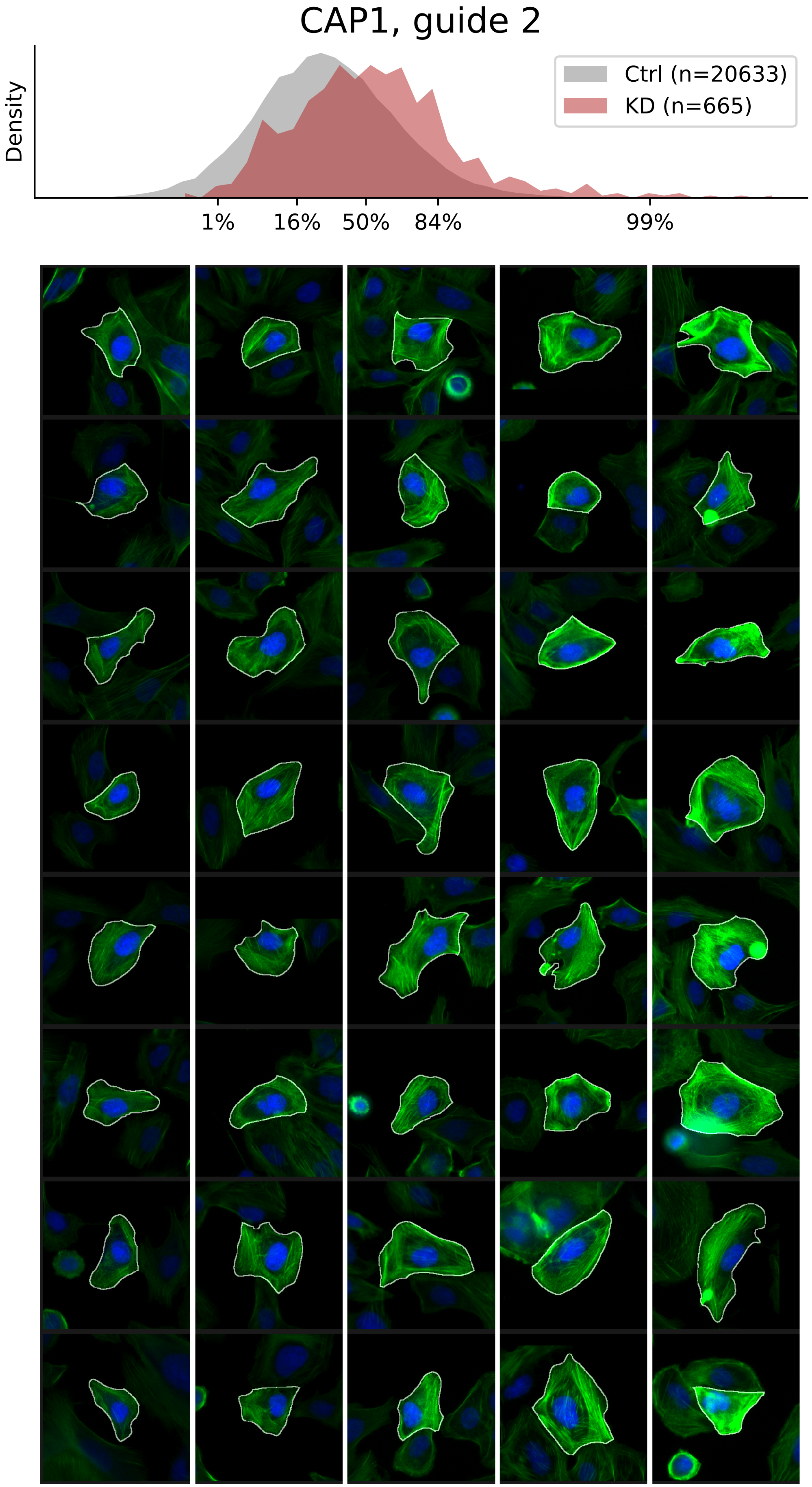

### VIEWS_fPC_CAPZA1_rep2.pdf

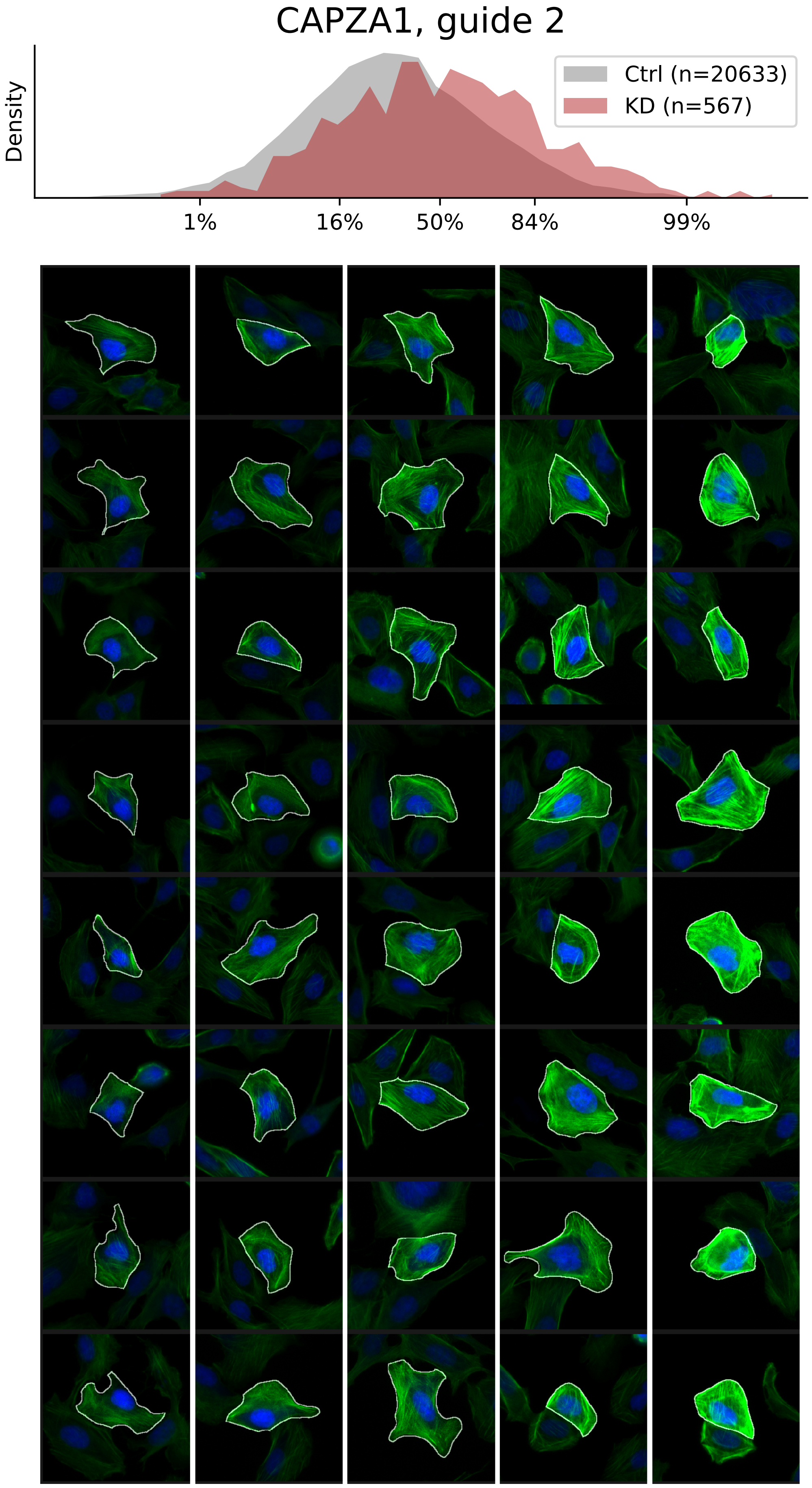

### VIEWS_fPC_CAPZA1_rep3.pdf

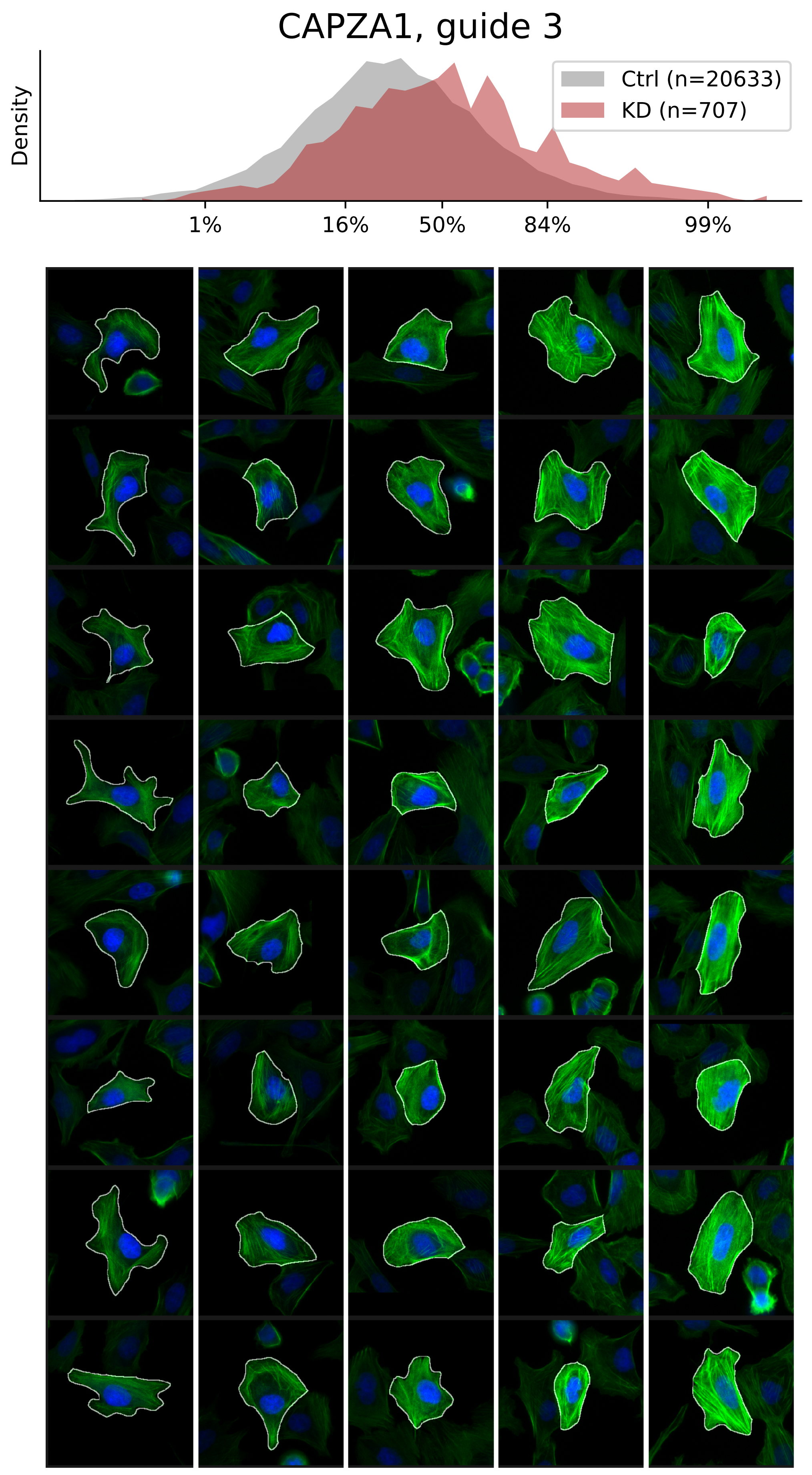

### VIEWS_fPC_CAPZB_rep2.pdf

#### CAPZB, guide 2

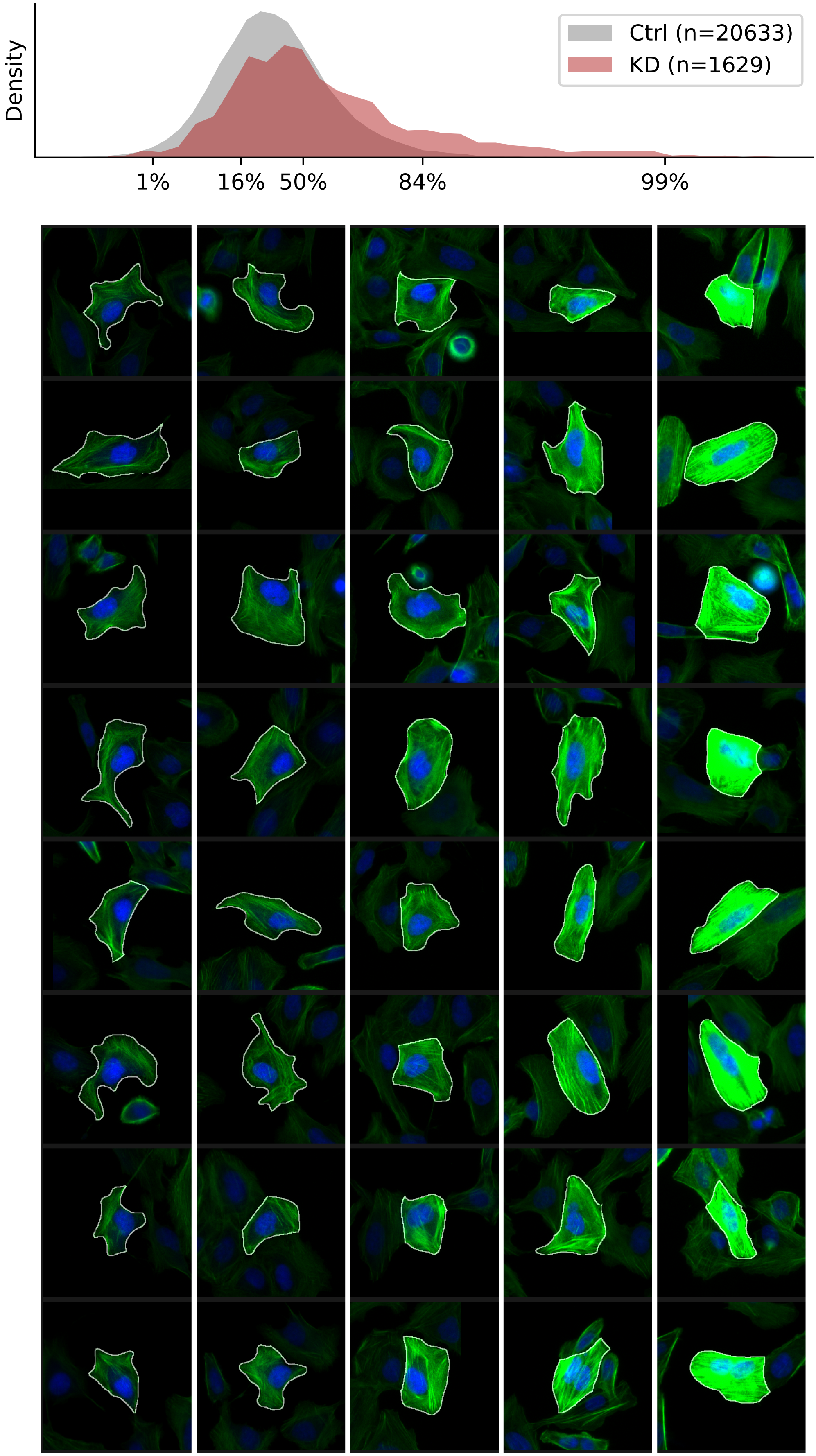

### VIEWS_fPC_CAPZB_rep3.pdf

#### CAPZB, guide 3

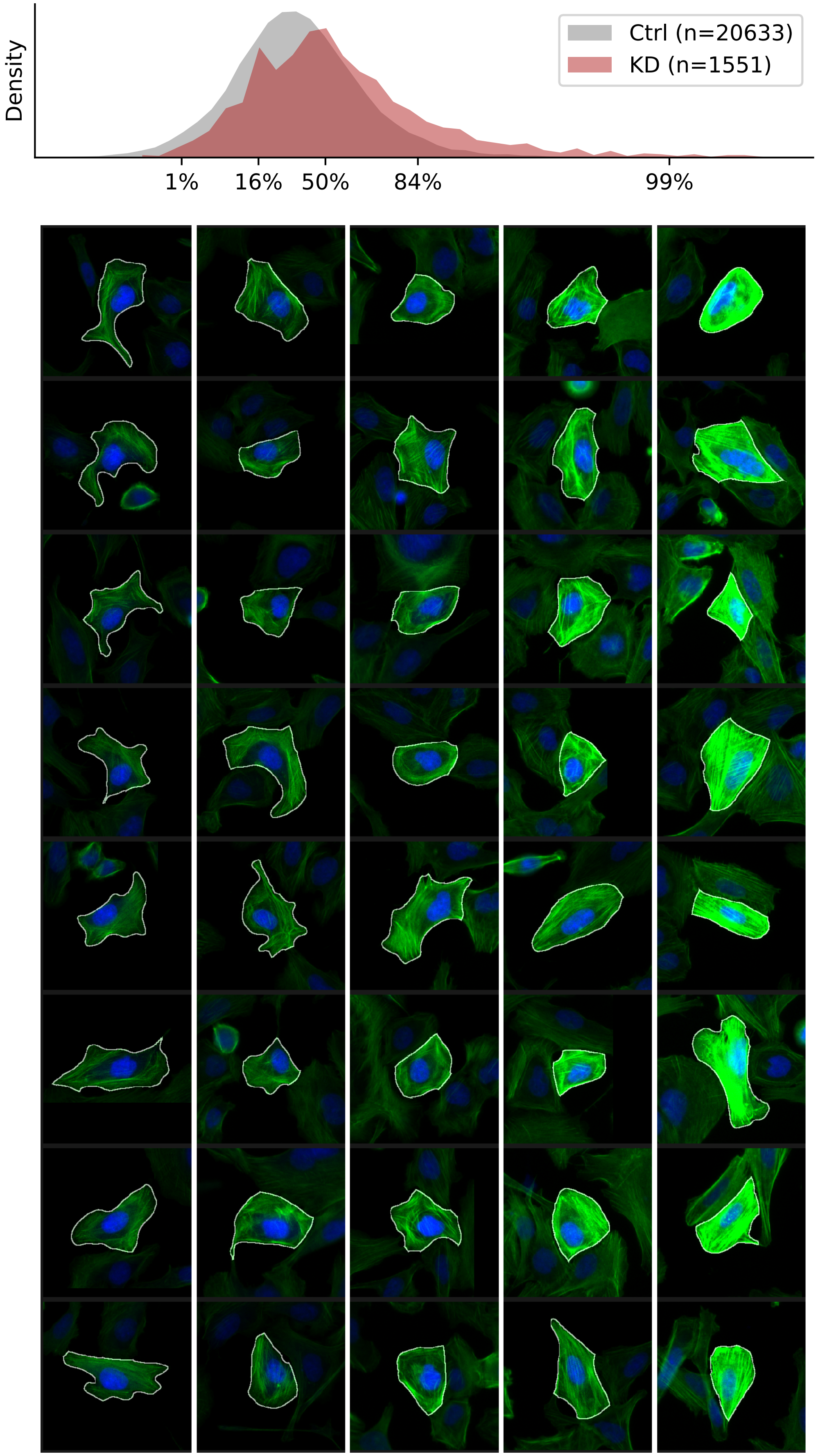

### VIEWS_fPC_CTTNBP2NL_rep2.pdf

#### CTTNBP2NL, guide 2

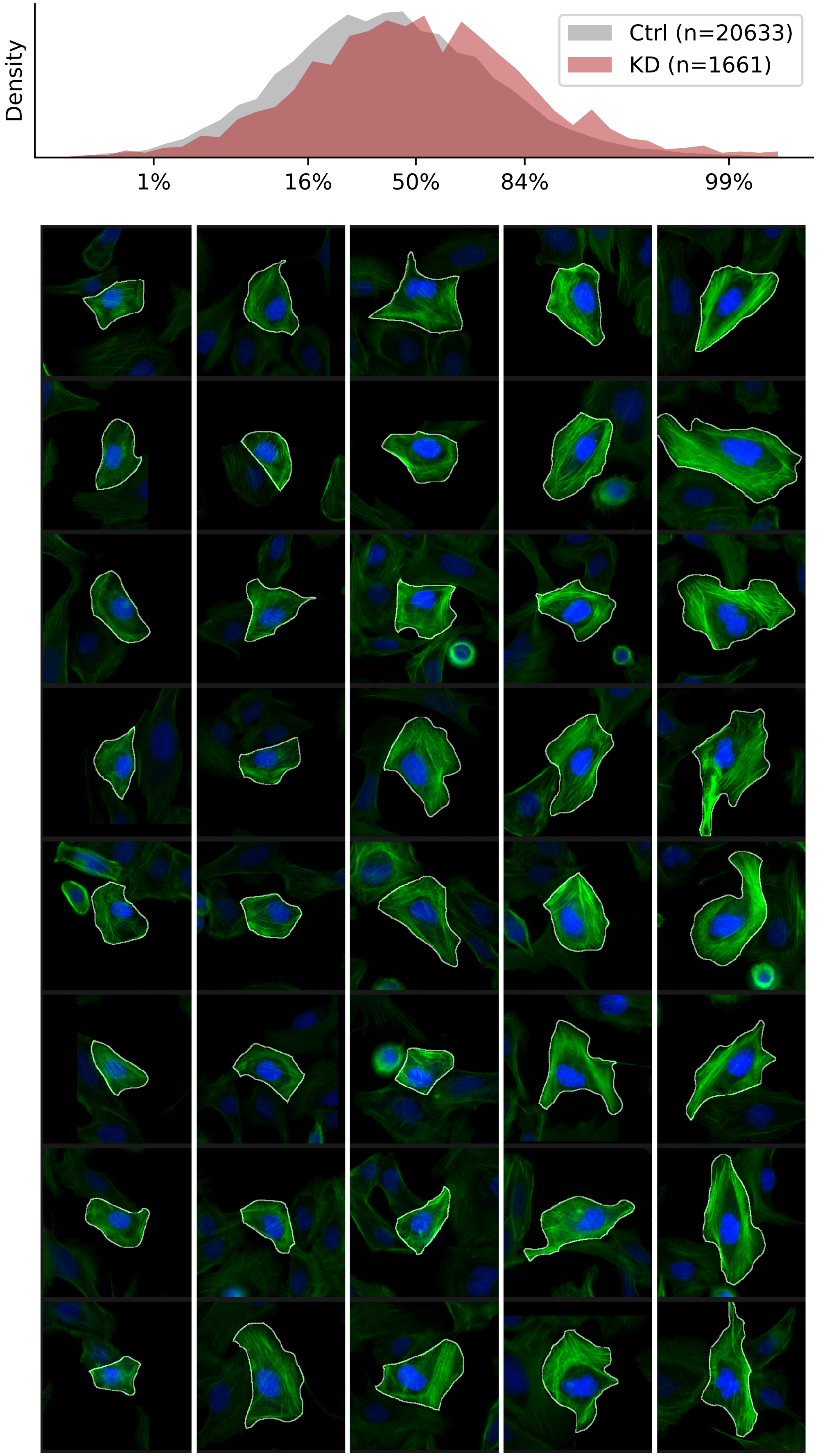

### VIEWS_fPC_CTTNBP2NL_rep3.pdf

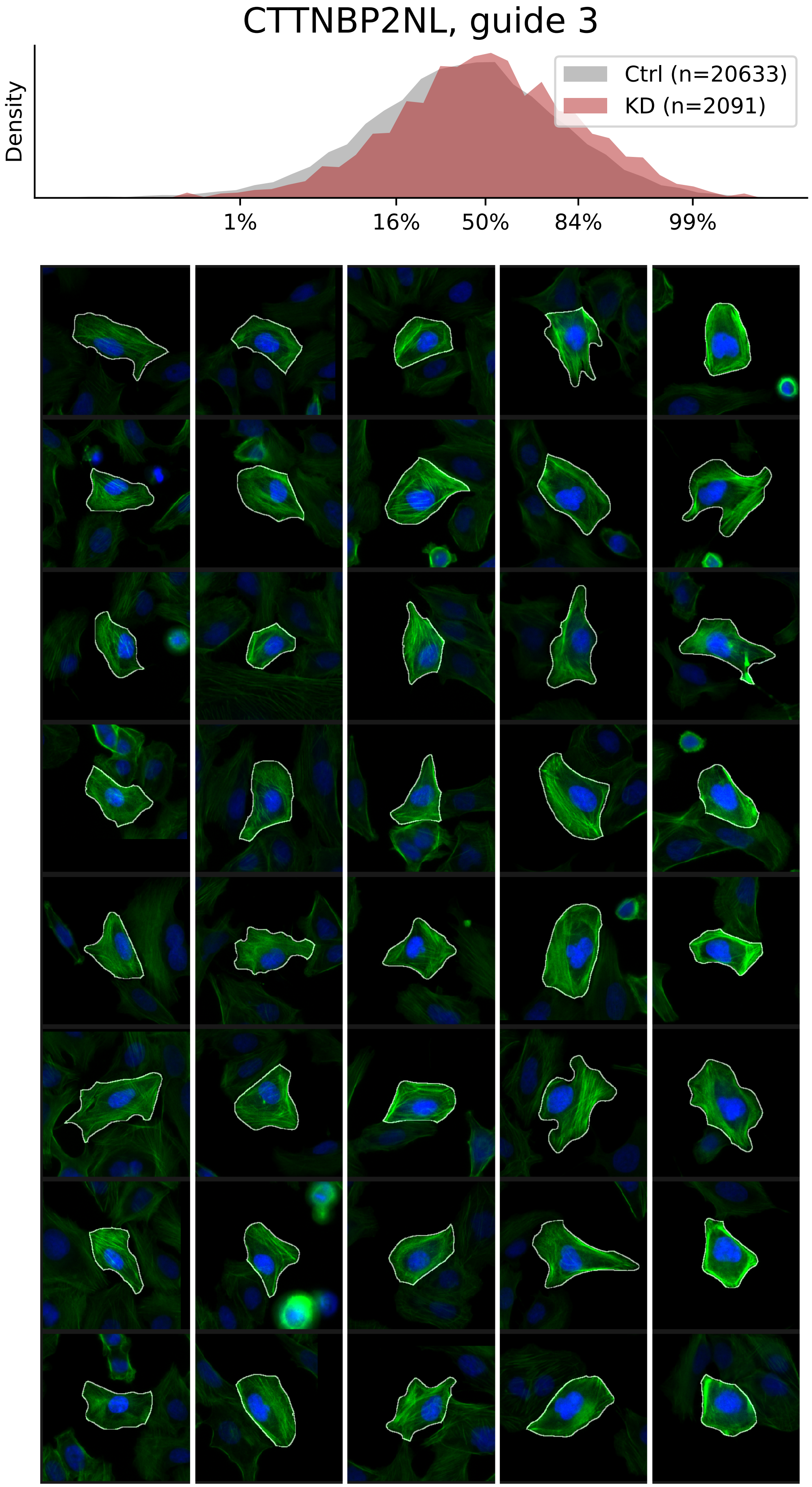

### VIEWS_fPC_FLII_rep1.pdf

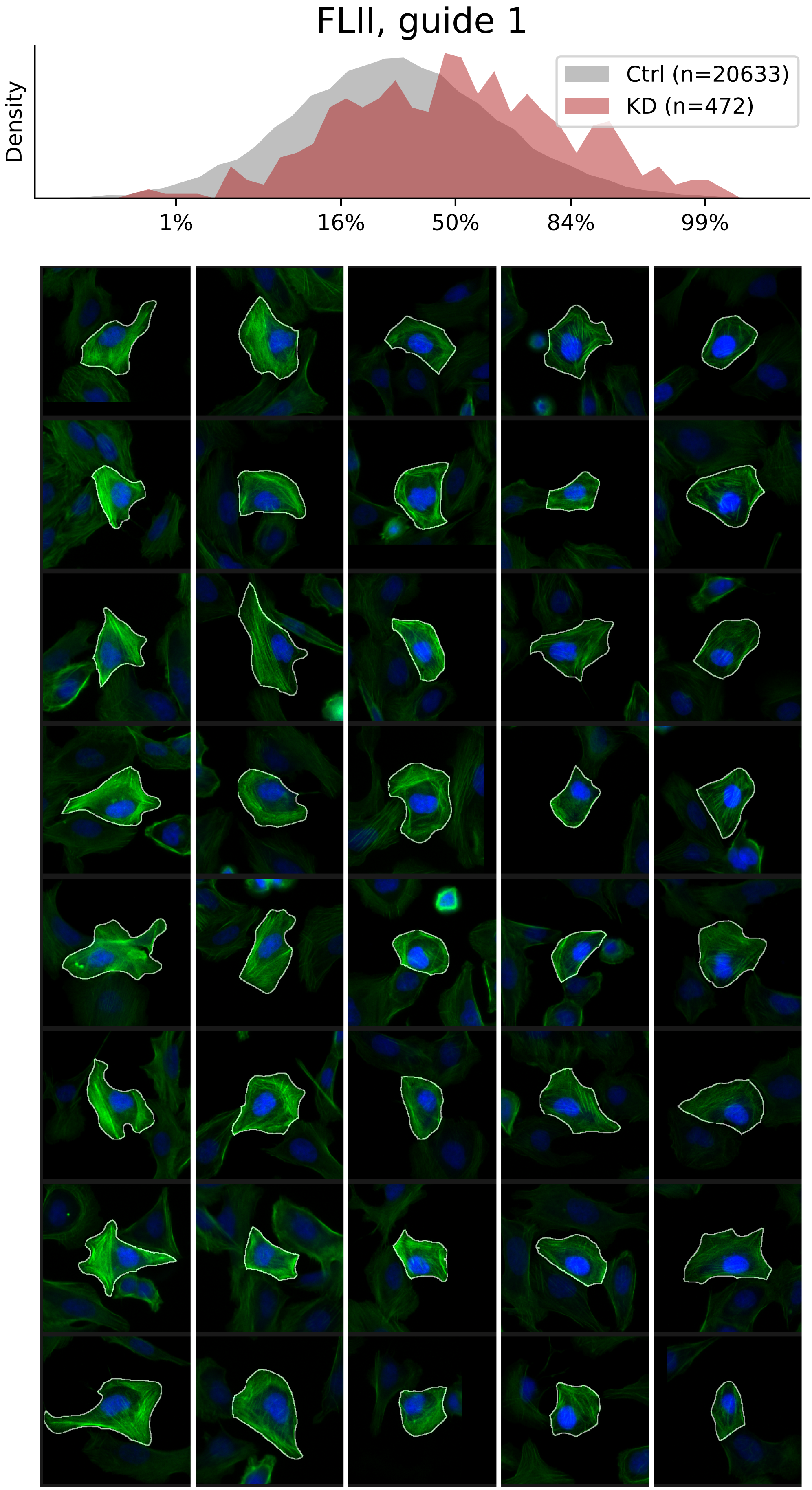

### VIEWS_fPC_FLII_rep2.pdf

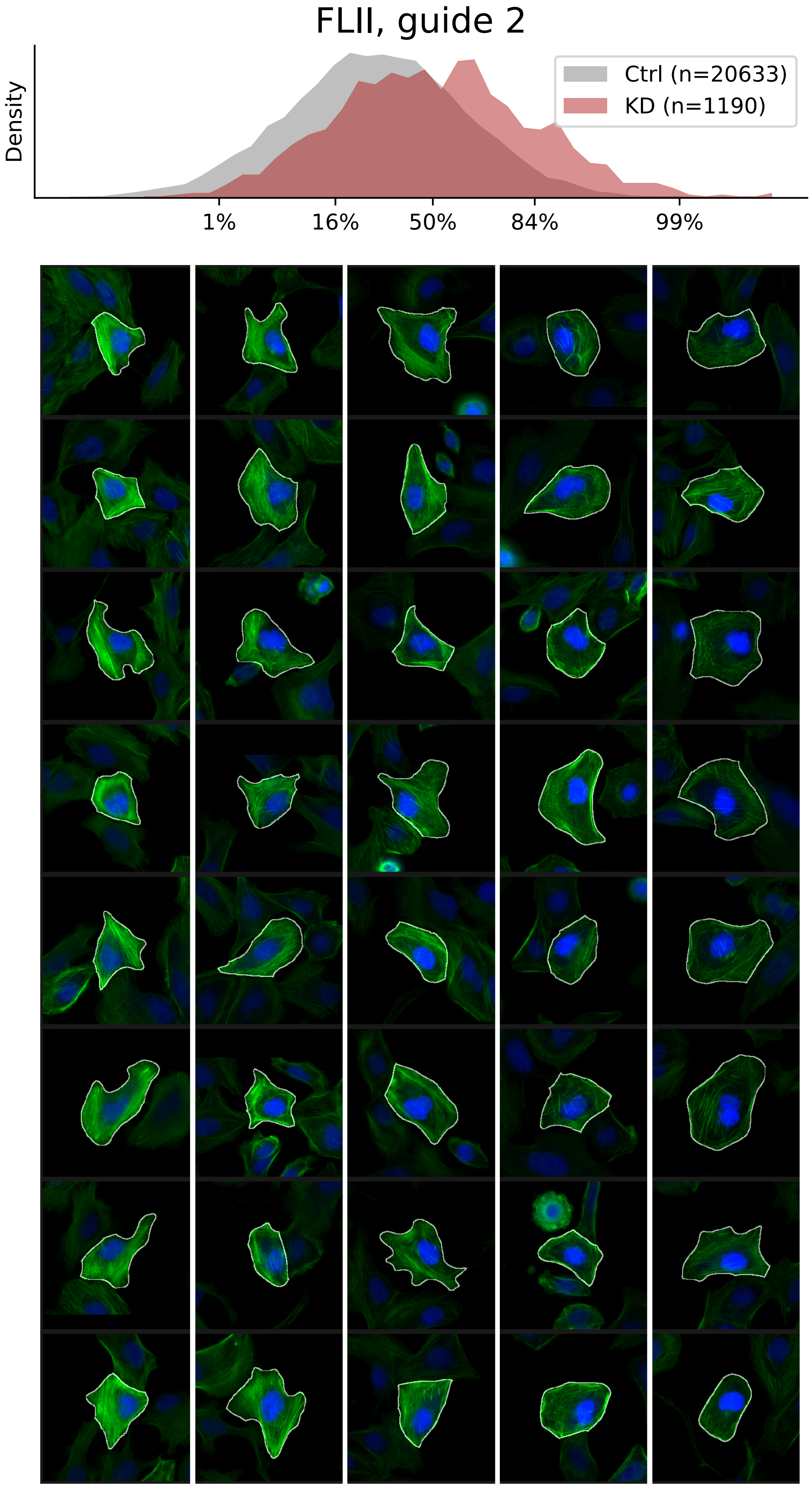

### VIEWS_fPC_FLII_rep3.pdf

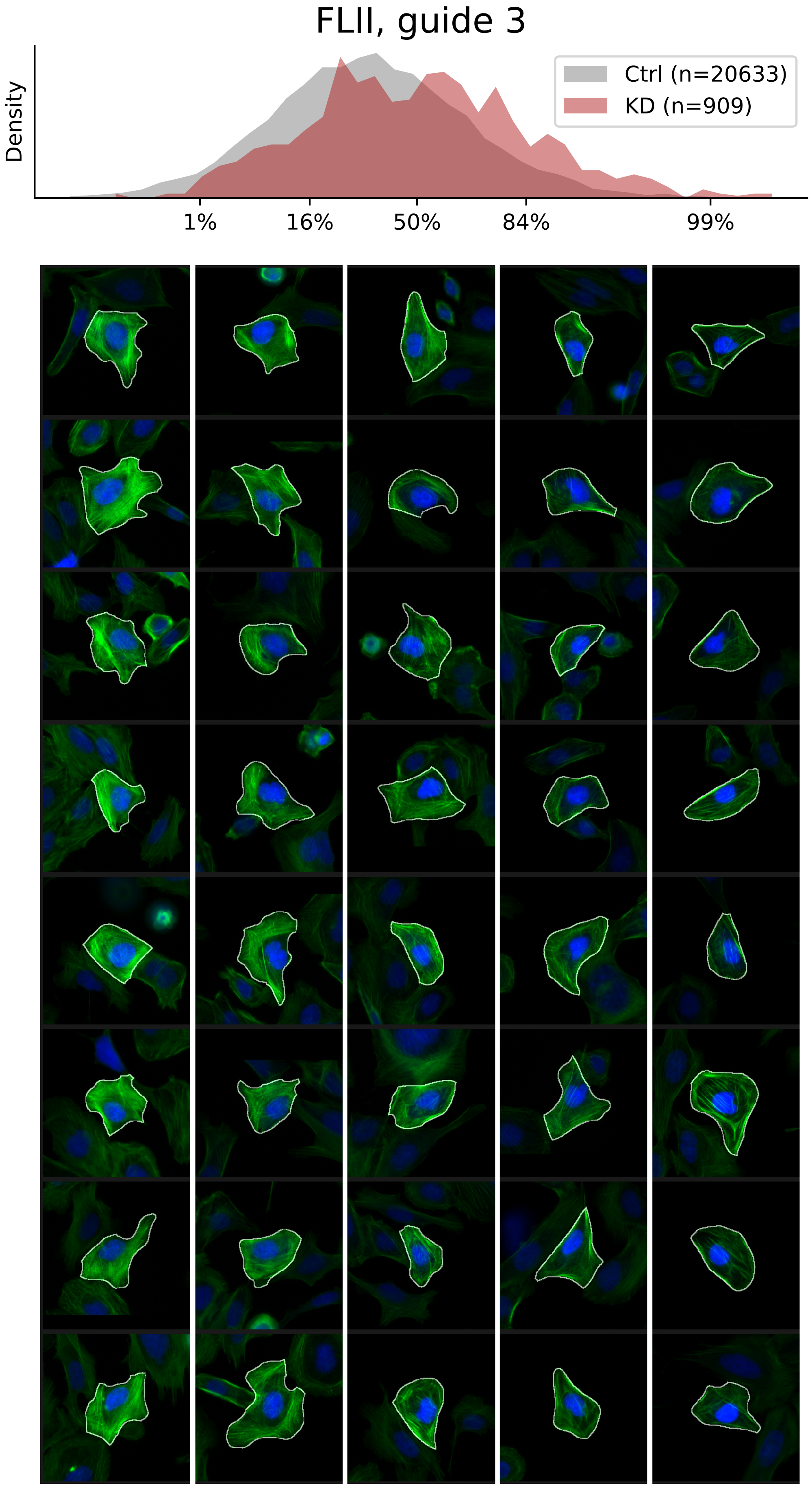

### VIEWS_fPC_GNA12_rep2.pdf

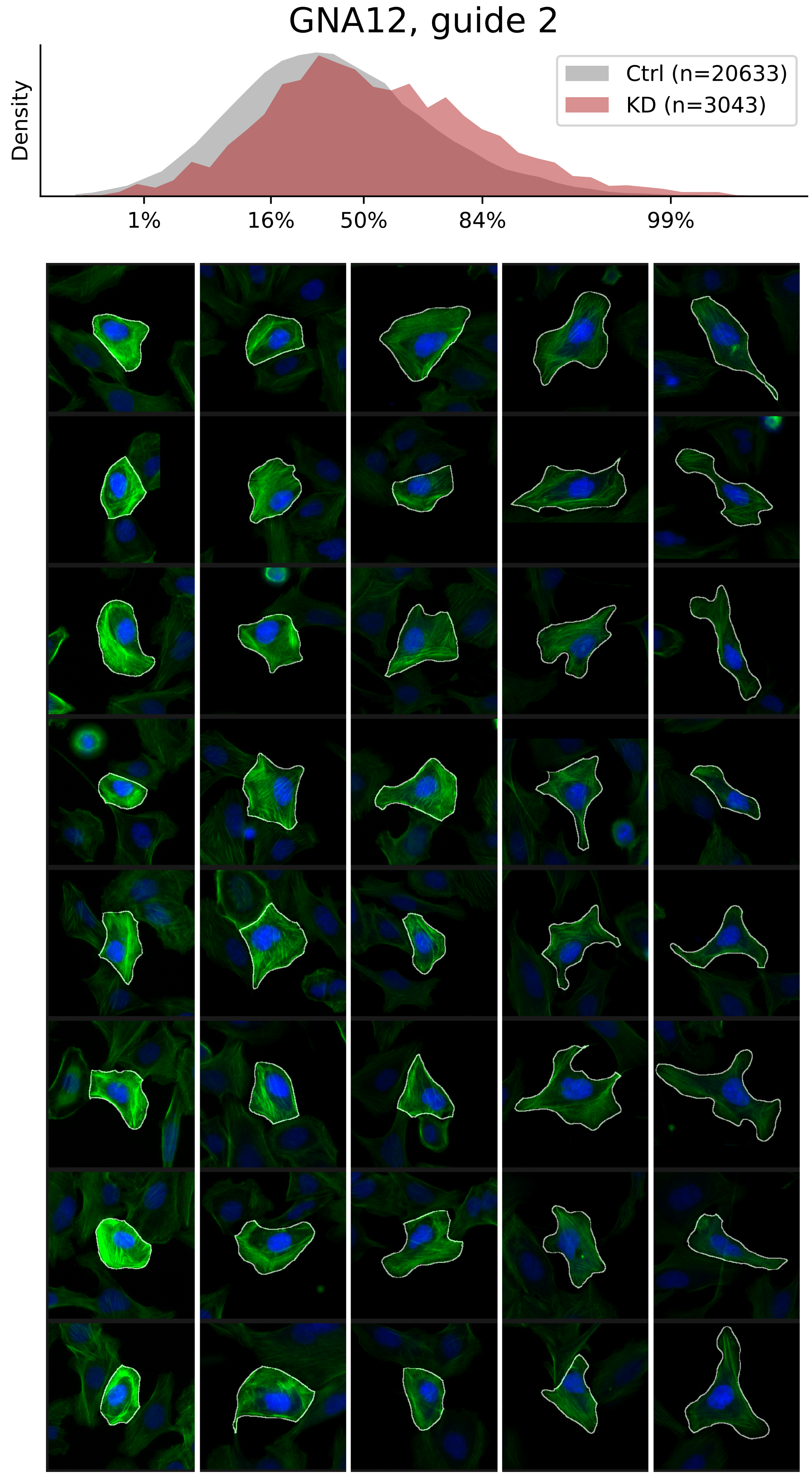

### VIEWS_fPC_ILK_rep2.pdf

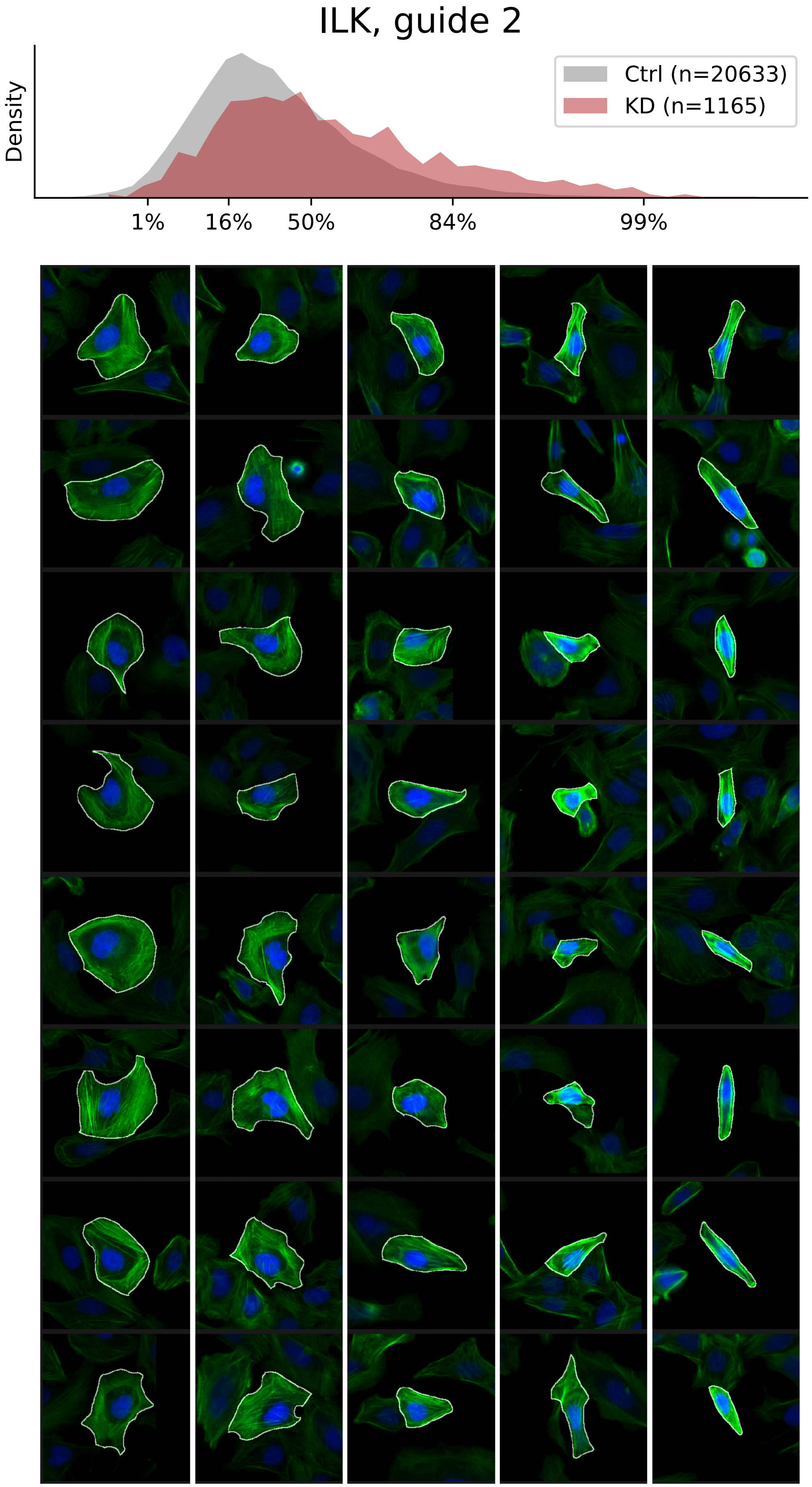

### VIEWS_fPC_ILK_rep3.pdf

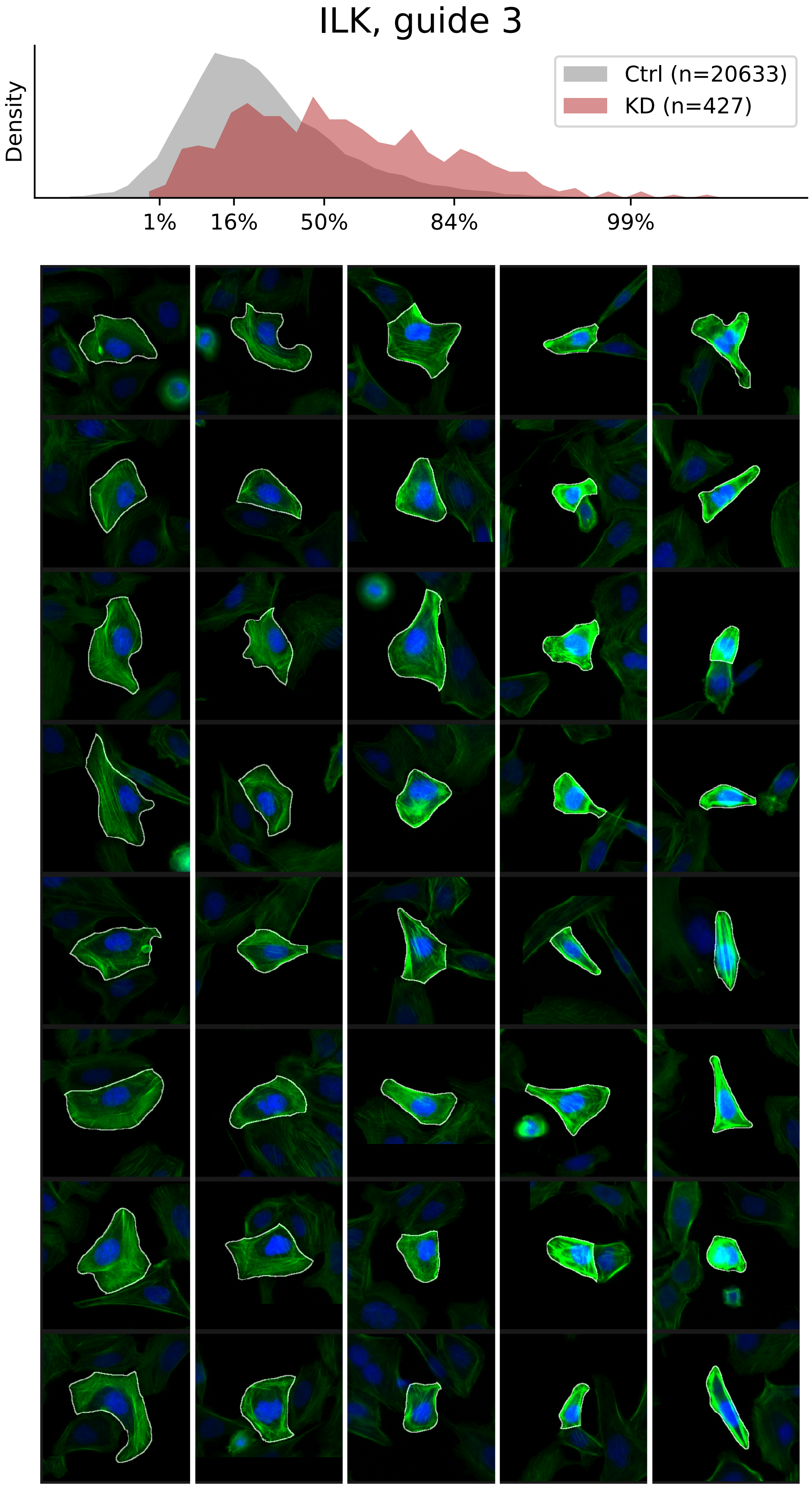

### VIEWS_fPC_LMNA_rep1.pdf

##### LMNA, guide 1

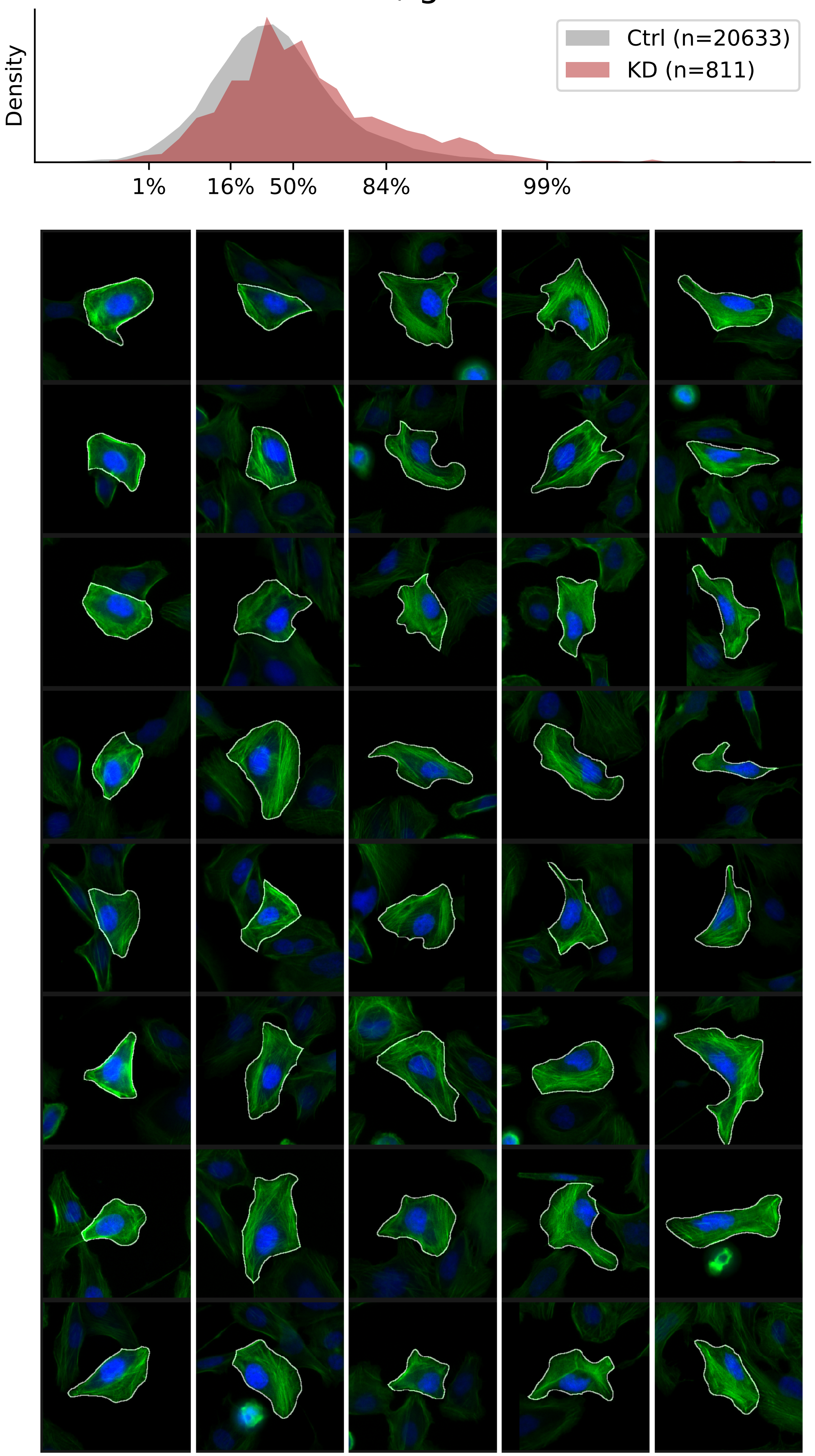

### VIEWS_fPC_LMNA_rep2.pdf

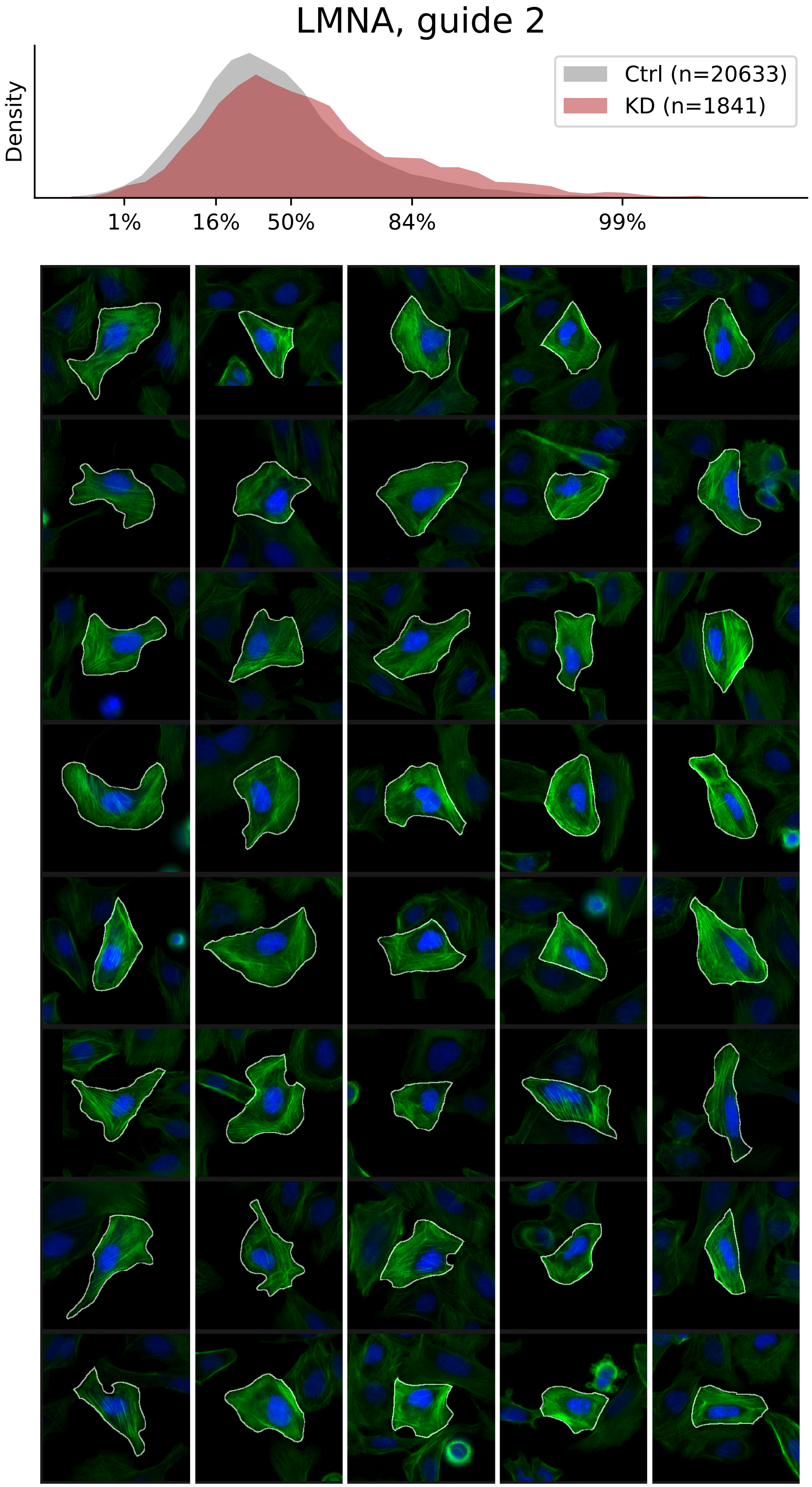

### VIEWS_fPC_LMNA_rep3.pdf

#### LMNA, guide 3

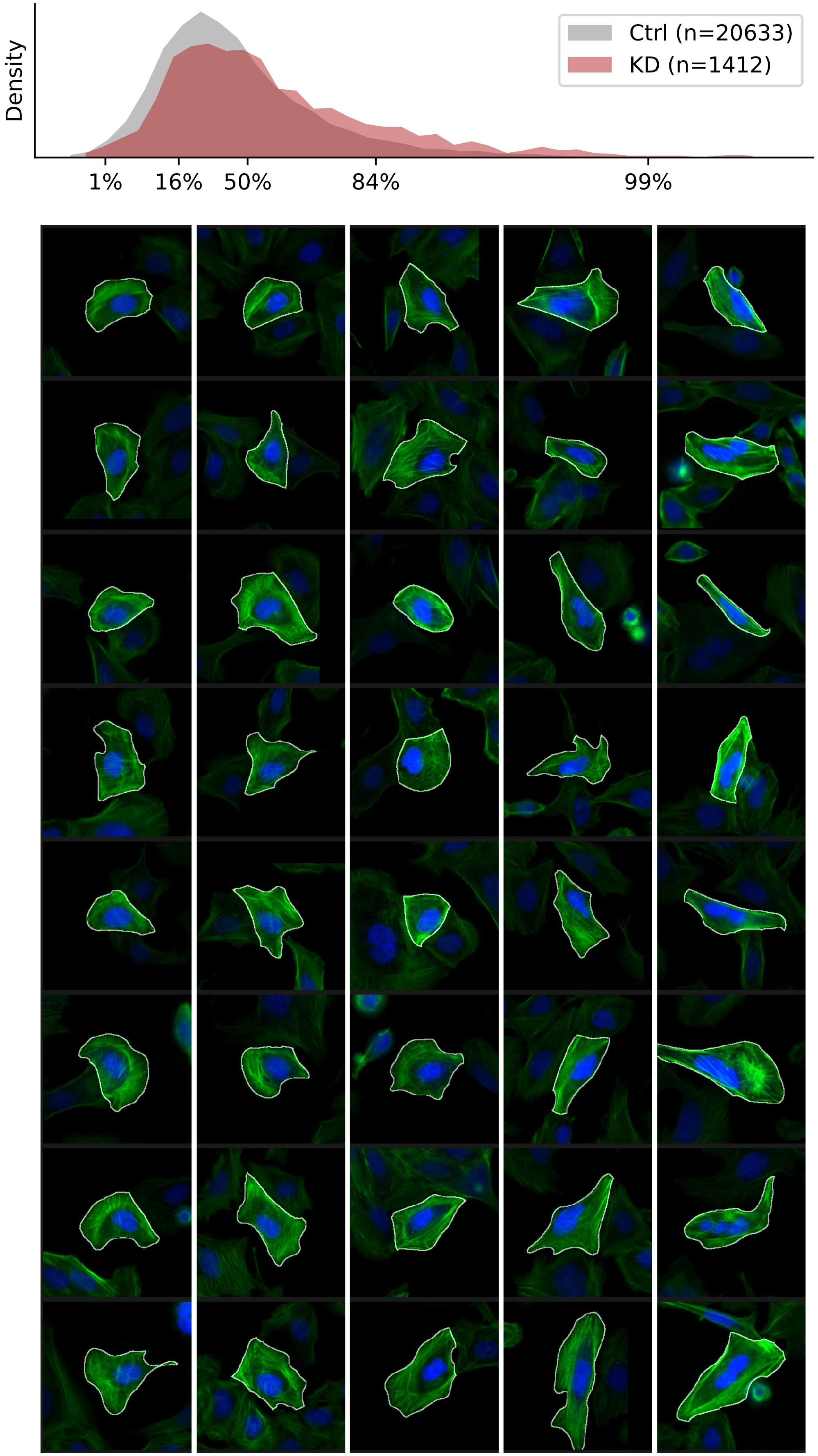

### VIEWS_fPC_MAEA_rep1.pdf

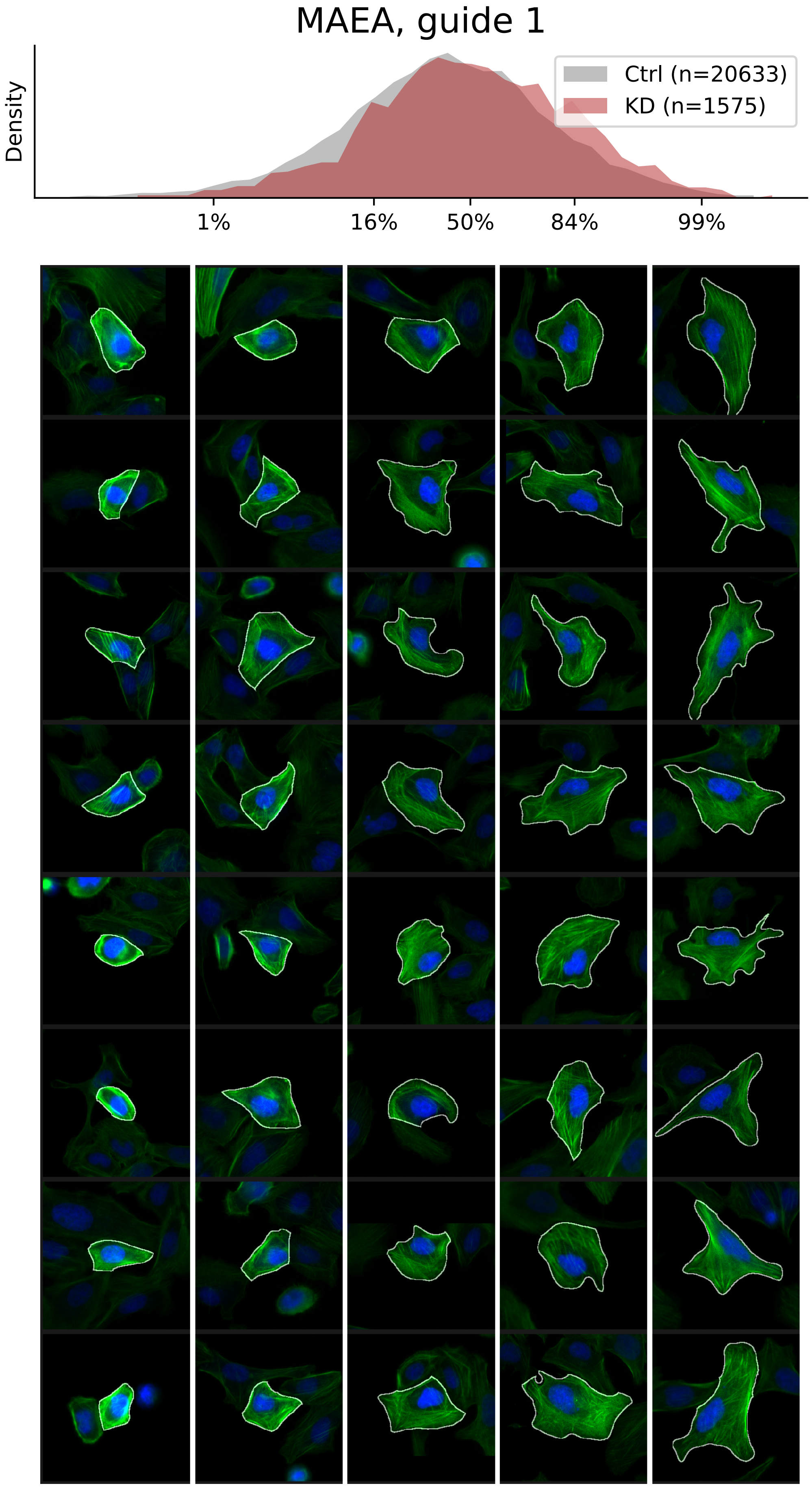

### VIEWS_fPC_MAEA_rep2.pdf

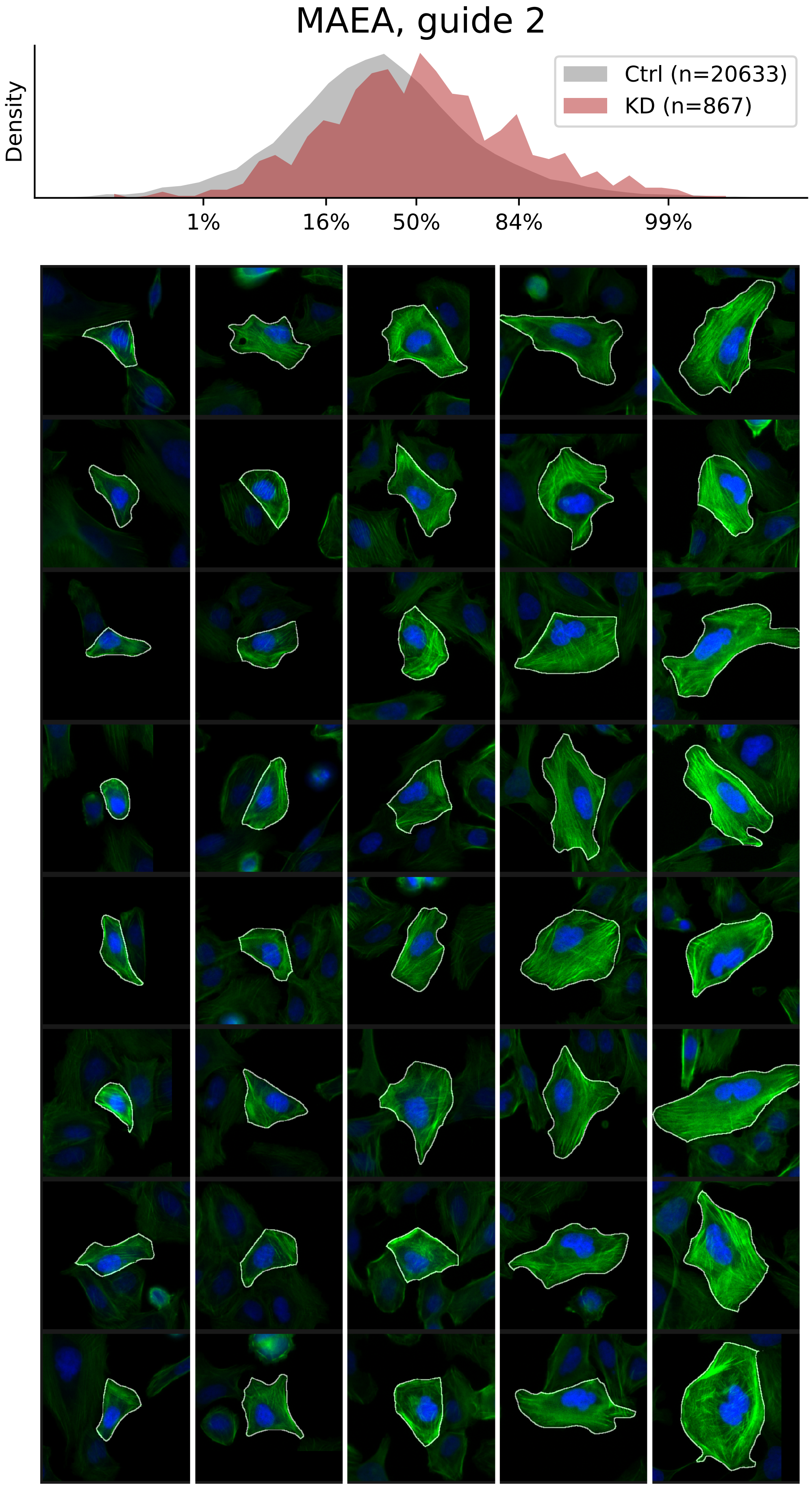

### VIEWS_fPC_MYO1B_rep1.pdf

#### MYO1B, guide 1

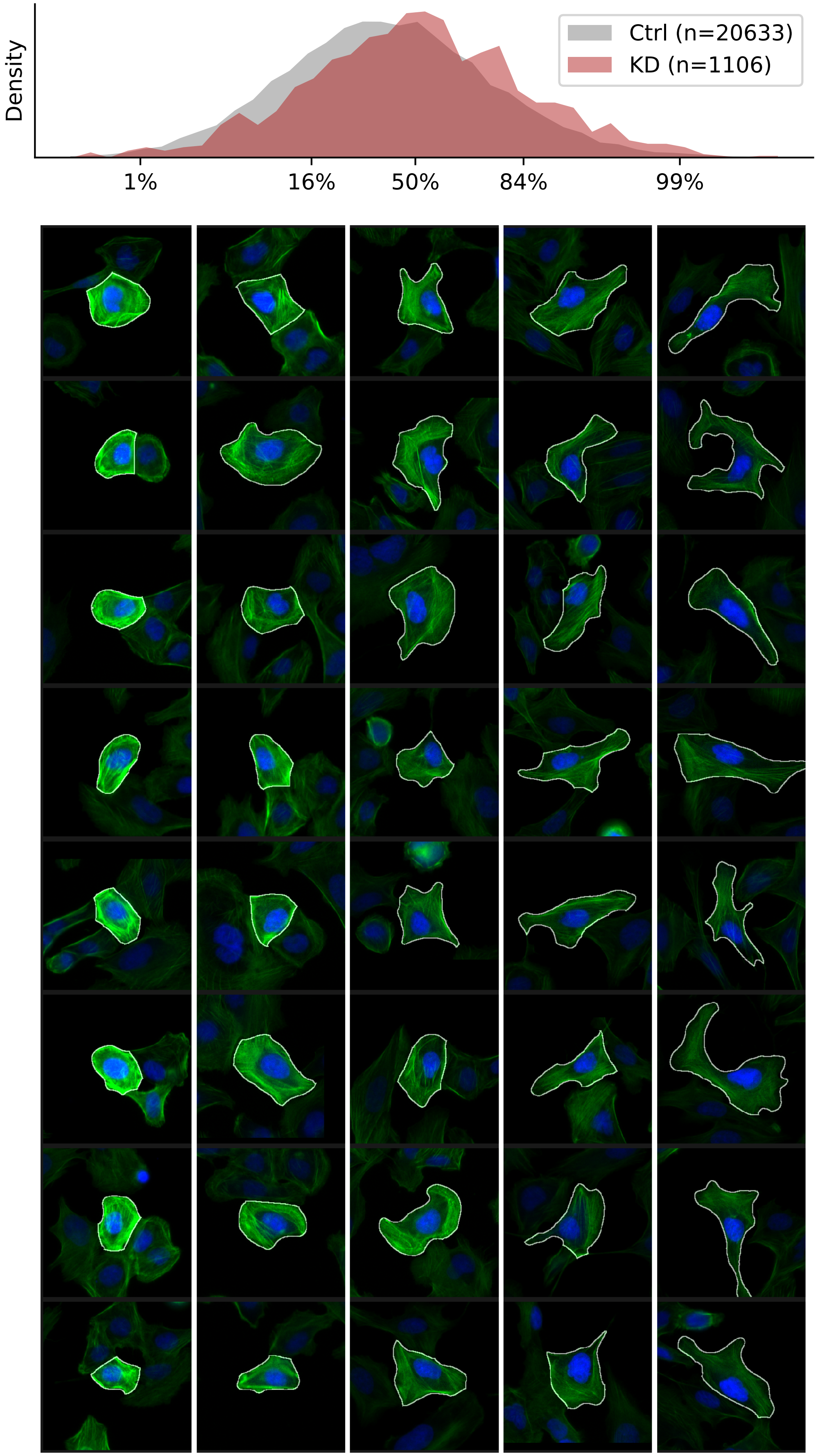

### VIEWS_fPC_MYO1B_rep2.pdf

#### MYO1B, guide 2

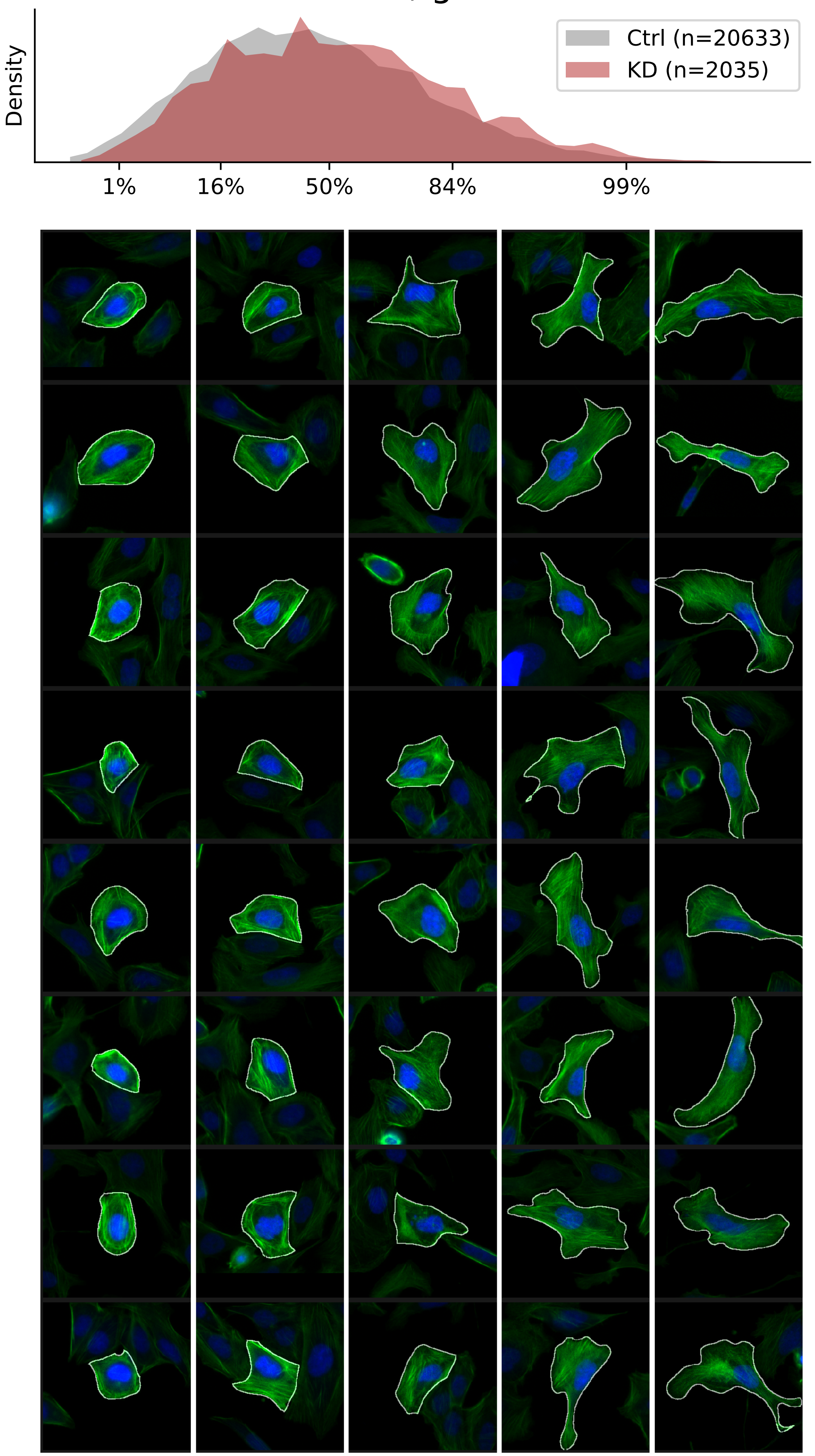

### VIEWS_fPC_PXN_rep1.pdf

#### PXN, guide 1

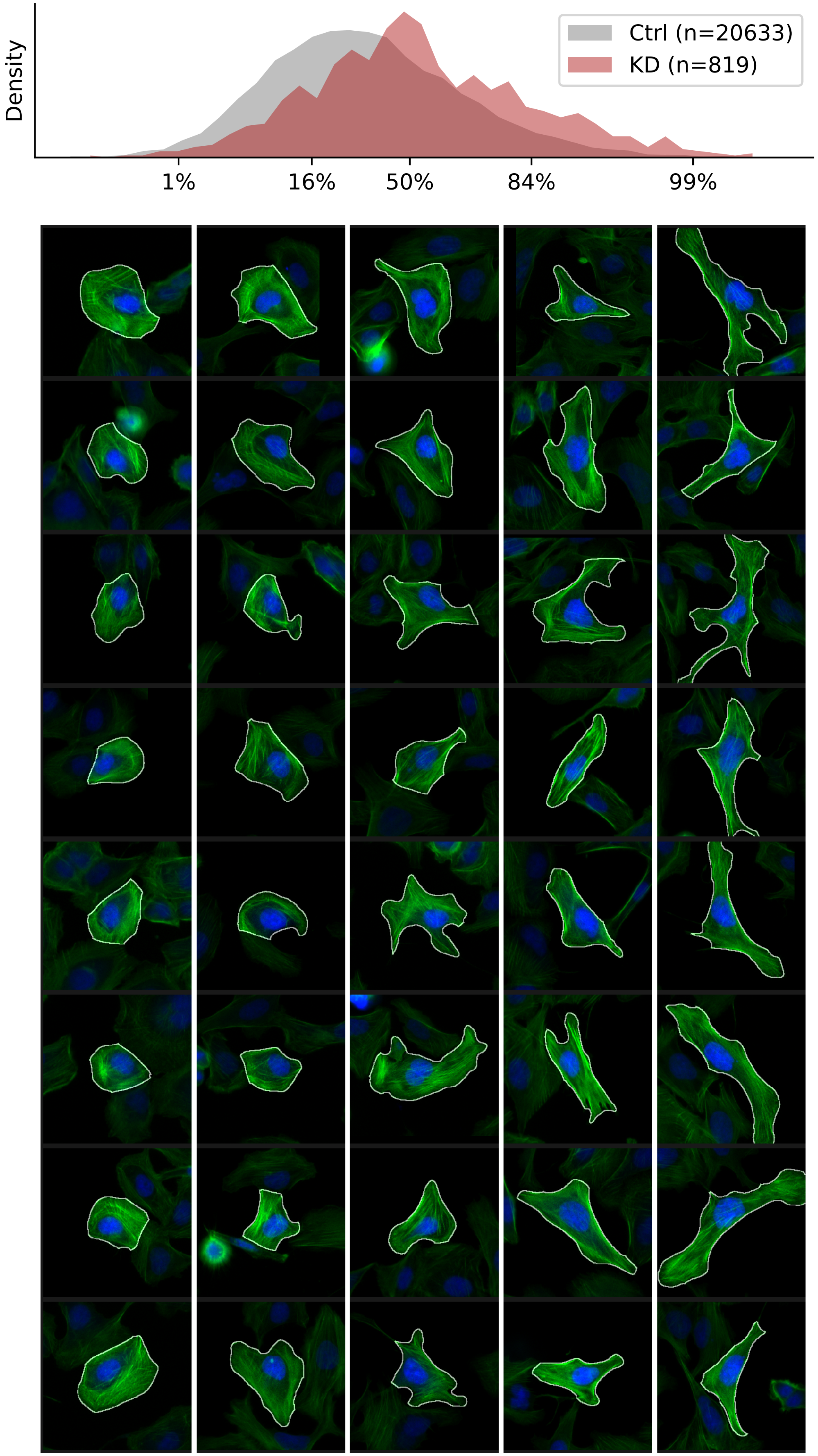

### VIEWS_fPC_PXN_rep3.pdf

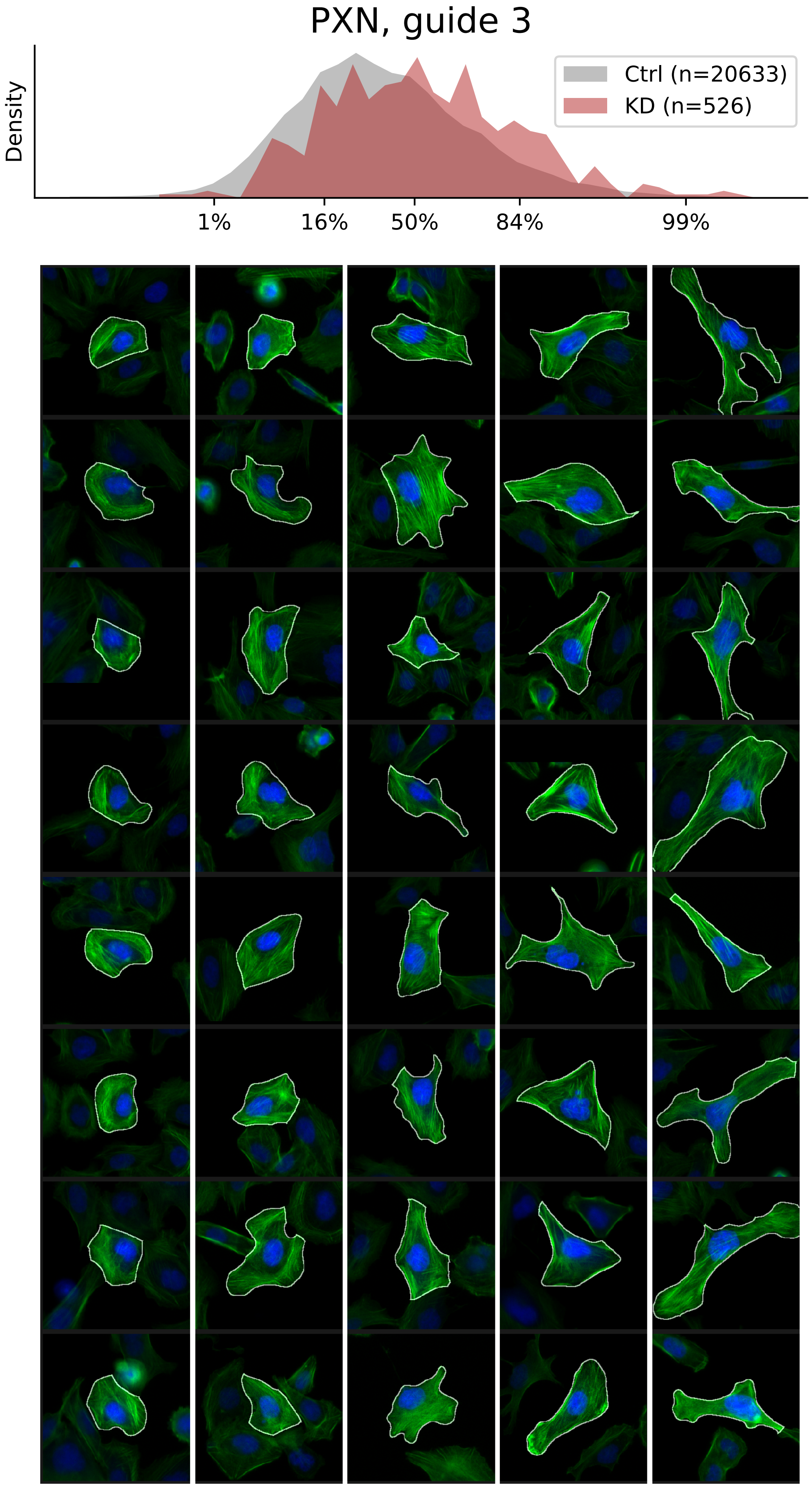

### VIEWS_fPC_ROCK2_rep1.pdf

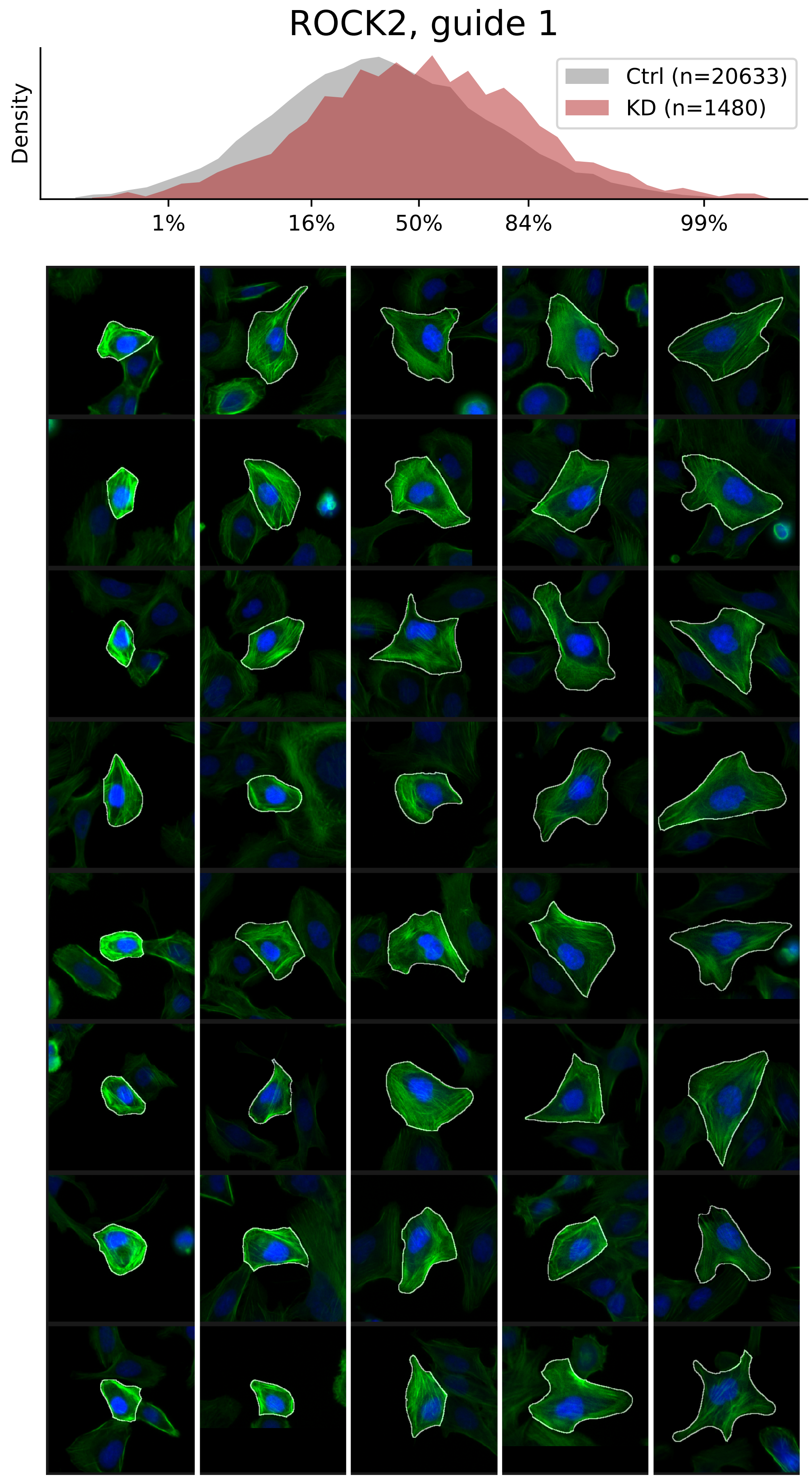

### VIEWS_fPC_ROCK2_rep2.pdf

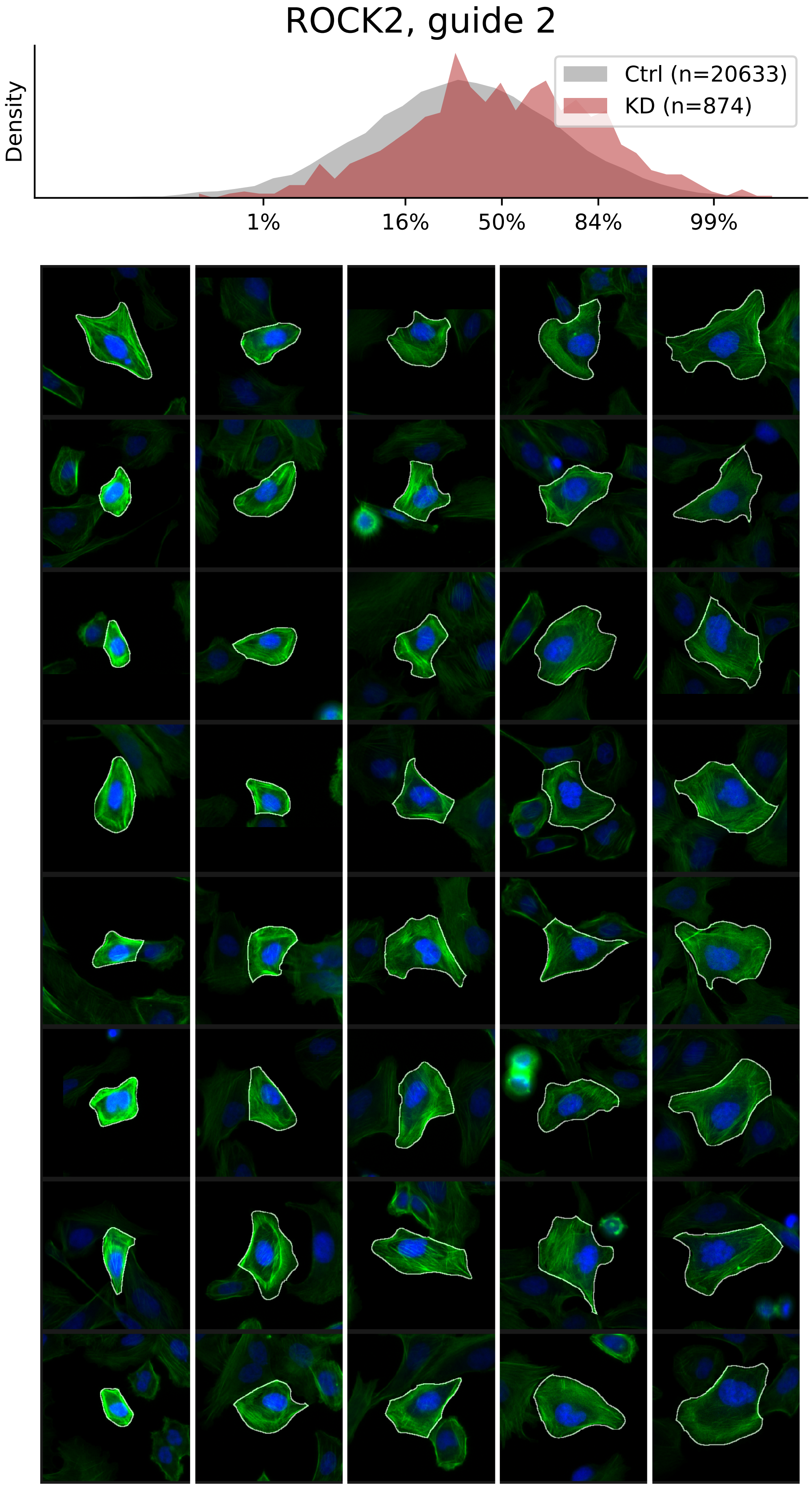

### VIEWS_fPC_SETD3_rep1.pdf

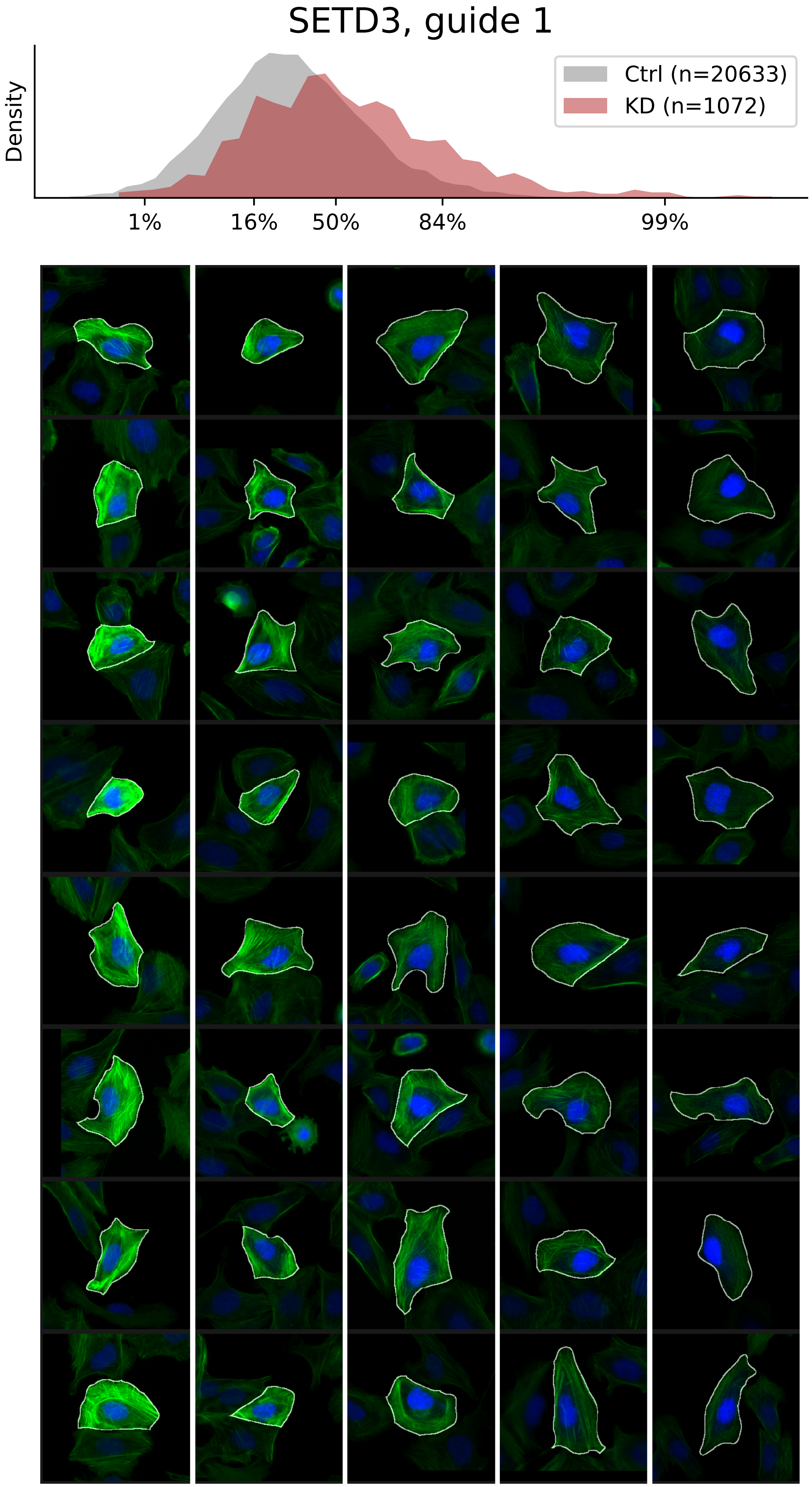

### VIEWS_fPC_SETD3_rep2.pdf

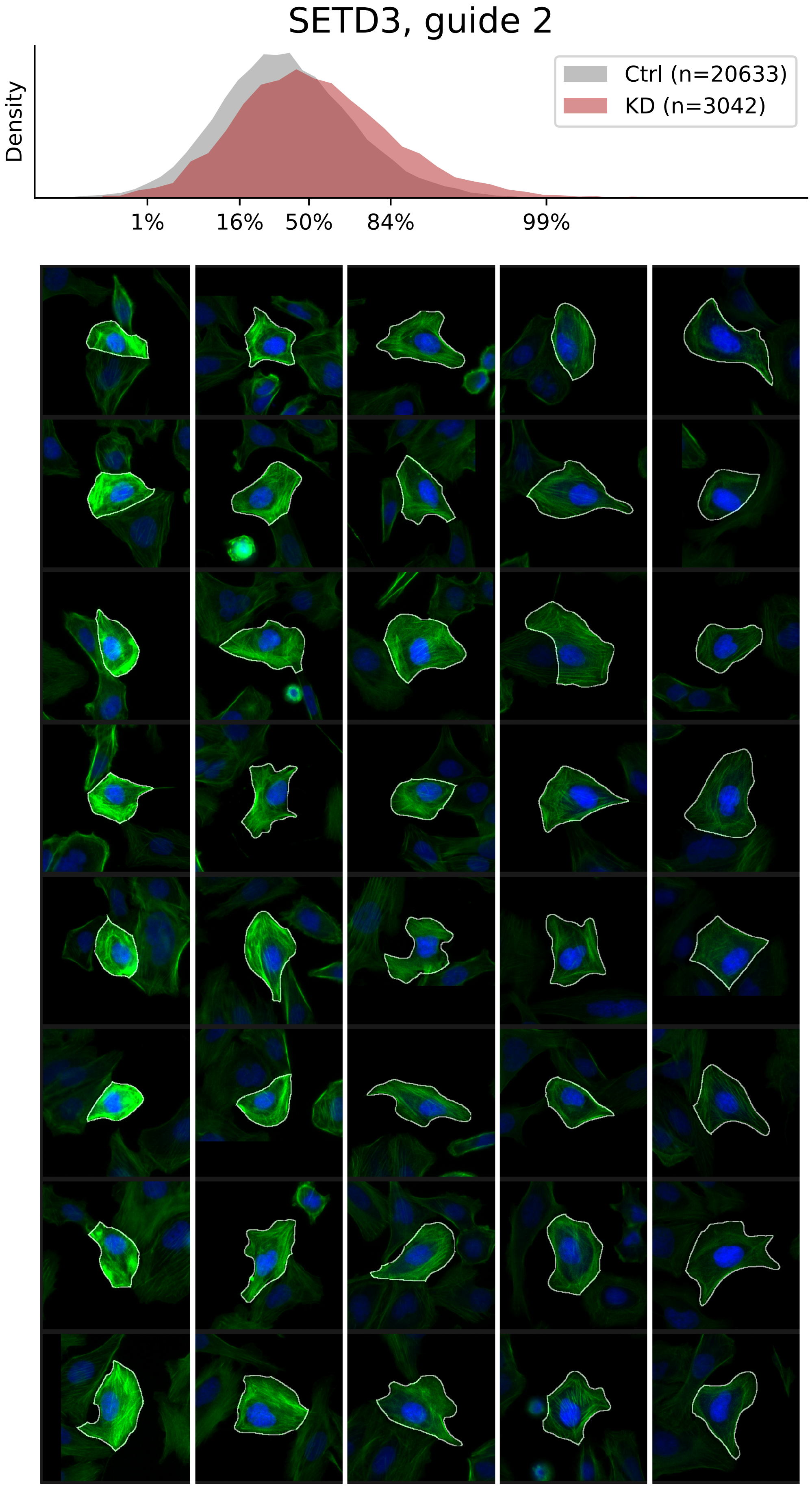

### VIEWS_fPC_SETD3_rep3.pdf

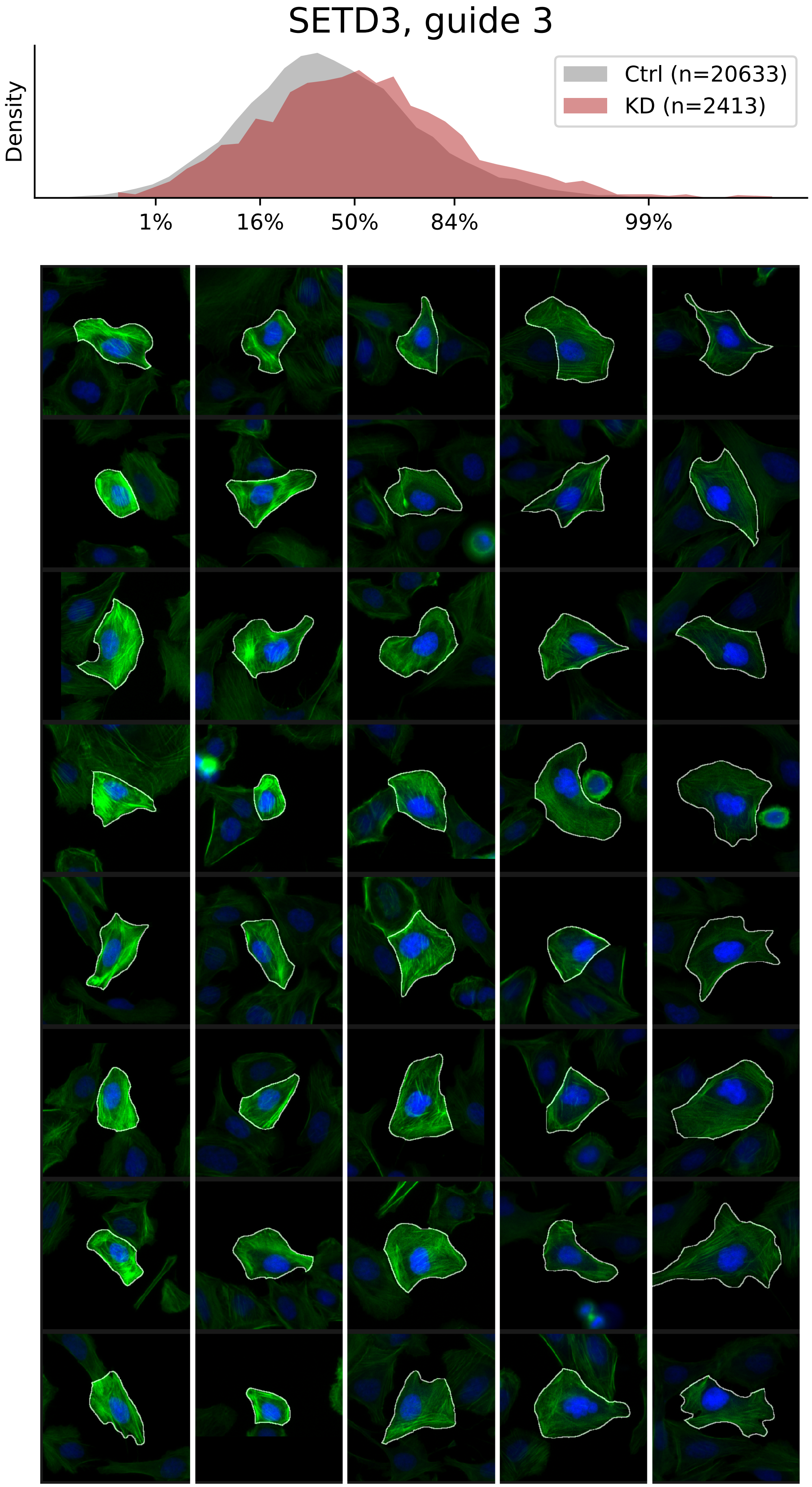

### VIEWS_fPC_TGFBR1_rep2.pdf

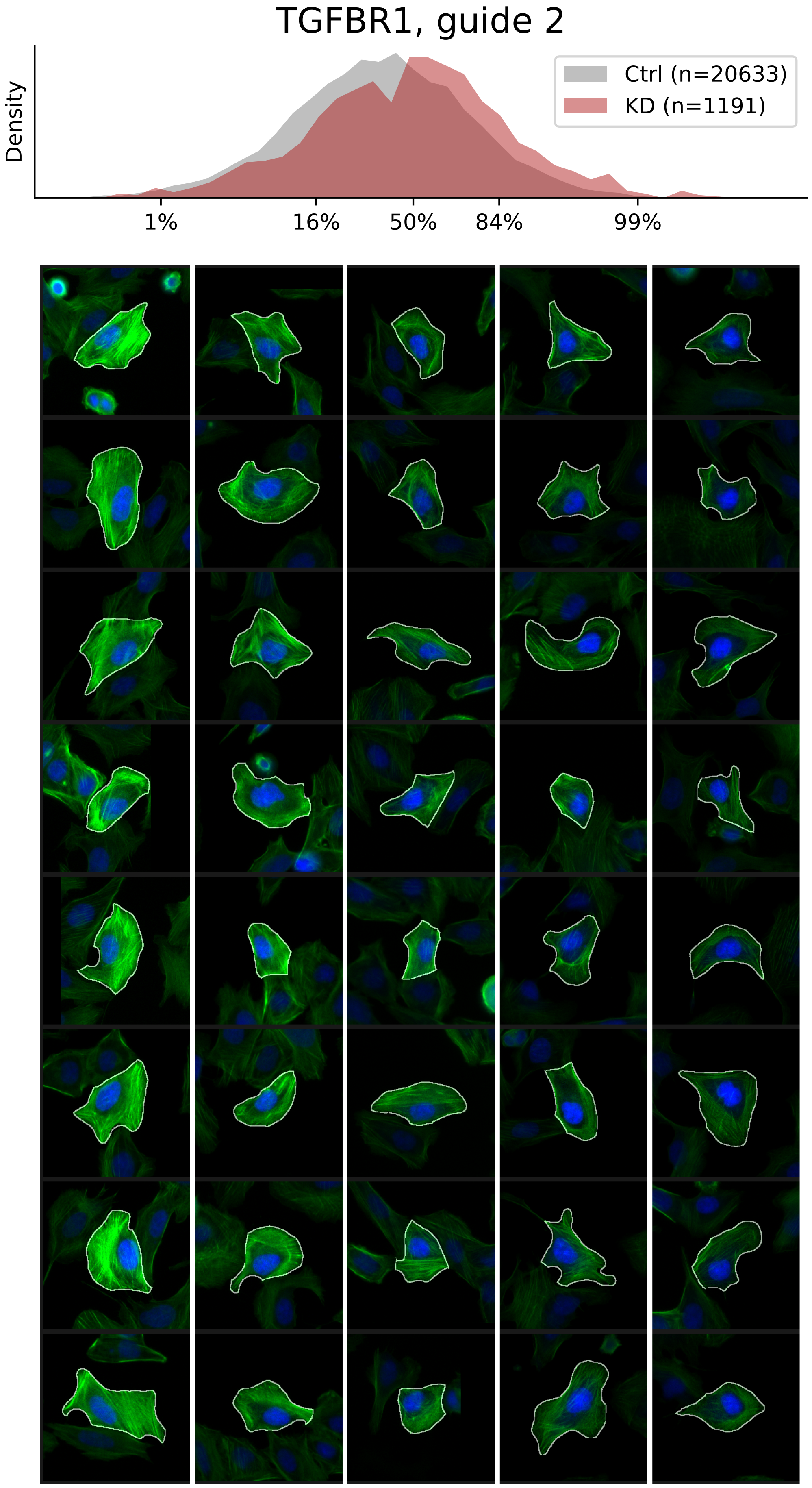

### VIEWS_fPC_TGFBR1_rep3.pdf

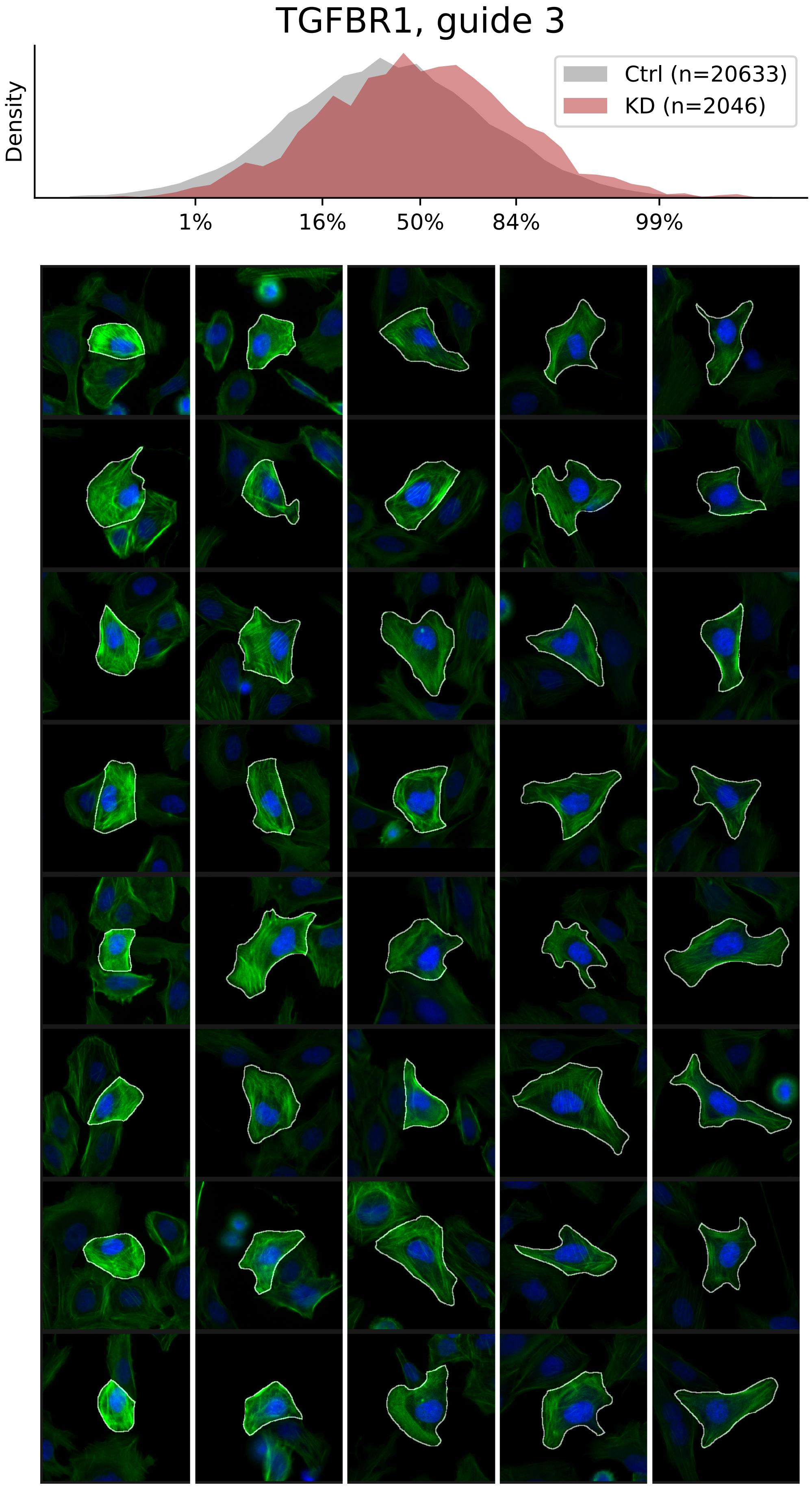

### VIEWS_fPC_TPM3_rep1.pdf

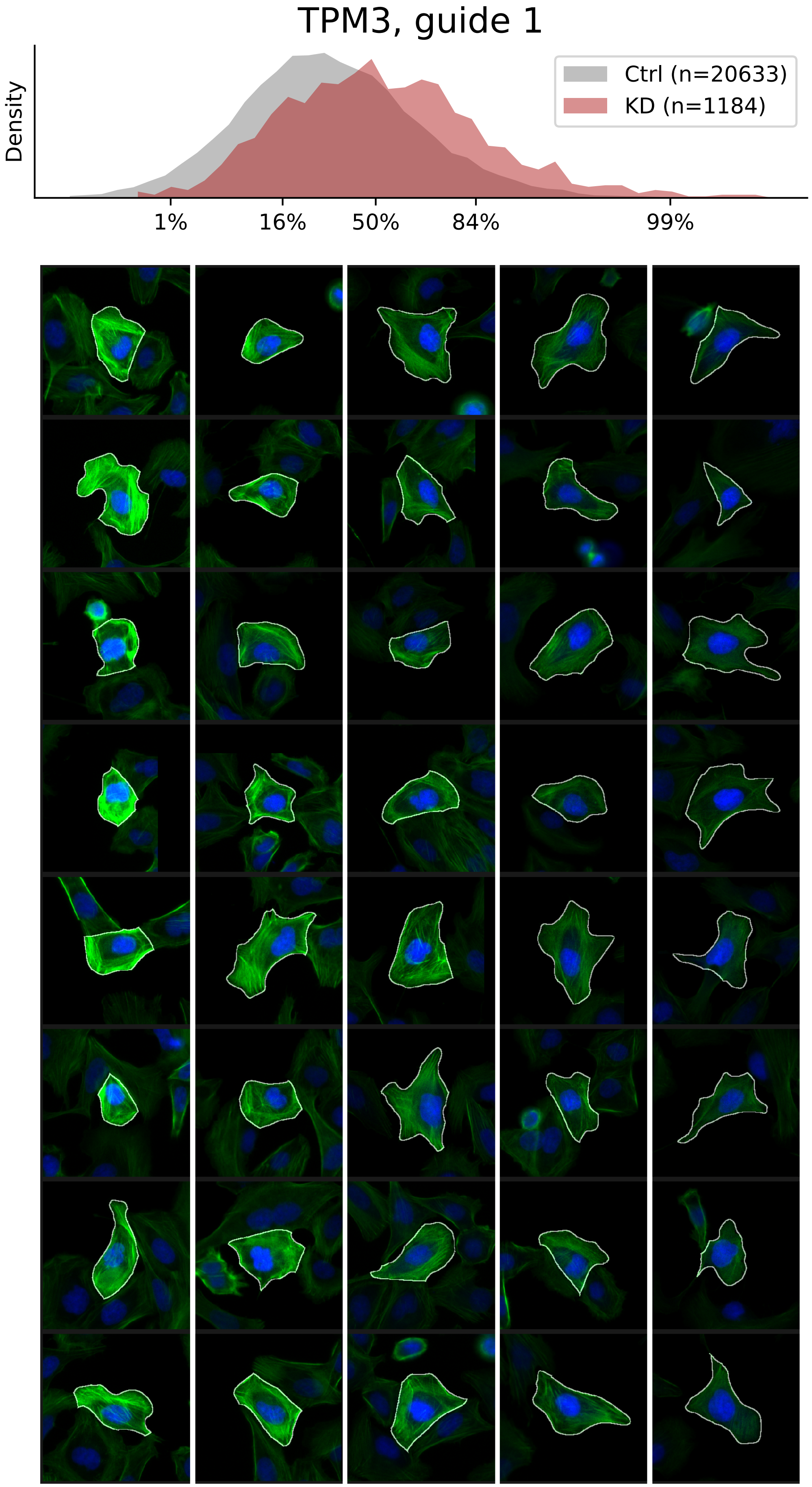

### VIEWS_fPC_TPM3_rep3.pdf

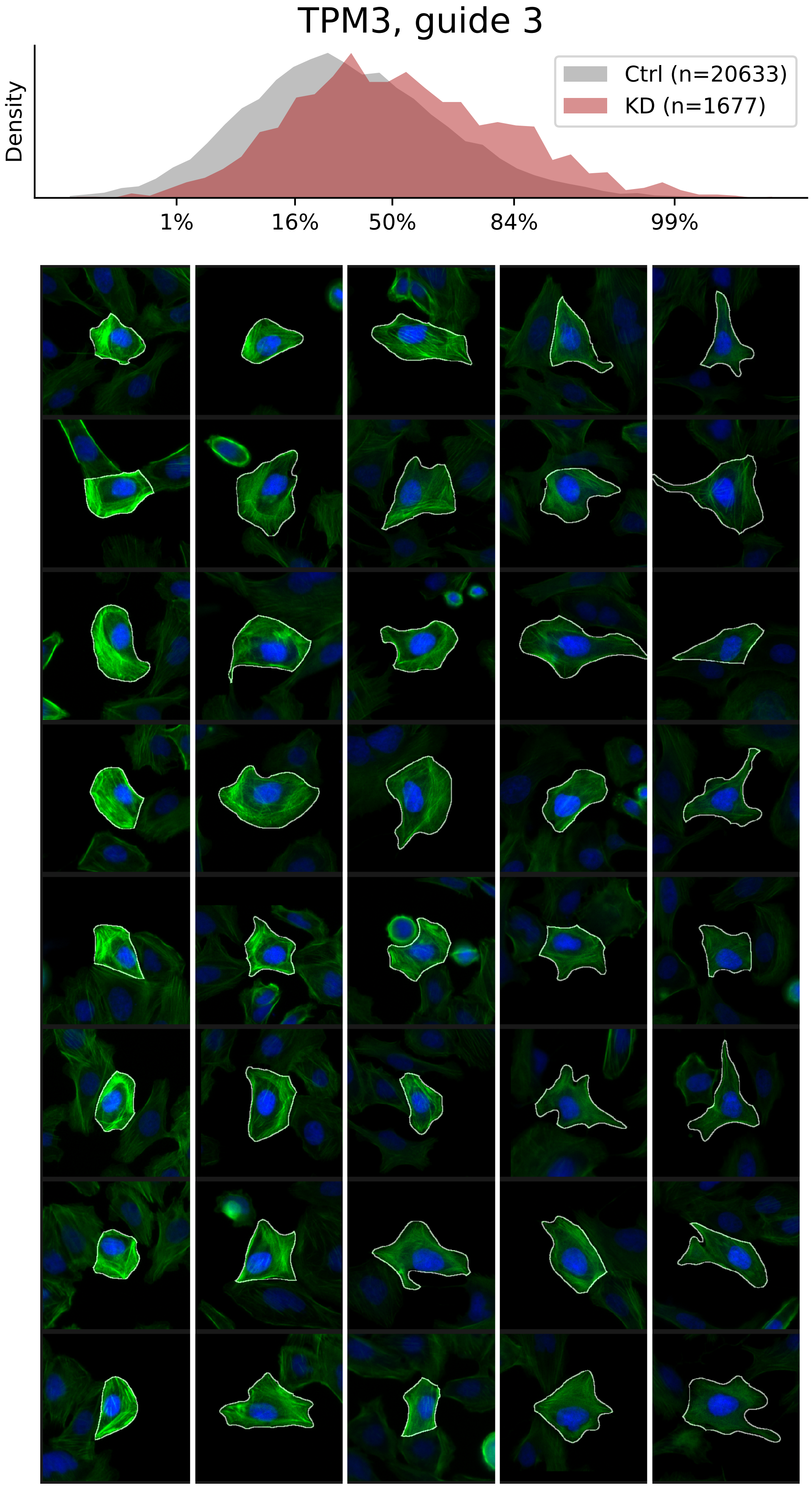

### VIEWS_iAD-LD-verified_ACTR3_rep2.pdf

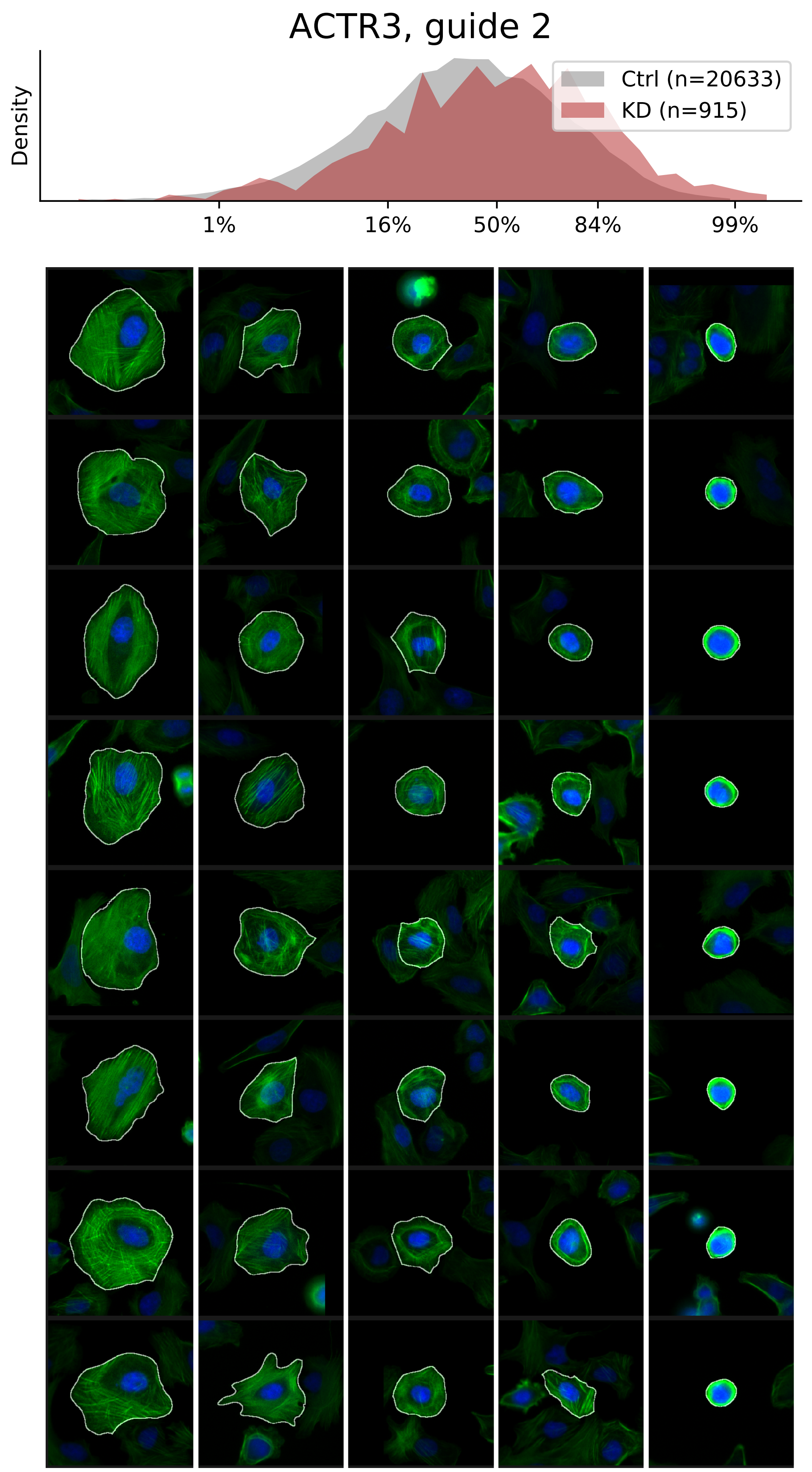

### VIEWS_iAD-LD-verified_GNA12_rep2.pdf

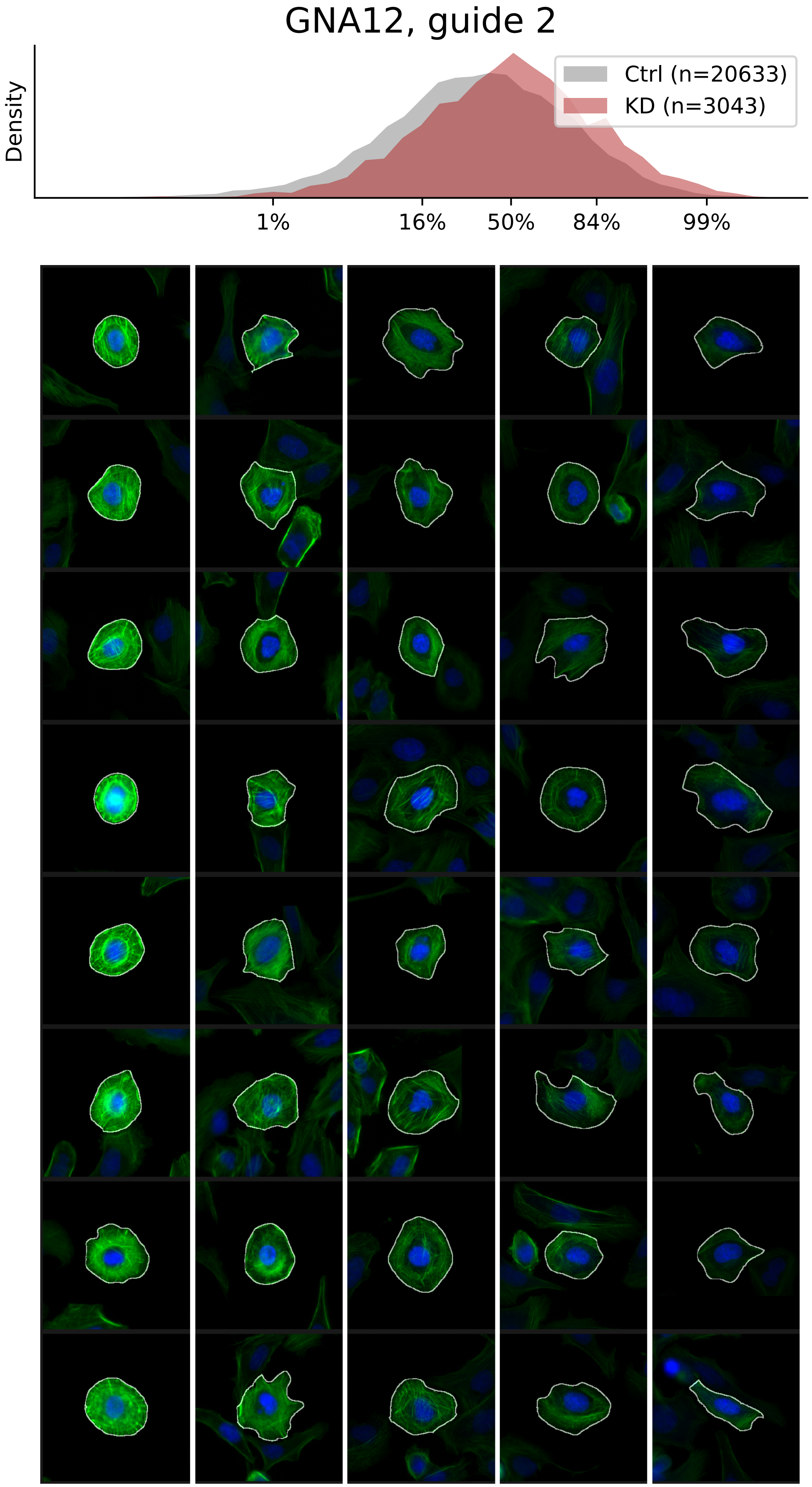

### VIEWS_iAD-LD-verified_MESDC1_rep3.pdf

#### MESDC1, guide 3

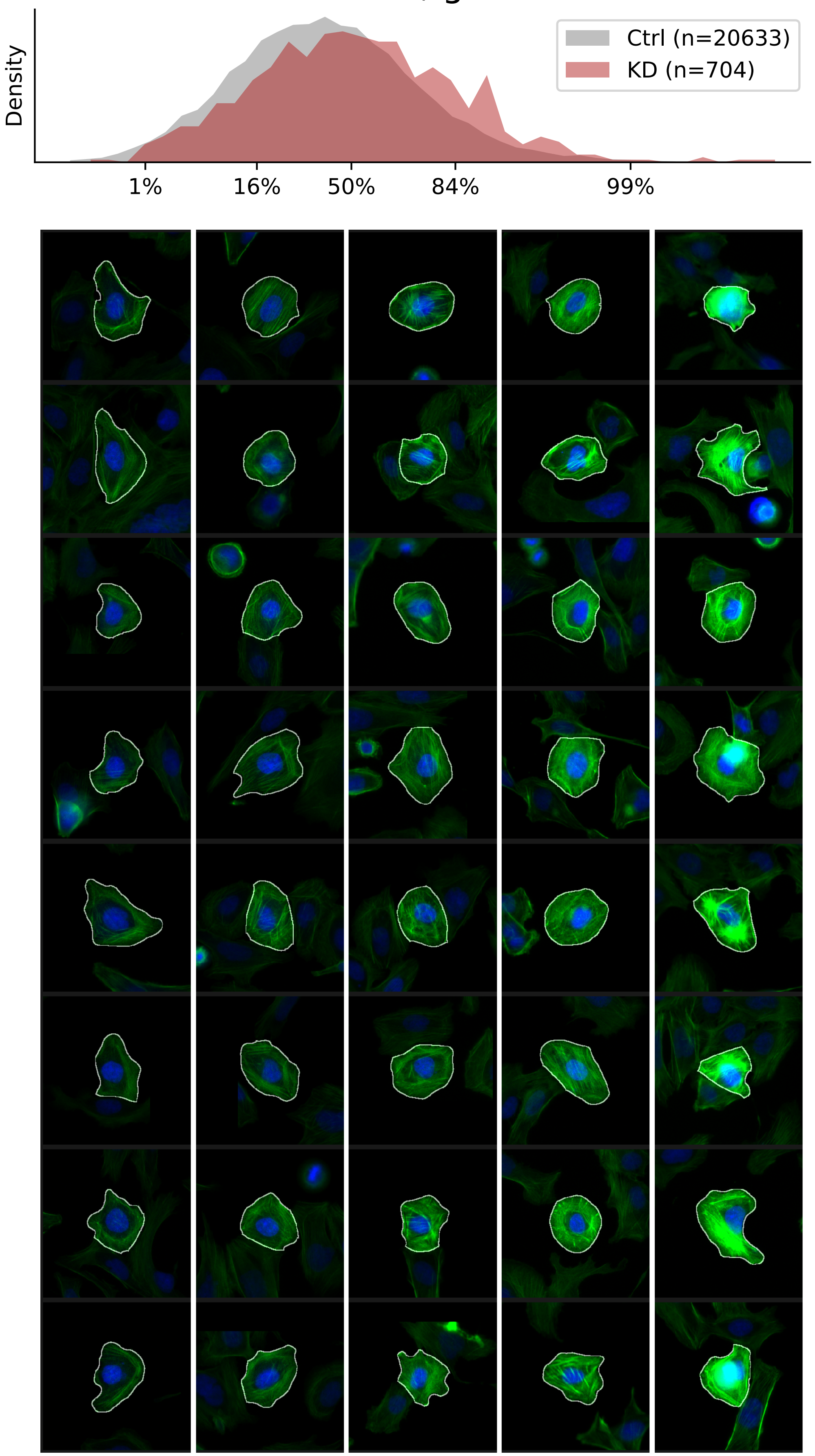

### VIEWS_iAD-LD-verified_MYO9B_rep1.pdf

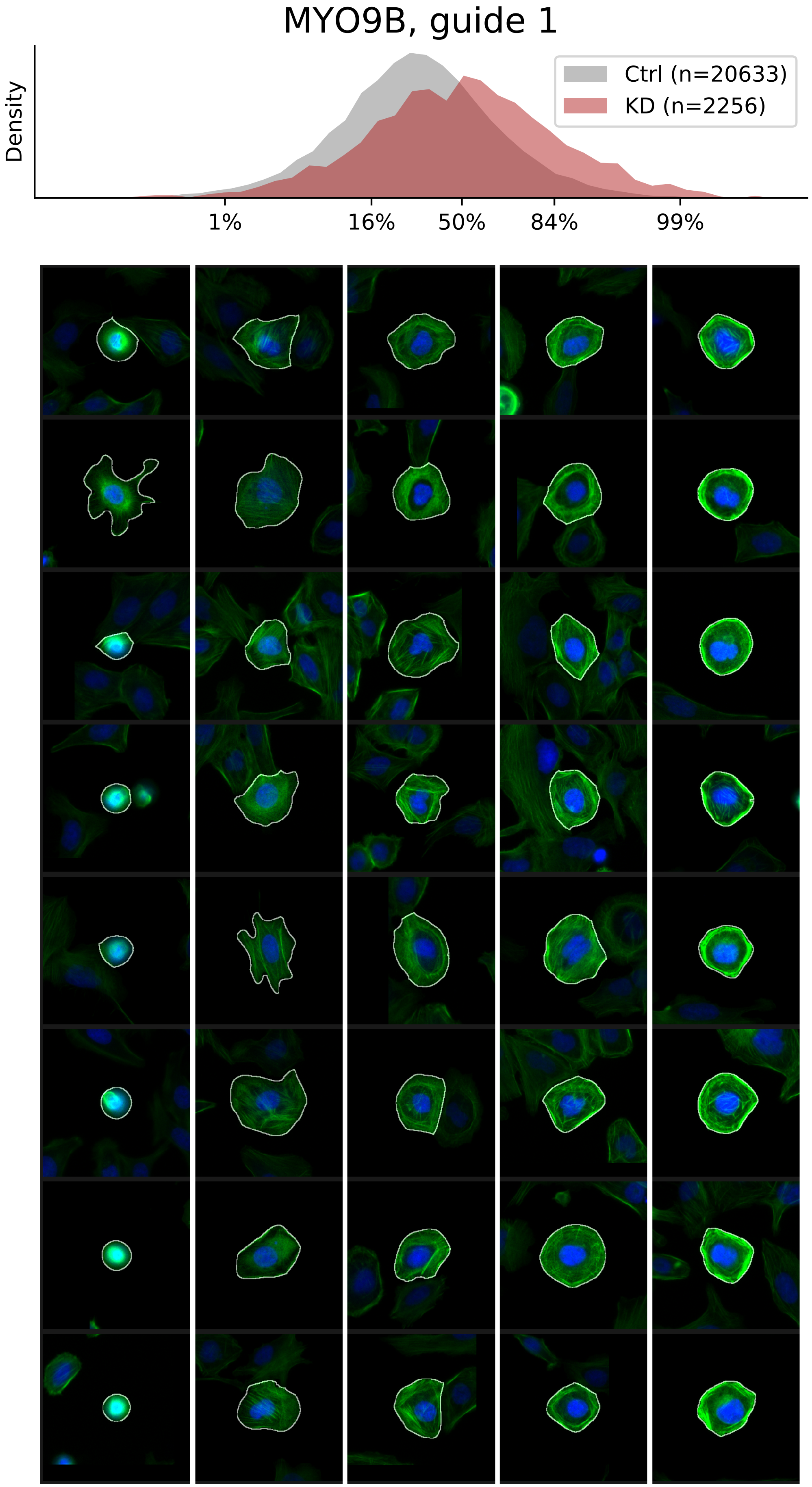

### VIEWS_iAD-LD-verified_RAC1_rep3.pdf

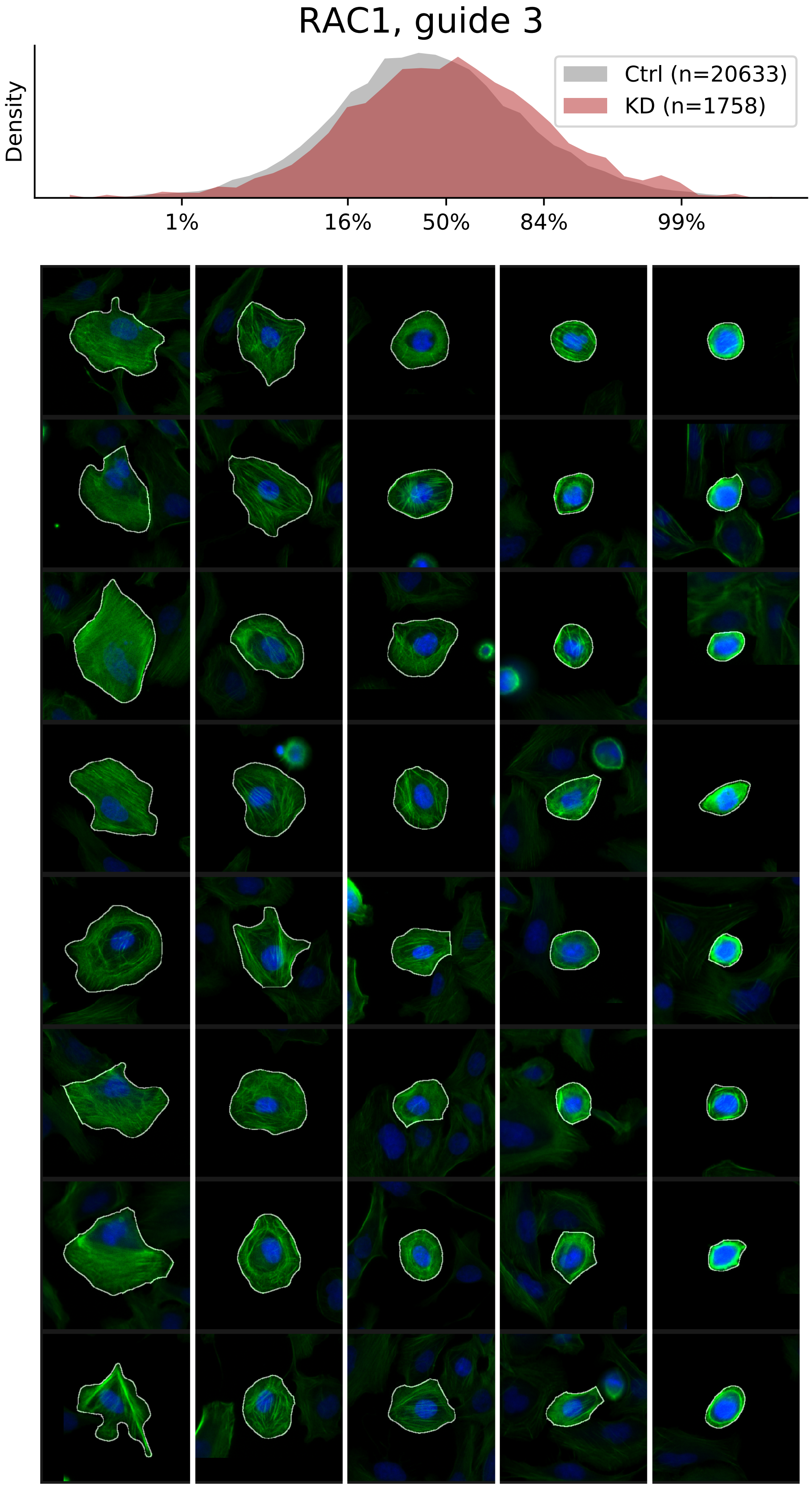

### VIEWS_iAD-LD-verified_SEPT7_rep1.pdf

#### SEPT7, guide 1

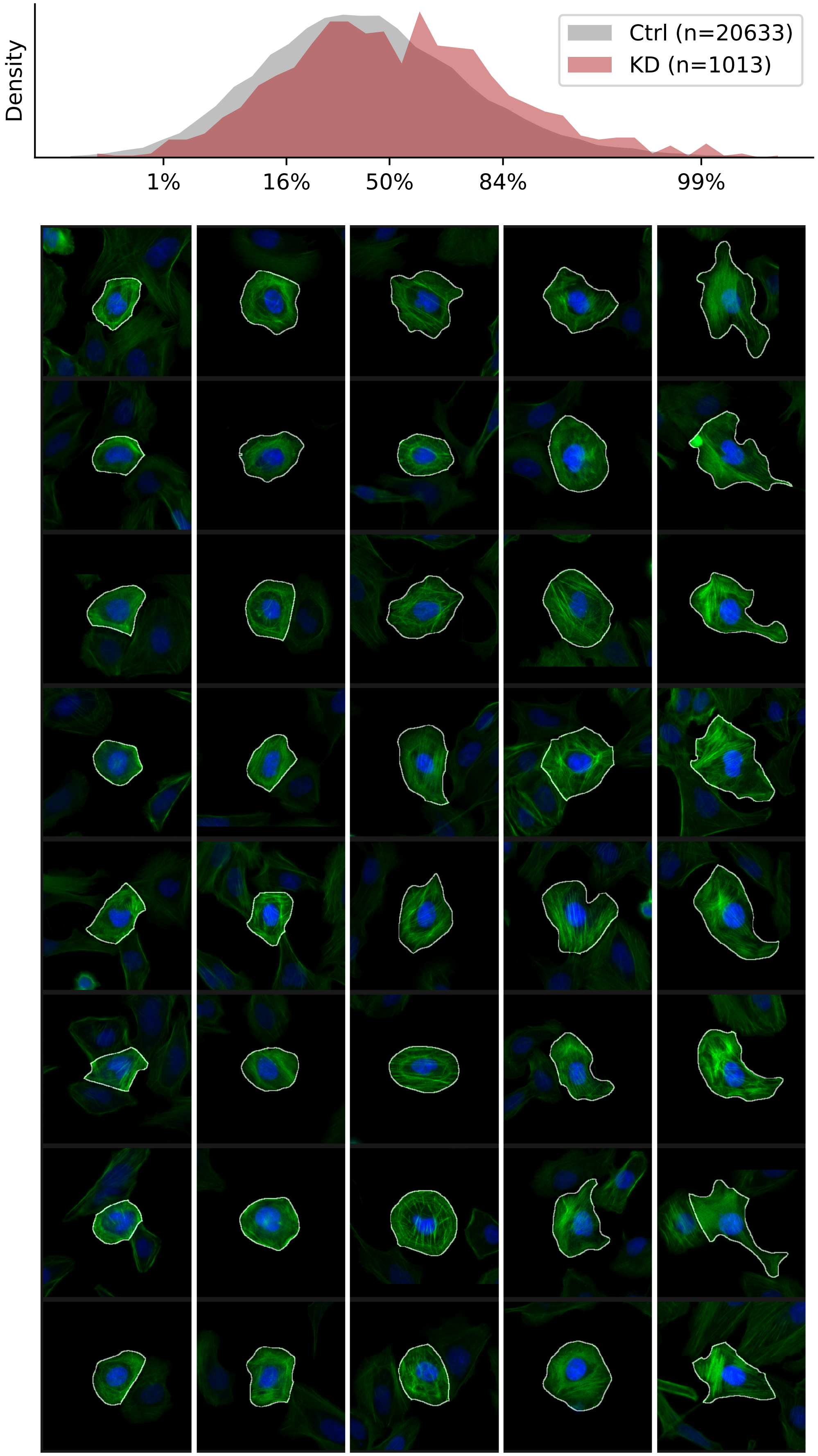

### VIEWS_iAD-LD-verified_TGFBR1_rep3.pdf

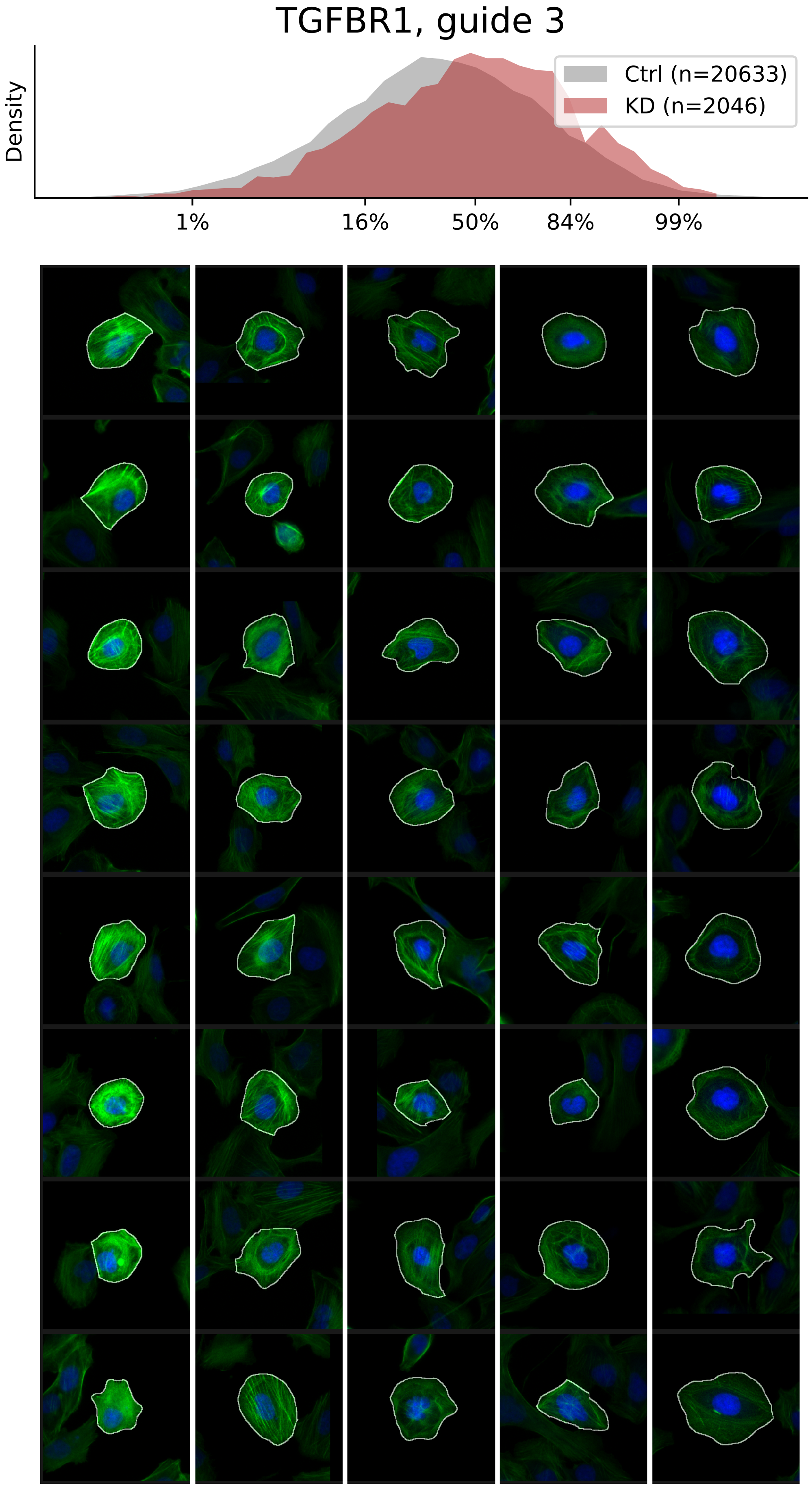

### VIEWS_iAD-LD-verified_TMOD3_rep2.pdf

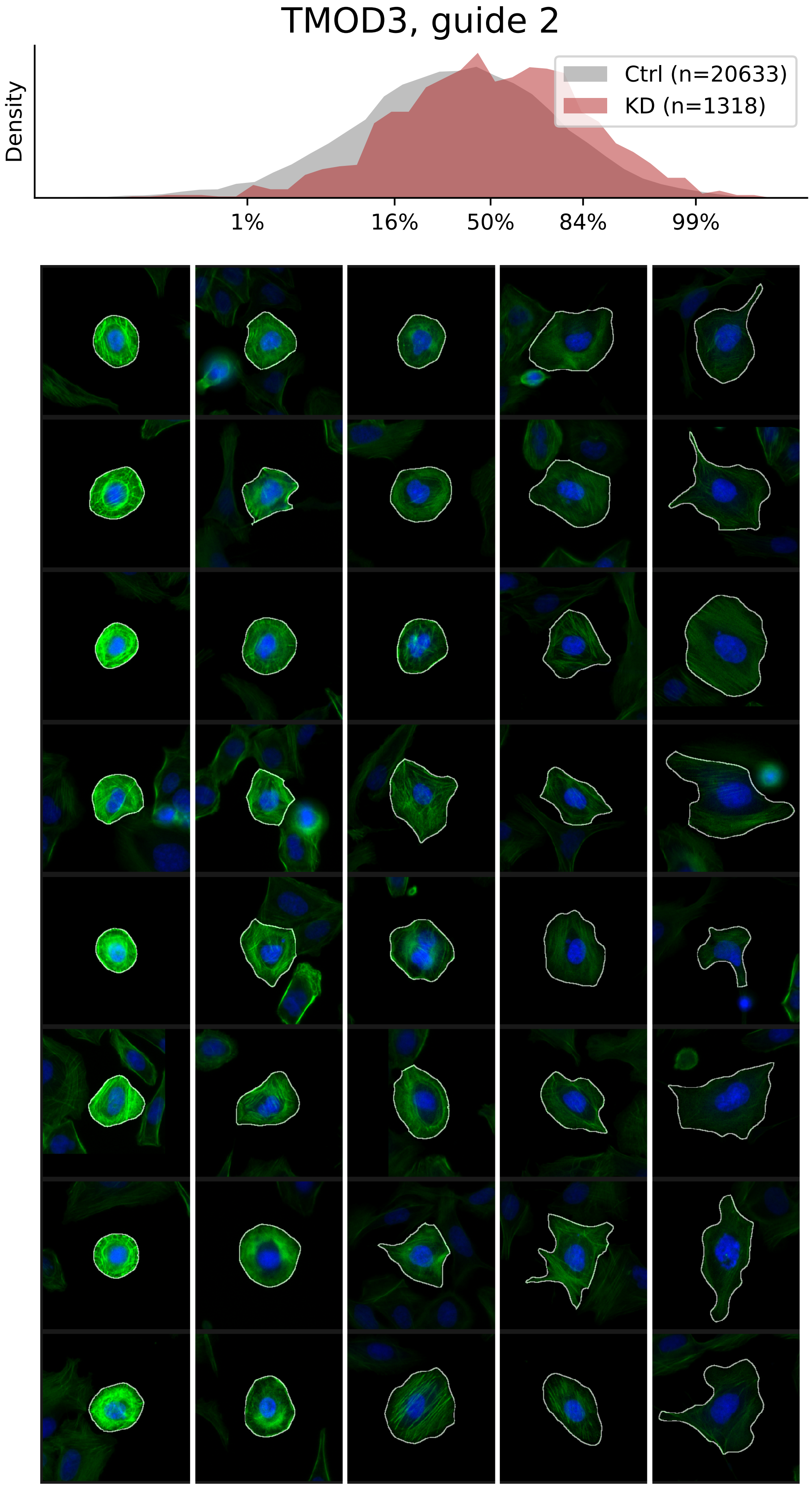

### VIEWS_iAD-LD-verified_TUBA1B_rep3.pdf

#### TUBA1B, guide 3

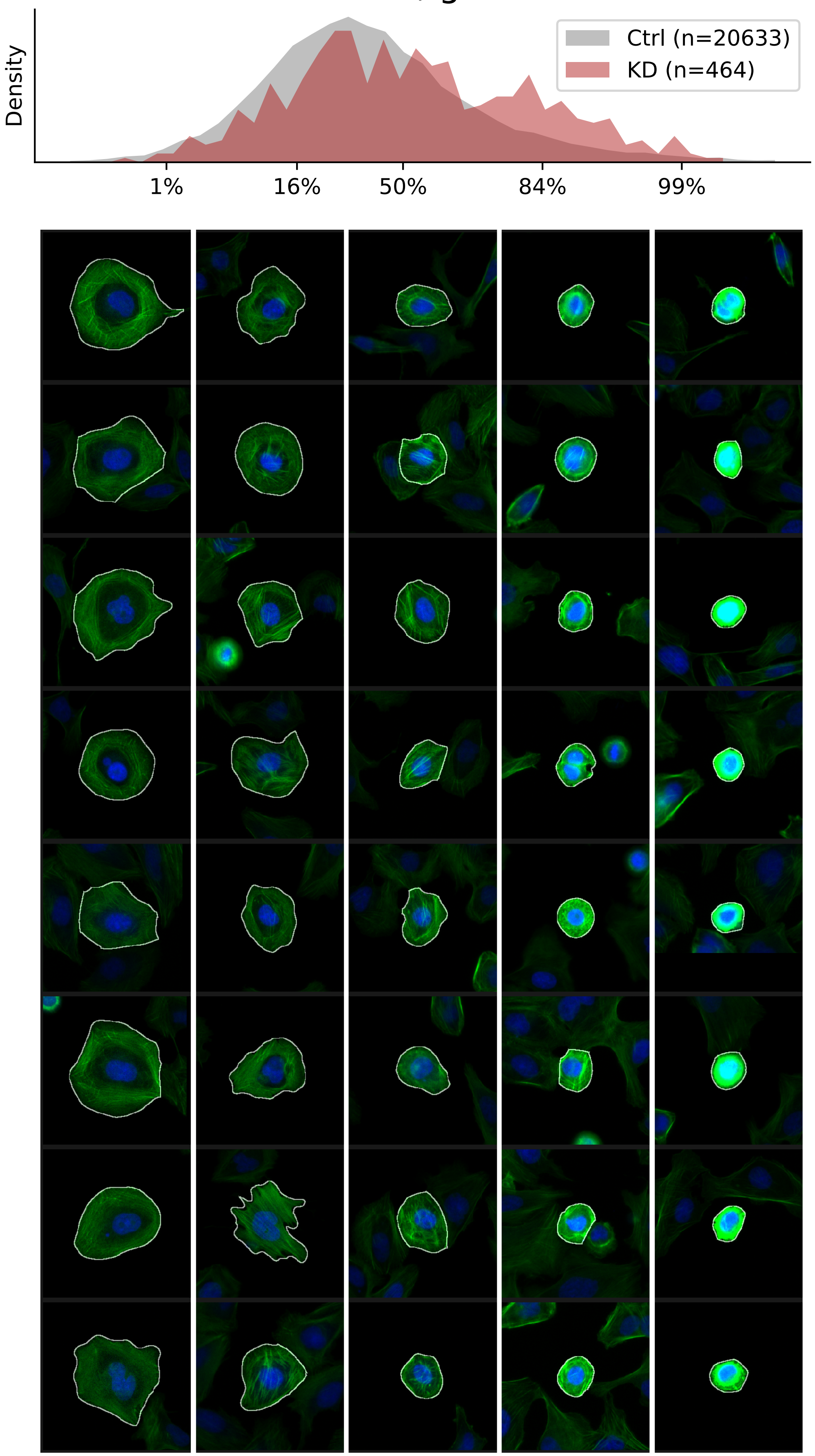

### VIEWS_iAD_ACTR2_rep1.pdf

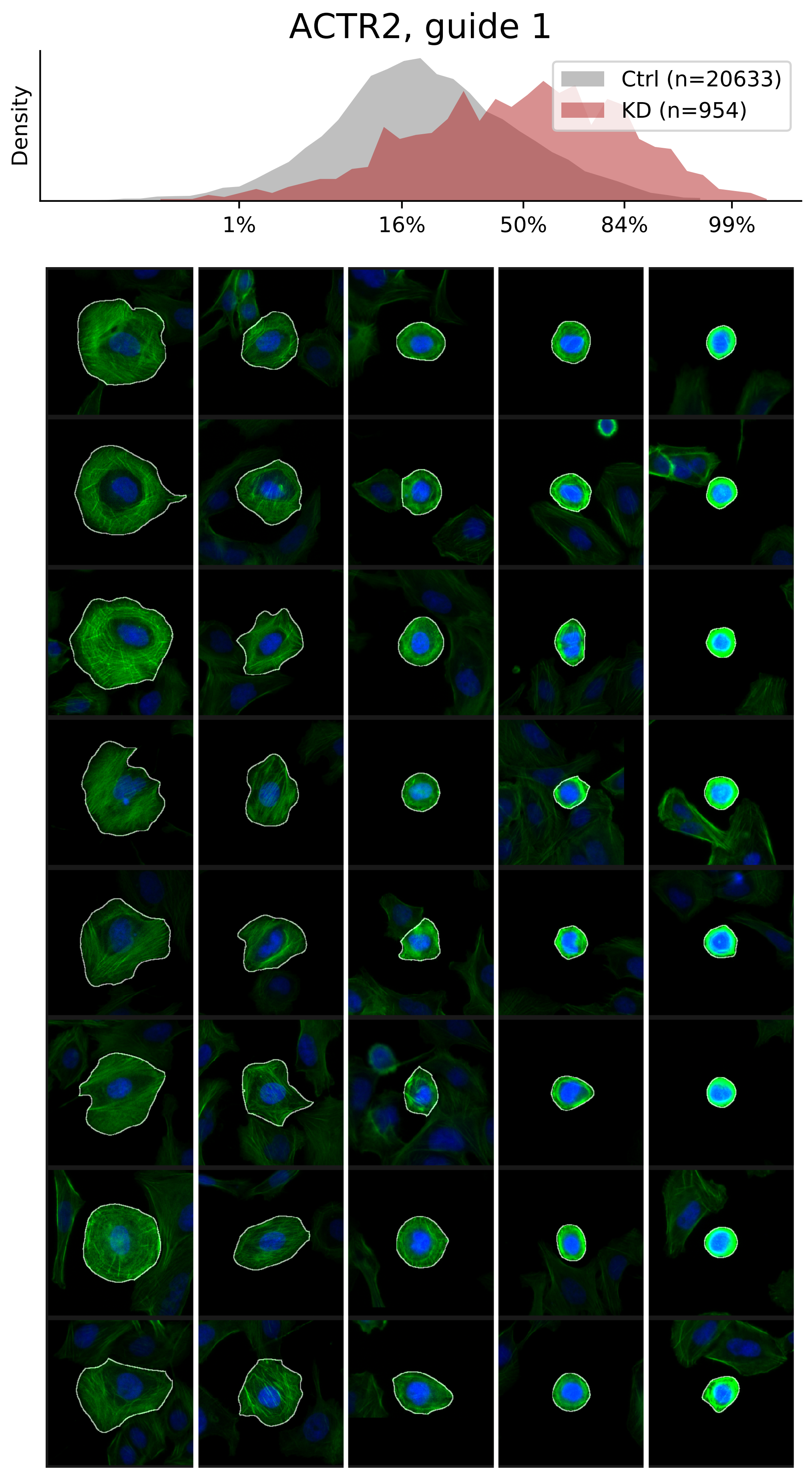

### VIEWS_iAD_ACTR2_rep3.pdf

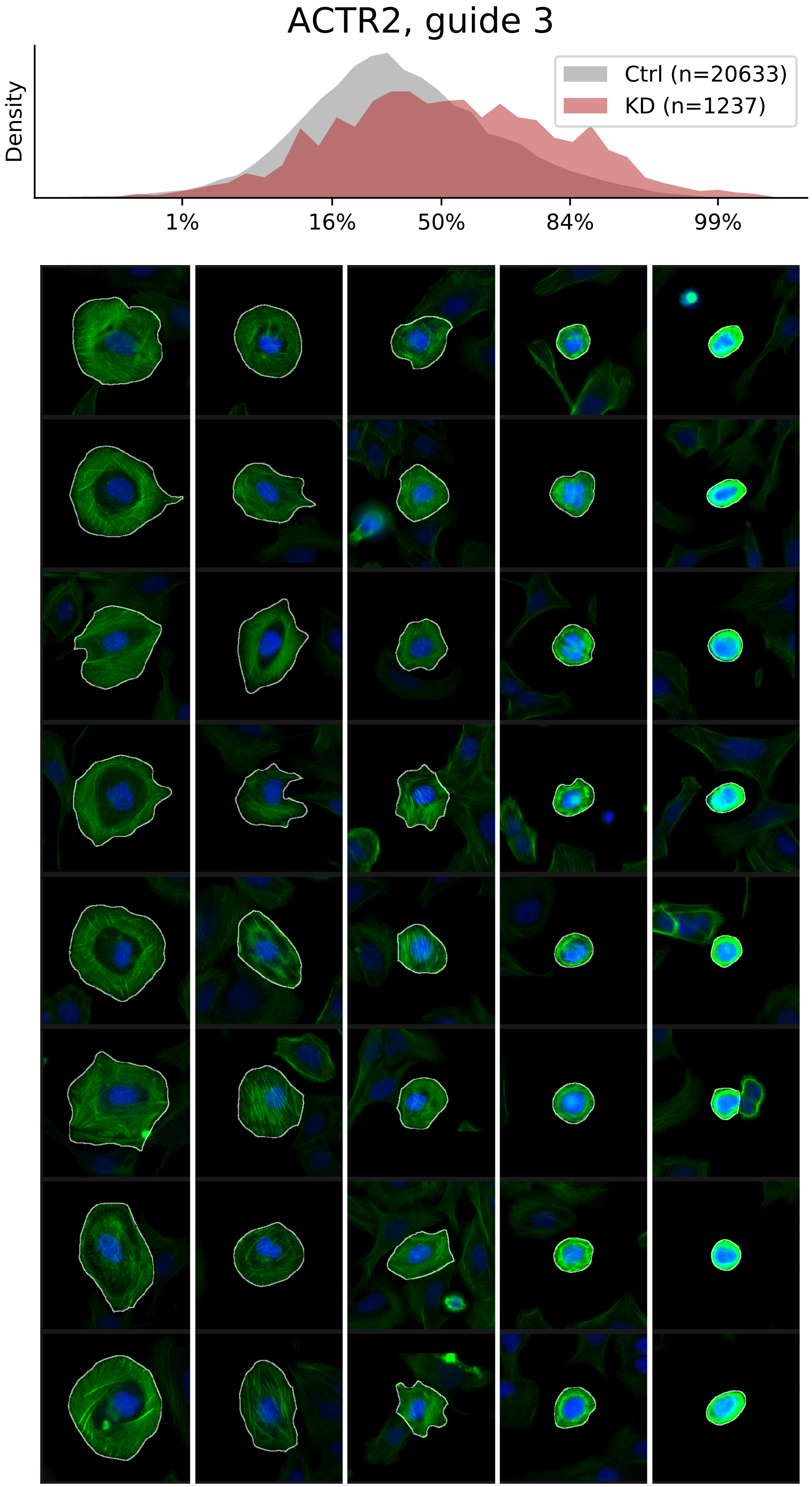

### VIEWS_iAD_BCAR1_rep1.pdf

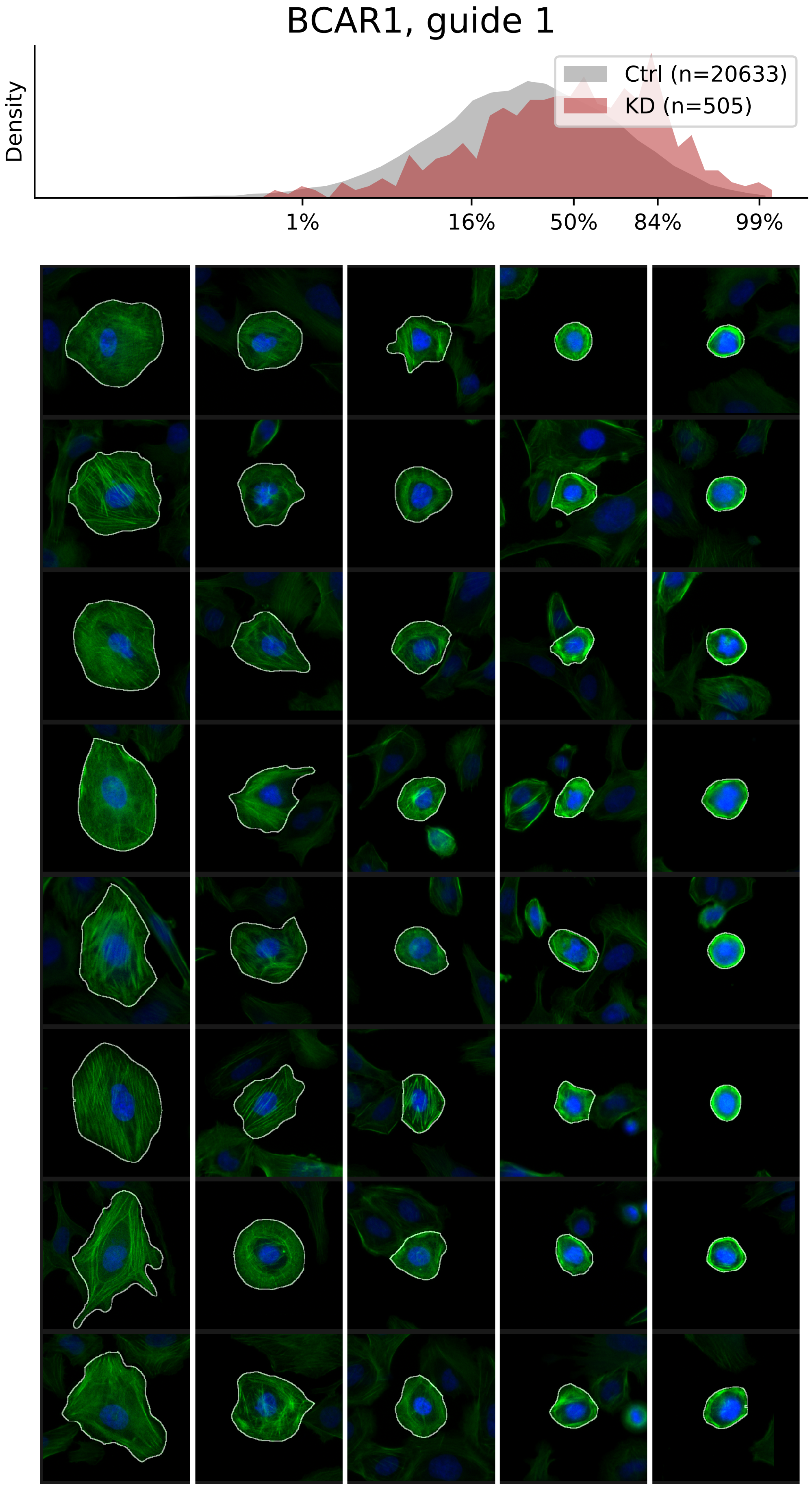

### VIEWS_iAD_BCAR1_rep2.pdf

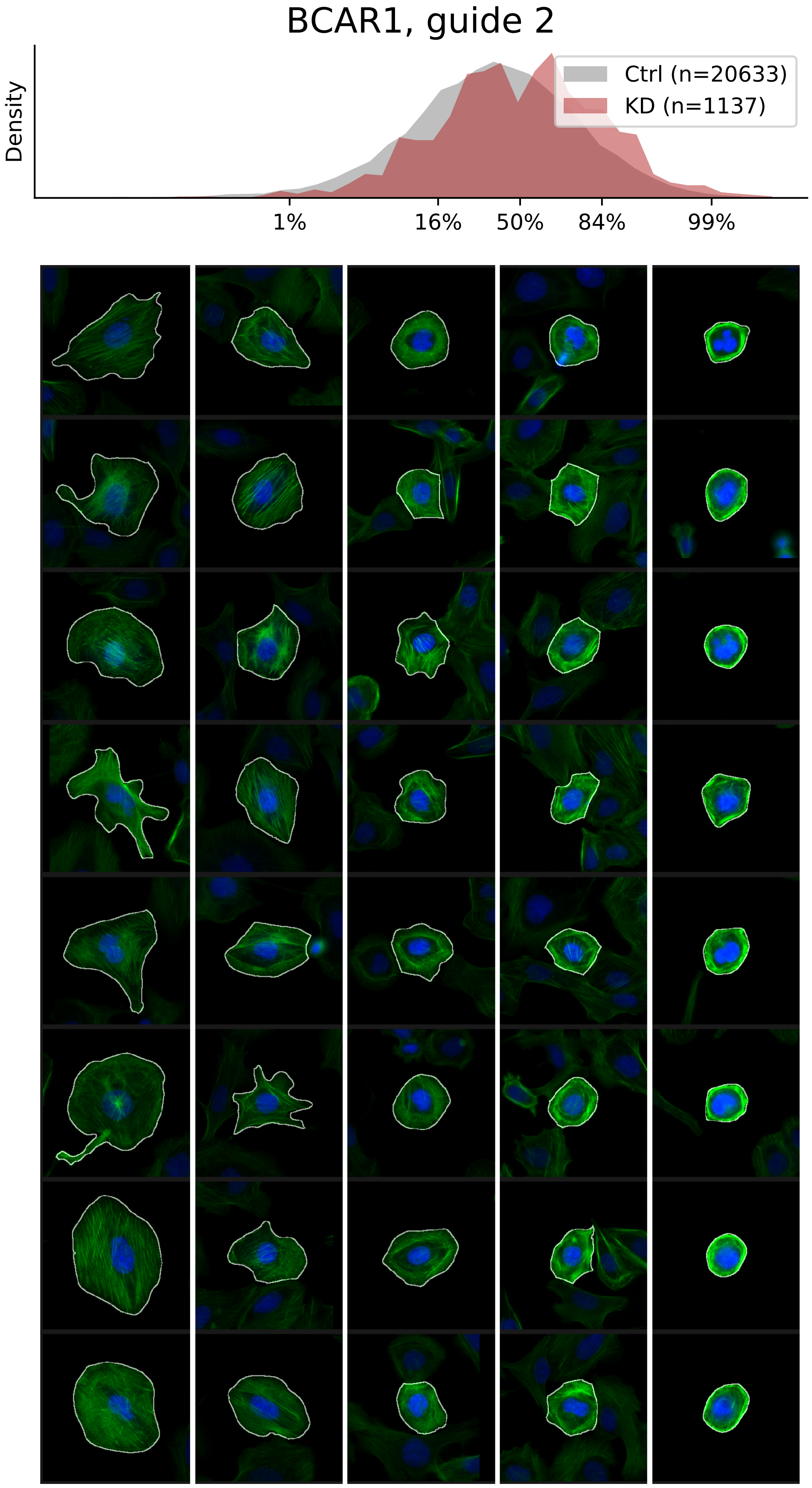

### VIEWS_iAD_ILK_rep2.pdf

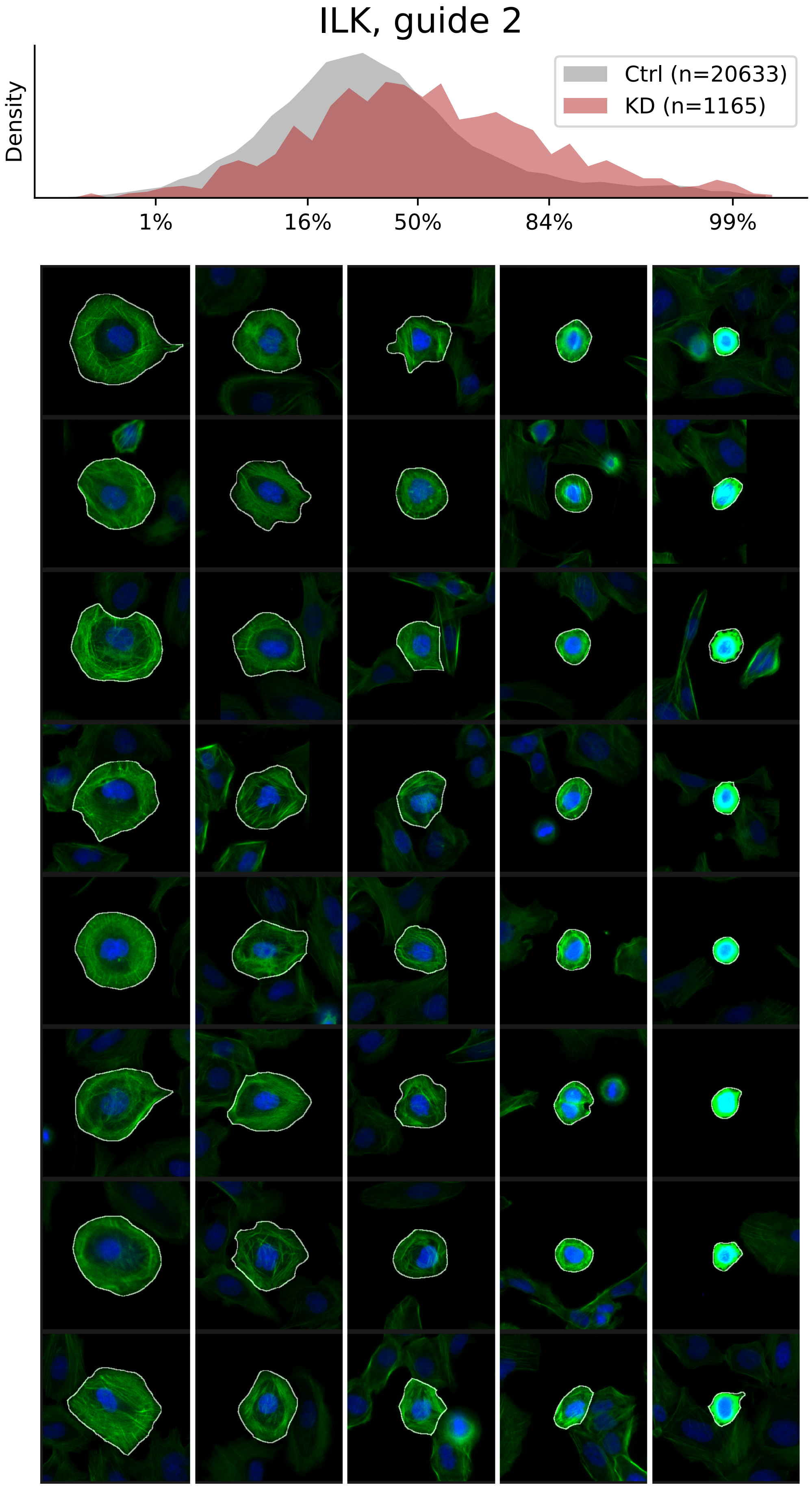

### VIEWS_iAD_ILK_rep3.pdf

#### ILK, guide 3

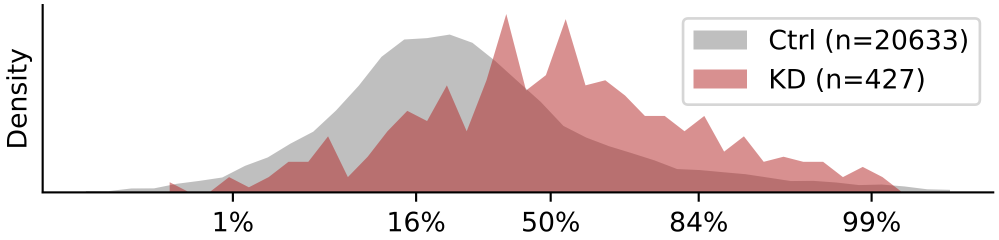


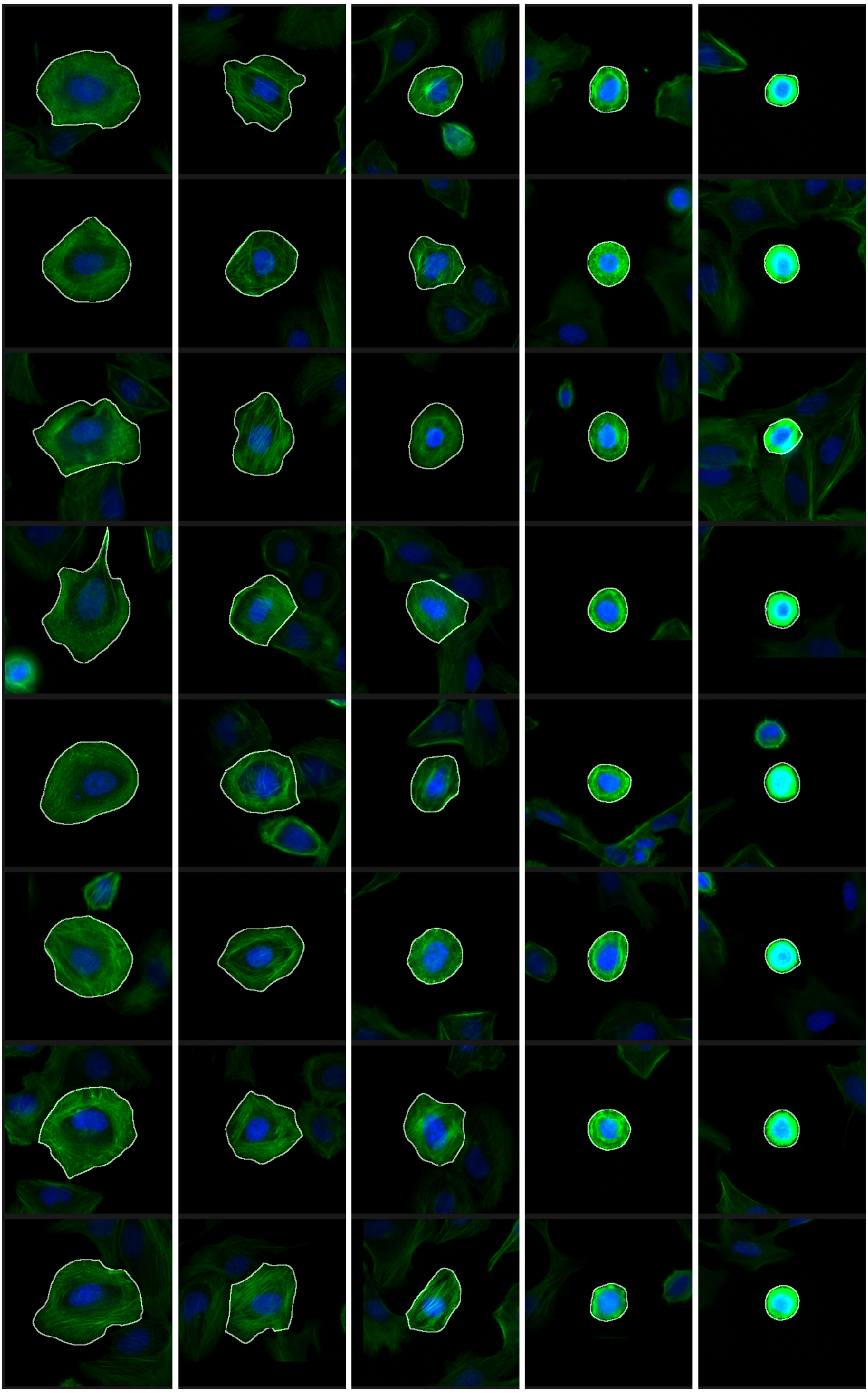

### VIEWS_iAD_LIMK2_rep2.pdf

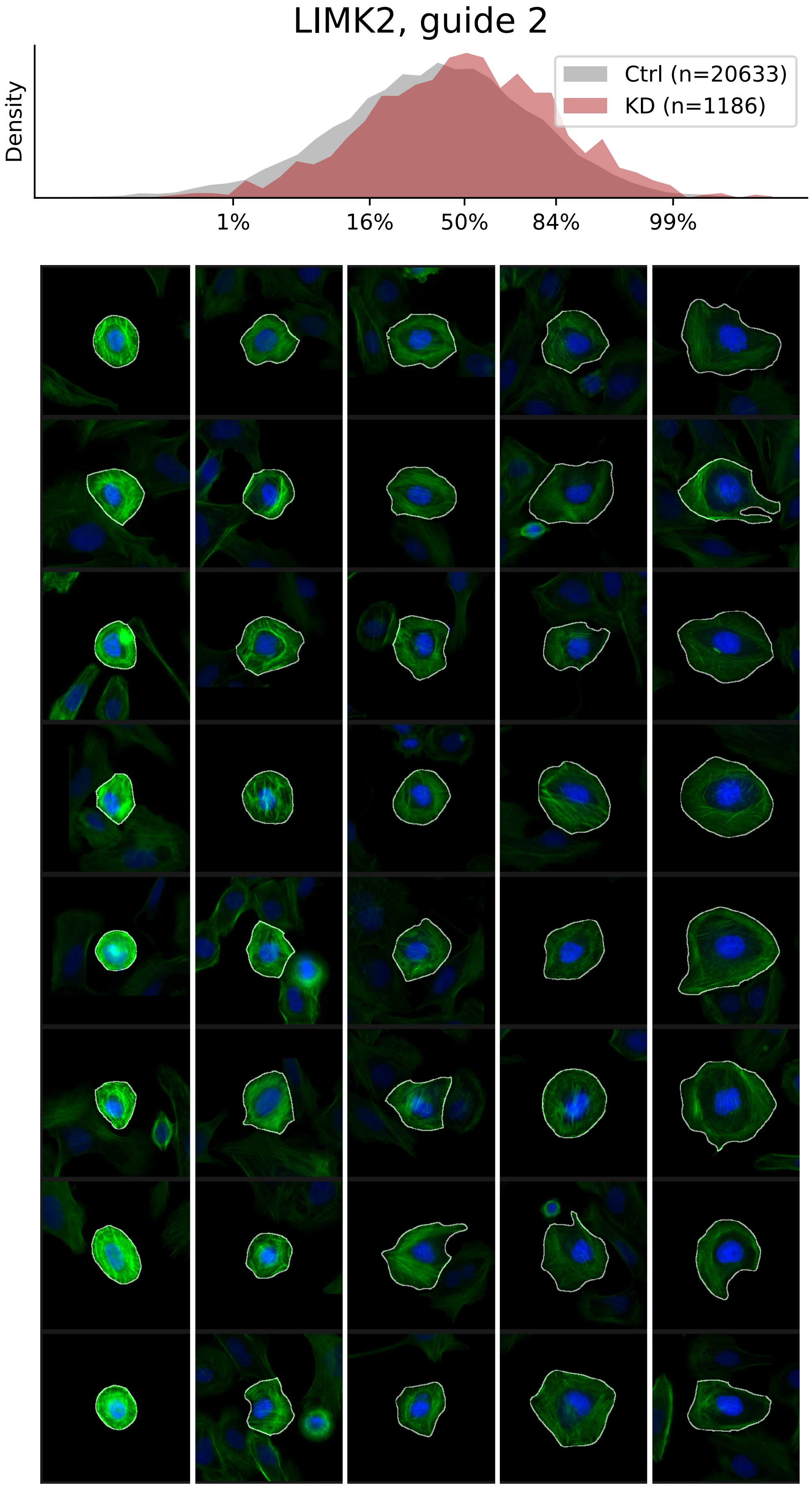

### VIEWS_iAD_LIMK2_rep3.pdf

#### LIMK2, guide 3

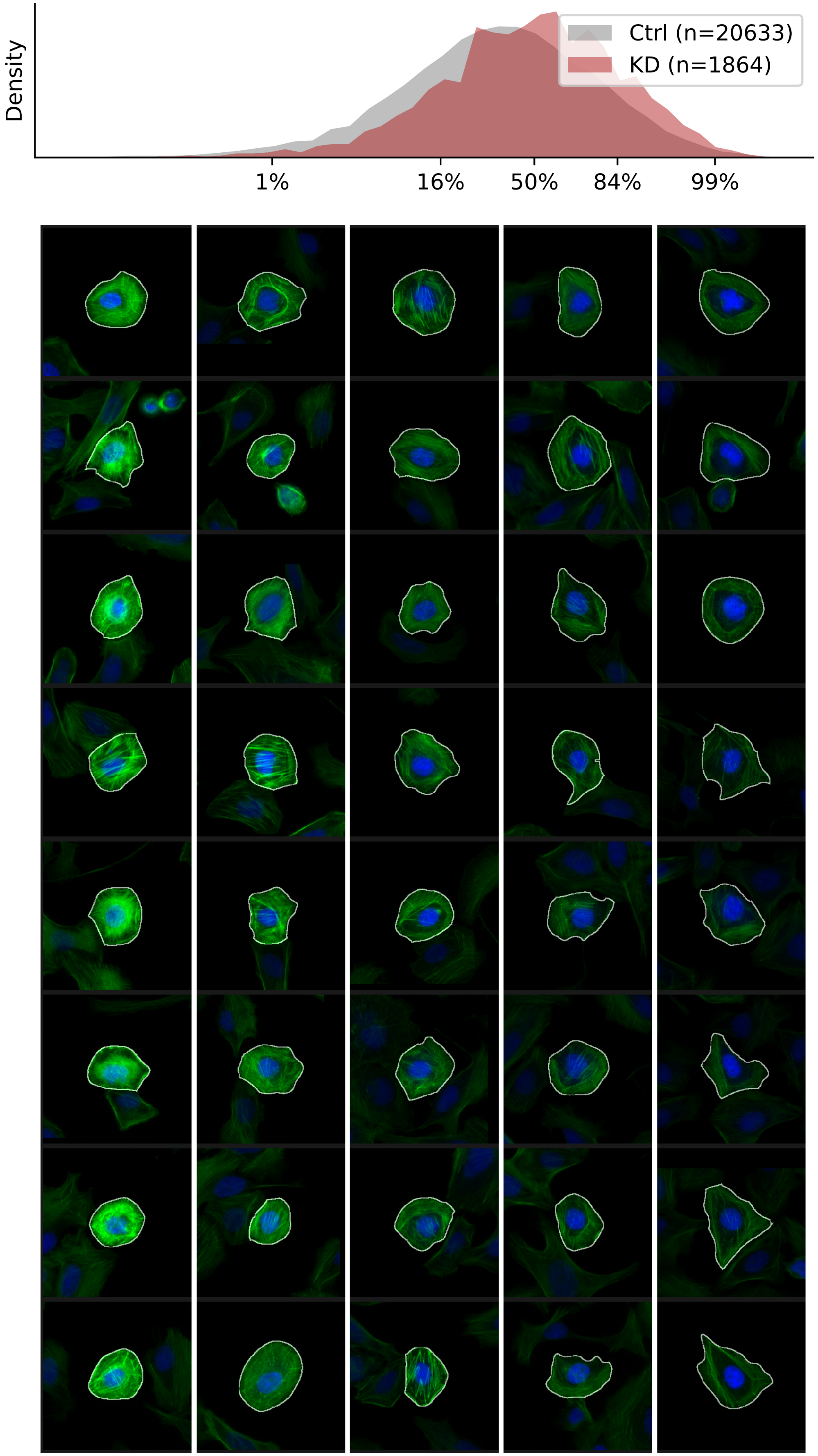

### VIEWS_iAD_MTOR_rep1.pdf

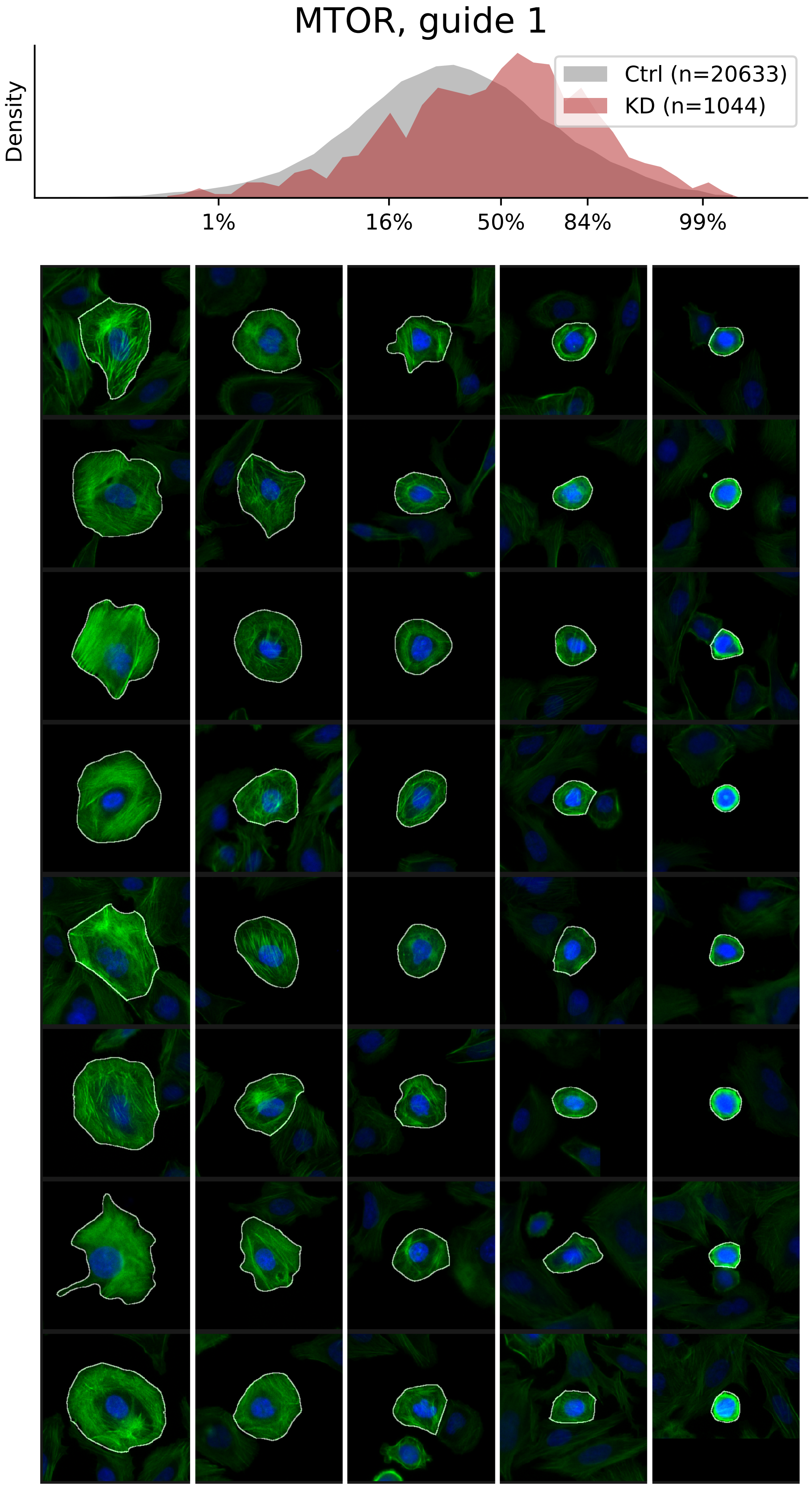

### VIEWS_iAD_MTOR_rep2.pdf

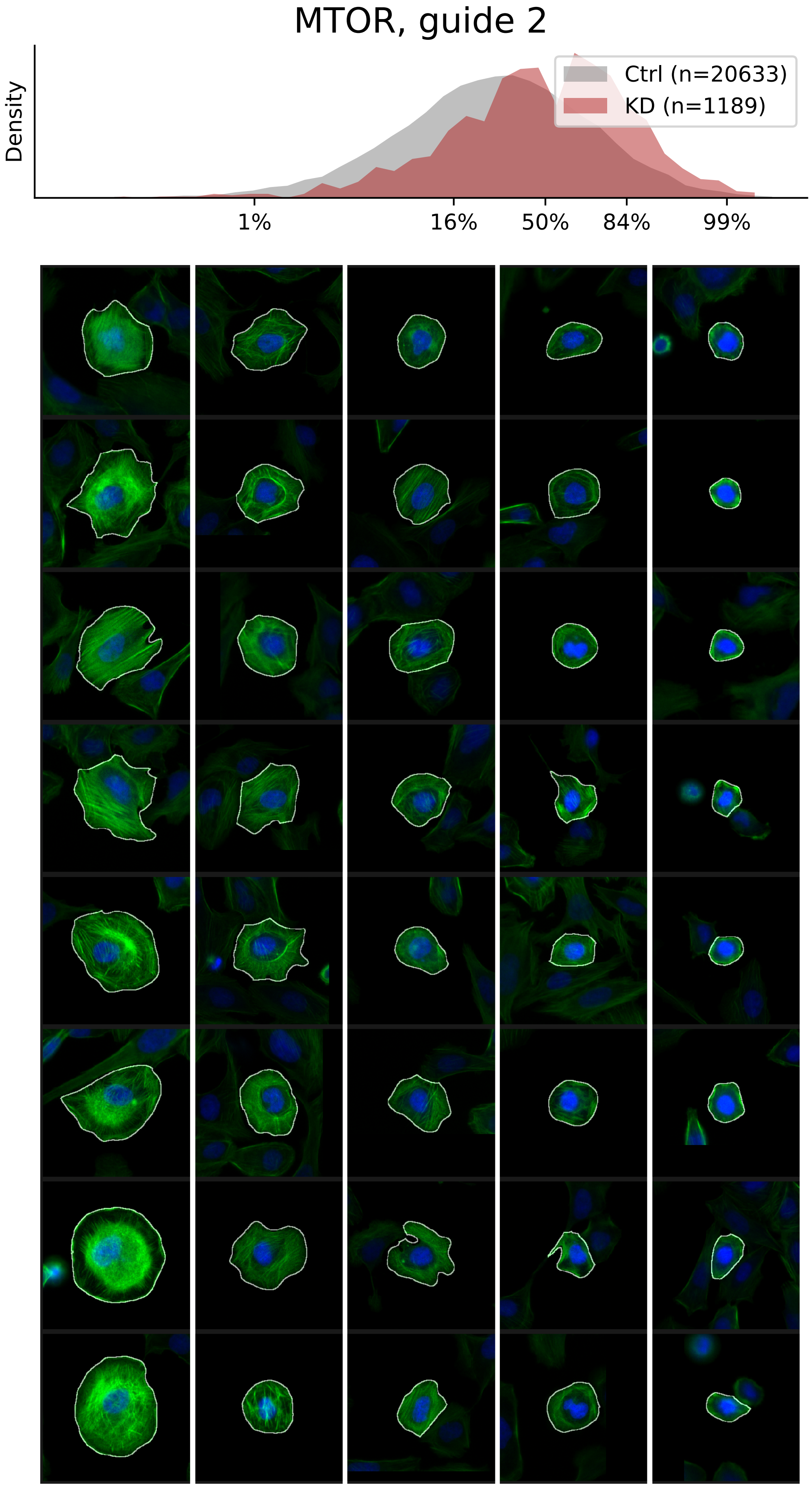

### VIEWS_iAD_MTOR_rep3.pdf

#### MTOR, guide 3

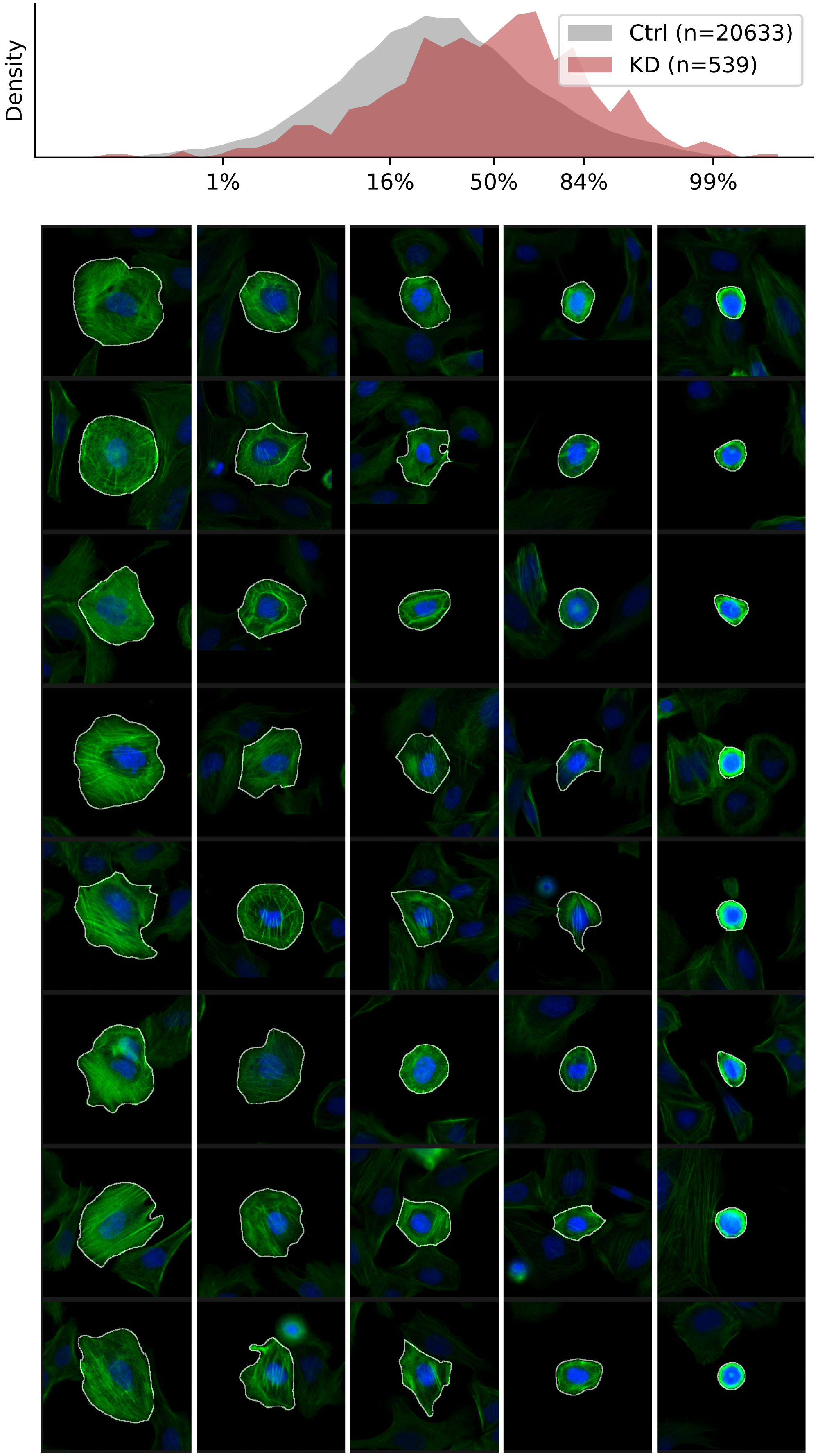

### VIEWS_iAD_MYH9_rep1.pdf

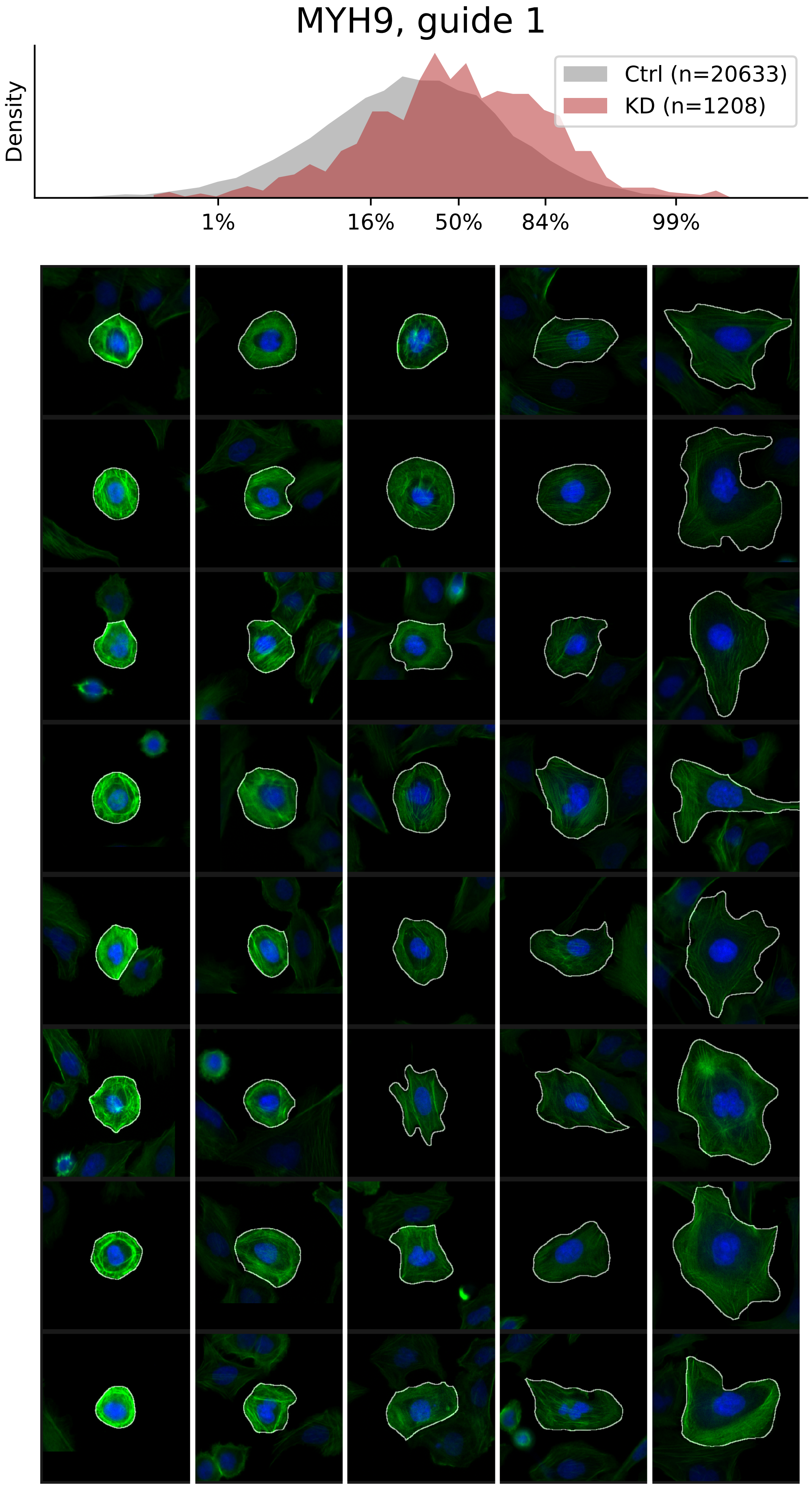

### VIEWS_iAD_MYH9_rep3.pdf

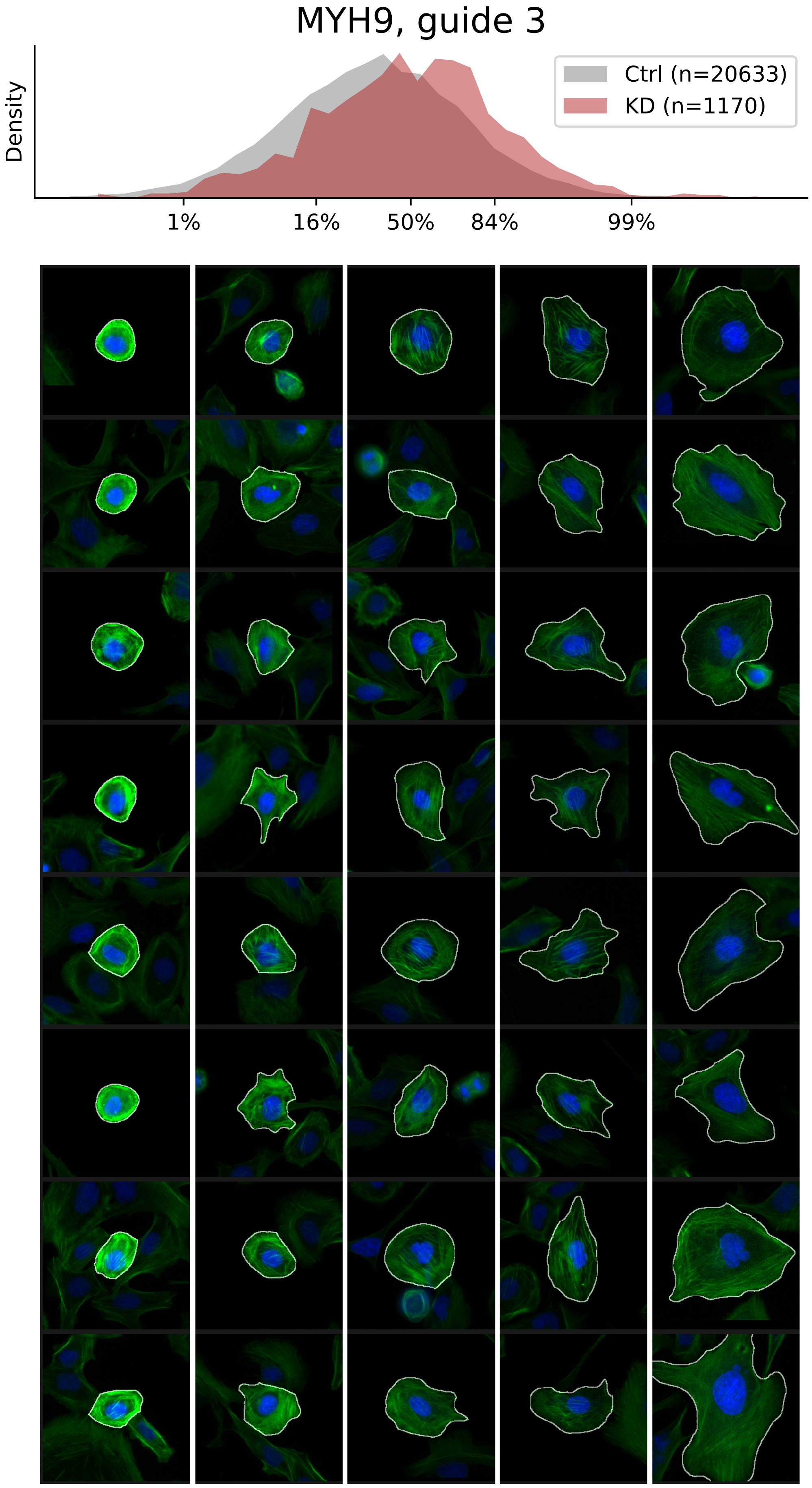

### VIEWS_iAD_PIP5K1A_rep1.pdf

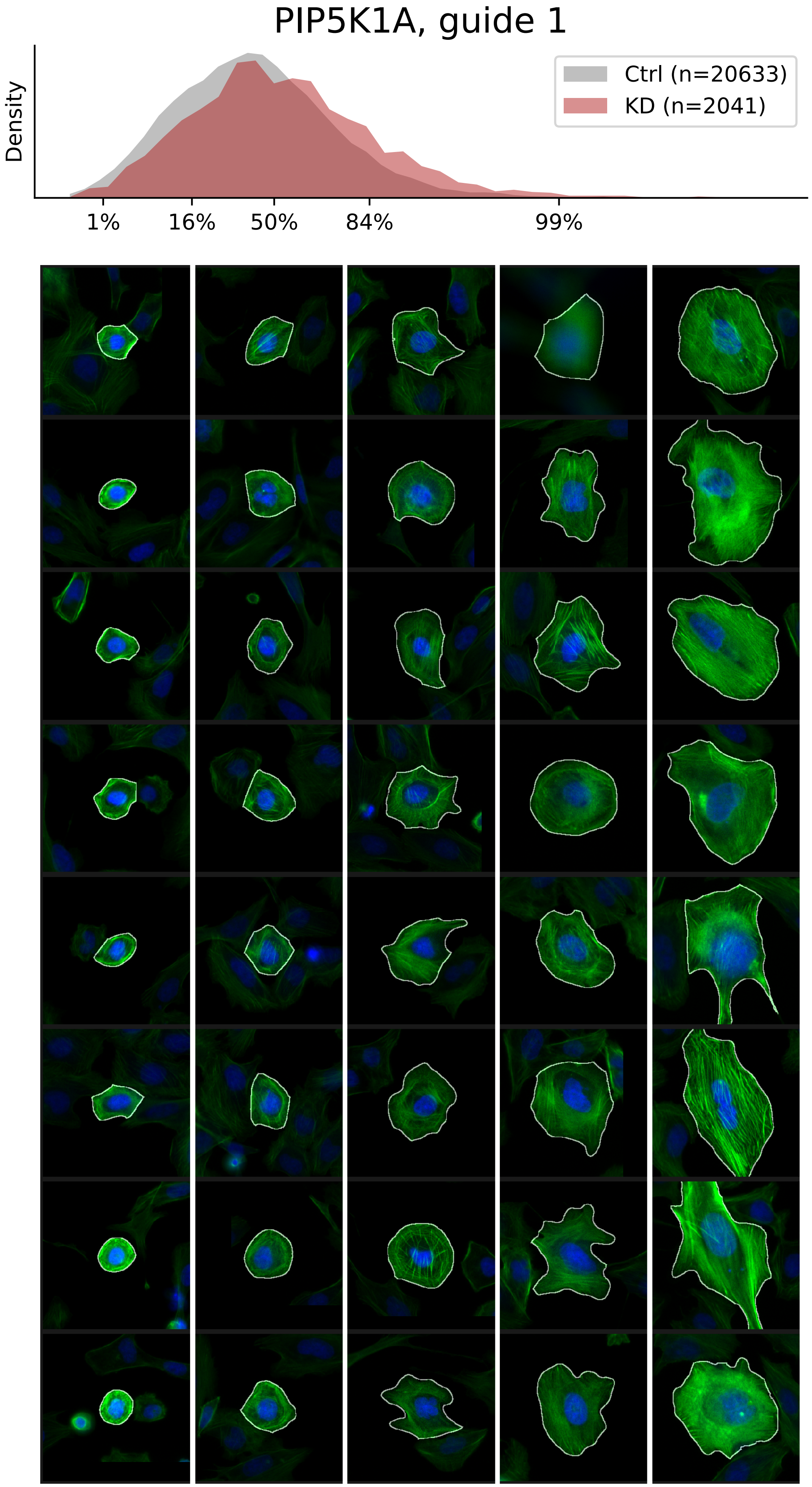

### VIEWS_iAD_PIP5K1A_rep2.pdf

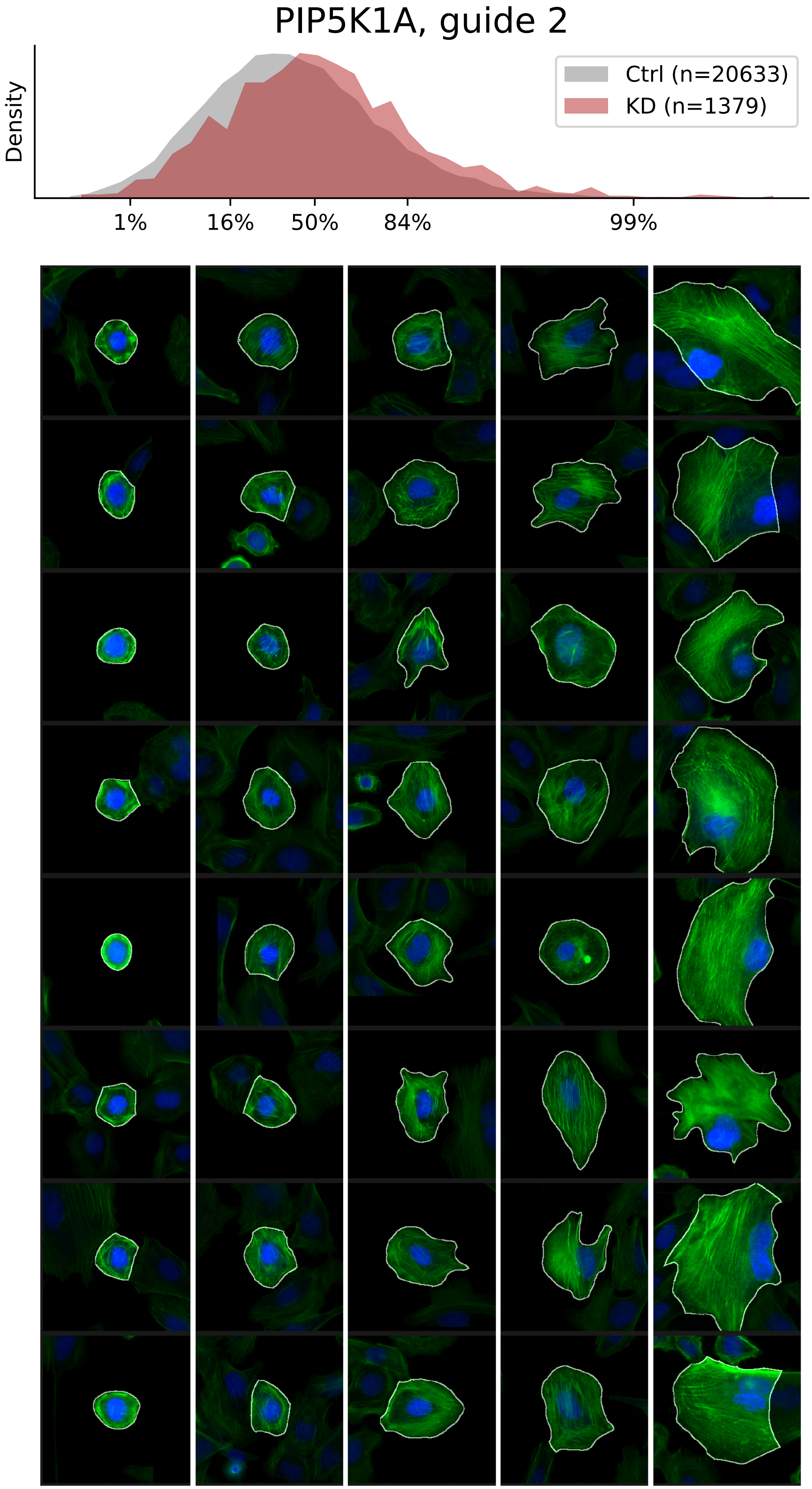

### VIEWS_iAD_PIP5K1A_rep3.pdf

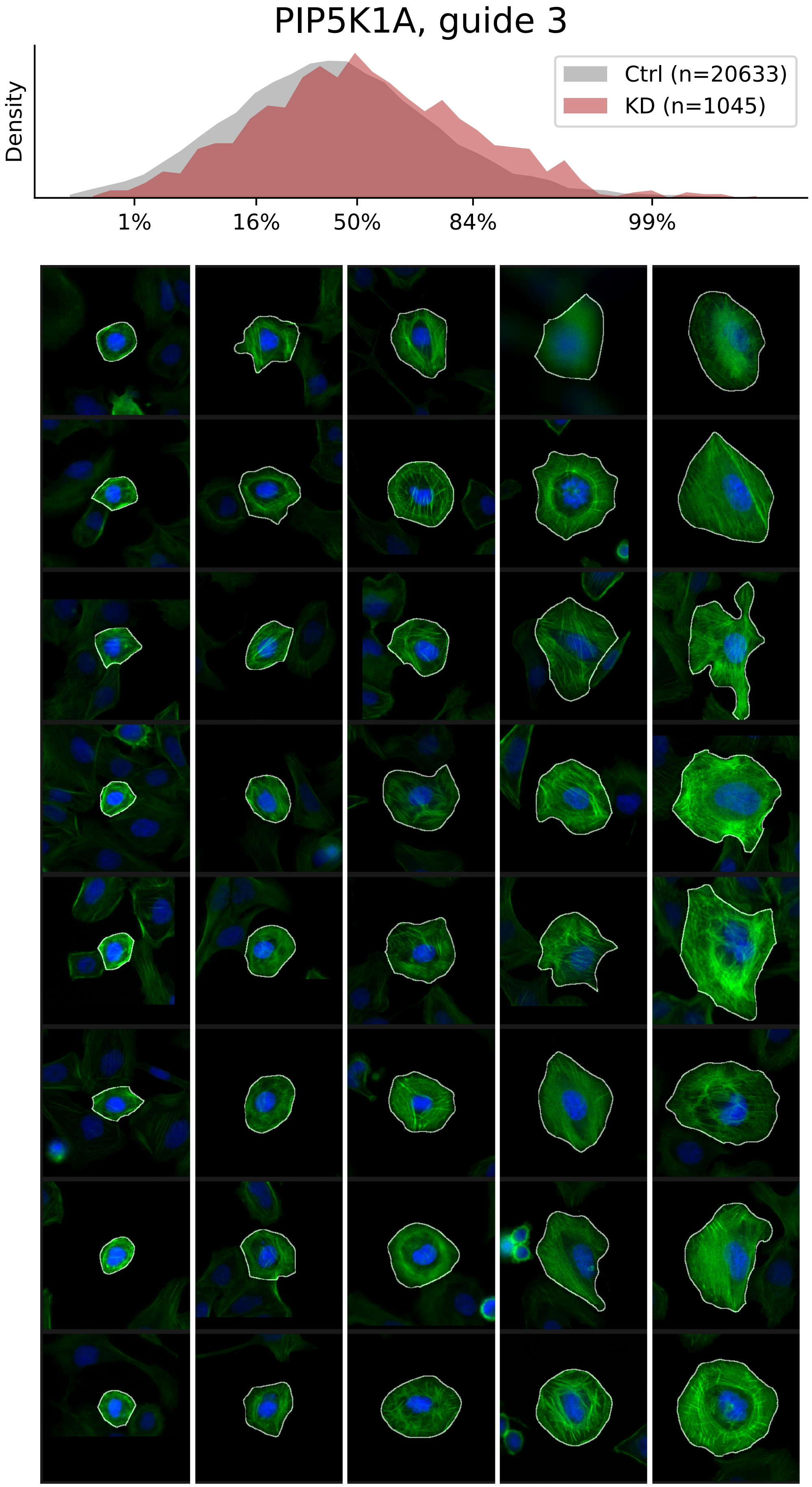

### VIEWS_iAD_TLN1_rep1.pdf

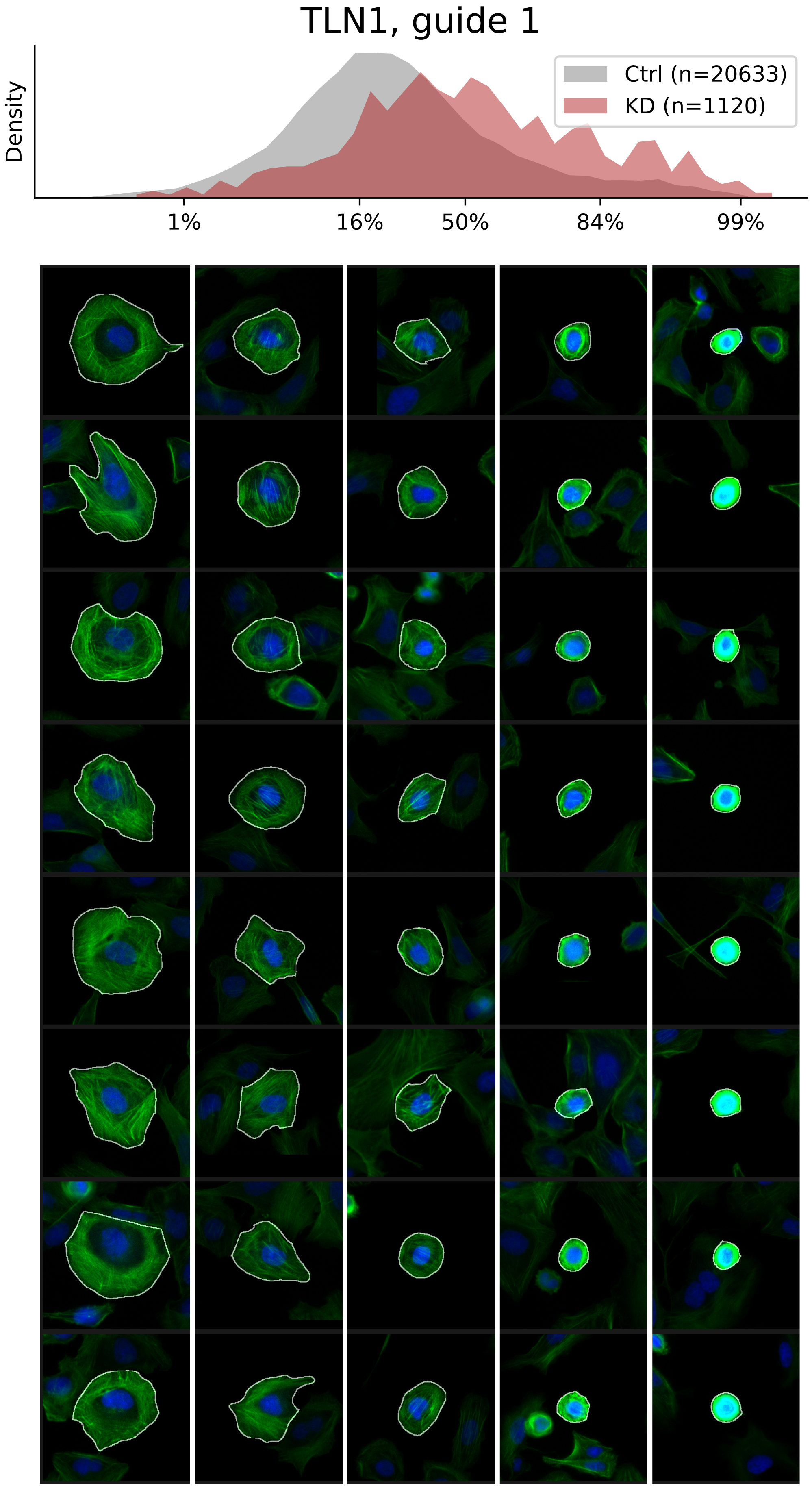

### VIEWS_iAD_TLN1_rep2.pdf

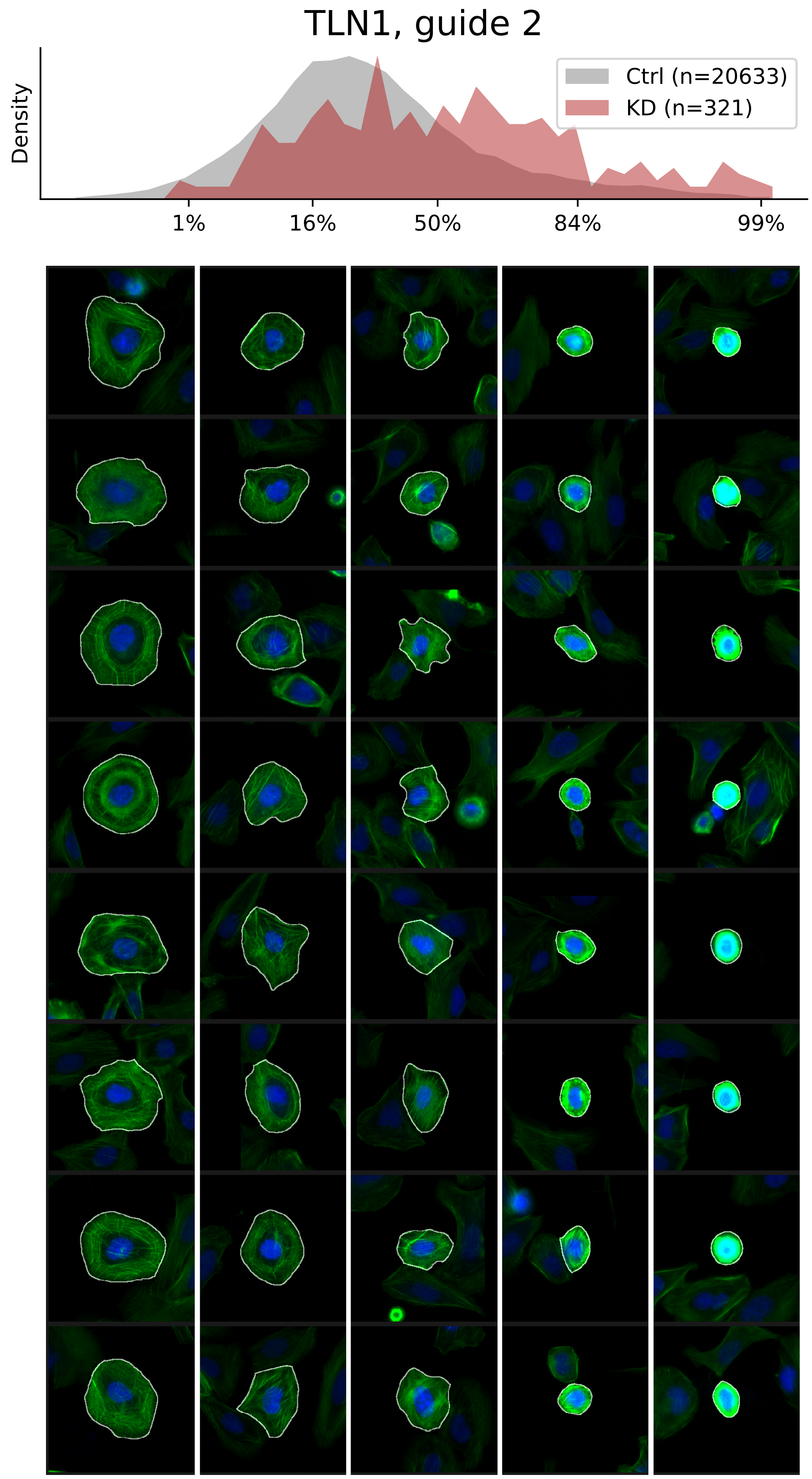

### VIEWS_iAD_TLN1_rep3.pdf

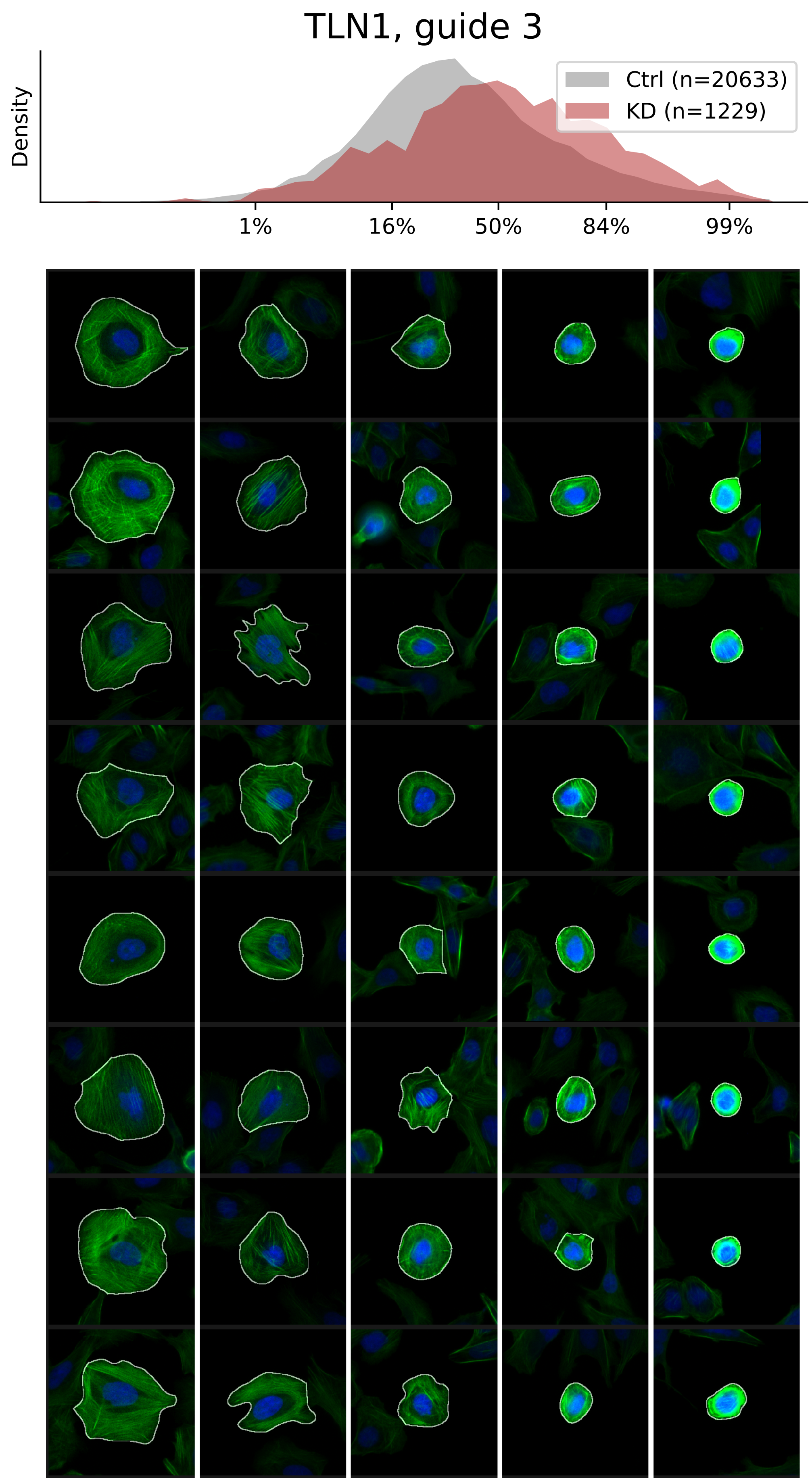

### VIEWS_iAD_TPM3_rep1.pdf

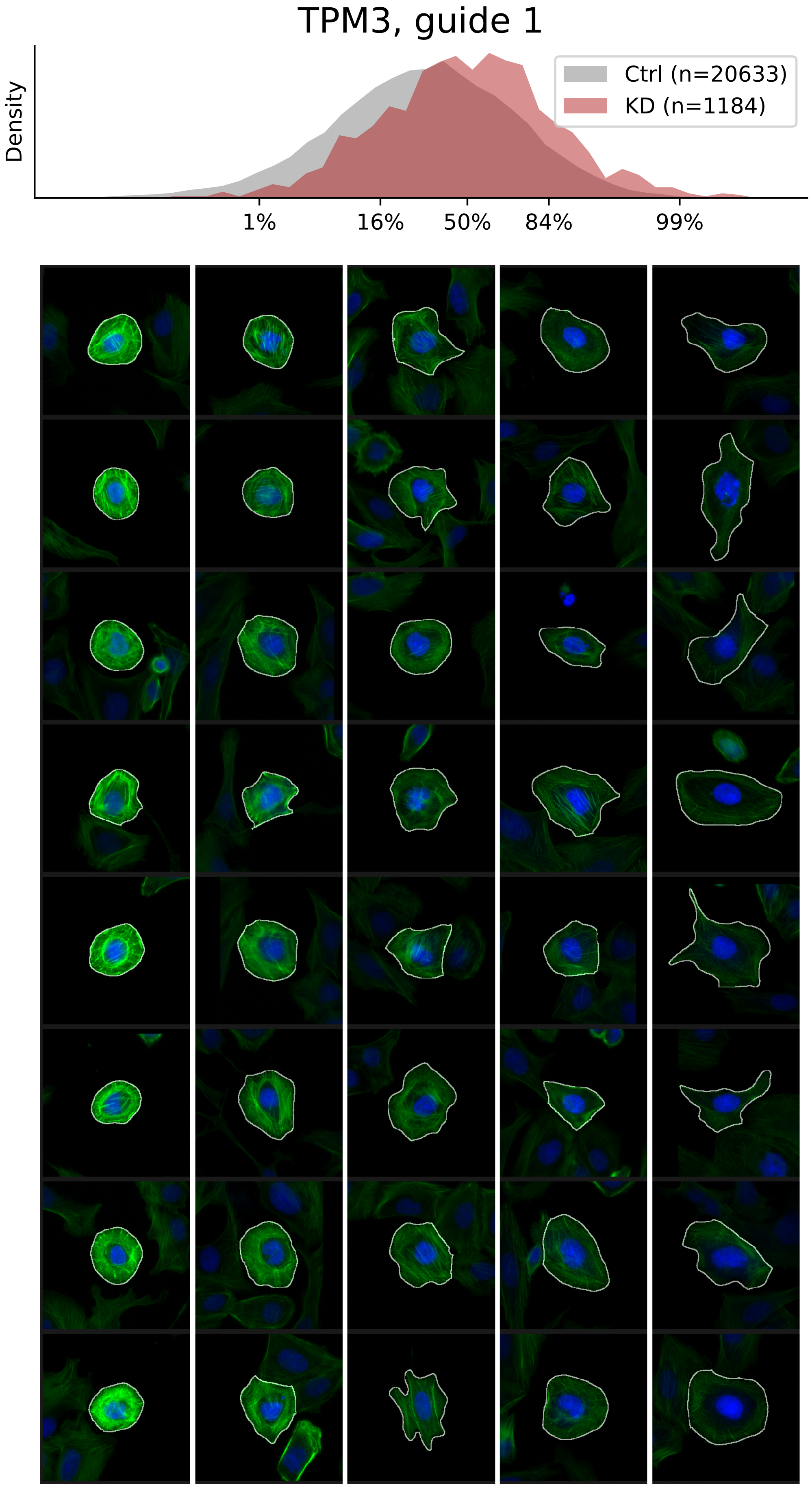

### VIEWS_iAD_TPM3_rep2.pdf

#### TPM3, guide 2

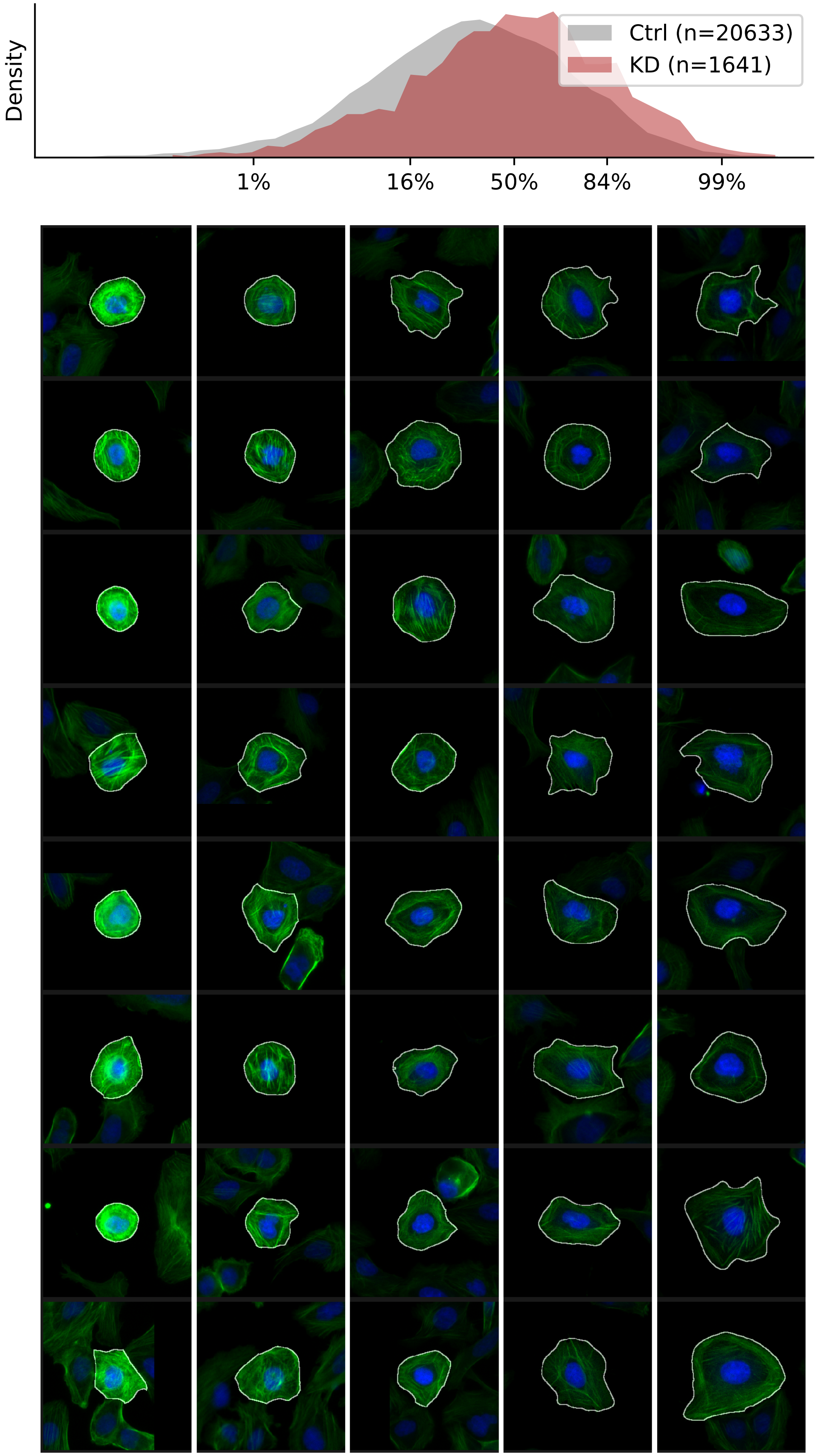

### VIEWS_iAD_TPM3_rep3.pdf

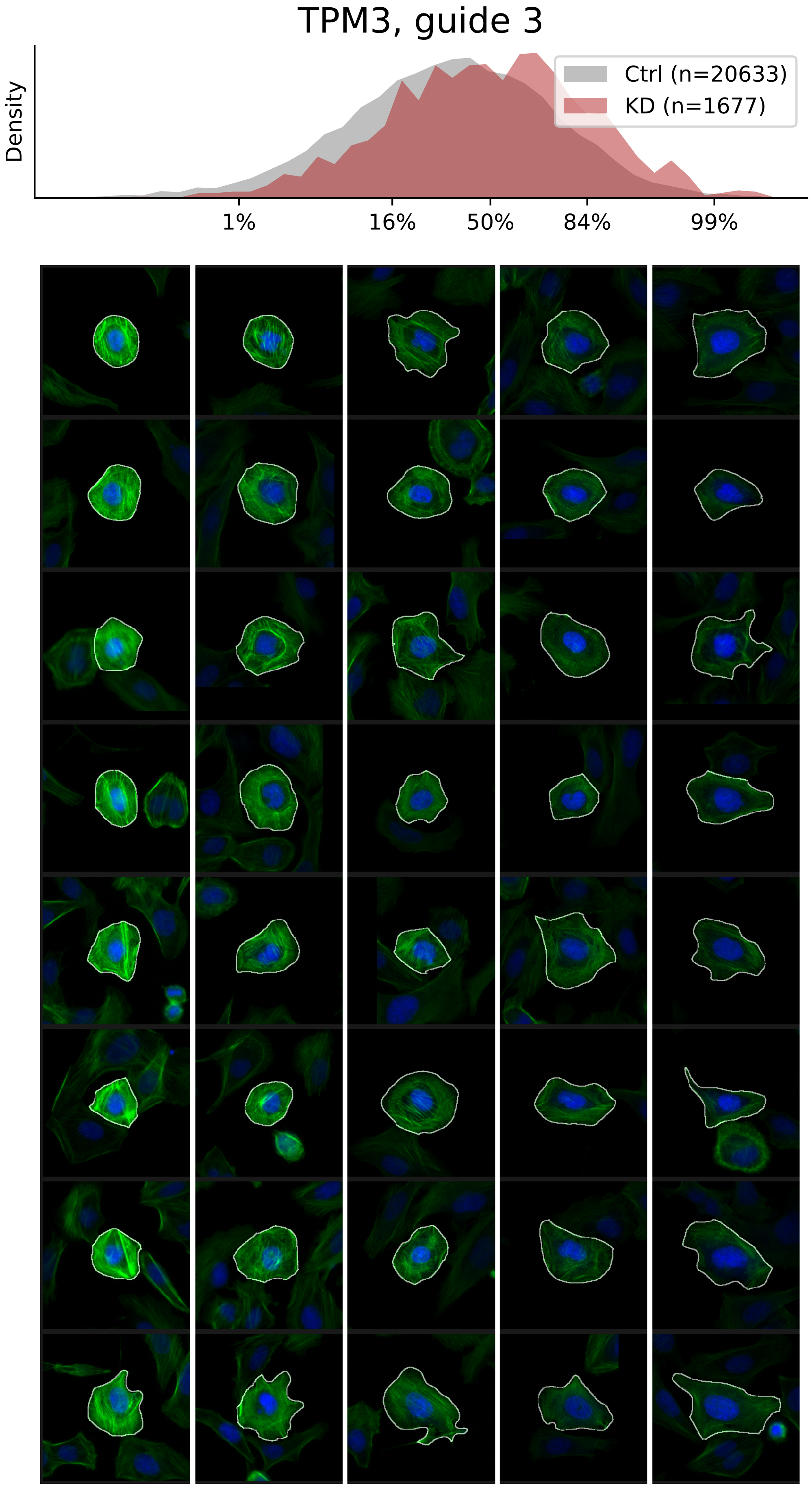
